# Modulation of the 14-3-3σ/C-RAF “auto”inhibited complex by molecular glues

**DOI:** 10.1101/2025.07.30.667769

**Authors:** Markella Konstantinidou, Holly R. Vickery, Marloes A.M. Pennings, Johanna M. Virta, Emira J. Visser, Sean D. Bannier, Mrudula Srikanth, Sabine Z. Cismoski, Lucy C. Young, Maxime C.M. van den Oetelaar, Frank McCormick, Christian Ottmann, Luc Brunsveld, Michelle R. Arkin

## Abstract

Molecular glues, compounds that bind cooperatively at protein-protein interfaces are revolutionizing chemical biology and drug discovery, allowing the modulation of traditional “undruggable” targets. Here, we focus on the native protein-protein interaction (PPI) of C-RAF, a key component of the MAPK signaling pathway, with the scaffolding protein 14-3-3. Although extensive drug discovery efforts have focused on the MAPK pathway, its central role in oncology and developmental disorders (RASopathies), still requires alternative approaches, moving beyond direct kinase inhibition. Indeed, stabilization of native PPIs is a relatively unexplored territory in this pathway. The function of C-RAF is regulated on multiple levels including dimerization, phosphorylation and complex formation with the hub protein 14-3-3. 14-3-3 prevents C-RAF activation by molecular recognition and binding at the phospho-serine 259. We used a fragment-merging approach to design a molecular glue scaffold that would bind to the composite surface of the 14-3-3/C-RAF “auto”inhibited complex. The synthesized molecular glues stabilized the 14-3-3/C-RAF complex up to 300-fold in biophysical assays; their glue-based mechanism of action was confirmed with several crystal structures of ternary complexes. Selectivity among the other RAF isoforms and other RAF phosphorylation sites was evaluated with biophysical assays. The best compounds showed excellent selectivity among a broad panel of 80 14-3-3 clients. Validation in cell assays showed on-target engagement, enhanced phosphorylation levels of the C-RAF pS259 site, reduced RAF dimerization and reduced ERK phosphorylation. Overall, this approach enables chemical biology studies on a C-RAF site that is intrinsically disordered prior to 14-3-3 binding and has not been targeted previously. These molecular glues will be useful as chemical probes and starting points for further drug discovery efforts to elucidate the effect of native PPI stabilization in the MAPK pathway with applications in oncology and RASopathies.

## INTRODUCTION

RAF kinases are central regulators of the RAS-RAF-MEK-ERK signaling pathway (MAPK), which controls multiple cellular processes, including cell proliferation, differentiation, and survival.^1^ Frequent mutations, especially on RAS and the B-RAF isoform, lead to aberrant downstream signaling and occur in different types of cancer.^2,3^ Direct inhibitors for components of the MAPK pathway have been the focus of drug discovery efforts for decades. ^4,5^

Recently, single-molecule cryo-EM has started to elucidate the structural aspects of MAPK protein-protein interactions (PPIs) as well as the underlying dynamics of pathway activation, providing new opportunities for therapeutic interventions.^6–8^ The regulation of RAF kinases is controlled on different levels^9,10^ by phosphorylation, conformation, dimerization, and binding to 14-3-3 proteins^11,12^, adaptor proteins that recognize specific phospho-serine or phospho-threonine motifs on client proteins.^13,14^ The three RAF isoforms (A-, B-, C-) have highly conserved amino acid sequences and consist of three main domains: the CR1 domain on the N-terminus with the RAS binding domain (RBD) and the cysteine-rich domain (CRD), the CR2 which includes phosphorylation sites, and the C-terminus (CR3), which includes the kinase catalytic domain (CAD).^15,16^ The 14-3-3 binding sites are located in the conserved regions (CRs) on either side of the kinase catalytic domain (Fig 1A). The two 14-3-3 binding sites have opposite functions on the regulation of the pathway; the sites located on CR2 (pS214 for A-RAF, pS365 for B-RAF, pS259 for C-RAF) inhibit RAF activation, while sites located on the CR3 domain (pS582 for A-RAF, pS729 for B-RAF, pS621 for C-RAF) are activating.^17^

**Figure 1.**
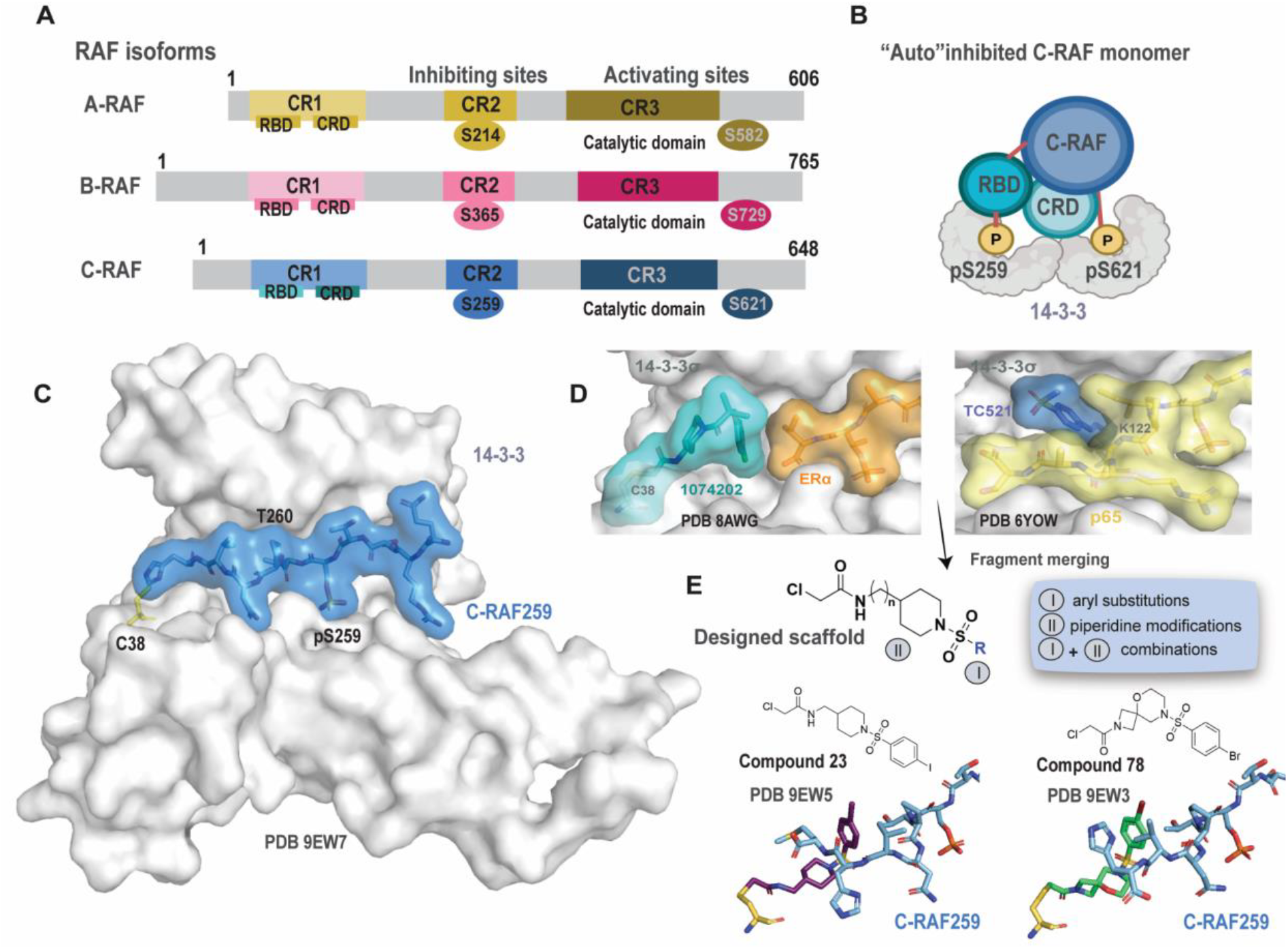
RAF isoforms, 14-3-3/C-RAF pS259 complex and fragment merging approach. A) Inhibiting and activating sites on A-, B- and C-RAF. B) Proposed model for the “auto”-inhibited C-RAF monomer complex bound to a 14-3-3 dimer. C) Crystal structure of 14-3-3σ (gray surface) bound to a phospho-C-RAF259 peptide (cyan surface). C38 (14-3-3) is shown as yellow sticks (PDB: 9EW7. D) Left: crystal structure of 1074202/14-3-3σ/ERα (PDB 8AWG). Right: crystal structure of TC521/14-3-3/p65 (PDB 6YOW). E) Top: Fragment merging approach towards a designed scaffold for the 14-3-3σ/C-RAF pS259 complex. Bottom: Representative examples and their binding mode (14-3-3 is omitted for clarity, C38 is shown as yellow sticks, and C-RAF as cyan sticks).

Although cryo-EM structures with full length C-RAF are not yet reported, C-RAF activation is hypothesized to proceed in a similar manner as B-RAF.^18^ A recent single-molecule FRET (sm-FRET) study on C-RAF supports this hypothesis.^19^ In analogy to the B-RAF cryo-EM structures, in the proposed model C-RAF is maintained in the “auto”inhibited, closed state via interactions of the N- and C-terminal domains with a 14-3-3 dimer (Fig 1B). The pS259 site (corresponding to the B-RAF pS365 site) and the pS621 site (corresponding to the B-RAF pS729 site) are each bound in an amphipathic, phosphopeptide-binding groove in the 14-3-3 dimer. Upon pathway activation, C-RAF is recruited to the membrane and interacts with RAS via the RBD domain. This is followed by the release of the CRD domain from the “auto”inhibited complex and its subsequent interaction with RAS and the membrane. This conformational change exposes the inhibitory C-RAF pS259 site to the phosphatase SHOC2-PP1 for dephosphorylation.^20,21^ RAF is then activated through dimerization of the catalytic domains, stabilized through binding to 14-3-3 via the C-terminal phosphorylation sites only (pS621 for C-RAF and pS729 for B-RAF). Homodimers and heterodimers between all three RAF isoforms are reported.^22,23^

Despite the sequence similarity in RAF isoforms, distinct roles are starting to emerge. Recent reports show that C-RAF depletion inhibits tumor growth but not ERK signaling in K-RAS mutant cells,^23^ whereas pan-RAF inhibition affects PI3K-signaling in multiple myeloma.^24^ The roles of hetero-dimerization is also an active area of research. The current thinking is that the B-RAF/C-RAF heterodimer is the primary species involved both in native and oncogenic signaling.^25,26^ Notably, the dimerization of C-RAF with A-RAF is also thought to be involved in K-RAS driven tumor growth^23^ and RAF heterodimers are an important mechanism linked to RAF inhibitor resistance (including ‘paradoxical activation’) in cancer therapy.^27,28^

Here, our aim is to disrupt the transition from the closed C-RAF monomer to the open, active dimer by stabilizing the “auto”inhibited C-RAF pS259/14-3-3 complex (Fig 1C) with molecular glues (MGs)^29^, as an alternative approach to direct C-RAF kinase inhibition. Although direct, selective inhibition of C-RAF has not yet been achieved clinically, a recent study utilizing bio-orthogonal ligand tethering showed that selective C-RAF inhibition promotes paradoxical activation, thus potentially leading to resistance mechanisms.^26^ To date, reported stabilizers of the “auto”inhibited C-RAF pS259/14-3-3 complex are the natural product cotylenin-A^30^ (CN-A, Fig S1) and disulfide-tethered fragments identified by our lab^31^. These disulfide fragments show two distinct binding conformations in crystal structures, although neither appear to make specific contacts with the residues C-terminal to C-RAF259^31^ (Fig S1).

Here, we used a fragment merging approach to design molecular glues for the unique composite surface of the 14-3-3/C-RAF259 complex, aiming at specific ligand – protein interactions, including the formation of a covalent bond with C38 (on 14-3-3σ) and polar interactions with T260 (+1 residue on C-RAF259 phosphorylation site). The most potent stabilizer, compound **23** resulted in 280-fold stabilization of the 14-3-3σ/C-RAF259 complex in fluorescence anisotropy assays, had an EC_50_ of 0.18 μM in a cellular 14-3-3/C-RAF NanoBRET assay and partially blocked N-RAS/C-RAF dimer formation in cells. Compound **78**, although less potent than **23**, showed higher selectivity for C-RAF over the other RAF isoforms in cell assays. Thus, a fragment-based development of MGs provided a tunable selectivity that will help elucidate specific roles of RAF complexes in normal and oncogenic signal transduction.

## RESULTS AND DISCUSSION

### SAR development and biophysical assays

Our scaffold design strategy combined a chloroacetamide piperidine moiety (inspired by analog 1074202, cysteine-targeting 14-3-3/ERα stabilizer^32^) with fragments bearing a sulfonyl group (inspired by TC-521, a lysine-targeting 14-3-3/p65 stabilizer^33^) (Fig 1D). We hypothesized that the sulfonyl group in the new scaffold would serve to induce a pronounced conformational bend in the MG scaffold and act as a flexible handle for potential polar interactions with T260. Chemical modifications were then performed to optimize the designed scaffold (Fig 1E). Modification (I), in close proximity to the peptide, included optimizing the linkage of the two merged fragments and substitutions on the aryl ring. Modification (II) focused on replacing the piperidine ring with spiro-cycles and fused ring systems with varying sizes and orientations. Lastly, combinations of modifications (I) and (II) aimed at exploring synergistic effects.

In analogy to our previous work, two orthogonal assays were used for screening^32^ (Fig S2). Briefly, the mass spectrometry (MS) assay monitored the formation of the covalent bond between electrophiles and the native cysteine (C38) on 14-3-3σ. Dose responses of compounds were performed in the absence and presence of the C-RAF259 phospho-peptide to distinguish between neutral binders and cooperative molecular glues. The experiment was performed as a time-course, with measurements every 8 hours, after an 1h incubation. Additionally, the compounds were tested in a fluorescence anisotropy (FA) assay in the presence of 5-carboxyfluorescein-(FAM)-labeled C-RAF259 peptide. For effective MGs, a significant increase in anisotropy was observed in the overnight measurement. The full dose-response data for all compounds are available in the SI (tables S2-S5, Fig S5-S8, Fig S15-S18, Fig S25-S32, Fig S37-S39). The dose-response data from both assays were also visually compared in the form of bar graphs. For the MS data, the bar graphs represented the % bound at 1 μΜ compound concentration and 100 nM 14-3-3σ (10:1 ratio) in the absence or presence of peptide. For FA experiments, the EC_50_ values from the overnight measurement were calculated and were plotted as bar graphs showing the positive log EC_50_ value (pEC_50_). Compounds acting as MGs were expected to show an increase in both types of bar graphs.

The first key modification on the newly designed scaffold focused on the appropriate linkage of the two merged fragments. Chemically, benzyl or aryl sulfonyl chlorides could be reacted with the appropriate piperidine moiety. Initial screening data from both MS and FA assays showed that benzyl analogs were inactive (compounds **6-9**, Fig S3, S5-S7), whereas for phenyl analogs a SAR began to emerge as substituents were added on the phenyl ring (Fig S4, S5-S7). Various substituents in *o*-/*m*-/*p*-positions were added, including electron-withdrawing and electron-donating groups. As a general trend, electron-donating groups were not tolerated, whereas halogens in the *p*-position significantly improved potency, stabilization, and cooperativity (Fig 2A-2B, Fig S4-S7, tables S2-S3). In the MS assay, analogs **11** (*p*-F) and **12** (*p*-Cl) showed binding in the presence of the C-RAF259 peptide and only very low binding in the apo screen (no peptide), indicating cooperative binding. In FA compound titrations, **11** and **12** showed low micromolar EC_50_ values (17 ± 3 and 3 ± 1 μM, respectively) (table S3, Fig S7). The two compounds were further validated as 14-3-3σ/C-RAF259 molecular glues in FA protein titrations. 14-3-3σ was titrated into 10 nM FAM-labeled C-RAF259 peptide in the presence of DMSO or saturating concentration of the compounds (100 μΜ). Compounds **11** and **12** decreased the dissociation constant for the 14-3-3s/C-RAF259 peptide complex by 4-fold and 22-fold, respectively (Fig S8, table S3). Increasing the linker length of the warhead or replacement of the phenyl ring with a pyridine ring led to loss of potency and these modifications were not investigated further (compounds **13**-**14**, Fig S4-S7).

**Figure 2.**
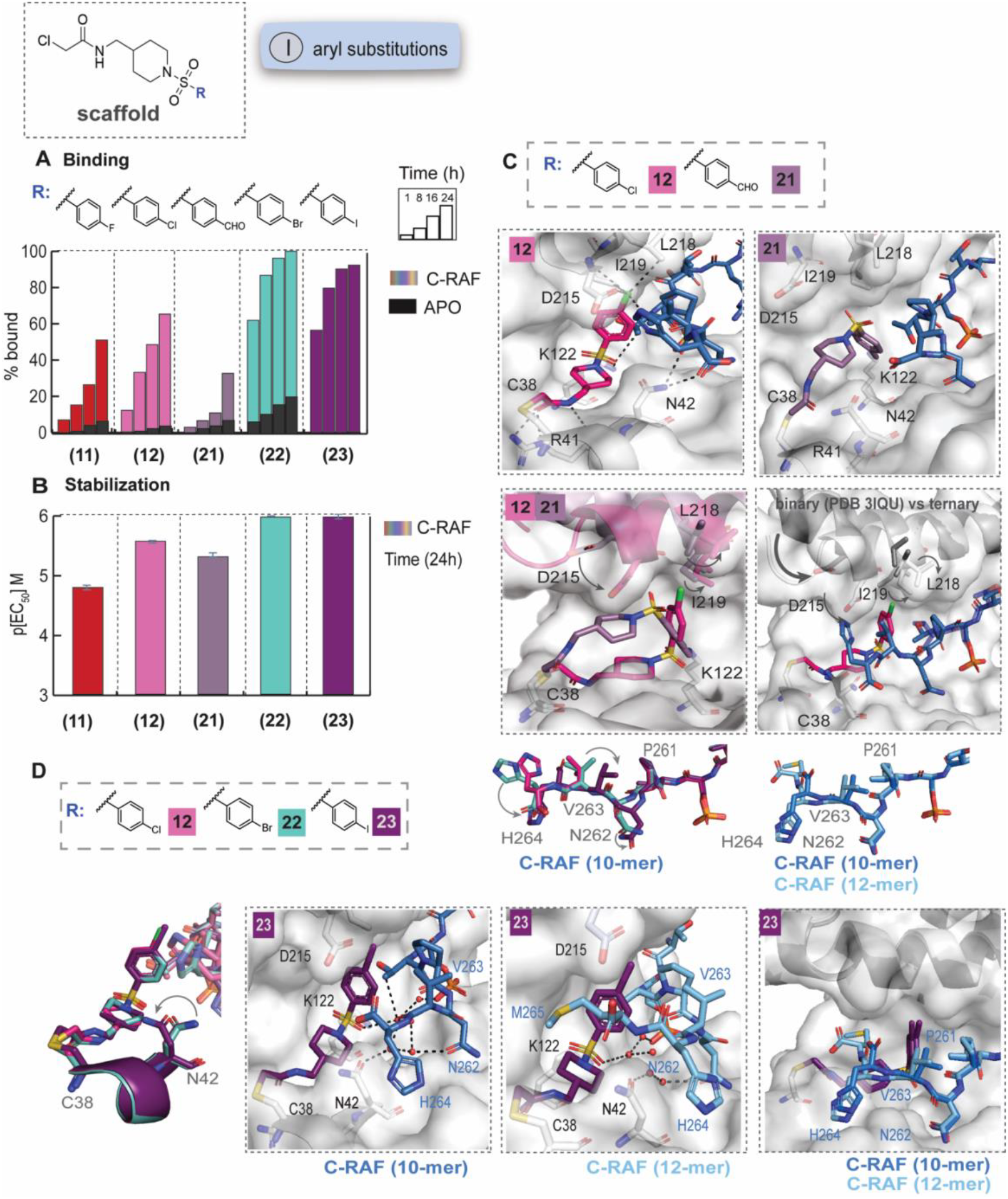
SAR and crystal structures of selected phenyl analogs. A) MS bar graphs at 1 μΜ. For each compound, time course experiments were performed with measurements at 1h, 8h, 16h and 24h. C-RAF259 data are shown with different colors for each compound, and apo data in black. B) Bar graphs of FA compound titration pEC_50_ values after overnight incubation. C-RAF259 data are shown with different colors for each compound. C) Top: Crystal structures of **12** (pink sticks) and **21** (light purple sticks) with 14-3-3σ (white surface) and C-RAF pS259 10-mer peptide (blue sticks). Bottom: overlays of **12** with **21** (left), and **12** with a binary complex (PDB 3IQU, right). D) Left: crystal structures of **12** (pink sticks)/**22** (turquoise sticks)/**23** (purple sticks) with 14-3-3σ and C-RAF259 (overlay). Middle: detailed interactions of **23**/14-3-3σ/C-RAF259 10-mer (left) and 12-mer (right). Interacting water molecules are shown as red spheres. Left: overlay of the structures.

To test whether the observed stabilization was unique to halogen substituents or other electron donating groups could be tolerated, the *p*-formyl analog **21** was synthesized. The two covalent warheads of this analog could potentially interact simultaneously with C38 and K122 on 14-3-3σ. The analog was weak in the MS assay and despite the low EC_50_ value (5 ± 1.0 μM) in the FA compound titrations (Fig 2A, 2B, Fig S4-S7), it resulted in only 9-fold stabilization of the 14-3-3s/C-RAF259 complex in the FA protein titrations (Fig S8, table S3). Based on these observations, the presence of halogens seemed to be crucial, and thus bulkier halogens were introduced (*p*-Br **(22)** and *p*-I **(23)**). Both compounds showed faster binding in the MS assay (Fig 2A), and remarkably the *p*-I analog showed almost no binding to 14-3-3a alone; thus, **23** binding was highly cooperative for the 14-3-3σ/C-RAF259 complex. In the FA assay both compounds showed low micromolar EC_50_’s (1 ± 0.5 μΜ). Consistent with high cooperativity, in protein titrations, the *app*K^d^ of the 14-3-3σ/C-RAF259 decreased from 8427 nM to 47 nM in the presence of **22** and to 30 nM in the presence of **23**. Thus, **22** and **23** stabilized the 14-3-3σ/C-RAF 259 complex by 179- and 280-fold, respectively (table S3, Fig S8).

Crystal structures of the ternary complexes with 14-3-3σ and a 10-mer C-RAF259 phospho-peptide were solved for **12, 21, 22** and **23** (Fig 2C, D, Fig S9-S10). Overall, **12, 22**, and **23** form compact, L-shaped molecules that form multiple interactions with both the C-RAF peptide and 14-3-3.

Taking **12** as an example, in addition to the covalent bond with C38 of 14-3-3, two hydrogen bonds were formed between the warhead amide, R41 of 14-3-3a (2.7Å), and the backbone carbonyl of C38 (3.3Å). N42 of 14-3-3 directly interacted with the backbone H264 of C-RAF via two hydrogen bonds (3.0Å). The sulfonyl group of **12** interacted with K122 of 14-3-3 via a charged-assisted hydrogen bond (3.2Å) and with T260 (+1 position) of C-RAF via a hydrogen bond (3.4Å).

Based on our past designs^32^ and the conformation of the aldehyde fragment TC521 (Fig 1D), we anticipated that the halogenated/formylated aryl ring would point these substitutions down into a pocket that includes K122 on 14-3-3σ. Instead, **12** (*p*-Cl) adopted a surprising upward conformation at the 14-3-3/peptide interface (Fig 2C, Fig S9) with the halogen pointing into a shallow pocket towards the top of the 14-3-3 binding groove, potentially allowing the formation of halogen bonds with D215, L218, I219 of 14-3-3 or V263 of C-RAF to increase the stabilization of the complex. In stark contrast, **21** (*p*-CHO) showed a different binding mode, interacting with C38 and K122 via the two covalent warheads. While this conformation more closely mirrored our prediction, no additional interactions were observed with 14-3-3 or C-RAF (Fig 2C). An overlay of the two crystal structures showed a significant conformational change of helix 9 of 14-3-3 for **12** (Fig 2C). The helix moved to a downward position, closer to the compound, thus “clamping” the compound in the 14-3-3 binding grove and causing conformational changes to D215, L218 and I219. An overlay with a previously published binary structure further confirmed that the movement of the helix was compound-induced (Fig 2C).

Crystal structures of **22** (*p*-Br) and **23** (*p*-I) showed an upward binding mode, consistent with **12** (Fig 2D, Fig S10). The key ligand-protein and ligand-peptide interactions were maintained. Different water-networks were formed for **22** and **23**. For **22**, N42 formed a water-network that interacted with D215 of 14-3-3. For **23**, a more extensive water-network was observed, which facilitated additional interactions between the compound, N42 (14-3-3) and the C-RAF residues and extended to the phospho-S259 residue. Interestingly, an overlay of the three structures showed that the C-RAF residues (N262, V263, H264) also adopted different conformations with bigger changes in the case of **23** (Fig 2D, Fig S10).

Notably, the C-RAF sequence is internal and extends well beyond H264. Although shorter peptides are usually favored for crystallography, we hypothesized that longer peptides might be a better representation of the binding mode for full-length C-RAF. To this end, we solved a binary complex of 14-3-3σ with a 12-mer C-RAF peptide and an additional ternary complex for **23** (Fig 2D, Fig S11). The presence of the compound significantly improved the density observed for the C-RAF sequence and allowed the visualization of M265, which “wrapped” around the main core of the compound, thus significantly covering the binding groove. An overlay of the two ternary complexes for **23** with the 10mer and 12-mer peptides showed similar conformations for N262, V263, and H264; the latter was oriented toward the outer side of the binding grove in both structures. Consistently, helix 9 adopted a downward conformation in both structures with **23** (Fig 2D).

Structure-guided optimization led to the synthesis of double- and triple-halogenated derivatives to investigate additional electronic and steric effects at the protein-peptide interface (Fig 3A,3B, Fig S12, Fig S5-7). While keeping the electron-withdrawing groups in the *p*-position, the introduction of additional *o*-F substituents was less favorable than *m*-F substituents. In *m*-position bulkier substituents than the F-group significantly decreased both the binding in the MS assay and the stabilization in the FA assay, indicating steric hindrance (tables S2-S3). Analogs **29** (*m*-F, *p*-Cl), **32** (*m*-F, *p*-CF_3_) and **37** (*m*-F, *p*-Br) were representative examples of the observed SAR (Fig 3A,3B, Fig S12). Consistent with previous observations, analog **37** bearing a *p*-Br group showed fast and cooperative binding in the MS assay. In the FA compound titrations analogs **29, 32** and **37** showed similar EC_50_ values (between 1-5 μΜ). In protein titrations, **37** was slightly more potent than **29** and **32** (*app*K_d_ 92 nM compared to 100 nM and 140 nM, respectively). Triple-substituting analogs were also tested, however in protein titrations, led to lower stabilization compared to **37** (table S3, Fig S8).

**Figure 3.**
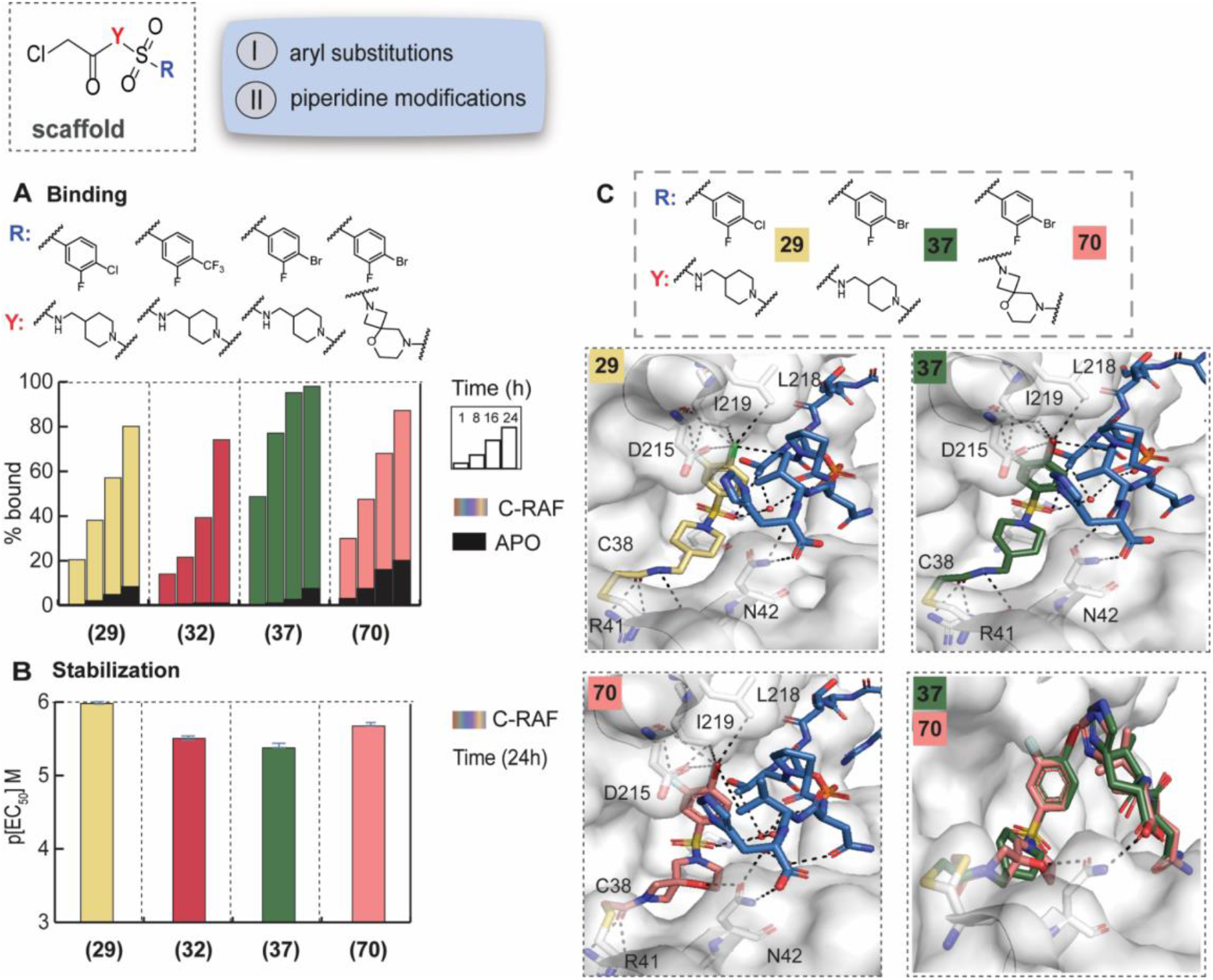
SAR and crystal structures of selected phenyl and spiro analogs. A) MS bar graphs at 1 μΜ. For each compound, time course experiments were performed with measurements at 1h, 8h, 16h and 24h. C-RAF259 data are shown with different colors for each compound, and apo data in black. B) Bar graphs of FA compound titration pEC_50_ values after overnight incubation. C-RAF259 data are shown with different colors for each compound. C) Top: Crystal structures of **29** (yellow sticks) and **37** (green sticks) with 14-3-3σ (white surface) and C-RAF pS259 10-mer peptide (blue sticks). Bottom: crystal structure of **70** (salmon sticks) with 14-3-3σ/C-RAF259 10-mer and overlay with **37**. Interacting water molecules are shown as red spheres.

Crystal structures with 14-3-3σ and 10-mer C-RAF259 were solved for **29, 32** and **37** (Fig 3C, Fig S13). In all cases, the compounds showed similar conformations with helix 9 adopting the downward conformation, as previously observed. The additional *m*-F group was positioned inwards and interacted with I219 of 14-3-3 (3.2Å). This orientation supported the observed SAR, where larger substituents in *m-*position were not tolerated. In *p*-position, the bulkier *p*-Br group formed the most favorable interactions, which correlated with faster kinetics and slightly improved stabilization in the biophysical assays. Although other electron-withdrawing groups in *p*-position could potentially interact with the same amino acid residues, differences in the size of the halogens (steric effects), as well as differences in electrostatic effects and σ holes^34^ had a direct effect on the biophysical assays.

Thus, the (*m*-F, *p*-Br)-substitution pattern of **37** was kept constant in the next modification, which focused on the piperidine ring (Fig S14-S18). Substituted piperidines, spirocycles and fused ring systems were investigated. Most of these changes had a negative impact on the potency and in certain cases, on the cooperativity, due to increased apo binding in the absence of C-RAF. One notable exception was analog **70**, with a spiro-ring bearing an oxygen atom. Although in the MS assay, it showed less binding than the linear analog **37**, in the FA assay it showed promising stabilization (EC_50_ = 2 μΜ, *app*K_d_ 58 nM, 95-fold stabilization) (Fig 3A,3B, Fig S14-S18). An overlay of the crystal structures of **37** and **70**, showed that the spiro-ring was slightly shifted, compared to the piperidine ring, and formed a hydrogen bond with N42 (14-3-3) at 3.0Å. An extended water-network was observed, similar to analog **23**, reaching the phospho-S259 residue (Fig 3C, S19).

Regarding the warhead position, two variations of the chloroacetamide electrophile were tested (structures **74** and **75** in table S1). Both analogs were completely inactive in both assays, indicating potentially steric hindrance and unfavorable geometry that prevented access to C38. Warhead modifications were not pursued further, instead we focused our attention on potential synergistic effects, by combining favorable aryl ring substituents with spiro-cycles (Fig 4A,4B, Fig S20, Fig S15-S18). Analogs **78** and **79**, bearing the same oxo-spiro ring as **70** and *p*-Br and *p*-I substituents respectively, showed less binding in the MS assay, which correlated with their poor solubility in the assay conditions. In the FA assay, the two analogs showed potent stabilization with EC_50_ = 2 μΜ in compound titrations and *app*K_d_ 35-40 nM (fold-stabilization 210-240, table S3, Fig S15-S18). To determine whether the aryl ring or the spiro ring were more important for binding and stabilization we synthesized analogs **80** and **83**. Replacing the favorable halogen bonds with a *p*-CF_3_ group in analog **80**, while maintaining the spiro-ring dramatically reduced the potency, indicating that the interactions between the aryl ring and V263 (C-RAF) were more significant than the hydrogen bond between the spiro ring and N42 (14-3-3). Similarly, analog **83**, with the same spiro-ring lacking the oxygen, showed slightly mer C-RAF peptide, which was more challenging. We obtained crystals with **78** and **86**, and in contrast to their binding mode with the shorter peptide, in this case 14-3-3 helix 9 was in the upward conformation. Additionally, less density was visible for C-RAF residues with the longer peptide (V263 instead of H264 for the shorter peptide), hinting potentially to unfavorable steric effects from the spiro-cycle. The overall observations imply that the movement of helix 9 was not entirely compound-induced, but it could also be affected by the length of the peptide used for crystallography. Thus, the significance of some of these interactions needs to be interpreted with caution, since the positioning of helix 9 affects the formation of compound-protein interactions.

**Figure 4.**
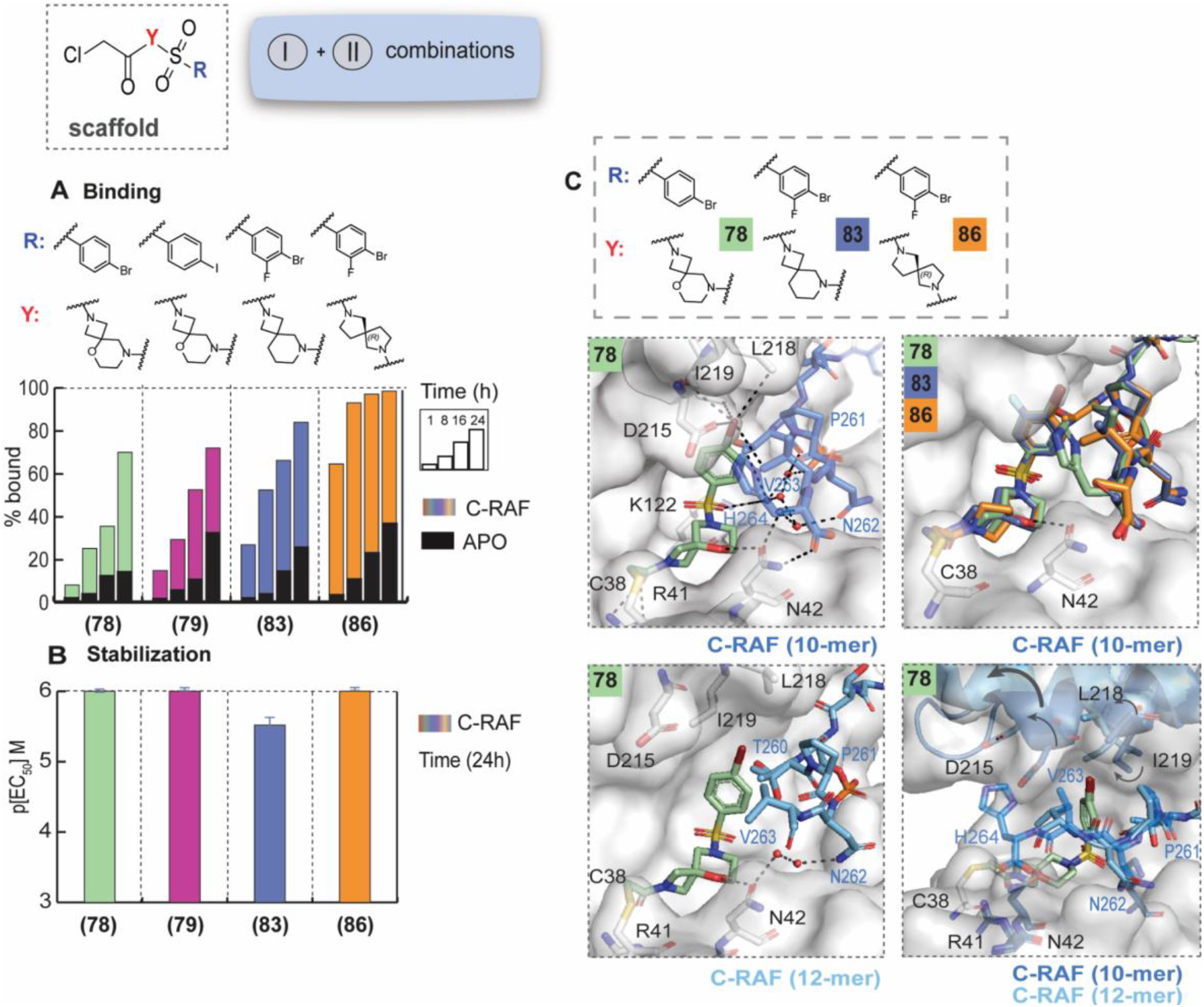
SAR and crystal structures of aryl and spiro/fused rings combinations. A) MS bar graphs at 1 μΜ. For each compound, time course experiments were performed with measurements at 1h, 8h, 16h and 24h. C-RAF259 data are shown with different colors for each compound, and apo data in black. B) Bar graphs of FA compound titration pEC_50_ values after overnight incubation. C-RAF259 data are shown with different colors for each compound. C) Top: Crystal structure of **78** (light green sticks) with 14-3-3σ/C-RAF259 10-mer and overlay with the **83** and **86** crystal structures. Bottom: crystal structure of **78** (light green sticks) with 14-3-3σ/C-RAF259 12-mer (left) and overlay of the two ternary complexes (right).

### Selectivity assessment with biophysical assays

After establishing SAR for C-RAF259, we aimed at evaluating potential selectivity; first with the A-RAF and B-RAF inhibiting sites, then with the activating sites and finally in a broader selectivity panel of diverse 14-3-3 clients. We selected 11 compounds for biophysical assays (**22, 23, 29, 32, 37, 70, 78, 79, 80, 83, 86** and compound **8** as an inactive control, as it lacked an effect on C-RAF259).

The sequences of the RAF isoform inhibiting sites are largely conserved in the regions recognized by 14-3-3; A-RAF214 and C-RAF259 have the same sequence (+1 to +6 amino acids, “TPNVHM” in the C-terminus to pS259), whereas B-RAF365 differs at the +1 position (A instead of T, Fig 5A). The +6/+7 residues for B-RAF also differ compared to A-RAF214 and C-RAF259. More differences occur in the sequences N-terminal to the phosphoserines, which are further away from the compound binding site and were expected to have a minor effect on the selectivity (detailed peptide sequences in the Methods). Based on the binding mode of the compounds, which interacted with the +1 (T), +4 (V) and +6 (M) residues of C-RAF, we thus expected similar binding/stabilization to A-RAF and lower binding/stabilization in the case of B-RAF.

**Figure 5.**
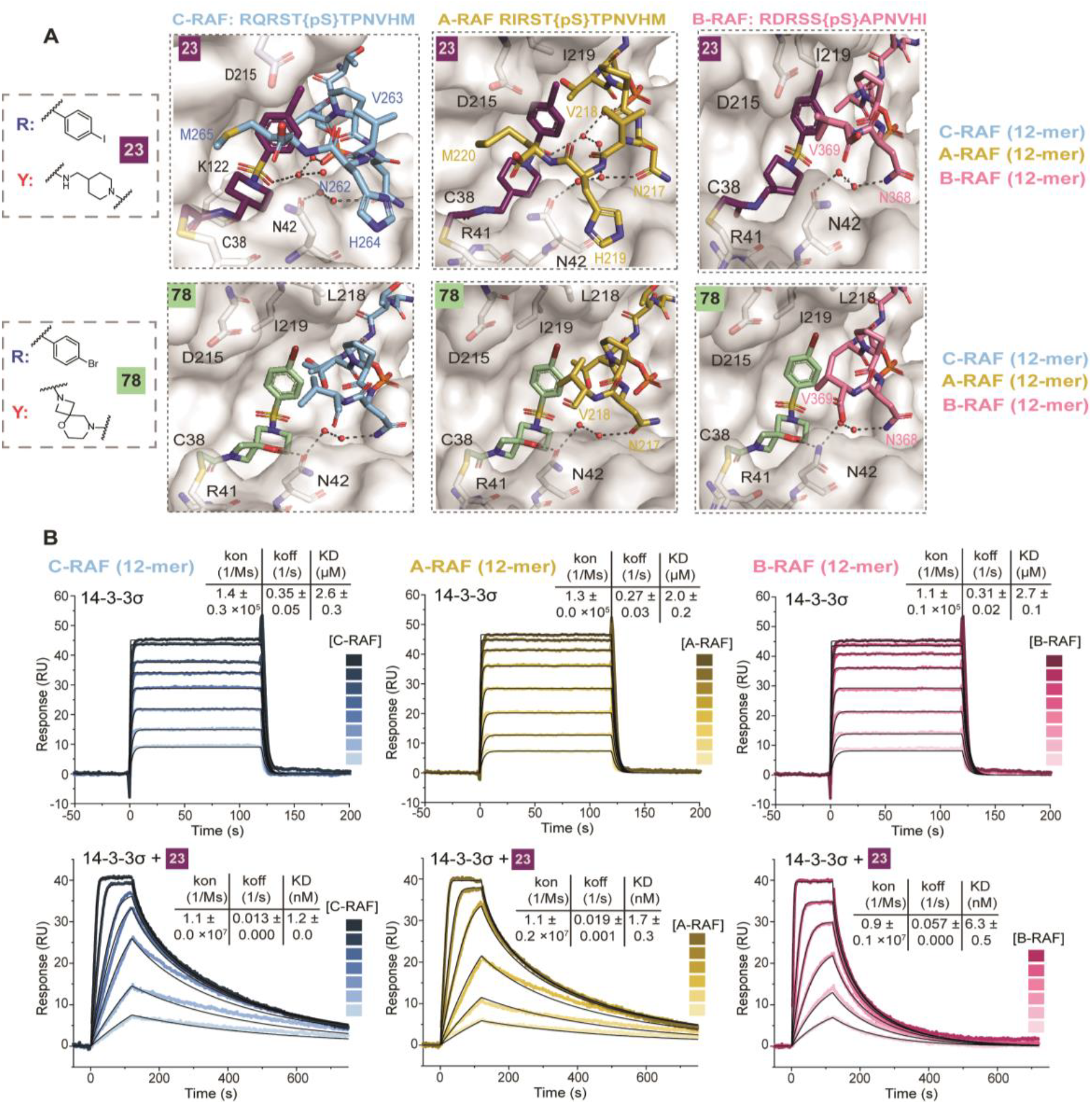
Crystallography and SPR with RAF isoforms. A) Crystal structures of **23** (top) and **78** (bottom) with 14-3-3σ/C-RAF-259 (left), 14-3-3σ/A-RAF214 (middle) and 14-3-3σ/B-RAF365 (right) 12-mer peptides. Interacting water molecules are shown as red spheres. B) SPR data of C-RAF (blue), A-RAF (yellow), and B-RAF (pink), 12-mer peptides binding to 14-3-3σ (top panel), and 14-3-3σ covalently bound to molecular glue **23** (parameters shown mean ± SD, n=2).

Initially we tried to assess selectivity from a structural perspective, using 12-mer peptides for crystallography. We solved the 14-3-3 binary complexes for A-RAF214, B-RAF365 and ternary complexes with **22, 23, 78** and **86** for both peptides (Fig 5A, S22-S24) and compared them to the C-RAF259 structures. For the binary complexes, 14-3-3 helix 9 was in the upward conformation, similar to C-RAF259. The observed density for the RAF peptides extended to the +4 residue (V218 for A-RAF214 and V369 for B-RAF365). In the presence of piperidine-containing compounds **22** and **23** with A-RAF214, the +6 M residue was visible and helix 9 was in the downward conformation. For the spiro-cycles **78** and **86** the visible density did not extend beyond the +4 V residue and helix 9 was in the upward conformation. We noticed similarities between the ternary complexes with ARAF and C-RAF, especially for **22** and **23** with a water network extending to the phosphate group. The key hydrogen bonds with backbone C38, R41, N42 (14-3-3) and the interactions with V218 and T215 (A-RAF) were consistent. For B-RAF365, for all four compounds the visible density stopped at the +4 V residue and helix 9 was in the downward conformation with the exception of **78**. The hydrogen bonds with backbone C38, R41, N42 (14-3-3) were consistent, however the compounds lacked specific interactions with the +1 A residue, which is typically considered significant for molecular recognition. As mentioned for C-RAF259, the importance of the movement of helix 9 was ambiguous, so instead we tried to correlate the ligand-protein and ligand-peptide interactions with biophysical data.

14-3-3 is considered to recognize between 2 and 20 amino acid residues on the client.^35^ However, the ideal phoshopeptide length, representative of the recognition motif for various 14-3-3 clients has not been thoroughly investigated. Typically, the +1 residues were expected to be the main driving force for molecular recognition, but in the ternary complexes of our glues with 14-3-3/CRAF259, the +4 and +6 residues were also significantly contributing to binding. Thus, taking into account the differences in C-RAF259 ternary complexes with the 10- and 12-mer peptides, we reconsidered whether this effect might also extend to the biophysical assays and further tested the compounds with peptides with different lengths. Up to this point, to establish the SAR for C-RAF we used an 11-mer acetylated peptide for the MS assay and a 10-mer FAM-labeled peptide for the FA assay. For a better comparison with crystallography, we tested the compounds again using 15-mer C-RAF peptides, before comparing them to A-RAF and B-RAF peptides. In the MS assay with the 15-mer C-RAF peptide, we noticed similar or increased binding for the piperidine-containing compounds (**22, 23, 29, 32, 37**), but a significant decrease in binding for the spiro-cycles (**78, 79, 80, 83, 86**) (table S4, Fig S25). In FA protein titrations, **22** and **23** appeared more potent with *app*K_d_ of 5 nM and 2400-fold stabilization with the 15-mer peptide (30-47 nM, 179-280-fold stabilization with the 10-mer peptide). Compounds **29, 32** and **37** showed similar *app*K_d_ values with both peptides (126 – 154 nM), whereas, in agreement with the MS data, the spiro-cycles (**78, 79, 80, 83, 86**) showed 2-3-fold weaker stabilization (*app*K_d_ 110 – 334 nM) (table S5, Fig S26). The data with the longer peptide were thus in support of the crystallography data; for the spiro-cycles fewer peptide residues were visible compared to compounds like **22**, where the density extended to the +6 M and the peptide tightly wrapped around the compound. This observation indicates potential steric hindrance of the longer RAF peptides with the spirocyclic compounds and testifies to additional opportunities for MG affinity and selectivity regulation.

To determine compound selectivity, we compared the 15-mer C-RAF data to 15-mer A-RAF. In the MS assay, the piperidine-containing compounds (**22, 23, 29, 32, 37**) showed significant binding, comparable or slightly higher to C-RAF, whereas low binding was observed for the spirocycles (table S4, Fig S27). In FA protein titrations, there were variations in the *app*K_d_ values, however it was difficult to establish a clear trend (table S5, Fig S28). We also noticed that although the A-RAF and C-RAF 15-mer peptides had similar K_d_ values (8-10 μΜ), the *app*K_d_ in the presence of DMSO differed between A-RAF and C-RAF (3 μΜ and 12 μΜ, respectively). This difference did not affect ranking of the compounds with the individual peptides but resulted in higher fold-stabilization values for C-RAF.

In the case of B-RAF365 we tested two different peptides: a 10-mer and a 15-mer and the results varied considerably, indicating that the 10-mer peptide was probably not a good representation of this PPI. In both assays, the compounds had a very small effect when the 10-mer peptide was used (table S4, S5, Fig S29-S30). The 15-mer B-RAF showed an increased compound binding compared to the 10-mer. When comparing the two B-RAF peptides with A- and C-RAF 15-mer peptides, less binding was observed with the MS assay for the piperidine analogs and consistently low binding for the spiro-compounds (table S4, Fig S31). A similar effect was observed in the FA assay, where the compounds showed lower stabilization of B-RAF in comparison to the A- and C-RAF 15-mer peptides; **22** and **23** showed *app*K_d_ of 5 nM and 448-fold-stabilization; or in other words, they were 3-fold weaker than C-RAF. Consistently, lower fold-stabilization was observed with the spiro-cycles (table S5, Fig S32). These observations were in good agreement with the crystal structures, where the compounds lacked interactions with the +1 residue and less peptide density was visible compared to A- and C-RAF.

The kinetic fundamentals behind RAF isoform selectivity were evaluated using surface plasmon resonance (SPR) experiments. 14-3-3σ, tagged with a Twinstrep-tag, was immobilized on a Strep-Tactin XT coated SPR chip. Subsequently, a 2-fold dilution series of acetylated 12-mer C-RAF, A-RAF, and B-RAF peptides were injected. The analysis of these peptides’ binary interactions with 14-3-3σ revealed K_d_ values ranging between 2.0 – 2.7 μM, determined through both kinetic- and affinity-based data fitting (Fig 5B, S33). The affinities of A-RAF and B-RAF were consistent with results from FA experiments, while for C-RAF, the K_d_ value measured by SPR was lower compared to FA (2.6 μM versus 10 μM, respectively), though this difference might be due to the fast k_off_, which makes precise measurement difficult. To compare the kinetic parameters of the binary interactions with those of the molecular glue-bound state of 14-3-3σ, the covalent bond between the chloroacetamide warhead of the compounds and C38 of 14-3-3σ was formed by overnight incubation in the presence of the C-RAF peptide. This resulting complex was immobilized on the chip and extensively washed to remove the C-RAF peptide. Afterwards, the kine ics of RAF peptides binding to the 14-3-3σ/molecular glue complex were analyzed. In the presence of **23**, the association rate (k_on_) increased 100-fold for all three RAF isoforms compared to their binary interactions, suggesting that the 14-3-3σ/**23** interface is recognized more rapidly by the RAF peptides than 14-3-3σ alone. Furthermore, the k_off_ significantly decreased in the presence of **23**, yielding *appK*_*d(compound)*_ values in the low nanomolar range (Fig 5B, S34-S36). Notably, the presence of the molecular glue resulted in mass-transport limitations, driven by the high association rate and peptide rebinding. These effects were accounted for during the fitting process (Table S6). The affinity increase was most pronounced for A-RAF and C-RAF (1000-fold and 2000-fold, respectively), whereas a weaker increase (400-fold) was observed for B-RAF, aligning with the selectivity measured in FA experiments. Since the shift in k_on_ was similar for all RAF isoforms, the difference in affinity among isoforms was attributed to a faster k_off_ for B-RAF compared to A-RAF and C-RAF. This indicated that the interactions of +1 and +6 residues (+1A and +6I for B-RAF, +1T and +6M for A-, C-RAF) with **23** were crucial for decreasing the k_off_, thereby increasing the residence time of RAF binding to 14-3-3σ. In addition, **22** induced a comparable change in k_on_ for A-RAF and C-RAF binding, relative to **23**. However, the k_off_ was increased, leading to a reduced apparent affinity, in line with FA experiments. While no notable difference was observed between **22** and **23** for B-RAF, the apparent affinity of B-RAF to 14-3-3/**22** complex remained the weakest among the RAF isoforms (Fig. S34-S36).

Regarding the activating A-RAF582, B-RAF729 and C-RAF621 sites, all isoforms contain the same +1 to +4 residues (EPSL). The +1 glutamic acid, in a published binary complex with C-RAF^621^, formed an ionic bond with K122 of 14-3-3 (PDB: 4IEA). We hypothesized that this bond could not be disrupted by the compounds, and it would also prevent the compounds binding, due to repulsive electrostatic interactions with the sulfonyl group of the compounds. This hypothesis was easily confirmed with FA protein titrations, where no stabilization was observed (Fig S37-S39).

To evaluate selectivity more broadly *in vitro*, we tested **22** and **23** in a selectivity panel that included 80 peptides representing a diverse set of 14-3-3 clients (Fig S40). Briefly, the 14-3-3σ protein concentration was set at 20% of the maximum effect for each individual peptide. The compounds were incubated overnight at calculated EC_90_ concentrations (under this assay’s conditions: 10.5 μM for **22** and **23**) in the presence of 14-3-3 and 100 nM client peptides. A client peptide with a high 14-3-3σ concentration (50 μM, maximum effect) was used as reference for normalization. As a qualitative output, clients were classified as hits when a positive change in anisotropy of more than 15 units (ca. 15%) was observed. Both compounds showed remarkable selectivity; for **23** only four clients emerged as hits (SOS1, TAZ, KC1A, TSC2), and six for **22** (SOS1, TAZ, ARGH2, H31, KC1A, TSC2). Strikingly, there were no apparent similarities in the peptide sequence for these off-target hits, except for the +2 P, which is a very common residue in 14-3-3 clients (Fig S41A). The lack of similarities in the peptide sequences suggested that conformational adaptivity was a key factor in the formation of ternary complexes and molecular recognition among different 14-3-3 clients. FA compound titrations were then performed for the common hits SOS1, TAZ, KC1A, TSC2 (Table S7, Fig S41B) and FA protein titrations were performed for SOS1, TAZ and TSC2 (Table S8, Fig S41C). The assay conditions and the C-RAF259 peptide length differed from the ones used to establish the SAR, leading to an EC_50_ = 5 µM (vs 1 µM routinely seen with the standard conditions). Nevertheless, the compounds showed greater fold-stabilization and lower EC_50_ values for C-RAF, with the second-most prominent hit being TAZ.

### Evaluation in cell assays

We further evaluated 10 compounds in cellular assays; the same compounds tested for RAF isoform selectivity, excluding the weak analog **80**. Compound **8**, as previously, served as negative control. A NanoBRET assay was developed to measure binding of full-length 14-3-3σ to C-RAF in HEK293T cells.^36^ HEK293T cells only weakly express 14-3-3σ, and show relatively low levels of pathway activity in normal growth conditions; thus, the transfected C-RAF and 14-3-3σ would be monitored without interference from endogenous expression. C-RAF was tagged with NanoLuciferase and 14-3-3σ was tagged with HaloTag/fluorescent HaloTag ligand. Stabilization of the interaction with a molecular glue was expected to lead to an increase in the BRET signal. The tested compounds resulted in a range of increased fold-stabilization (up to 1.6) and cellular EC_50_ values between 0.1 and 1.2 μΜ (Fig S42A, table S9). Piperidine-containing compounds **22** and **23** showed similar EC_50_ values (0.20 and 0.18 μΜ respectively) and fold-stabilization of 1.5 and 1.6-fold, respectively (Fig 6A). For the spiro-cycles **70** and **78**, cellular fold-stabilization was lower (1.3 and 1.4, respectively), in good agreement with the signal window observed in FA data. No increase in the BRET signal was observed for the negative control compound **8**. The stabilization effect was specific to C38 of the σ isoform of 14-3-3 – the molecular glues did not increase the BRET signal when the native C38 was mutated to N, the residue in the other 14-3-3 isoforms (Fig S42B).

**Figure 6.**
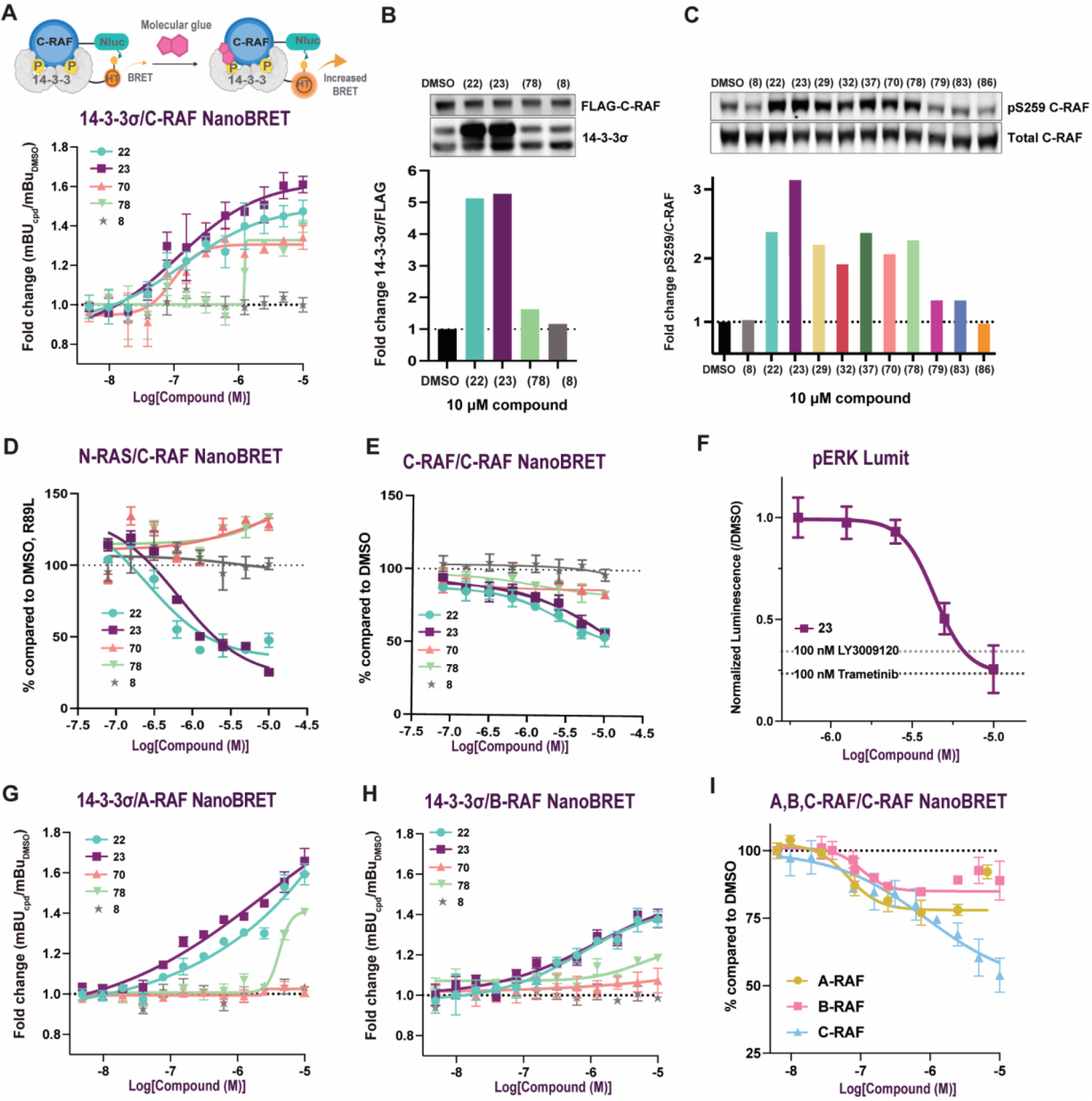
Evaluation of MGs in cell assays. A) 14-3-3σ/C-RAF NanoBRET schematic (left) and dose-response curves in HEK293T cells. B) Co-IP western blots and quantification. C) Protection of phosphorylation of the C-RAF pS259 site in HEK293T cells. D) N-RAS/C-RAF NanoBRET; 100% was considered the DMSO mBU and 0% the mBU value of NRAS/C-RAF R89L, which does not bind to C-RAF. E) C-RAF/C-RAF homodimers NanoBRET dose-response curves. F) Lumit immunoassay for pERK levels in dose-responses with **23**. G) 14-3-3σ/A-RAF NanoBRET. H) 14-3-3σ/B-RAF NanoBRET. I) NanoBRET for heterodimer formation for A-RAF/C-RAF and B-RAF/C-RAF, compared to C-RAF homodimers NanoBRET for **23**.

Binding of full-length 14-3-3σ/C-RAF was further validated in HEK293T cells through co-immunoprecipitation (co-IP) at 10 μΜ compound. NRAS Q61L, an oncogenic mutation that causes NRAS to be in its GTP-bound active state, FLAG-C-RAF, and HA-14-3-3σ were transfected and IP was performed with anti-FLAG beads. We measured a 5.3-fold increase for **23**, 5.1-fold for **22** and 1.6-fold for **78** in a 14-3-3σ pull-down with FLAG-C-RAF normalized to FLAG band intensity (Fig 6B). This effect was not observed without transfecting C-RAF, as shown for **22** (Fig S43A).

In addition to binding assays, we quantified the effect of the molecular glues on a 14-3-3 specific function – the protection of phosphorylated residues on the client proteins from phosphatases (Fig 6C, table S9). By western blots in HEK293T cells, with 10 μΜ compound we measured 2.4-fold and 3.2-fold increases in phosphorylation of the C-RAF259 site relatively to total C-RAF for **22** and **23**, respectively. The spirocycles **70** and **78** showed a slightly smaller effect (2.1-fold and 2.2-fold, respectively). In MIA PaCa-2 cells, a KRAS G12C dependent cell line, compounds **22** and **23** showed a 1.4-fold and 1.5-fold increase in endogenous C-RAF pS259 levels relative to total C-RAF (Fig S43B, table S9).

As mentioned in the introduction, the activation mechanism of C-RAF is assumed to proceed in a similar manner as B-RAF, where the first step is the translocation to the membrane and the interaction with RAS. To study the effect of the molecular glues on this step of the pathway, we developed a NanoBRET assay to measure the interaction between N-RAS Q61L and C-RAF in HEK293T cells (Fig 6D). For the quantification of the N-RAS/C-RAF NanoBRET, we set the DMSO-treated cell-signal as 100% and the signal for NRAS/C-RAF R89L, a C-RAF mutation that prevented C-RAF from interacting with RAS, as the 0% control.^28,37^ The stabilization of the interaction between 14-3-3σ and C-RAF259 by the molecular glues was expected to compete for the interaction of N-RAS/C-RAF. As expected, treatment with **23** resulted in 75% signal reduction of the N-RAS Q61L/C-RAF interaction and 52% with **22**. For **70** and **78** no clear trend was observed. Compound **8** did not affect N-RAS Q61L/C-RAF binding.

The final activation step of the pathway includes the dimerization of RAF kinase domains. Dimer formation is quite complex and different RAF homo- and hetero-dimers can occur, depending on the cell type or cancer type. We aimed to study potential effects of the molecular glues on RAF-dimer formation using a NanoBRET assay. The C-RAF/C-RAF NanoBRET assays were performed with N-RAS Q61L to ensure pathway activation^38^ (Fig 6E). We observed 43% decrease in C-RAF homodimer formation with **22**, 47% decrease with **23**, and ∼20% for **70** and **78**.

To test the downstream MAPK effects of stabilizing 14-3-3/C-RAF, we developed a Lumit immunoassay to analyze the levels of pERK. The secondary antibody pair, lysis buffer, and primary antibody concentration were optimized for the ERK/pERK antibodies used (data not shown). 14-3-3σ and C-RAF were transfected in a 1:1 ratio in HEK293T cells. Cells were treated with **23**, a MEK inhibitor (Trametinib, 100 nM), or a pan-RAF inhibitor (LY3009120, 100 nM) in 0.5% FBS FluoroBrite DMEM for 24 hours, followed by EGF stimulation (100 ng/μL, 30 minutes) before performing the Lumit immunoassay protocol. We corrected for cell number using the fluorogenic live cell GF-AFC substrate. Data was normalized to DMSO-treated samples. Compound **23** dosed at 10 μM inhibited the phosphorylation of ERK to a similar extent as the MEK inhibitor (Trametinib) and slightly increased the inhibition of pERK compared to the pan-RAF inhibitor (LY3009120); the IC_50_ value for **23** was 4.4 μM (Fig 6F). The Lumit data suggested stabilization of 14-3-3/C-RAF was an effective mechanism to modulate the MAPK pathway.

We developed additional NanoBRET assays to quantify compound binding to 14-3-3σ/A-RAF and 14-3-3σ/B-RAF and determined RAF selectivity in a cellular context (Fig 6G, 6H, Fig S44, table S10). Compounds **22** and **23** showed binding in the 14-3-3σ/A-RAF NanoBRET, giving 1.6-fold and 1.7-fold increase, respectively, at 10 µM; dose-response data did not plateau. **70** was inactive and **78** showed an effect only in the two highest compound concentrations. Similarly, in the 14-3-3σ/B-RAF NanoBRET, increased binding was observed in the presence of **22** and **23** (1.40-fold increase) and a minor effect for **70** and **78**. Quantitively, **22** and **23** showed comparable fold-stabilization for A- and C-RAF, whereas **70** and **78** showed significant selectivity for C-RAF, especially in lower compound concentrations. In all cases, binding was dependent on C38, as the compounds were inactive with the C38N mutant (data not shown). We also tested the formation of C-RAF heterodimers, A-RAF/C-RAF and B-RAF/C-RAF. Compound **23** exhibited inhibition of heterodimers, decreasing the A-RAF/C-RAF interaction by 23% and the B-RAF/C-RAF dimer by 15% (Fig 6I).

The compound effect on the phosphorylation protection of the inhibiting A-RAF pS214 site was also quantified and compared to the effect on the C-RAF pS259 site (Fig S45, table S10). Compound **23** resulted in 1.3-fold increase in the case of A-RAF and 3.2-fold increase in the case of C-RAF. Compound **70** did not increase the A-RAF phosphorylation, whereas it showed 2.1-fold increase for C-RAF. A similar experiment was performed for the B-RAF pS365 site; however, the quantification of the Western blots was inconsistent, potentially due to the antibody used (data not shown).

Taken together, the cell data suggests on-target activity on the C-RAF259/14-3-3σ complex for the most potent compounds **22** and **23**, a protective effect on C-RAF259 phosphorylation, disruption of C-RAF active kinase dimers, and decrease in phosphorylation of the downstream MAPK target pERK. Regarding SAR, the presence of a piperidine ring was more favorable in cell assays, compared to the spirocycles, supporting our crystal structures, where interactions occur with M265 (+6 residue). Interestingly, while the piperidine containing **22** and **23** favored C-RAF over A- and B-RAFs, the spirocycles showed an even stronger selectivity for C-RAF.

## CONCLUSIONS

In summary, we describe a fragment-merging approach for the development of cell-active molecular glues targeting the inhibitory 14-3-3σ/C-RAF259 complex. Validation of the compounds in multiple biophysical assays, from mass spectrometry to fluorescence anisotropy and surface plasmon resonance allowed the quantification of binding, stabilization, and kinetics, elucidating the factors involved in the formation of cooperative ternary complexes. The establishment of SAR for the 14-3-3σ/C-RAF259 complex was achieved with biophysical assays using phospho-peptides as representations of the full-length client. MS and FA assays were in good agreement in establishing SAR for a given peptide length for C-RAF259. However, we found that the longer C-RAF peptides led to differentiated results, especially in the case of the spirocyclic compounds, where the rigid part of the compound was in close proximity to the flexible region of the peptides. These effects were not exclusively steric, based on space-filling models; rather, we hypothesized that entropy and/or less optimal interactions with the +4 and +6 peptide residues were also involved. The peptide length also had a strong effect on compound evaluation for B-RAF, indicating again that residues involved in molecular recognition can extend well beyond the +1 – here, as far as the +4 and +6 residues. By comparing data using peptides of different lengths, we aimed to emphasize that for 14-3-3 clients where the phosphorylation site lies on a highly dynamic loop, conformational adaptivity can have a significant effect both on the biophysical assays and crystallography. Thus, in cases where the sequence is internal the length should be chosen carefully.

From crystal structures with different RAF peptide lengths, we observed differences not only in the peptide conformation, which was expected, but also in the positioning of helix 9 of 14-3-3. The movement was not always compound- or peptide-specific, but it could have a significant effect on the formed ligand-protein interactions. It is noteworthy that the downward positioning of this helix is consistent with a cryo-EM structure of 14-3-3/B-RAF (PDB 6NYB), indicating that it is unlikely to be a crystallographic observation (Fig S46). Full length, inhibitory 14-3-3/C-RAF cryo-EM structures are still missing, thus a direct comparison with our crystal structures is not feasible yet.

We further evaluated the 10 most potent compounds from the biophysical assays in cell assays with the full-length RAF clients. We noticed good potency correlation in most of the cases. Compounds **22** and **23** were the most potent in cell assays and stabilized both the 14-3-3/C-RAF and 14-3-3/A-RAF complexes, whereas spirocycles **70** and **78** showed greater selectivity for C-RAF in the NanoBRET assay and the protection of phosphorylation. Taken together, our cell data indicate that the molecular glues showed on target activity, stabilizing the 14-3-3σ/C-RAF interaction and protecting the pS259 site from dephosphorylation. Mechanistically, we see the expected inhibition of the N-RAS/C-RAF interaction, as well as C-RAF dimerization in an activated pathway for the most potent analogs. These data indicate that the molecular glues targeted the “auto”inhibited 14-3-3/C-RAF monomer, thus shifting the equilibrium away from the active form of C-RAF. The effect on the structurally similar A-RAF isoform was chemotype-dependent, with piperidine compounds **22** and **23** being less selective compared to spirocycles **70** and **78**. The observed differences in A-RAF stabilization are intriguing, considering the sequence similarity with C-RAF on the 14-3-3 binding grove. Possible explanations could be differences in the N-terminus, dynamics, or regulation on a cellular level. The similarity of the A-RAF isoform could explain the observed inhibition of A-RAF/C-RAF complexes with **23** whereas the B-RAF/C-RAF dimer was minimally affected.

Quantifying the downstream effects of stabilizing 14-3-3/C-RAF, **23** prevented the phosphorylation of ERK to similar levels as other modes of MAPK modulation. However, **23**, was slower to modulate the pathway than Trametinib or LY3009120, taking 18 to 24 hours compared to 3 hours. We hypothesize that the time delay is due to the slow reaction of the covalent warhead, rather than the intrinsic biology of 14-3-3/C-RAF stabilization. The kinetics do complicate the biological analysis, since the MAPK pathway includes multiple feedback loops and varied dimer formation. These might explain why we do not see full inhibition of NRAS/C-RAF, nor do we observe on-target cell death in cancer cells like MIA PaCa-2. Park et al,^39^ solved the cryo-EM structure of K-RAS and MEK bound to 14-3-3/BRAF, in which BRAF pS365 (the equivalent to CRAF pS259) is still phosphorylated and bound to 14-3-3. Interestingly, our data indicate that 14-3-3/C-RAF stabilization can significantly inhibit binding to RAS, suggesting that multiple mechanisms of RAF activation (and/or inhibition) are possible.

Additionally, recent literature suggests that C-RAF has roles independent from the MAPK signaling pathway, which we have not explored. For instance, Venkatanarayan *et al*. showed that C-RAF depletion inhibits tumor growth but not ERK signaling in KRAS mutant cells, and upon C-RAF loss, A-RAF dimers promote ERK signaling, leading to cell-cycle arrest.^23^ Sanclemente *et al*. reported that in KRASG12V/Trp p53 mutant lung tumors, systematic abrogation of C-RAF expression did not inhibit canonical MAPK signaling, which in contrast to MEK inhibitors, resulted in limited toxicities.^40^ For these types of tumors, significant levels of phospho-MEK and phospho-ERK expression were retained, despite the elimination of C-RAF expression in most of their cells.^40^ Alternatively, the C-RAF preference of our glues may limit phenotypic changes. Müller *et al*. showed that selective knockdown of a single RAF isoform had at best partial effects on the phosphorylation levels of MEK1/2 and ERK1/2 indicating functional redundancy and/or compensatory mechanisms between the RAF isoforms, while pan-RAF activity affected PI3K-dependent signaling.^24^ Deciphering the precise effect of these molecular glues on the C-RAF biology and broader pathway changes is beyond the scope of the current study. We believe, however, that our molecular glues will be useful chemical biology tools that extend beyond direct inhibition and focusing on regulation of proteinprotein interactions. Additionally, our cell-active compounds could be useful starting points for medicinal chemistry campaigns aiming either at pan-RAF/14-3-3 glues or RAF-isoform specific glues to further elucidate pathway biology.

## ASSOCIATED CONTENT

### Supporting Information

Experimental methods, supplementary figures and tables, synthetic procedures, compound characterization, NMR spectra, crystallography data (PDF). This material is available free of charge via the Internet at http://pubs.acs.org.

## AUTHOR INFORMATION

### Present Addresses

‡ Christian Ottmann: Ambagon Therapeutics, de Lismortel 31, 5612AR Eindhoven, the Netherlands

## Author Contributions

M.K. performed the design and synthesis of the compounds, the MS and FA assays. H.R.V co-developed the NanoBRET assays and performed the cell assays. M.A.M.P. solved the majority of the crystal structures and performed the SPR assay. J.M.V codeveloped the NanoBRET assays, developed the Lumit immunoassay, and performed cell assays. E.J.V solved initial crystal structures. S.D.B. and M.S. performed the selectivity panel. S.Z.C. co-developed the Lumit immunoassay. L.C.Y. assisted in the development of cell assays. M.C.M. v.d. O. assisted in crystallography setup. F. M. provided feedback and guidance for the cell assays. C.O., L.B. and M.R.A. designed and supervised the project. M.K. wrote the manuscript with contributions from all authors.

‡ Markella Konstantinidou, Holly R. Vickery, Marloes A.M. Pennings contributed equally.

## Funding Sources

This research was funded by the Ono Pharma Foundation Breakthrough Science Initiative Award and NIH/NIGMS GM147696 and the Netherlands Organization for Scientific Research (NWO) through Gravity program 024.001.035, ECHO grant 711.018.003, Nanotorch OCenW.M20.200.

## Notes

The authors declare the following competing interest: M.R.A., C.O., and L.B. are founders of Ambagon Therapeutics.

## ACKNOWLEDGMENT

We thank Amanda Paulson for the automated mass spec data processing infrastructure in the SMDC. We thank Loes Stevers for useful discussions and feedback, and Maurits Overmans and Yan Ni for technical support. We acknowledge DESY (Hamburg, Germany), a member of the Helmholtz Association HGF, for the provision of experimental facilities. Parts of this research were carried out at PETRA III and we would like to thank Johanna Hakanpää for assistance in using beam P11. Beamtime was allocated for proposals 11010503, 11012310, 11011126, 11010888 and 11012787. Further, we acknowledge the European Synchrotron Radiation Facility (ESRF) for provision of synchrotron radiation facilities, and we would like to thank Romain Talon, Daniele de Sanctis, Shibom Basu, Matthew Bowler and David Flot, for assistance and support in using beamlines ID23-1, ID23-2, ID30A and ID30B (mx2268). The authors also thank Diamond Light Source for beamtime (proposal mx19800-27), and the staff of beamline I03 for assistance with crystal testing and data collection. We want to acknowledge Leon Kraakman, Christin Radon, and Anja Drescher from Cytiva for advice on fitting the SPR data.

## TOC figure

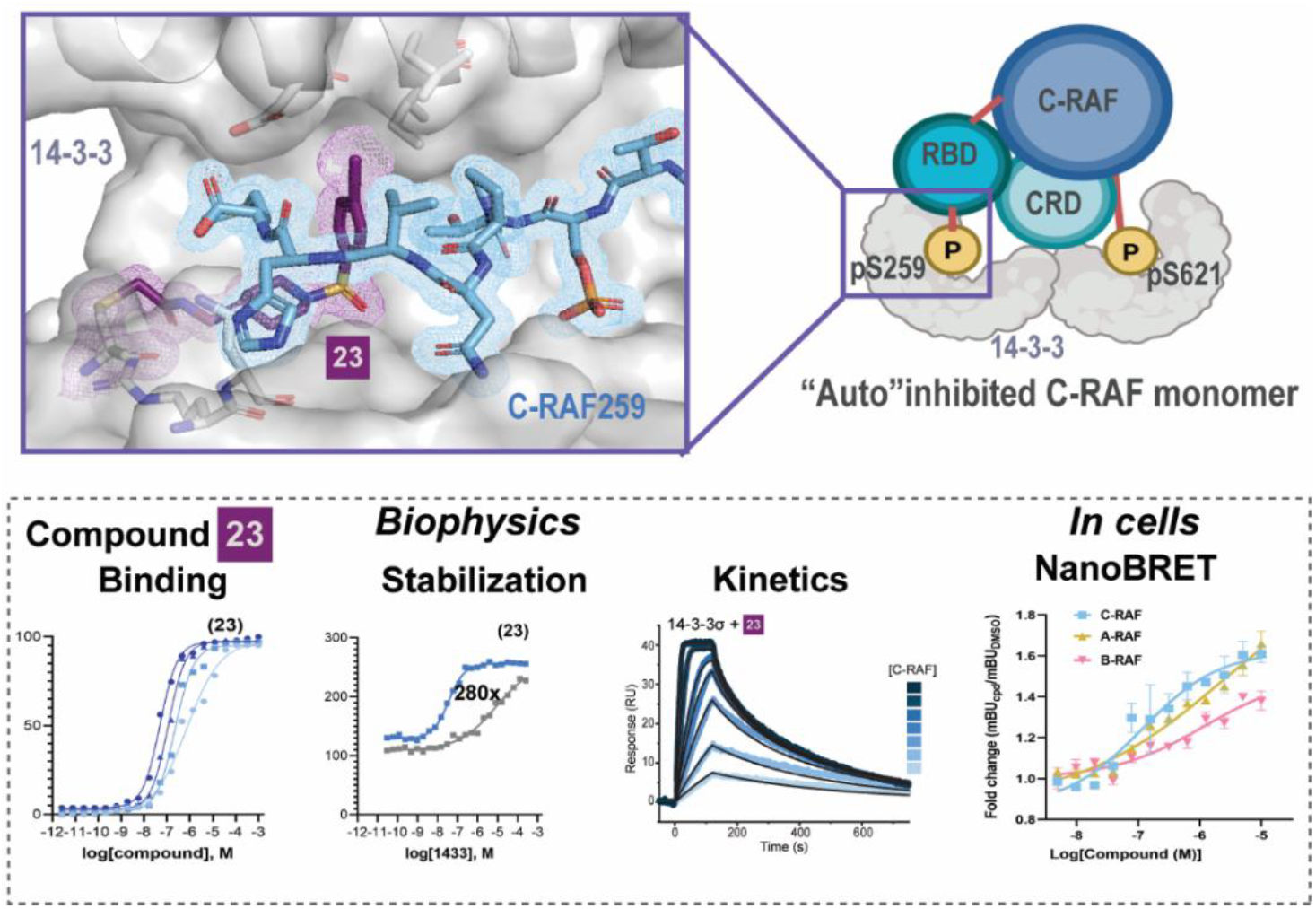

## REFERENCES

(1) Dhillon, A. S.; Hagan, S.; Rath, O.; Kolch, W. MAP Kinase Signalling Pathways in Cancer. Oncogene 2007, 26 (22), 3279–3290. 10.1038/sj.onc.1210421.

(2) Fernandez-Medarde, A.; Santos, E. Ras in Cancer and Developmental Diseases. Genes & Cancer 2011, 2 (3), 344–358. 10.1177/1947601911411084.

(3) Davies, H.; Bignell, G. R.; Cox, C.; Stephens, P.; Edkins, S.; Clegg, S.; Teague, J.; Woffendin, H.; Garnett, M. J.; Bottomley, W.; Davis, N.; Dicks, E.; Ewing, R.; Floyd, Y.; Gray, K.; Hall, S.; Hawes, R.; Hughes, J.; Kosmidou, V.; Menzies, A.; Mould, C.; Parker, A.; Stevens, C.; Watt, S.; Hooper, S.; Wilson, R.; Jayatilake, H.; Gusterson, B. A.; Cooper, C.; Shipley, J.; Hargrave, D.; Pritchard-Jones, K.; Maitland, N.; Chenevix-Trench, G.; Riggins, G. J.; Bigner, D. D.; Palmieri, G.; Cossu, A.; Flanagan, A.; Nicholson, A.; Ho, J. W. C.; Leung, S. Y.; Yuen, S. T.; Weber, B. L.; Seigler, H. F.; Darrow, T. L.; Paterson, H.; Marais, R.; Marshall, C. J.; Wooster, R.; Stratton, M. R.; Futreal, P. A. Mutations of the BRAF Gene in Human Cancer. Nature 2002, 417 (6892), 949–954. 10.1038/nature00766.

(4) Zhang, W. BRAF Inhibitors: The Current and the Future. Current Opinion in Pharmacology 2015, 23, 68–73. 10.1016/j.coph.2015.05.015.

(5) Drosten, M.; Barbacid, M. Targeting the MAPK Pathway in KRAS-Driven Tumors. Cancer Cell 2020, 37 (4), 543–550. 10.1016/j.ccell.2020.03.013.

(6) Kondo, Y.; Ognjenović, J.; Banerjee, S.; Karandur, D.; Merk, A.; Kulhanek, K.; Wong, K.; Roose, J. P.; Subramaniam, S.; Kuriyan, J. Cryo-EM Structure of a Dimeric B-Raf:14-3-3 Complex Reveals Asymmetry in the Active Sites of B-Raf Kinases. Science 2019, 366 (6461), 109–115. 10.1126/science.aay0543.

(7) Park, E.; Rawson, S.; Li, K.; Kim, B.-W.; Ficarro, S. B.; Pino, G. G.-D.; Sharif, H.; Marto, J. A.; Jeon, H.; Eck, M. J. Architecture of Autoinhibited and Active BRAF-MEK1-14-3-3 Complexes. Nature 2019, 575 (7783), 545–550. 10.1038/s41586-019-1660-y.

(8) Martinez Fiesco, J. A.; Durrant, D. E.; Morrison, D. K.; Zhang, P. Structural Insights into the BRAF Monomer-to-Dimer Transition Mediated by RAS Binding. Nat Commun 2022, 13 (1), 486. 10.1038/s41467-022-28084-3.

(9) Cutler, R. E.; Stephens, R. M.; Saracino, M. R.; Morrison, D. K. Autoregulation of the Raf-1 Serine/Threonine Kinase. Proc. Natl. Acad. Sci. U.S.A. 1998, 95 (16), 9214–9219. 10.1073/pnas.95.16.9214.

(10) Chong, H.; Guan, K.-L. Regulation of Raf through Phosphorylation and N Terminus-C Terminus Interaction. Journal of Biological Chemistry 2003, 278 (38), 36269–36276. 10.1074/jbc.M212803200.

(11) Tzivion, G.; Luo, Z.; Avruch, J. A Dimeric 14-3-3 Protein Is an Essential Cofactor for Raf Kinase Activity. Nature 1998, 394 (6688), 88. 10.1038/27938.

(12) Garnett, M. J.; Rana, S.; Paterson, H.; Barford, D.; Marais, R. Wild-Type and Mutant B-RAF Activate C-RAF through Distinct Mechanisms Involving Heterodimerization. Molecular Cell 2005, 20 (6), 963–969. 10.1016/j.molcel.2005.10.022.

(13) Aitken, A. 14-3-3 Proteins: A Historic Overview. Semin Cancer Biol 2006, 16 (3), 162–172. 10.1016/j.semcancer.2006.03.005.

(14) Sluchanko, N. N. Reading the Phosphorylation Code: Binding of the 14-3-3 Protein to Multivalent Client Phosphoproteins. Biochemical Journal 2020, 477 (7), 1219–1225. 10.1042/BCJ20200084.

(15) Roskoski, R. RAF Protein-Serine/Threonine Kinases: Structure and Regulation. Biochemical and Biophysical Research Communications 2010, 399 (3), 313–317. 10.1016/j.bbrc.2010.07.092.

(16) Wellbrock, C.; Karasarides, M.; Marais, R. The RAF Proteins Take Centre Stage. Nat Rev Mol Cell Biol 2004, 5 (11), 875–885. 10.1038/nrm1498.

(17) Lavoie, H.; Therrien, M. Regulation of RAF Protein Kinases in ERK Signalling. Nat Rev Mol Cell Biol 2015, 16 (5), 281–298. 10.1038/nrm3979.

(18) García-Alonso, S.; Mesa, P.; Ovejero, L. D. L. P.; Aizpurua, G.; Lechuga, C. G.; Zarzuela, E.; Santiveri, C. M.; Sanclemente, M.; Muñoz, J.; Musteanu, M.; Campos-Olivas, R.; Martínez-Torrecuadrada, J.; Barbacid, M.; Montoya, G. Structure of the RAF1HSP90-CDC37 Complex Reveals the Basis of RAF1 Regulation. Molecular Cell 2022, 82 (18), 3438-3452.e8. 10.1016/j.molcel.2022.08.012.

(19) Okamoto, K.; Sako, Y. Two Closed Conformations of CRAF Require the 14-3-3 Binding Motifs and Cysteine-Rich Domain to Be Intact in Live Cells. Journal of Molecular Biology 2023, 435 (6), 167989. 10.1016/j.jmb.2023.167989.

(20) Young, L. C.; Hartig, N.; Boned Del Río, I.; Sari, S.; Ringham-Terry, B.; Wainwright, J. R.; Jones, G. G.; McCormick, F.; Rodriguez-Viciana, P. SHOC2MRAS-PP1 Complex Positively Regulates RAF Activity and Contributes to Noonan Syndrome Pathogenesis. Proc Natl Acad Sci U S A 2018, 115 (45), E10576–E10585. 10.1073/pnas.1720352115.

(21) Liau, N. P. D.; Johnson, M. C.; Izadi, S.; Gerosa, L.; Hammel, M.; Bruning, J. M.; Wendorff, T. J.; Phung, W.; Hymowitz, S. G.; Sudhamsu, J. Structural Basis for SHOC2 Modulation of RAS Signalling. Nature 2022, 609 (7926), 400–407. 10.1038/s41586-022-04838-3.

(22) Rushworth, L. K.; Hindley, A. D.; O’Neill, E.; Kolch, W. Regulation and Role of Raf-1/B-Raf Heterodimerization. Molecular and Cellular Biology 2006, 26 (6), 2262–2272. 10.1128/MCB.26.6.22622272.2006.

(23) Venkatanarayan, A.; Liang, J.; Yen, I.; Shanahan, F.; Haley, B.; Phu, L.; Verschueren, E.; Hinkle, T. B.; Kan, D.; Segal, E.; Long, J. E.; Lima, T.; Liau, N. P. D.; Sudhamsu, J.; Li, J.; Klijn, C.; Piskol, R.; Junttila, M. R.; Shaw, A. S.; Merchant, M.; Chang, M. T.; Kirkpatrick, D. S.; Malek, S. CRAF Dimerization with ARAF Regulates KRAS-Driven Tumor Growth. Cell Reports 2022, 38 (6), 110351. 10.1016/j.celrep.2022.110351.

(24) Müller, E.; Bauer, S.; Stühmer, T.; Mottok, A.; Scholz, C.-J.; Steinbrunn, T.; Brünnert, D.; Brandl, A.; Schraud, H.; Kreßmann, S.; Beilhack, A.; Rosenwald, A.; Bargou, R. C.; Chatterjee, M. Pan-Raf Co-Operates with PI3K-Dependent Signalling and Critically Contributes to Myeloma Cell Survival Independently of Mutated RAS. Leukemia 2017, 31 (4), 922–933. 10.1038/leu.2016.264.

(25) Hatzivassiliou, G.; Song, K.; Yen, I.; Brandhuber, B. J.; Anderson, D. J.; Alvarado, R.; Ludlam, M. J. C.; Stokoe, D.; Gloor, S. L.; Vigers, G.; Morales, T.; Aliagas, I.; Liu, B.; Sideris, S.; Hoeflich, K. P.; Jaiswal, B. S.; Seshagiri, S.; Koeppen, H.; Belvin, M.; Friedman, L. S.; Malek, S. RAF Inhibitors Prime WildType RAF to Activate the MAPK Pathway and Enhance Growth. Nature 2010, 464 (7287), 431– 435. 10.1038/nature08833.

(26) Morgan, C. W.; Dale, I. L.; Thomas, A. P.; Hunt, J.; Chin, J. W. Selective CRAF Inhibition Elicits Transactivation. J. Am. Chem. Soc. 2021, 143 (12), 4600– 4606. 10.1021/jacs.0c11958.

(27) Poulikakos, P. I.; Zhang, C.; Bollag, G.; Shokat, K. M.; Rosen, N. RAF Inhibitors Transactivate RAF Dimers and ERK Signalling in Cells with Wild-Type BRAF. Nature 2010, 464 (7287), 427–430. 10.1038/nature08902.

(28) Heidorn, S. J.; Milagre, C.; Whittaker, S.; Nourry, A.; Niculescu-Duvas, I.; Dhomen, N.; Hussain, J.; Reis-Filho, J. S.; Springer, C. J.; Pritchard, C.; Marais, R. Kinase-Dead BRAF and Oncogenic RAS Cooperate to Drive Tumor Progression through CRAF. Cell 2010, 140 (2), 209–221. 10.1016/j.cell.2009.12.040.

(29) Somsen, B. A.; Cossar, P. J.; Arkin, M. R.; Brunsveld, L.; Ottmann, C. 14-3-3 Protein-Protein Interactions: From Mechanistic Understanding to Their Small-Molecule Stabilization. ChemBioChem 2024, 25 (14). 10.1002/cbic.202400214.

(30) Molzan, M.; Kasper, S.; Röglin, L.; Skwarczynska, M.; Sassa, T.; Inoue, T.; Breitenbuecher, F.; Ohkanda, J.; Kato, N.; Schuler, M.; Ottmann, C. Stabilization of Physical RAF/14-3-3 Interaction by Cotylenin A as Treatment Strategy for RAS Mutant Cancers. ACS Chem Biol 2013, 8 (9), 1869–1875. 10.1021/cb4003464.

(31) Kenanova, D. N.; Visser, E. J.; Virta, J. M.; Sijbesma, E.; Centorrino, F.; Vickery, H. R.; Zhong, M.; Neitz, R. J.; Brunsveld, L.; Ottmann, C.; Arkin, M. R. A Systematic Approach to the Discovery of Protein– Protein Interaction Stabilizers. ACS Cent. Sci. 2023, 9 (5), 937–946. 10.1021/acscentsci.2c01449.

(32) Konstantinidou, M.; Visser, E. J.; Vandenboorn, E.; Chen, S.; Jaishankar, P.; Overmans, M.; Dutta, S.; Neitz, R. J.; Renslo, A. R.; Ottmann, C.; Brunsveld, L.; Arkin, M. R. Structure-Based Optimization of Covalent, Small-Molecule Stabilizers of the 14-3-3σ/ERα Protein–Protein Interaction from Nonselective Fragments. J. Am. Chem. Soc. 2023, 145 (37), 20328–20343. 10.1021/jacs.3c05161.

(33) Wolter, M.; Valenti, D.; Cossar, P. J.; Hristeva, S.; Levy, L. M.; Genski, T.; Hoffmann, T.; Brunsveld, L.; Tzalis, D.; Ottmann, C. An Exploration of Chemical Properties Required for Cooperative Stabilization of the 14-3-3 Interaction with NF-κB—Utilizing a Reversible Covalent Tethering Approach. J. Med. Chem. 2021, 64 (12), 8423–8436. 10.1021/acs.jmedchem.1c00401.

(34) Shinada, N. K.; De Brevern, A. G.; Schmidtke, P. Halogens in Protein–Ligand Binding Mechanism: A Structural Perspective. J. Med. Chem. 2019, 62 (21), 9341–9356. 10.1021/acs.jmedchem.8b01453.

(35) Milroy, L.-G.; Brunsveld, L.; Ottmann, C. Stabilization and Inhibition of Protein-Protein Interactions: The 14-3-3 Case Study. ACS Chem Biol 2013, 8 (1), 27–35. 10.1021/cb300599t.

(36) Vickery, H. R.; Virta, J. M.; Konstantinidou, M.; Arkin, M. R. Development of a NanoBRET Assay for Evaluation of 14-3-3σ Molecular Glues. SLAS Discovery 2024, 100165. 10.1016/j.slasd.2024.100165.

(37) Fabian, J. R.; Vojtek, A. B.; Cooper, J. A.; Morrison, D.. A Single Amino Acid Change in Raf-1 Inhibits Ras Binding and Alters Raf-1 Function. Proc. Natl. Acad. Sci. U.S.A. 1994, 91 (13), 5982–5986. 10.1073/pnas.91.13.5982.

(38) Hobbs, G. A.; Der, C. J.; Rossman, K. L. RAS Isoforms and Mutations in Cancer at a Glance. Journal of Cell Science 2016, 129 (7), 1287–1292. 10.1242/jcs.182873.

(39) Park, E.; Rawson, S.; Schmoker, A.; Kim, B.-W.; Oh, S.; Song, K.; Jeon, H.; Eck, M. J. Cryo-EM Structure of a RAS/RAF Recruitment Complex. Nat Commun 2023, 14 (1). 10.1038/s41467-023-40299-6.

(40) Sanclemente, M.; Francoz, S.; Esteban-Burgos, L.; Bousquet-Mur, E.; Djurec, M.; Lopez-Casas, P. P.; Hidalgo, M.; Guerra, C.; Drosten, M.; Musteanu, M.; Barbacid, M. C-RAF Ablation Induces Regression of Advanced Kras/Trp53 Mutant Lung Adenocarcinomas by a Mechanism Independent of MAPK Signaling. Cancer Cell 2018, 33 (2), 217-228.e4. 10.1016/j.ccell.2017.12.014.

