## Supplemental Information for "Modulation of the 14-3-3σ/C-RAF “auto”inhibited complex by molecular glues"

*[1]Department of Pharmaceutical Chemistry and Small Molecule Discovery Center (SMDC), University of California, San Francisco 94143, United States, [2] Laboratory of Chemical Biology, Department of Biomedical Engineering and Institute for Complex Molecular Systems (ICMS), Eindhoven University of Technology, 5600 MB Eindhoven, The Netherlands, [3]Ambagon Therapeutics, de Lismortel 31, 5612AR Eindhoven, the Netherlands, [4] Helen Diller Family Comprehensive Cancer Center, University of California, San Francisco, CA, USA*

[‡] Authors contributed equally

**Table of Contents**

| | |
|---|---|
| **1. Supplementary Methods** | p1 – 6 |
| **2. Supplementary Figures** | |
| Figures S1 – S46 | p7 – 41 |
| **3. Supplementary Tables** | |
| Table S1: Overview of molecular structures of compounds and assays | p42 – 45 |
| Table S2: MS data (apo, C-RAF259) | p46 – 47 |
| Table S3: FA compound and protein titrations (C-RAF259) | p47 – 48 |
| Table S4: MS data (A-RAF214, B-RAF365) | p48 |
| Table S5: FA protein titrations (A-RAF214, B-RAF365) | p49 |
| Table S6: SPR data | p50 – 51 |
| Table S7: FA compound titration follow-up from selectivity panel | p52 |
| Table S8: FA protein titration follow-up from selectivity panel | p52 |
| Table S9: Cell data for 14-3-3/C-RAF | p52 |
| Table S10: Cell data for 14-3-3/A-RAF and 14-3-3/B-RAF | p52 |
| Table S11: crystallography data collection parameters | p53 – 61 |
| **4. Synthetic Procedures** | p62 – 78 |
| **5. Representative NMR spectra** | p79 – 105 |
| **6. References** | p106 |

### 1. METHODS

**PROTEIN EXPRESSSION AND PURIFICATION**

The 14-3-3σ isoform (full-length for mass spectrometry and fluorescence anisotropy assays, ΔC for crystallography) with an N-terminal His6 tag was expressed in Rosetta™ 2(DE3)pLysS competent E. coli (Novagen) from a pPROEX HTb expression vector. After transformation following manufacturer's instructions, single colonies were picked to inoculate 30 mL precultures (LB), which were added to 1.5 L terrific broth (TB) medium after overnight growth at 37°C, 250 rpm. Expression was induced upon reaching $OD_{600}$ 1.9−2.1 by adding 400 µM IPTG. After overnight expression at 30 °C, 150 rpm, cells were harvested by centrifugation at 6,500 rpm, resuspended in lysis buffer (50 mM HEPES pH 7.5, 500 mM NaCl, 20 mM imidazole, 10% glycerol, 1 mM TCEP), and lysed by sonication. The His6-tagged protein was purified by Ni-affinity chromatography (Ni-NTA Agarose, Invitrogen) (Wash buffer 50 mM HEPES pH 7.5, 500 mM NaCl, 20 mM imidazole, 1 mM TCEP; Elution buffer 50 mM HEPES pH 7.5, 500 mM NaCl, 500 mM imidazole, 1 mM TCEP) and analyzed for purity by SDS-PAGE and Q-Tof LC/MS. The protein was buffer exchanged (Storage buffer 25 mM HEPES pH 7.5, 150 mM NaCl, 1 mM TCEP) and concentrated to ~16 mg/mL and aliquots flash-frozen for storage at −80 °C. The ΔC variant was truncated at the C-terminus after T231 to enhance crystallization and after the first Ni-affinity chromatography column, the construct was treated with TEV protease to cleave off the His6 tag during dialysis (25 mM HEPES, pH 7.5, 200 mM NaCl, 5% glycerol, 10 mM $MgCl_2$, 250 µM TCEP) overnight at 4 °C. The flow-through of a second Ni-affinity column was subjected to a final purification step by size exclusion chromatography (Superdex 75 pg 16/60 size exclusion column (GE Life Science) (SEC buffer: 25 mM HEPES pH 7.5, 100 mM NaCl, 10 mM $MgCl_2$, 250 µM TCEP). The protein was concentrated to ~60 mg/mL, analyzed for purity by SDS-PAGE and Q-Tof LC/MS and aliquots flash-frozen for storage at -80 °C.

For SPR, the 14-3-3 σ isoform (FL) with a N-terminal His6-tag and C-terminal Twinstrep-tag was expressed in BL21(DE3) competent E.coli (Novagen) from a pET28a expression vector. After transformation, following manufacturer's instructions, single colonies were picked to inoculate 30mL precultures (LB), which were added to 1.5 L terrific broth (TB) medium after overnight growth at 37 °C, 250 rpm. Protein expression was induced at $OD6_{00}$ ~ 0.8 by adding isopropyl β-D-1-thiogalactopyranoside (IPTG; 0.4 mM) and cells were harvested by centrifugation (10 min, 4 °C, 16,000 × *g*) after overnight expression (18 °C, 140 rpm). Pellets were resuspended in wash buffer (50 mM Tris pH 8.0, 300 mM NaCl, 12.5 mM imidazole and 2 mM β-mercaptoethanol (βME)). After homogenizing the cells (40 bar, Emulsiflex-C3 homogenizer), the soluble fraction was collected by centrifugation (30 min, 4 °C, 40,000 × *g*) and loaded onto a $Ni^{2+}$-affinity column pre-equilibrated with wash buffer. After a washing step (wash buffer + 20 mM imidazole), the bound protein was eluted with 200 mM imidazole. After Ni-affinity chromatography the elution fraction was loaded on a StrepTactin XT column (Iba lifesciences) (wash buffer 100 mM Tris pH 8, 150 mM NaCl, 1 mM EDTA; elution buffer 100 mM Tris pH 8, 150 mM NaCl, 1 mM EDTA, 50 mM biotin) and analyzed for purity by SDS-PAGE and Q-Tof LC/MS. The protein was buffer exchanged (storage buffer 25 mM HEPES pH 8, 100 mM NaCl, 10 mM MgCl2, 0.5 mM TCEP) and concentrated to 34.8 mg/mL and aliquots flash-frozen for storage at -80 °C.

**PEPTIDES**

Peptides for mass spectrometry assays were purchased from Elim Biopharmaceuticals, Inc. (Hayward,CA). Peptides for X-ray crystallography were purchased from GenScript Biotech Corp. Fluorescein-labeled (5-FAM) peptides for fluorescence anisotropy assays were purchased from Elim Biopharmaceuticals, Inc. (Hayward,CA) or GenScript Biotech Corp.

The following acetylated peptides were used in the MS assay:
C-RAF pS259 (11-mer): *Ac*-RQRST{pS}TPNVH-$CONH_2$,
C-RAF pS259 (15-mer): *Ac*-SQRQRST{pS}TPNVHMV-$CONH_2$,
A-RAF pS214 (15-mer) *Ac*-LQRIRST{pS}TPNVHMV-$CONH_2$,
B-RAF pS365 (10-mer): *Ac*-DRSS{pS}APNVH-$CONH_2$,
B-RAF pS365 (15-mer): *Ac*-GQRDRSS{pS}APNVHIN-$CONH_2$.
The following 5-FAM-labeled peptides were used in the FA assay:
C-RAF pS259 (10-mer): *5-FAM*-QRST{pS}TPNVH-$CONH_2$,
C-RAF pS259 (15-mer): *5-FAM*-SQRQRST{pS}TPNVHMV-$CONH_2$,
A-RAF pS214 (15-mer) *5-FAM*-LQRIRST{pS}TPNVHMV-$CONH_2$,

B-RAF pS365 (10-mer): *5-FAM*-DRSS{pS}APNVH-*$CONH_2$*,
B-RAF pS365 (15-mer): *5-FAM*-GQRDRSS{pS}APNVHIN-*$CONH_2$*,
A-RAF pS582 (15-mer): *5-FAM*-PKIERSA{pS}EPSLHRT-*$CONH_2$*,
B-RAF pS729 (15-mer): *5-FAM*-PKIHRSA{pS}EPSLNRA-*$CONH_2$*,
C-RAF pS621 (15-mer): *5-FAM*-PKINRSA{pS}EPSLHRA-*$CONH_2$.*
The following peptides were used for crystallography:
C-RAF pS259 (10-mer): QRST{pS}TPNVH,
C-RAF pS259 (12-mer): RQRST{pS}TPNVHM (used also for SPR),
A-RAF pS214 (12-mer): RIRST{pS}TPNVHM (used also for SPR),
B-RAF pS365 (12-mer): RDRSS{pS}APNVHI (used also for SPR).
The following peptides were used in the FA selectivity panel:
C-RAF pS259: FAM[Ahx]RQRST{pS}TPNVHMVS-*$CONH_2$*,
SOS1 pS1161: FITC-Ahx-PRRRPE{pS}APAESS-*$CONH_2$*,
TAZ pS89: FITC-Ahx-QHVRSH{pS}SPASLQLGTGA-*$CONH_2$*,
KC1A pS218: FAM-Ahx-LMYFNRT{pS}LPWQGLK-*$CONH_2$*,
TSC2 pS1254: FAM-Ahx-LYKSL{pS}VPAASTAK-*$CONH_2$*,
ARHG2 pS886: FAM-Ahx-PVDPRRR{pS}LPAGDAL-*$CONH_2$*.

**INTACT MASS SPECTROMETRY ASSAY**

Mass spectrometry dose response assays were performed on a Waters Acquity UPLC/ Xevo G2-XS Q-Tof mass spectrometer. A Waters UPLC Protein BEH-C4 Column (300 Å, 1.7 µm, 2.1 mm x 50 mm) was used to desalt the samples prior to application on the mass spectrometer. For 19-point MS dose responses, 50 mM compound stocks in DMSO were serially diluted in 3-fold increment in a master plate, then 1000 nL of the compound solutions were transferred in the assay plates. Master mixes containing 100 nM full-length wild type 14-3-3σ in the absence or presence of either 18 µM C-RAF259 or 16 µM A-RAF214 or 5 µM B-RAF365 were then dispensed into 384 well plates (Greiner Bio-One, catalog number 784201). Assay buffer was TRIS (10 mM, pH 8.0) and final volume per well was 50 µL, with final top concentration of compounds dose response series at 1 mM. The reaction mixtures were incubated for 1h at rt before subjected to MS. Four measurements (1h, 8h, 16h, 24h) were performed for time-course experiments. The injection volume for each sample was 6 µL. 24 µL of sample were needed for the time-course experiments, so the total volume in the assay plate was adjusted to 50 µL, to account for the dead volume in the injections. Data collection and automated processing followed a custom workflow, as previously described.[1] z-Plots were created using GraphPad Prism with the log(agonist) vs. response (variable slope, four parameters) fitting model.

**FLUORESCENCE ANISOTROPY MEASUREMENTS**

N-terminal fluorescein-labeled peptides (5-FAM), 14-3-3σ FL protein, the compounds (50 mM stock solution in DMSO) were diluted in buffer (10 mM HEPES, pH 7.5, 150 mM NaCl, 0.1% Tween20, 1 mg/mL Bovine Serum Albumin (BSA; Sigma-Aldrich). Final DMSO in the assay was less than 1%. Dilution series of 14-3-3 proteins or compounds were made in black, round-bottom 384-microwell plates (Greiner Bio-one 784900) in a final sample volume of 10 µL in triplicates.

For *compound titrations* echo acoustic dispensing was used to prepare 2-fold serial dilution series. A master mix containing 10 nM fluorescein-labeled C-RAF-pS259 peptide and 5 µM full length 14-3-3σ (concentration at $EC_{20}$ value of the protein-peptide complex) in buffer (10 mM HEPES, pH 7.5, 150 mM NaCl, 0.1% Tween20, 1 mg/mL BSA) was dispensed in the assay plates. The final volume per well was 10 µL, with final top concentration of compounds dose response series at 1mM. Each compound was measured in triplicates, in two independent experiments. Fluorescence anisotropy measurements were performed directly and after overnight incubation at room temperature. The high protein concentration used in this assay limits the sensitivity for potent compounds ($EC_{50}$ < 1µM). *Protein titrations* were made by titrating 14-3-3σ in a 2-fold dilution series (starting at 250 µM) to a mix of fluorescein-labeled peptide (10 nM) and DMSO or compound (100 µM). Fluorescence anisotropy measurements were performed after overnight incubation at room temperature. Protein titrations were more sensitive than compound titrations for highly potent compounds and the *app*$K_D$ values obtained were not limited by the protein concentration used. Fluorescence anisotropy values were measured using a Molecular Devices ID5 plate reader (filter set lex: 485 ± 20 nm, lem: 535 ± 25 nm; integration time: 50 ms; settle time: 0 ms; shake 5 sec, medium, read height 3.00 mm, G-factor = 1. Data reported are at endpoint. $EC_{50}$ and apparent $K_d$ values were obtained from fitting the data with a four-parameter logistic model (4PL) in GraphPad Prism 10 for Windows. Data was obtained and averaged based on two independent experiments.

**SELECTIVITY PANEL**

*For single point measurements*: N-terminal fluorescein-labeled peptides (5-FAM) and 14-3-3σ FL protein were diluted in buffer (10 mM HEPES, pH 7.4, 150 mM NaCl, 2 mM $MgCl_2$, 0.1% Tween20, 1 mg/mL Bovine Serum Albumin (BSA), 100 µM β-mercaptoethanol (β-ME)) transferred to black, round-bottom 384-microwell plates (Greiner Bio-one 784900). The concentration of the FAM-peptide panel was 100 nM, and the 14-3-3σ protein concentration was set at 20% of the maximum effect for each individual peptide. Compounds 22 and 23 were dispensed to the 384 well plate at 10.5 µM using the Tecan D300e digital dispenser. As reference a TAZ peptide with a high 14-3-3σ concentration (50 µM, maximum effect) was used. Plates were incubated overnight at room temperature, after which anisotropy values were measured using a BioTek Synergy H1 microplate reader (excitation at 485 nm, emission at 528 nm). The signal plotted is the anisotropy value with compound minus the anisotropy value without compound, giving the anisotropy change (ΔAU). *Compound titrations* were performed to follow-up the single point selectivity panel. A mastermix containing 10 nM of FAM-labeled peptides (C-RAF, SOS1, TAZ, KC1A, TSC2) and the corresponding 14-3-3σ concentration at $EC_{20}$ of the protein-peptide complex in buffer (10 mM HEPES, pH 7.4, 150 mM NaCl, 2 mM $MgCl_2$, 0.1% Tween20, 1 mg/mL BSA, 100 µM β-ME) was dispensed in assay plates. Compounds 22 and 23 were dispensed with the Tecan D300e digital dispenser in a 3-fold dilution series (starting at 250 µM). Plates were incubated overnight at room temperature, after which anisotropy values were measured using a BioTek Synergy H1 microplate reader (excitation at 485 nm, emission at 528 nm). *Protein titrations* were performed by titrating 14-3-3σFL protein in a 2-fold dilution series (starting at 300 µM) to a mix of 10 nM FAM-labeled peptides (CRAF, SOS1, TAZ, TSC-2) and 100 µM of compounds 22 and 23, or DMSO in buffer (10 mM HEPES, pH 7.4, 150 mM NaCl, 0.1% Tween20, 1 mg/mL BSA). After overnight incubation, anisotropy values were measured using a Tecan Infinite F500 plate reader (filter set excitation: 485±20 nm, emission: 535±25 nm, mirror: Dichroic 510, flashes: 20, integration time: 50 ms, settle time: 0 ms, gain: optimal).

**SURFACE PLASMA RESONANCE**

The SPR experiments were performed at 25°C using a Biacore X100 and a 200 nm Strep-Tactin XT derivatized linear polycarboxylate hydrogel chip, medium charge density (XanTec Bioanalytics). All proteins and peptides were dissolved in fresh running buffer prepared with ultrapure water and filtered through a 0.2 µm filter (10 mM HEPES pH 7.4, 200 mM NaCl, 50 µM EDTA, 0.005% P20). First the surface was conditioned with a 1 min injection of 3 M Guanidine HCl. Then, the recombinant 14-3-3σ-Twinstrep protein (250 nM) was captured on flow cell 2 of the sensor chip at a flow rate of 10 µL/min for 2 minutes, which resulted in a capture level of 1000 RU. For the ternary interaction, 14-3-3σ-Twinstrep protein (250 nM) was first incubated overnight with 1 µM Ac-C-RAF peptide and 20 µM compound prior to immobilization to the chip. The bound C-RAF peptide was washed away using running buffer flowed over the chip for 15 min. Flow cell 1 was left blank as a reference surface. After immobilization of the protein, the Biacore X100 was primed with running buffer. Multi-cycle kinetic measurements were conducted at a flow rate of 30 µL/min. A 2-fold dilution series of analyte (Ac-A-, B-, C-RAF peptides) in running buffer were injected over de sensor chip for 2 min, followed by dissociation of 3 min (binary interaction), 15 minutes (ternary interaction). For the binary interaction, the highest concentration of the RAF peptides was 50 µM, and for the ternary interactions this was 250 nM. Between cycles of one multi-cycle measurement, no regeneration step was performed due to complete dissociation of the analyte. After a measurement, the chip was regenerated by 2 times 30 sec injections of 3 M Guanidine HCl. The data was corrected by double subtracting to the reference surface (flow cell 1) and buffer injection and analyzed using 1:1 interaction fitting model with the BIA evaluation software (2020).

**X-RAY CRYSTALLOGRAPHY DATA COLLECTION AND REFINEMENT**

The 14-3-3σΔC protein, Ac-C-RAF/B-RAF/A-RAF and compounds (50 mM stock in DMSO) were dissolved in complexation buffer (25 mM HEPES pH=7.5, 2 mM $MgCl_2$ and 100 µM TCEP) and mixed in a 1:2:3 or 1:3:5 molecular stoichiometry (protein : peptide : compound) with a final protein concentration of 11 or 12 mg/mL. The complex was set-up for sitting-drop crystallization after overnight incubation at 4 °C, in a custom crystallization liquor (0.05 M HEPES (pH 7.1, 7.3, 7.5, 7.7), 0.19 M $CaCl_2$ 24-29% PEG400, and 5% (v/v) glycerol). Crystals grew within 10-14 days at 4 °C. Crystals were fished and flash-cooled in liquid nitrogen. X-ray diffraction (XRD) data were collected at the European Synchrotron Radiation Facility (ESRF Grenoble, France, beamline ID23-1, ID30A-3/MASSIF-3, or ID23-2).

Initial data processing was performed at ESRF using autoPROC after which pre-processed data was taken towards further scaling steps, molecular replacement, and refinement.

Data was processed using CCP4i2 suite (version 8.0.003). After indexing and integrating the data, scaling was done using AIMLESS. The data was phased with MolRep, using PDB 3IQU as template. The presence of co-crystallized ligands was verified by visual inspection of the Fo-Fc and 2Fo-Fc electron density maps in COOT (version 0.9.6). If electron density corresponding to the co-crystallized ligand was present, its structure and restraints were generated using AceDRG. After building in the ligand, model rebuilding and refinement was performed using REFMAC5. The PDB REDO server (pdb-redo.edu) or phenix.refine form Phenix software suite (version 1.19.2-4158) was used to complete the model building and refinement. The images were created using the PyMOL Molecular Graphics System (Schrödinger LLC, version 4.6.0). See SI table S11 for data collection and refinement statistics.

**NanoBRET ASSAYS**

NanoBRET assays were performed as previously described.[2] Briefly, eight constructs were generated, and combination and ratio tests were performed in HEK293T cells for development of the initial 14-3-3σ/C-RAF NanoBRET. All NanoBRET assays were performed in HEK293T cells at a 1:10 transfection ratio of NanoLuc:HaloTag plasmids. Compounds were tested in a 2-fold dilution series starting at 10 µM with C-RAF-NanoLuc (C-terminal tag)/14-3-3-HaloTag (C-terminal tag). 14-3-3σ/A-RAF and 14-3-3σ/B-RAF NanoBRET were performed with the same conditions and tag placements. N-RAS/C-RAF NanoBRET was performed with C-RAF-NanoLuc (C-terminal tag)/HaloTag-NRAS Q61L (N-terminal tag) with HA-14-3-3σ transfected in addition at a 1:1 ratio with C-RAF-NanoLuc. C-RAF/C-RAF NanoBRET used C-RAF-NanoLuc/HaloTag-C-RAF with HA-14-3-3σ and FLAG-N-RAS Q61L at a 1:1 ratio with C-RAF-NanoLuc.

Data was analyzed as previously described.[2] Briefly, milliBRET units (mBU) were calculated as a ratio of acceptor/donor and mean mBU values were corrected by subtracting the mean mBU of controls with no HaloTag ligand. Fold change was calculated by dividing $mBU_{compound}$ by $mBU_{DMSO\ control}$. For N-RAS/C-RAF NanoBRET, 100% was considered the DMSO mBU and 0% the mBU value of NRAS/C-RAF R89L.

**CELL CULTURE AND WESTERN BLOTS**

HEK293T (CRL-3216) and MIA PaCa-2 (CRL-1420) cells were purchased from ATCC. Both cell lines were grown in DMEM, high glucose (Gibco) with 10% Fetal Bovine Serum (FBS). All compound treatments were 24 hours. Anti-phospho-C-RAF pS259 (Invitrogen 44-502), anti-C-RAF (BD Biosciences 610152), anti-phospho-A-RAF pS214 (Thermo-Fischer BS-3005R), anti-A-RAF pS214 (OriGene Technologies TA803662), anti-phospho-B-RAF pS365 (Thermo-Fischer BS-3013R), anti-phospho-ERK (Cell Signaling Technology 4370S), anti-ERK (Santa Cruz Biotechnology sc-514302), anti-Vinculin (Cell Signaling Technology 13901S), anti-FLAG (Cell Signaling Technology 8146S), and anti-14-3-3σ (Invitrogen SD2070) antibodies were used for protein detection. Antibodies were used at a 1:1,000 dilution except anti-Vinculin, which was used at 1:2,000. Western blots were imaged on a LI-COR imaging system and analyzed using Image Studio.

**CO-IMMUNOPRECIPITATION**

HEK293T cells were dosed with 10 µM compound for 24 hours before lysis (Triton X-100). Clarified lysate was incubated with Pierce anti-FLAG magnetic beads (Thermo Fisher A36797) following the manufacturers protocol. Lysate and IP samples were analyzed using antibodies described above.

**LUMIT IMMUNOASSAY**

Lumit immunoassays were performed according to the manufacturer (Promega W1331). Briefly, HEK293T cells were transfected in a 1:1 ratio of HA-14-3-3σ:FLAG-C-RAF in 10% FBS DMEM (Gibco). 24 hours after transfection, cells were re-seeded in 384-well plates (10,000 cells per well) in 0.5% FBS FluoroBrite DMEM media (Gibco). Cells were treated with compound **23** in a 2-fold dilution series, starting at 10 µM, or LY3009120 (100 nM; Selleck Chemicals S7842) or Trametinib (100 nM; Selleck chemicals S2673) for 24 hours before performing the Lumit protocol. After compound treatment, media was replaced with 10 µL media supplemented with fluorogenic live cell GF-AFC substrate (50 µM) for normalization purposes and 100 ng/mL EGF to stimulate the MAPK pathway. Cells were incubated for 30 minutes before proceeding. The protocol was optimized for the antibodies used: digitonin (0.1%) lysis buffer, 100 ng/µL pERK antibody (Cell Signaling Technology 4376), 100 ng/µL ERK antibody (Santa Cruz Biotechnology 514302), and 0.1 µL/well of secondary antibodies (SmBiT mouse, LgBiT rabbit). The ratio of cell lysate:antibody:substrate was 2:2:1.

Cells were lysed for 40 minutes at 800 rpm before incubation with the antibody mix for 90 minutes (2 minutes at 400 rpm prior to incubation). After antibody incubation, substrate was added and incubated for 2 minutes at 400 rpm. Luminescence was then read on an EnVision Xcite 2105 plate reader. Luminescence was normalized against the total fluorescence from the GF-AFC (Luminescence/Total fluorescence). Values were then normalized to DMSO treated cells (experimental/DMSO). Data was analyzed using GraphPad Prism and fit using a 4PL model. Data represents the average of 3 technical replicates, confirmed over 3 biological replicates.

**DOCKING**
Computational design for SAR optimization and docking was performed with SeeSAR version 11.2.0; BioSolveIT GmbH, Sankt Augustin, Germany, 2022, www.biosolveit.de/SeeSAR

**SOFTWARE VERSIONS**
Prism (10.2.1), Illustrator (22.1 (64-bit)), Biorender (64-bit), Pymol (4.6.0), CCP4i2 (8.0.003), COOT (0.9.8.1), Phenix (1.19.2-4158)

**DATA AVAILABILITY** Supplementary figures and tables, synthetic procedures, compound characterization, NMR spectra, crystallography data (PDF). The structures were deposited in the protein data bank (PDB) with IDs: 8Q5C, 8Q55, 8QS4, 8QS3, 9EW5, 8QS2, 8QS5, 8QS6, 8QS7, 8QS8, 9EW3, 9EW1, 8S42, 8QS9, 8QSA, 9EW4, 9EW7, 9EW6, 8VSL, 8QSC, 8QSH, 8VSM, 8QSB, 8QSF, 8QSE, 8VSO, 8QSD, 8QSG.

### 2. SUPPLEMENTARY FIGURES

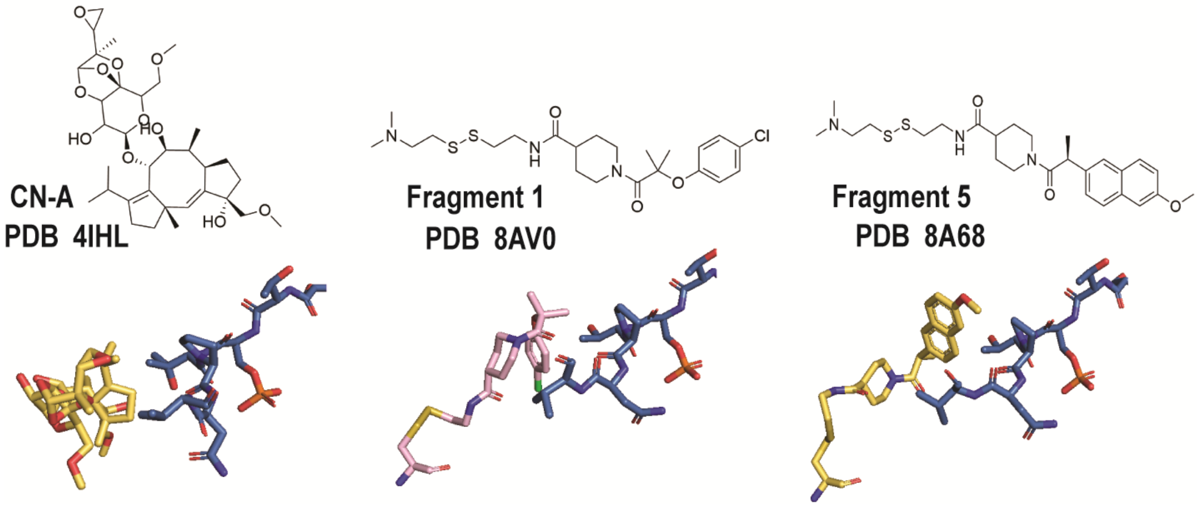

**Figure S1.** Crystal structures of known molecular glues bound to 14-3-3/C-RAF259 (14-3-3 is omitted for clarity, C38 is shown as yellow sticks, and CRAF as blue sticks.

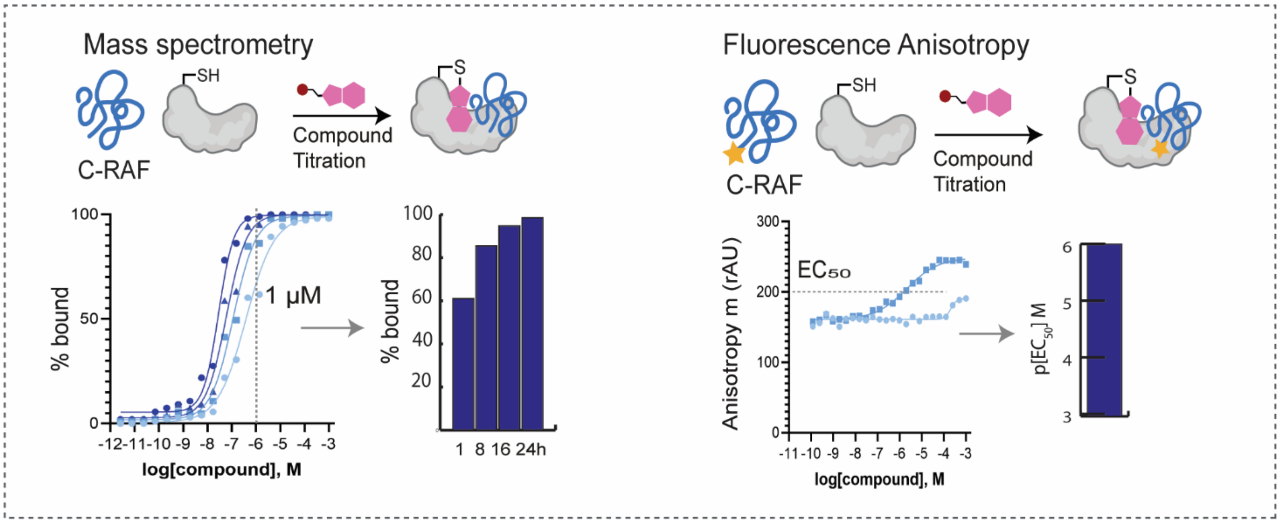

**Figure S2.** Overview of assays. Left: mass spectrometry time-course experiments were performed at 1h, 8h, 16h and 24h in the presence of C-RAF259 phosphopeptide. Bar graphs were used to represent % bound at 1 μM compound concentration (1:10 [protein]:[compound] ratio).Right: fluorescence anisotropy compound titrations were performed in the presence of FAM-labeled C-RAF259 phosphopeptide. $EC_{50}$ values were calculated from the overnight measurement. $pEC_{50}$ bar graphs represent the positive log $EC_{50}$ value.

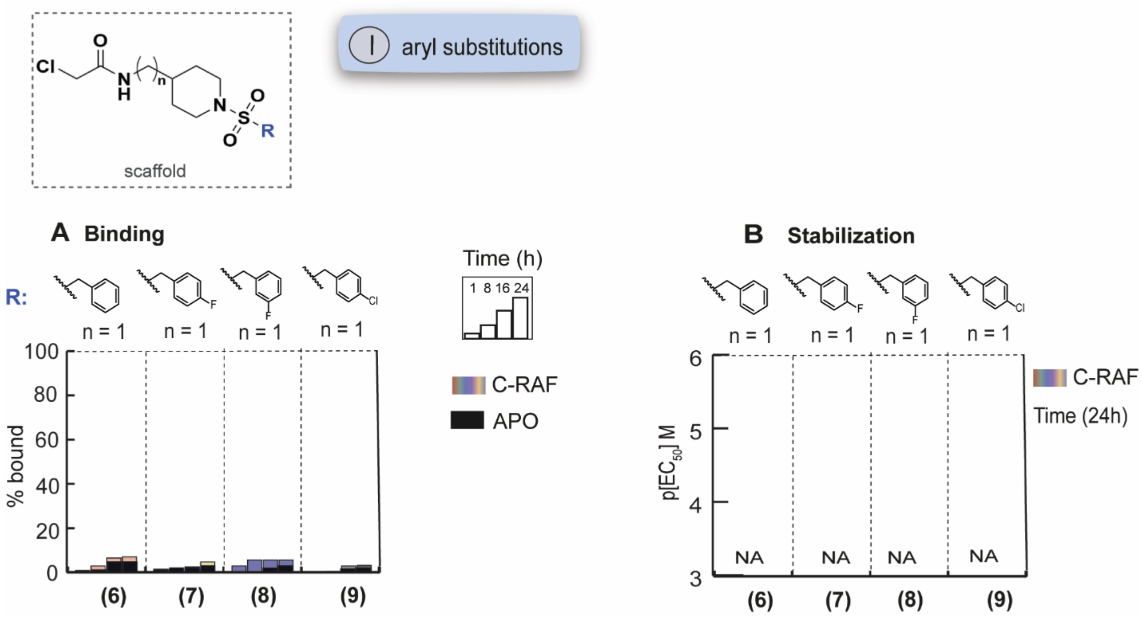

**Figure S3.** SAR of benzyl analogs; MS bar graphs (left) and FA bar graphs (right).

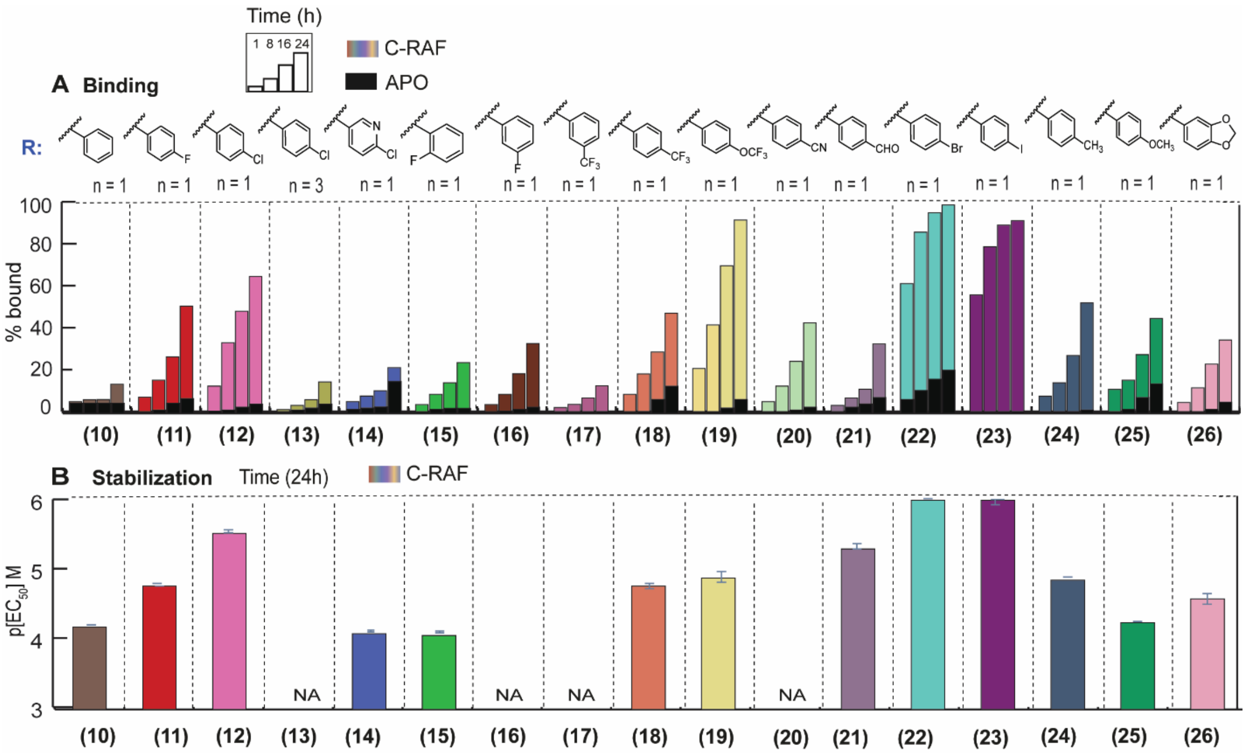

**Figure S4.** SAR of phenyl analogs. Top: MS bar graphs sat 1 μM. For each compound, time course experiments were performed with measurements at 1h, 8h, 16h and 24h. C-RAF259 data are shown with different colors for each compound, and apo data in black. Bottom: Bar graphs of FA compound titration $pEC_{50}$ values after overnight incubation. C-RAF259 data are shown with different colors for each compound. Inactive compounds are described as non-applicable (NA).

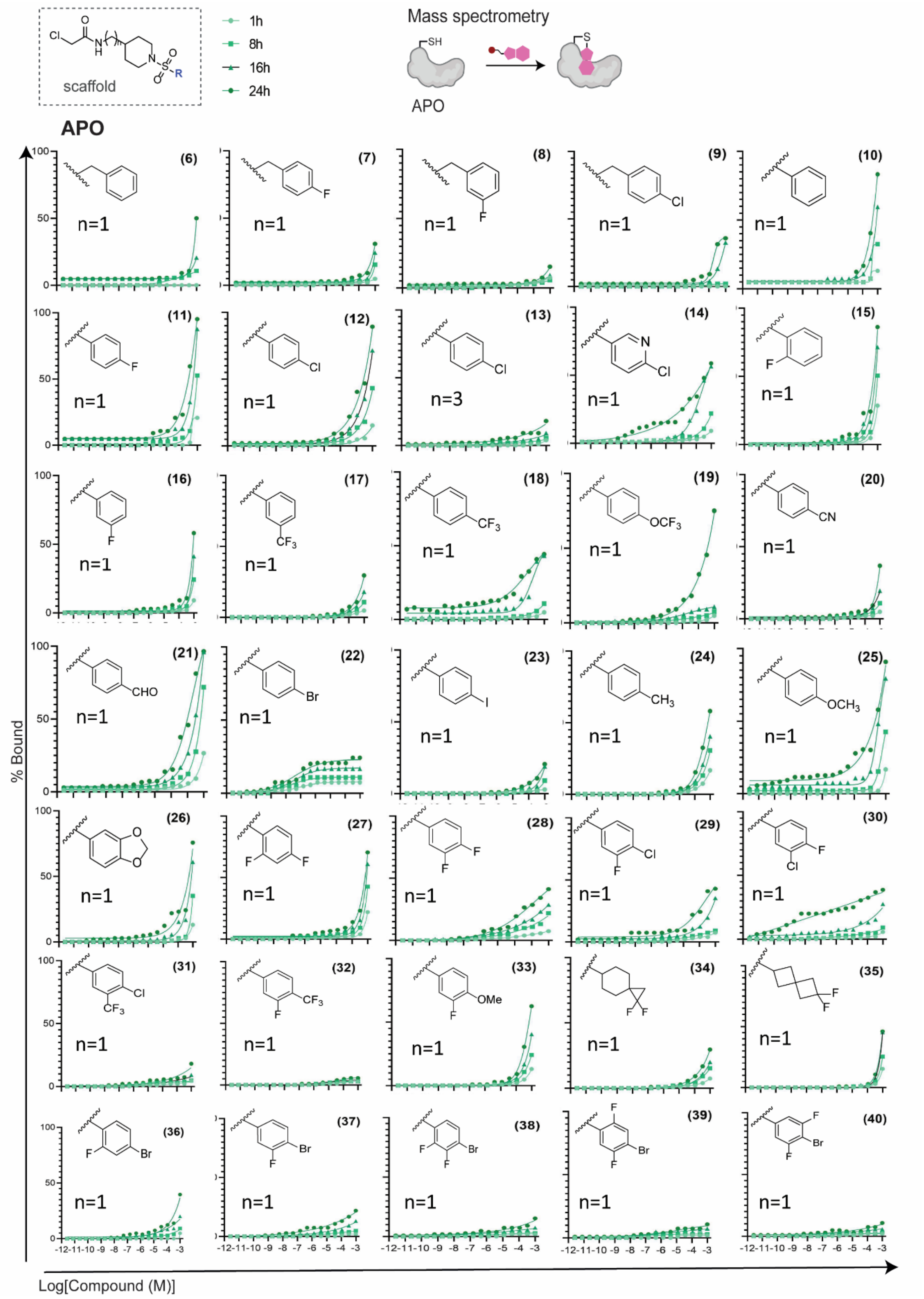

**Figure S5.** MS dose-response curves (apo) for aryl-ring modifications.

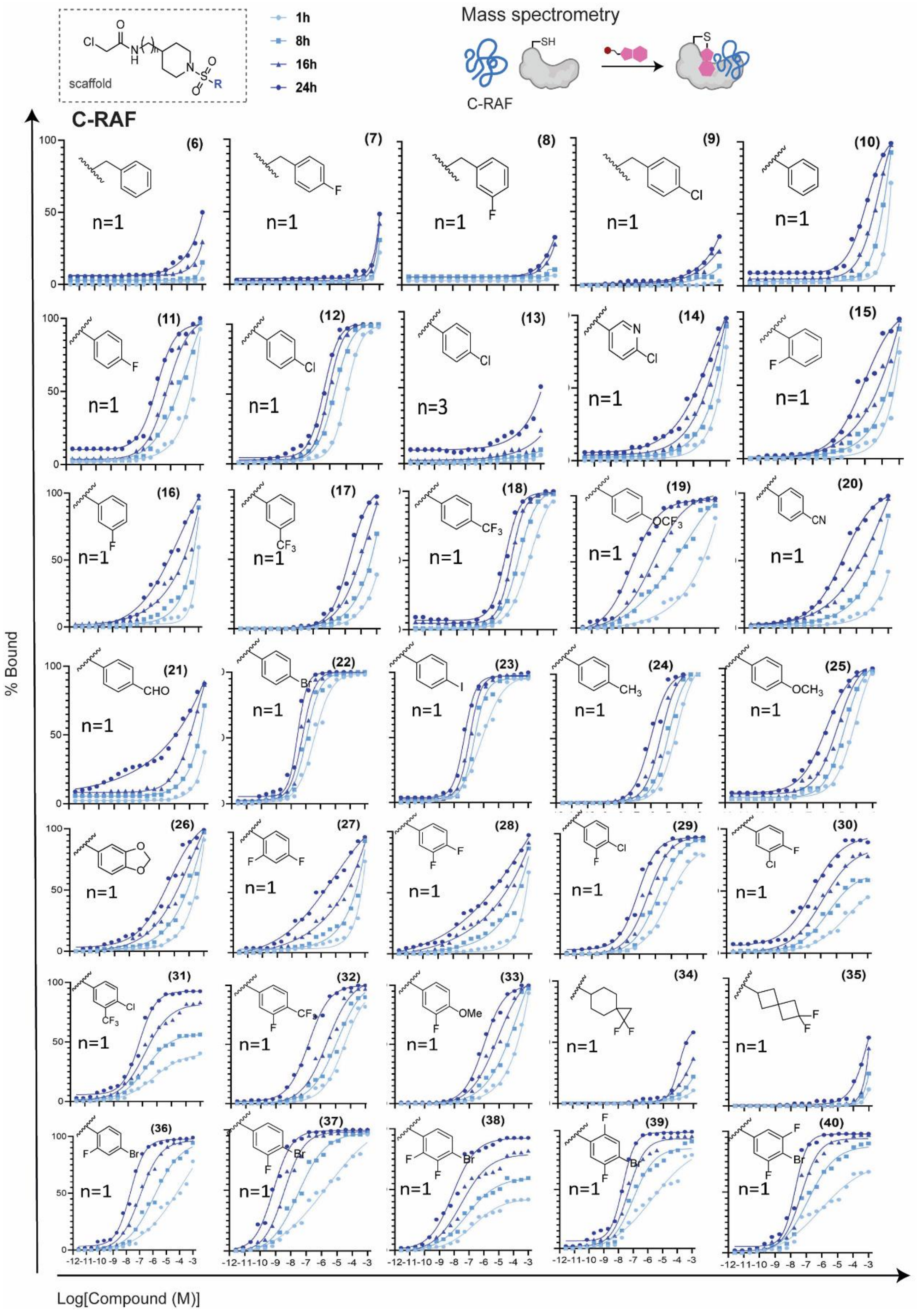

**Figure S6.** MS dose-response curves (C-RAF259) for aryl-ring modifications.

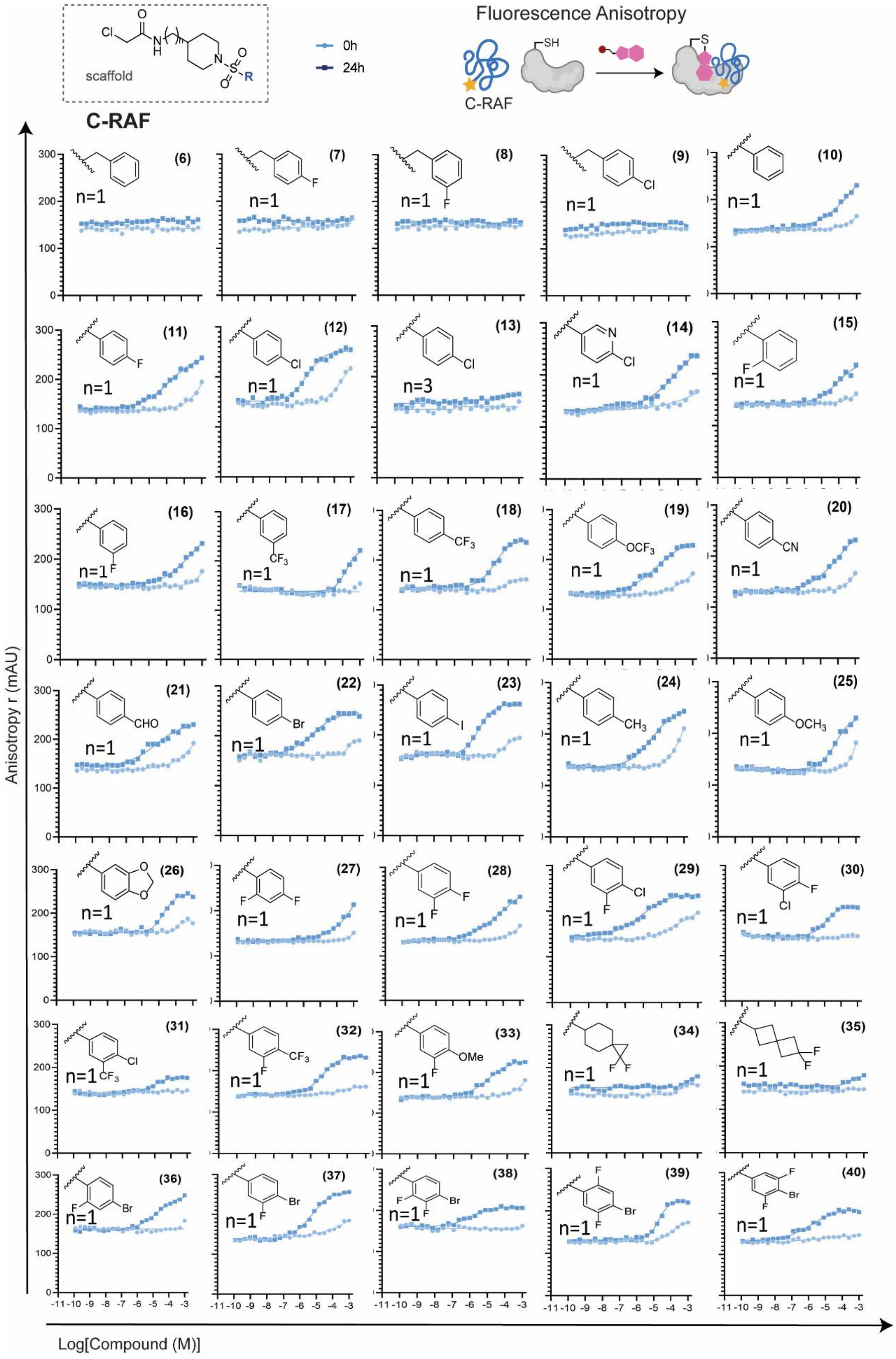

**Figure S7.** FA compound titrations (C-RAF259) for aryl-ring modifications.

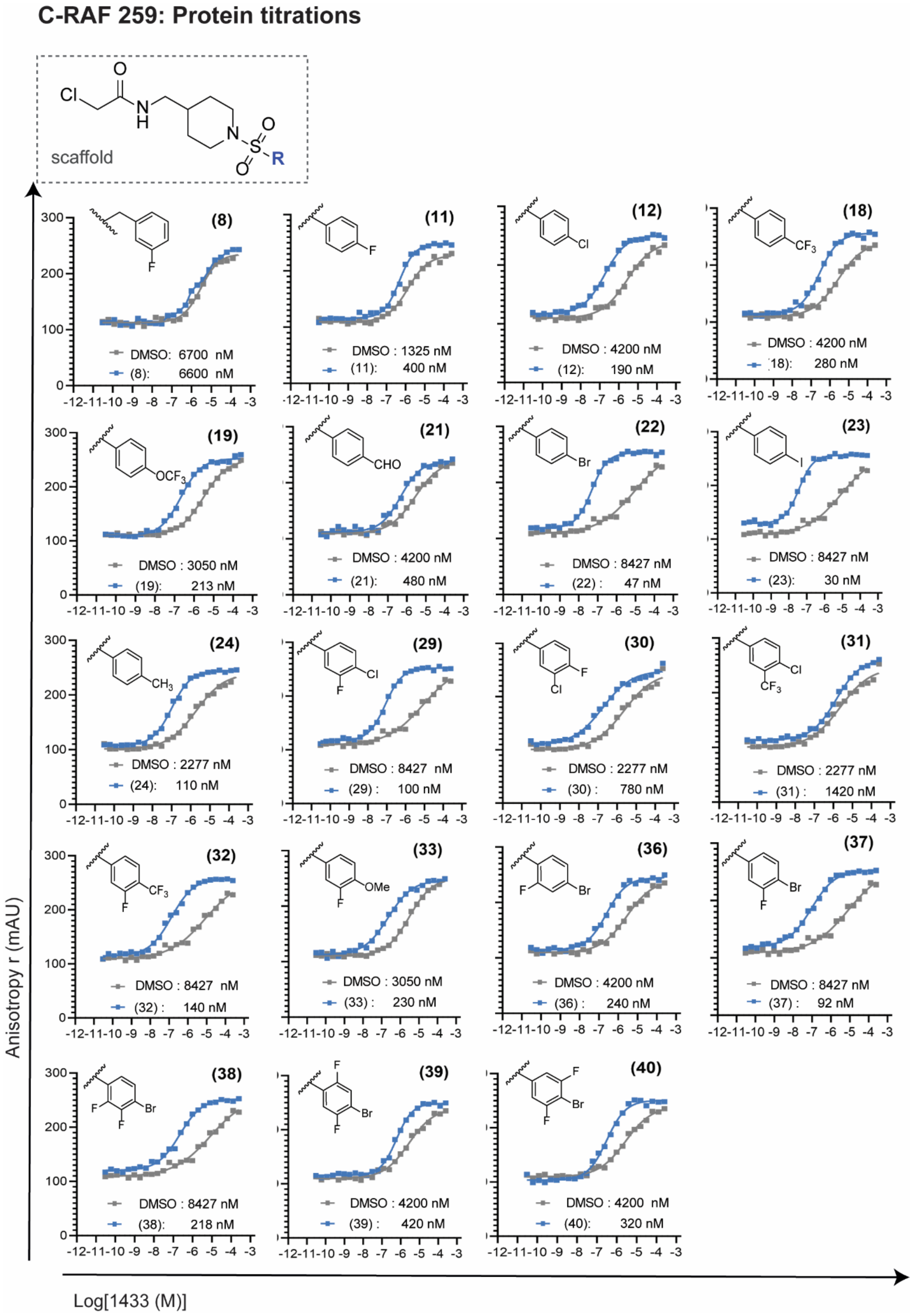

**Figure S8.** FA protein titrations (C-RAF259) for aryl-ring modifications.

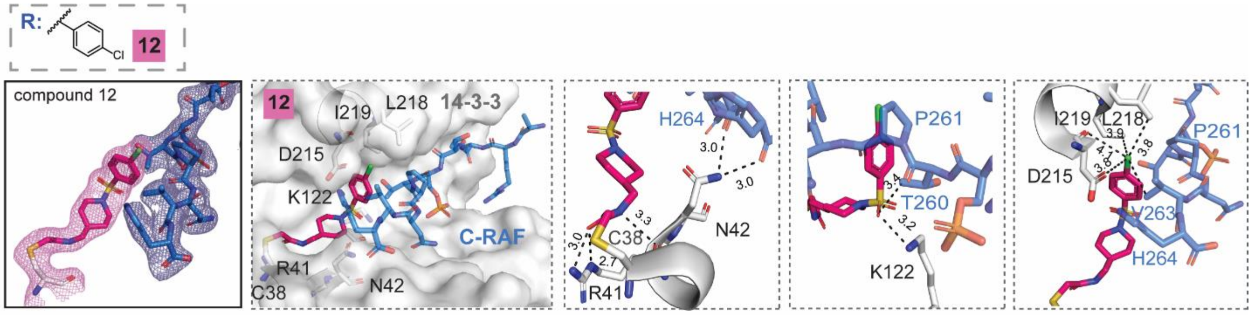

**Figure S9.** Density and crystal structure of **12** (magenta) in complex with C-RAF pS259 10-mer peptide and 14-3-3σ (white). Close-up views on interacting aminoacids. 2Fo-Fc electron density maps (mesh) were contoured at 1σ.

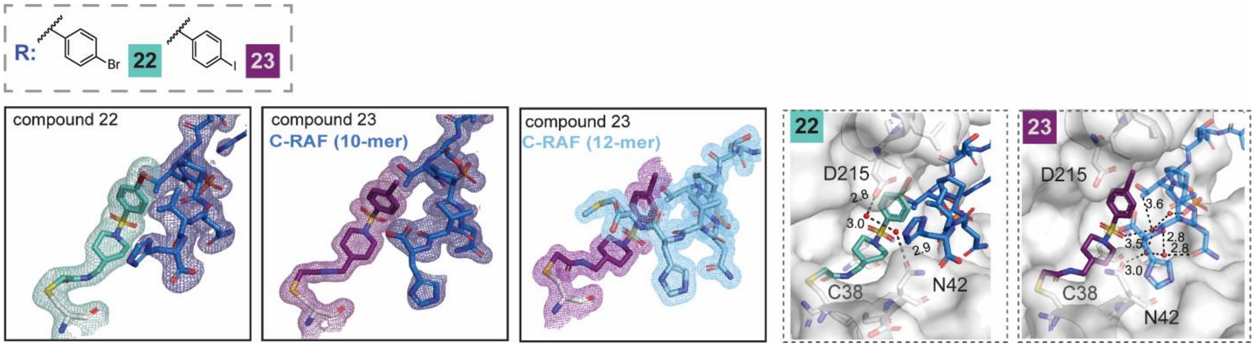

**Figure S10.** Density and crystal structures of **22** (turquoise) and **23** (purple) in complex with C-RAF pS259 10-mer peptide and 14-3-3σ. Peptide lengths for **23** as indicated in the figure. Close-up views on interacting aminoacids. Interacting water molecules are shown as red spheres. 2Fo-Fc electron density maps (mesh) were contoured at 1σ.

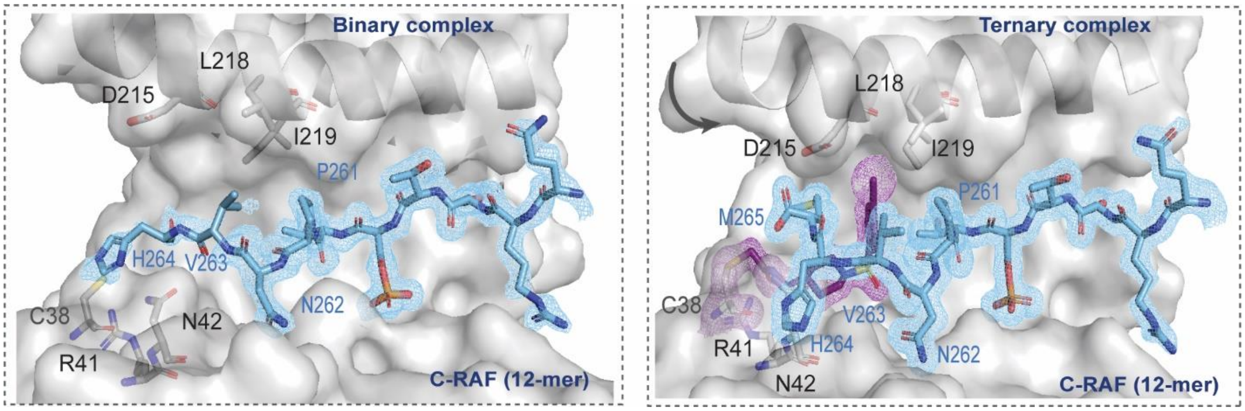

**Figure S11.** Left: Binary complex of 14-3-3σ with C-RAF pS259 12-mer peptide. Right: ternary complex of **23** (purple)/14-3-3σ/ C-RAF pS259 12-mer peptide (cyan sticks). Movement of helix 9 is indicated by a gray arrow.

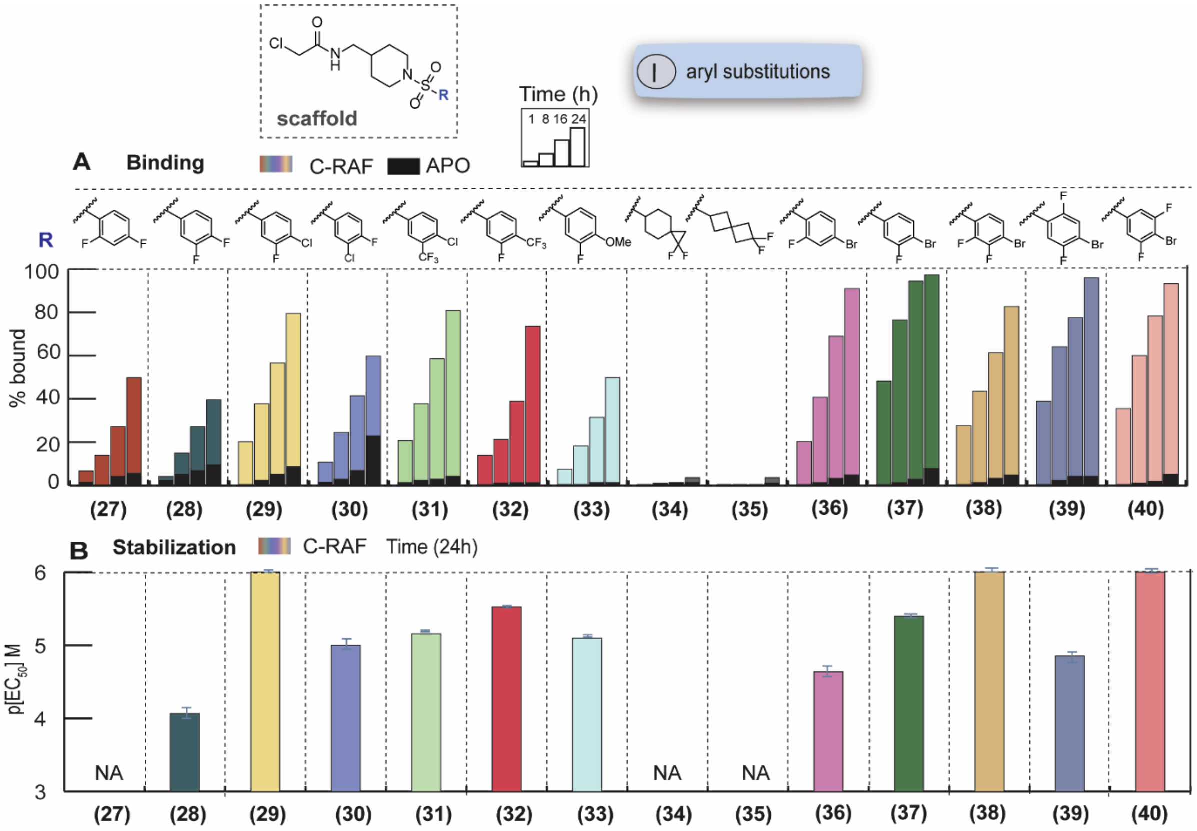

**Figure S12.** SAR of double- and triple- halogenated derivatives. Top: MS bar graphs sat 1 µM. For each compound, time course experiments were performed with measurements at 1h, 8h, 16h and 24h. C-RAF259 data are shown with different colors for each compound, and apo data in black. Bottom: Bar graphs of FA compound titration $pEC_{50}$ values after overnight incubation. C-RAF259 data are shown with different colors for each compound. Inactive compounds are described as non-applicable (NA).

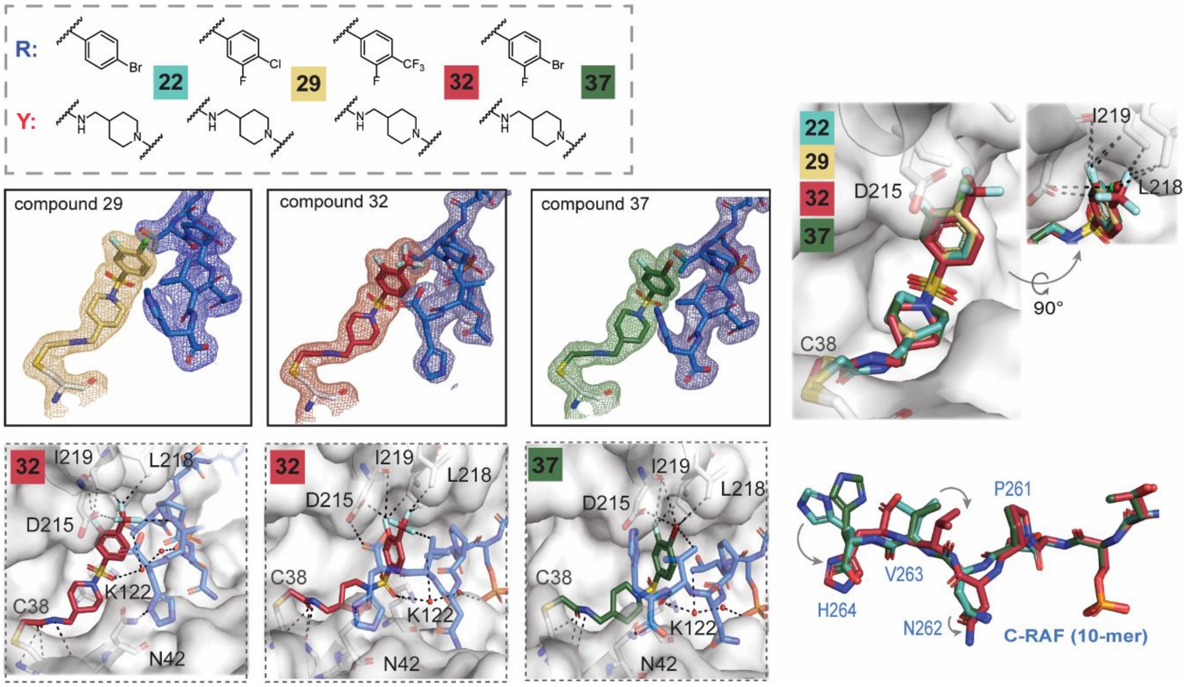

**Figure S13.** Density and crystal structures of **29** (yellow), **32** (orange) and **37** (green) in complex with C-RAF pS259 10-mer peptide and 14-3-3σ. Close-up views on interacting aminoacids and overlays. Interacting water molecules are shown as red spheres. 2Fo-Fc electron density maps (mesh) were contoured at 1σ.

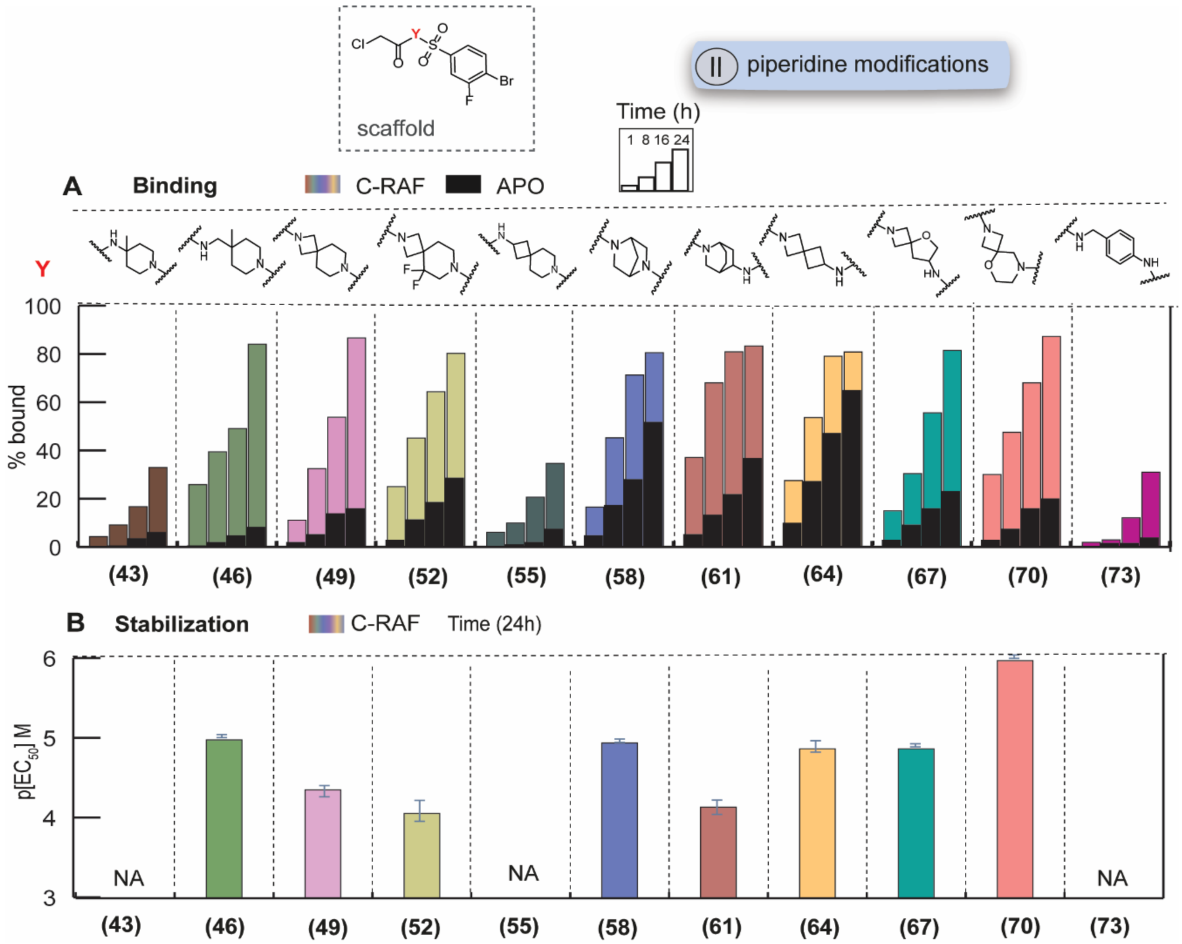

**Figure S14.** SAR of modifications on the piperidine position. Top: MS bar graphs sat 1 µM. For each compound, time course experiments were performed with measurements at 1h, 8h, 16h and 24h. C-RAF259 data are shown with different colors for each compound, and apo data in black. Bottom: Bar graphs of FA compound titration $pEC_{50}$ values after overnight incubation. C-RAF259 data are shown with different colors for each compound. Inactive compounds are described as non-applicable (NA).

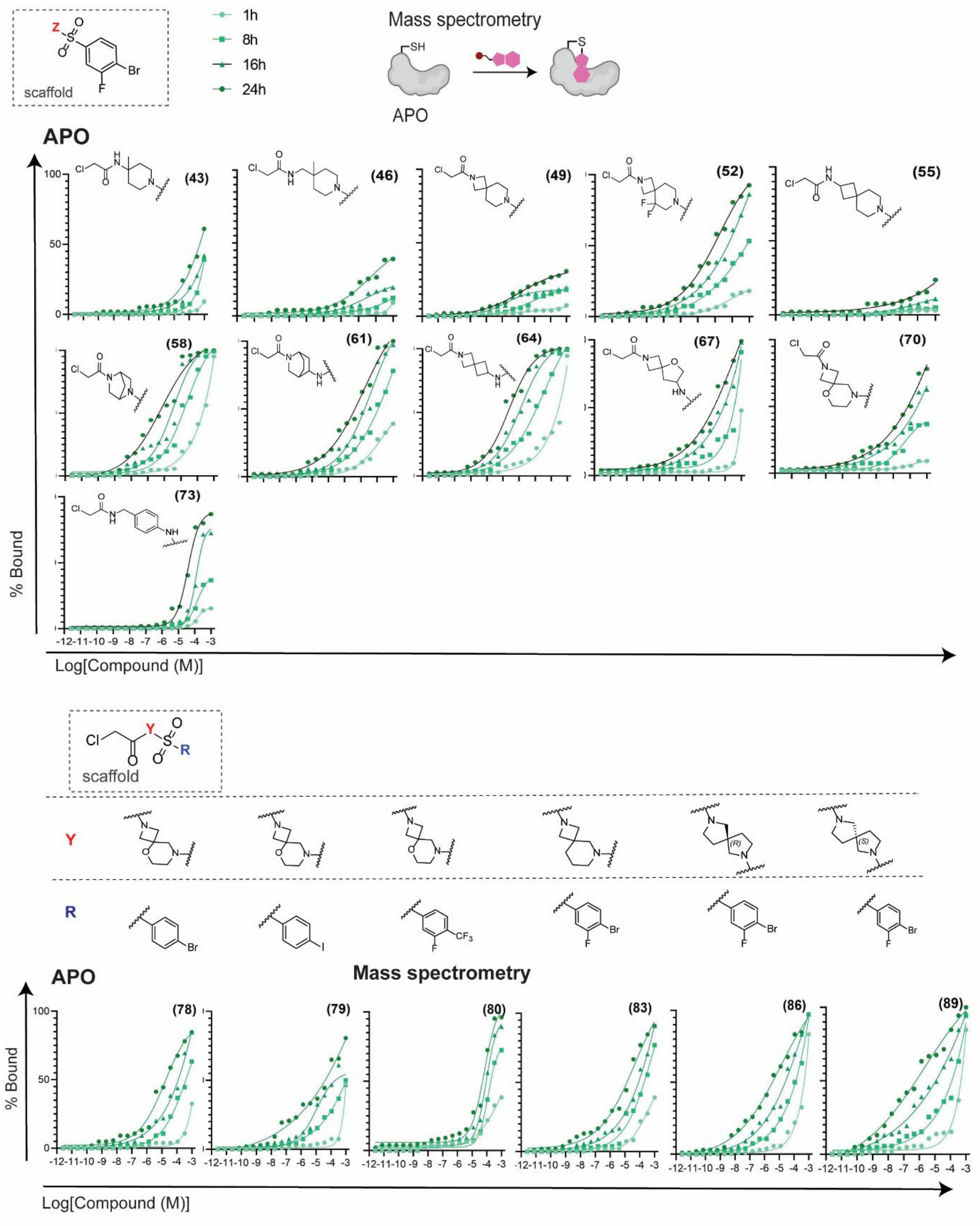

**Figure S15.** MS dose-response curves (apo) for piperidine modifications and combinations.

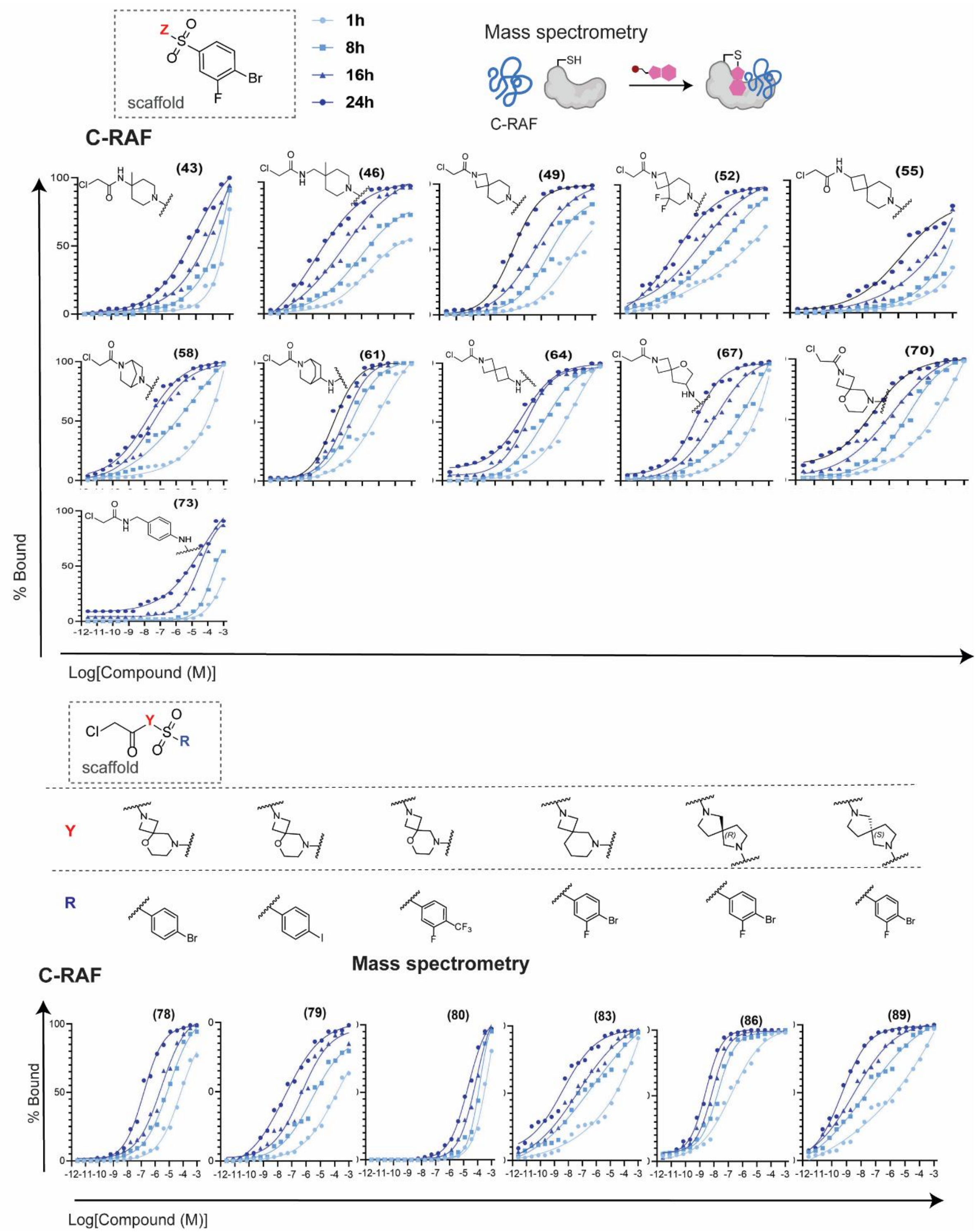

**Figure S16.** MS dose-response curves (C-RAF259) for piperidine modifications and combinations.

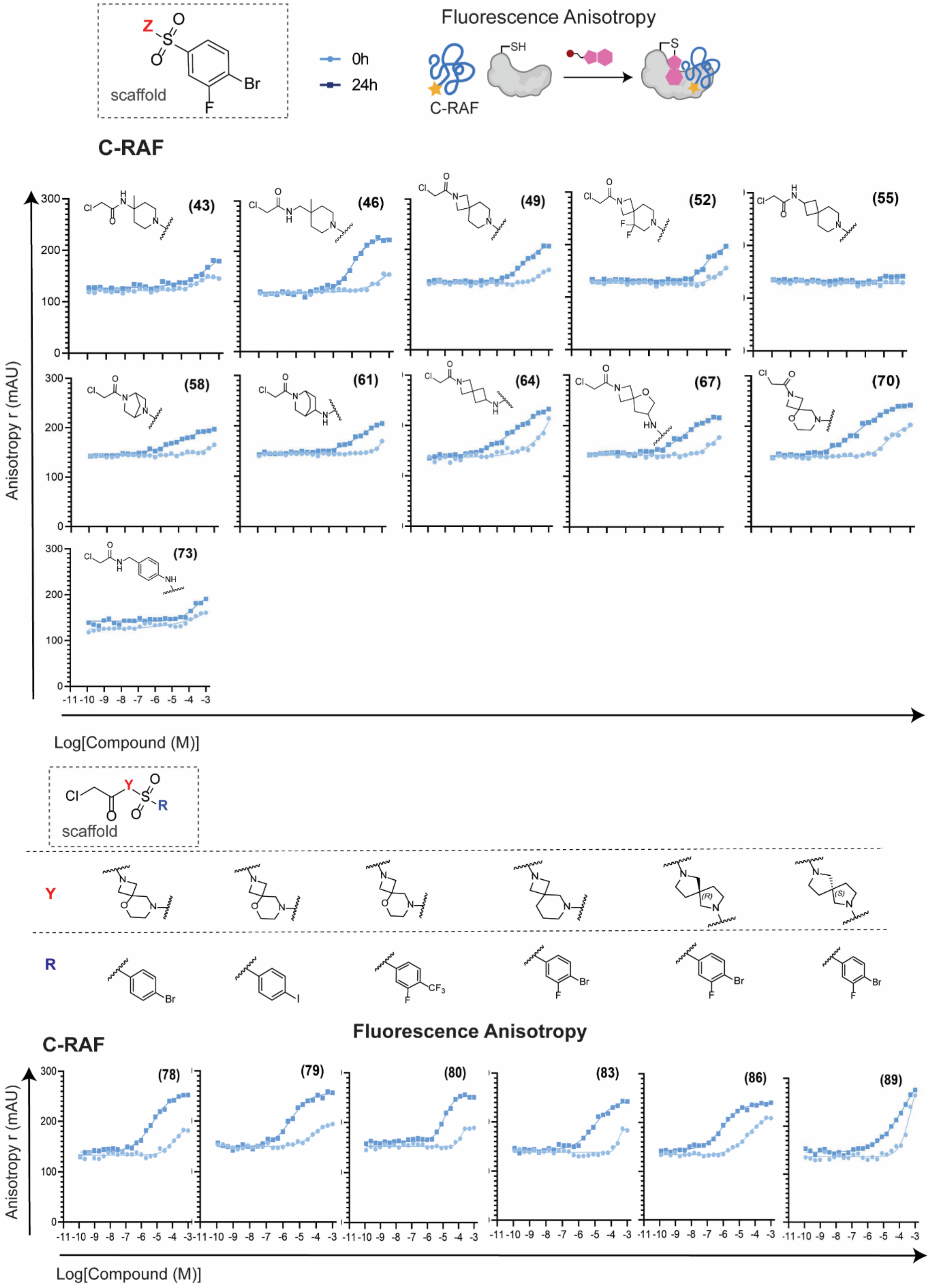

**Figure S17.** FA compound titrations (C-RAF259) for piperidine modifications and combinations.

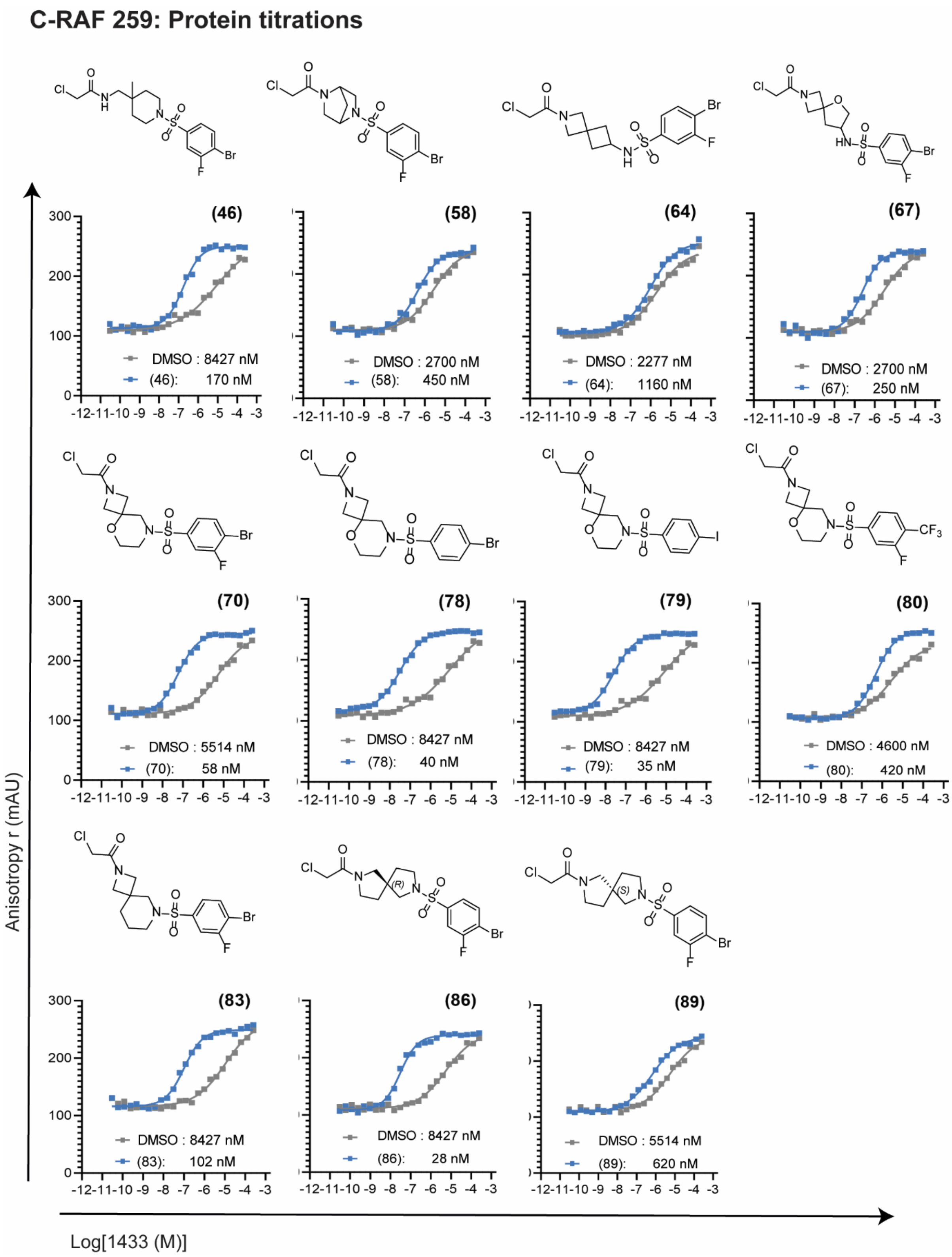

**Figure S18.** FA protein titrations (C-RAF259) for piperidine modifications and combinations.

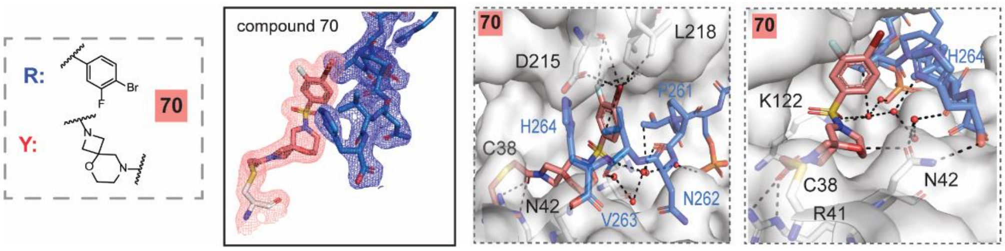

**Figure S19.** Density and crystal structure of **70** (salmon) in complex with C-RAF pS259 10-mer peptide and 14-3-3σ. Close-up views on interacting aminoacids Interacting water molecules are shown as red spheres. 2Fo-Fc electron density maps (mesh) were contoured at 1σ.

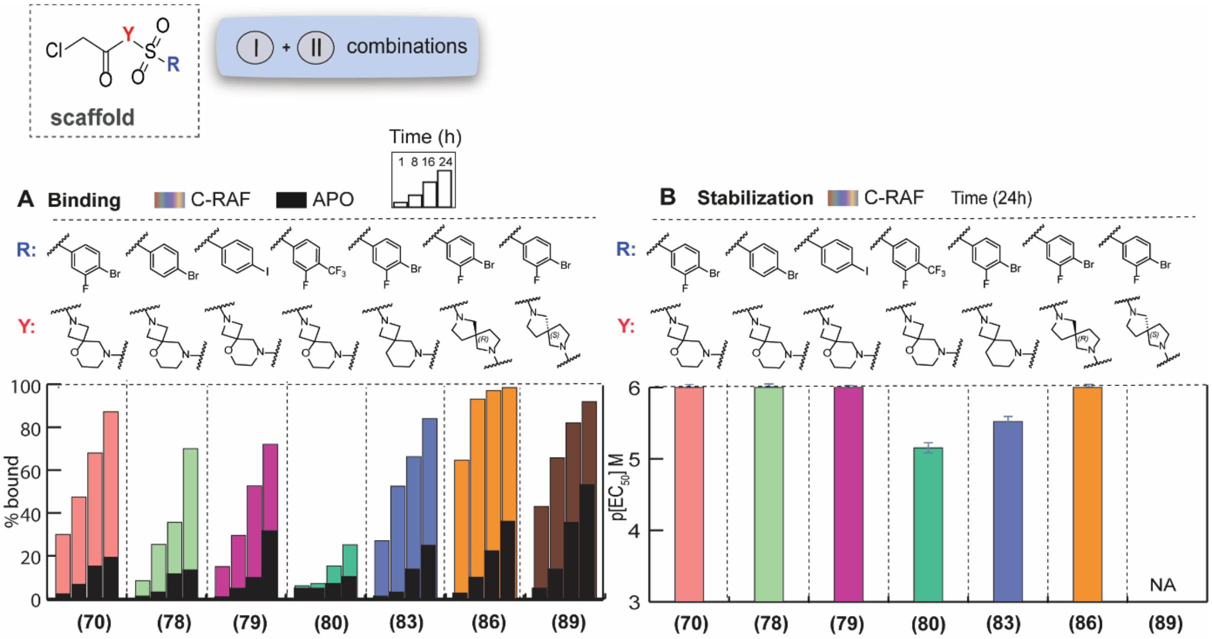

**Figure S20.** SAR of aryl and spiro/fused rings combinations. Left: MS bar graphs sat 1 µM. For each compound, time course experiments were performed with measurements at 1h, 8h, 16h and 24h. C-RAF259 data are shown with different colors for each compound, and apo data in black. Right: Bar graphs of FA compound titration $pEC_{50}$ values after overnight incubation. C-RAF259 data are shown with different colors for each compound. Inactive compounds are described as non-applicable (NA).

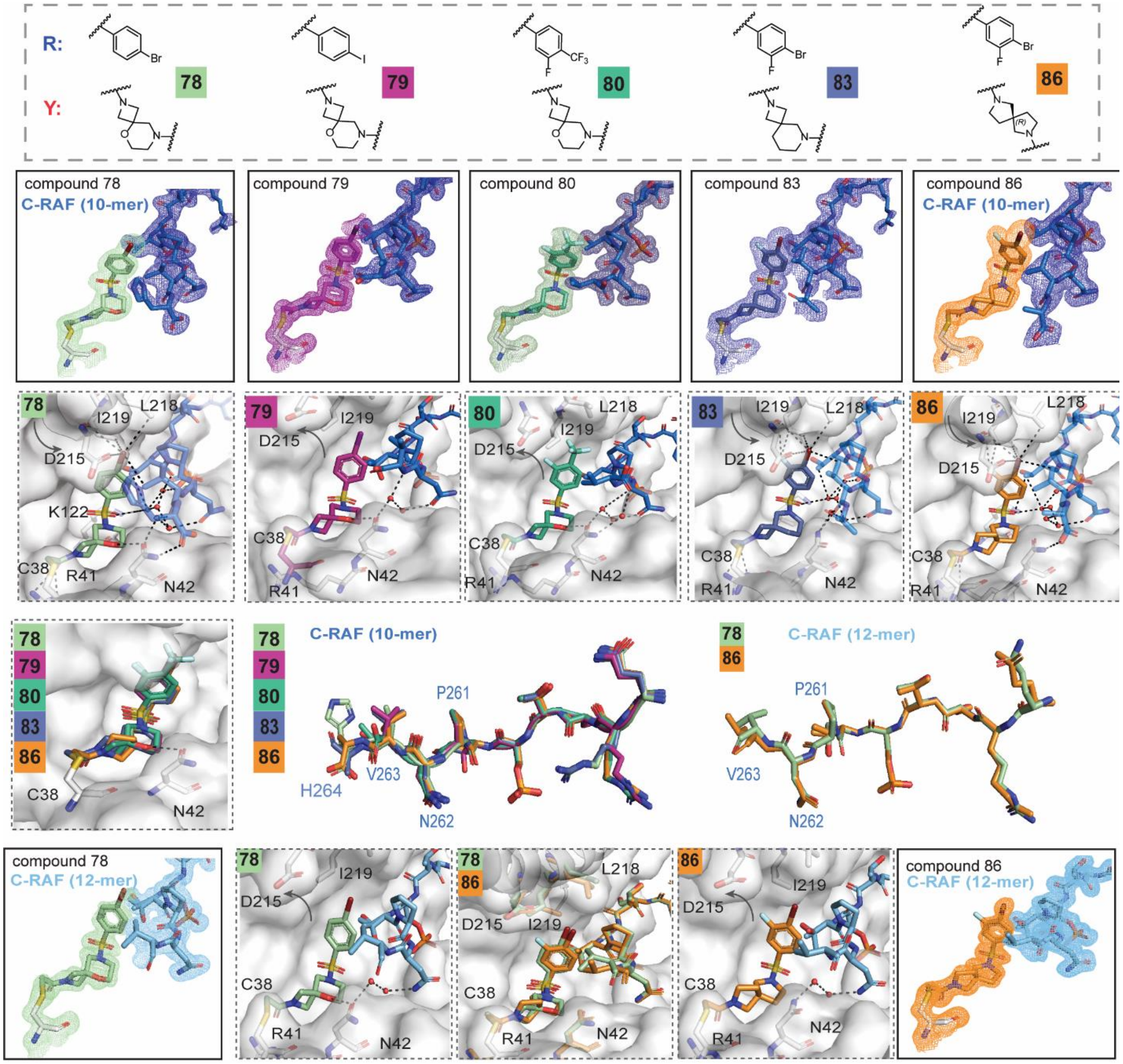

**Figure S21.** Top: Densities and crystal structures of **78** (light green sticks), **79** (lilac), **80** (teal), **83** (sky blue), **86** (orange) in complex with C-RAF pS259 10-mer peptide and 14-3-3σ. Close-up views on interacting aminoacids Interacting water molecules are shown as red spheres. 2Fo-Fc electron density maps (mesh) were contoured at 1σ. Bottom: overlays, crystal structures of **78** (light green sticks) and **86** (orange) in complex with C-RAF pS259 12-mer peptide and 14-3-3σ and close-up views of interacting aminoacids.

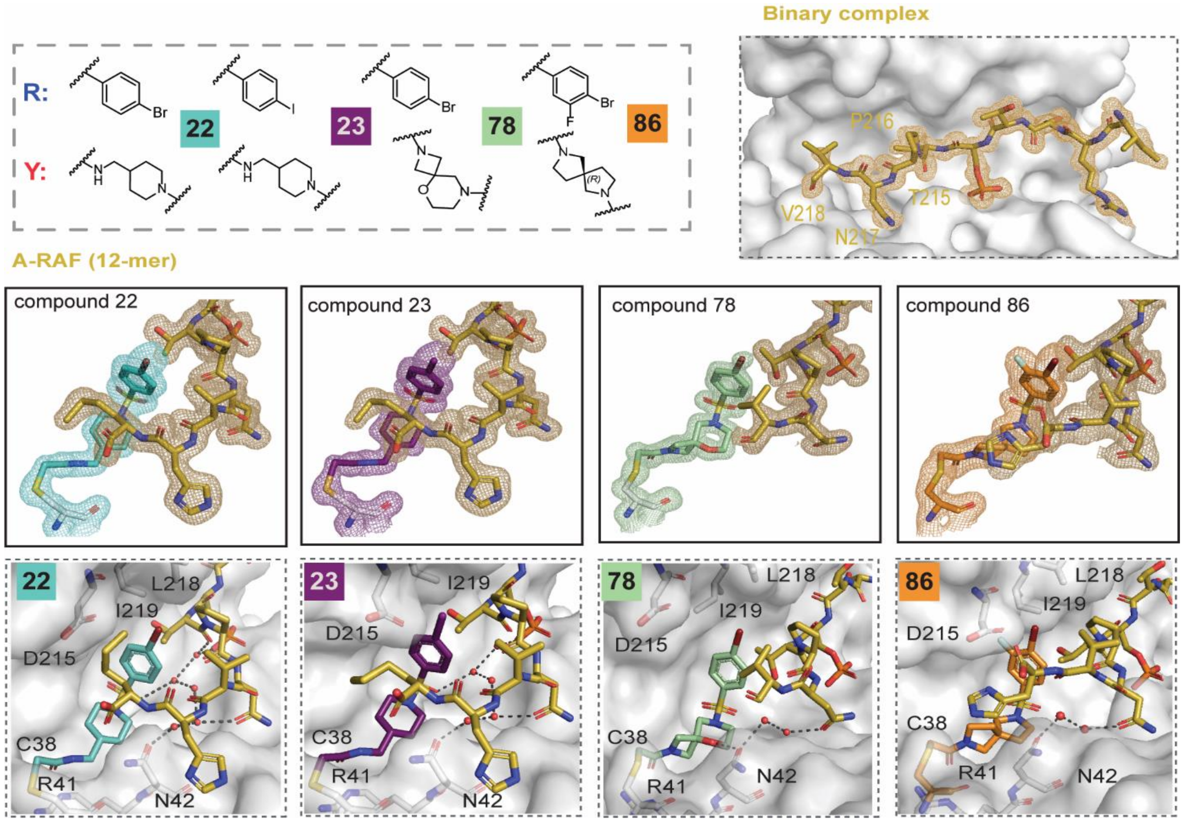

**Figure S22.** Densities and crystal structures with 14-3-3σ/A-RAF pS214 12-mer peptide. Top right: binary complex. Middle and bottom: ternary complexes of **22** (turquoise), **23** (purple), **78** (light green) and **86** (orange) with A-RAF pS214 12-mer peptide and 14-3-3σ. Close-up views on interacting aminoacids Interacting water molecules are shown as red spheres. 2Fo-Fc electron density maps (mesh) were contoured at 1σ.

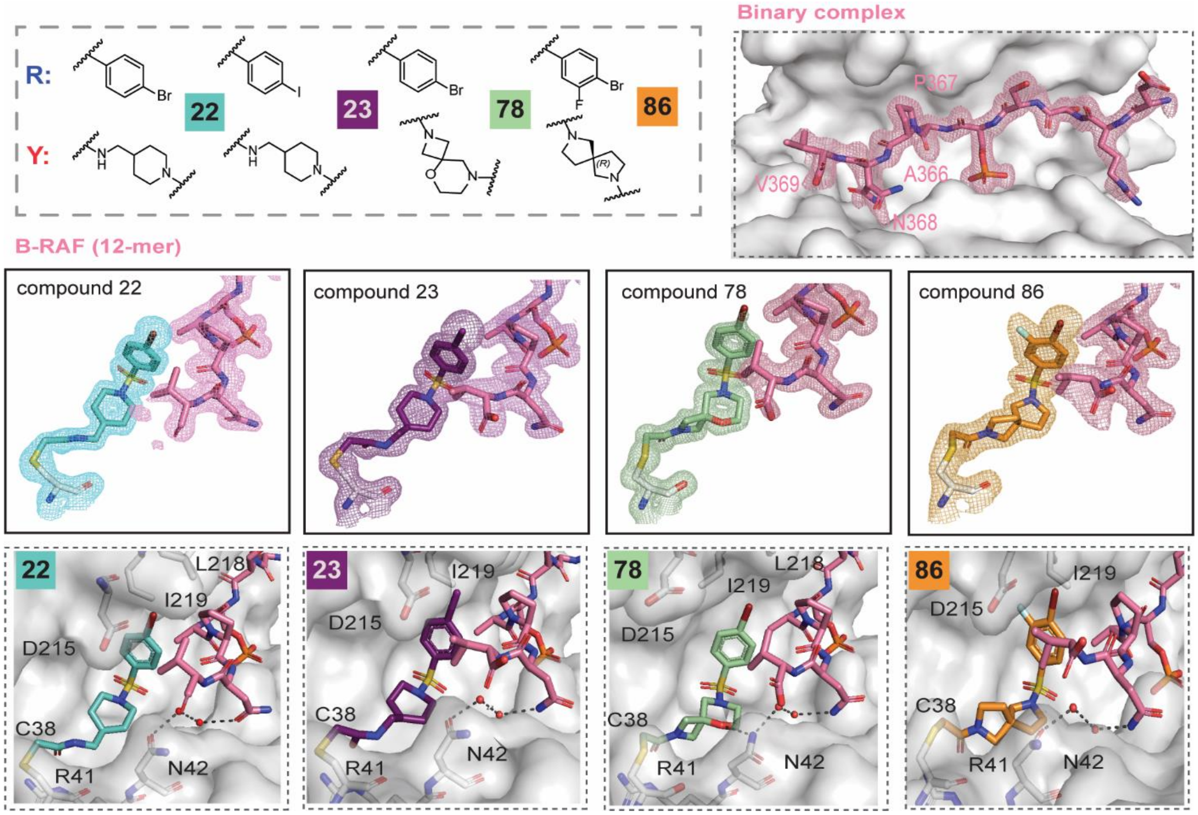

**Figure S23.** Densities and crystal structures with 14-3-3σ/B-RAF pS365 12-mer peptide. Top right: binary complex. Middle and bottom: ternary complexes of **22** (turquoise), **23** (purple), **78** (light green) and **86** (orange) with B-RAF pS365 12-mer peptide and 14-3-3σ. Close-up views on interacting aminoacids Interacting water molecules are shown as red spheres. 2Fo-Fc electron density maps (mesh) were contoured at 1σ.

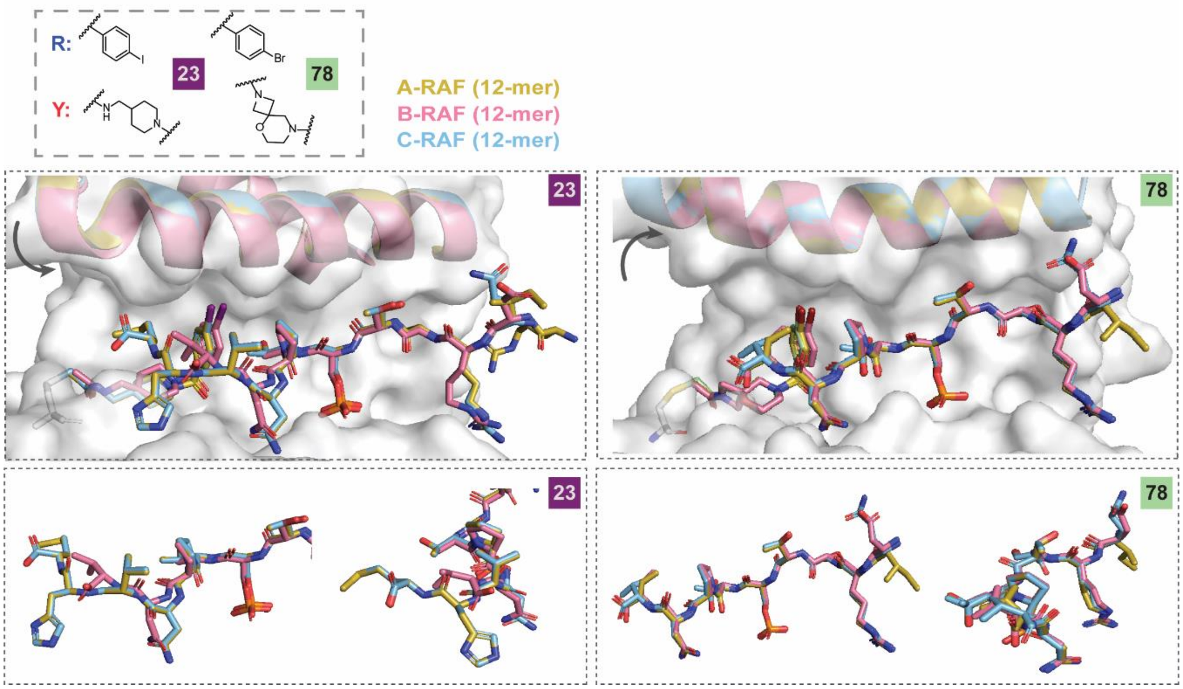

**Figure S24.** Overlays of ternary complexes for **23** (purple) and **78** (light green) with A-RAF pS214, B-RAF pS365 and C-RAF pS259. Close-up views on peptide conformations.

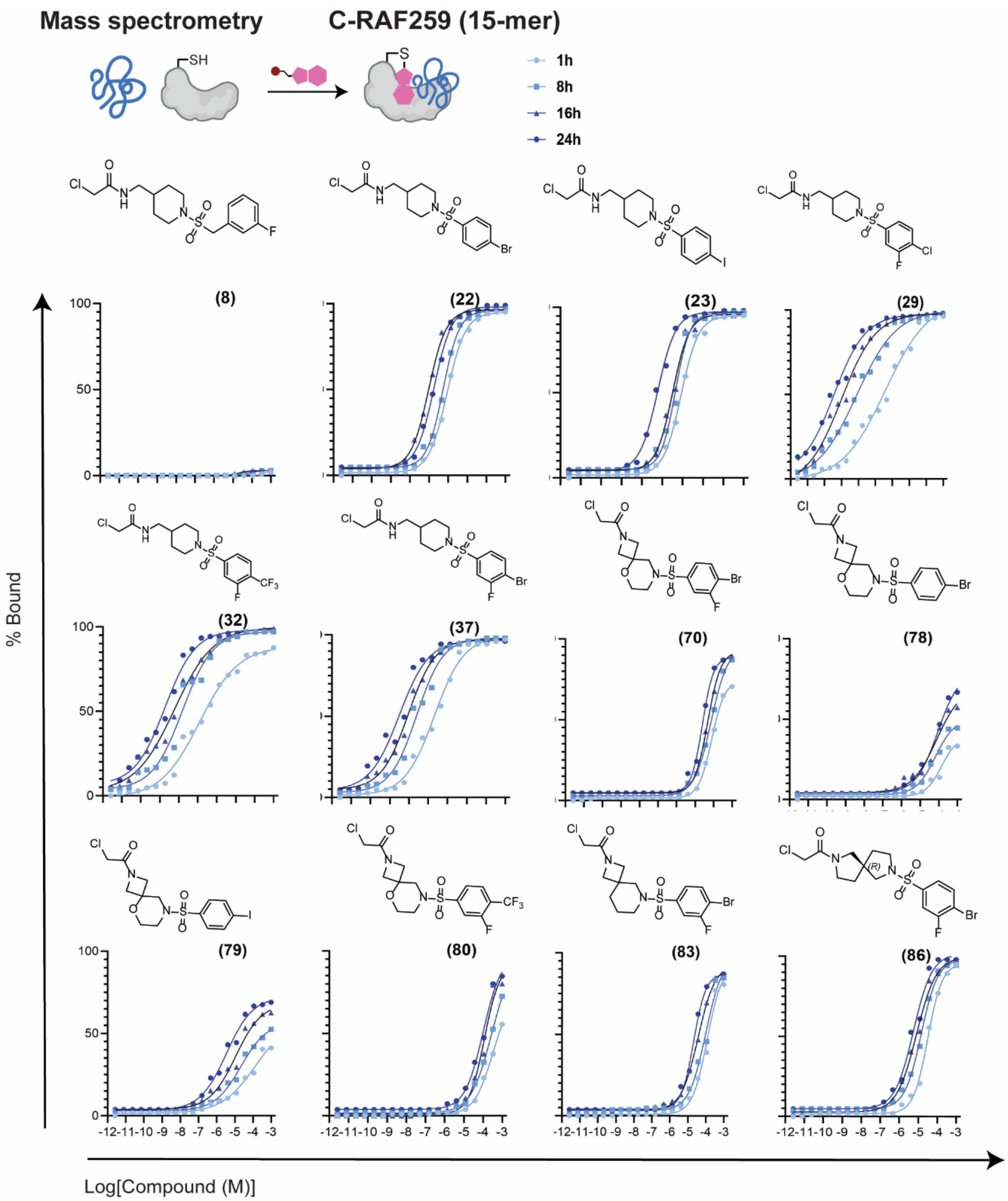

**Figure S25.** MS dose-response curves (C-RAF259 15-mer peptide).

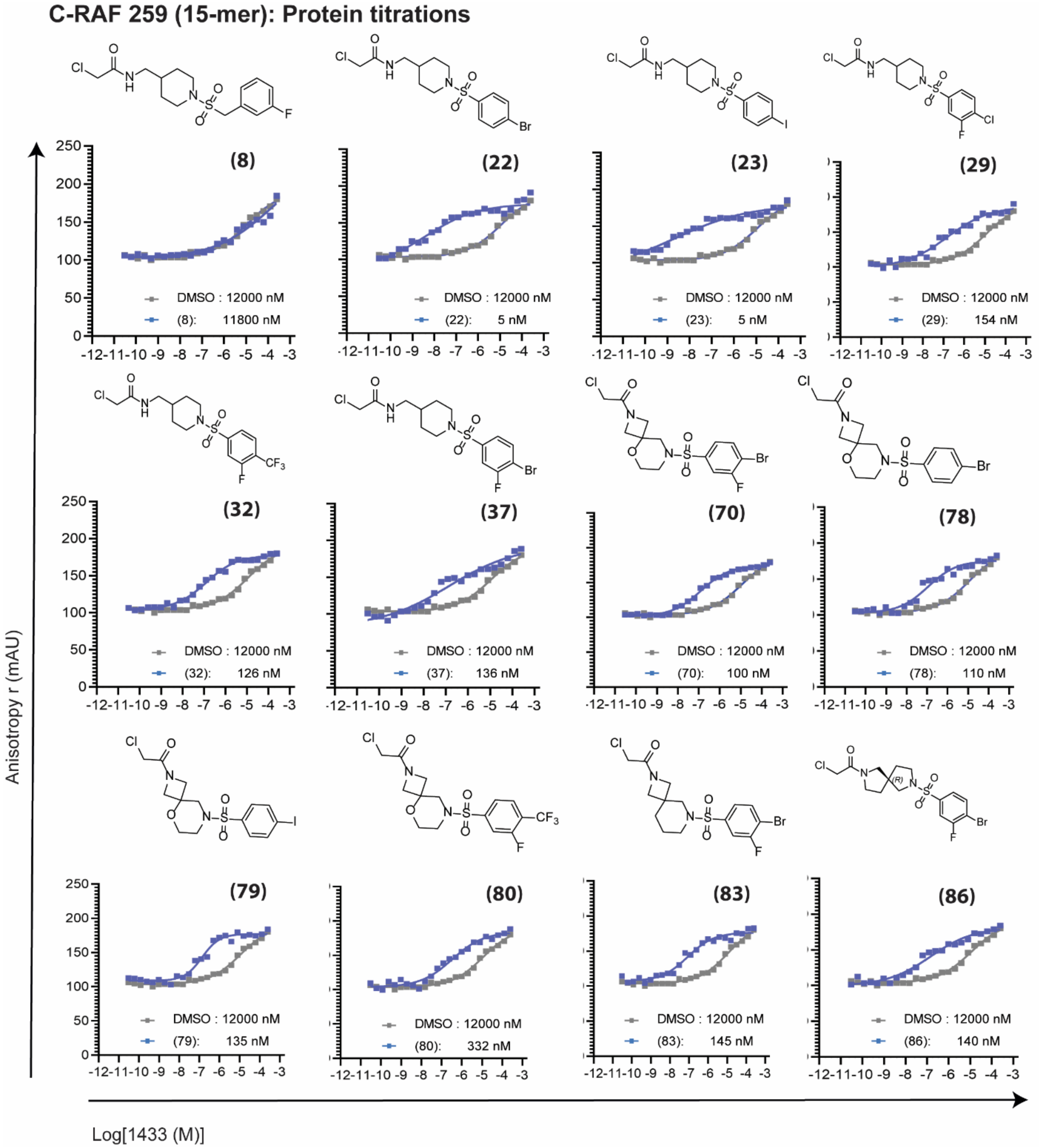

**Figure S26.** FA protein titrations (C-RAF259 15-mer peptide).

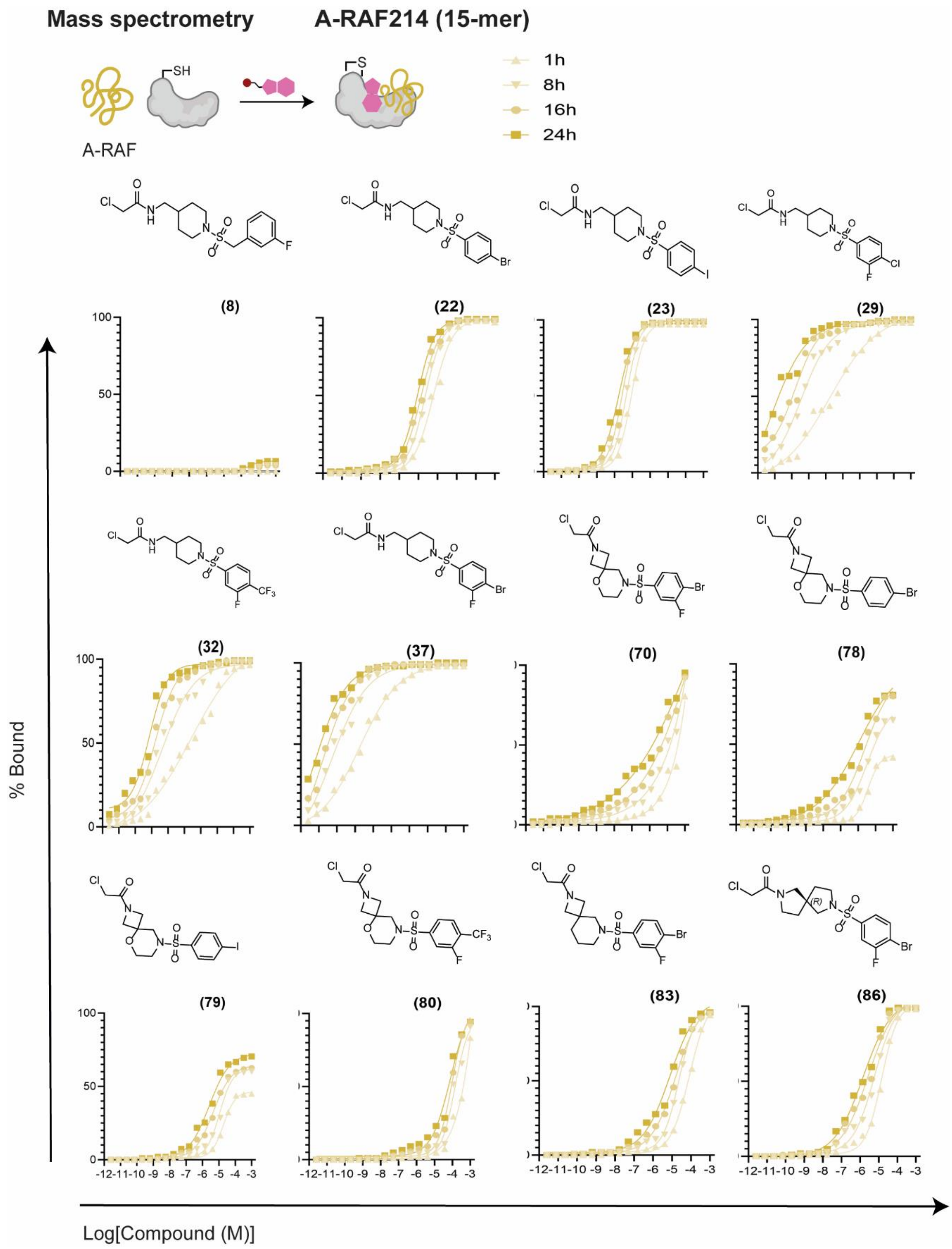

**Figure S27.** MS dose-response curves (A-RAF214 15-mer peptide).

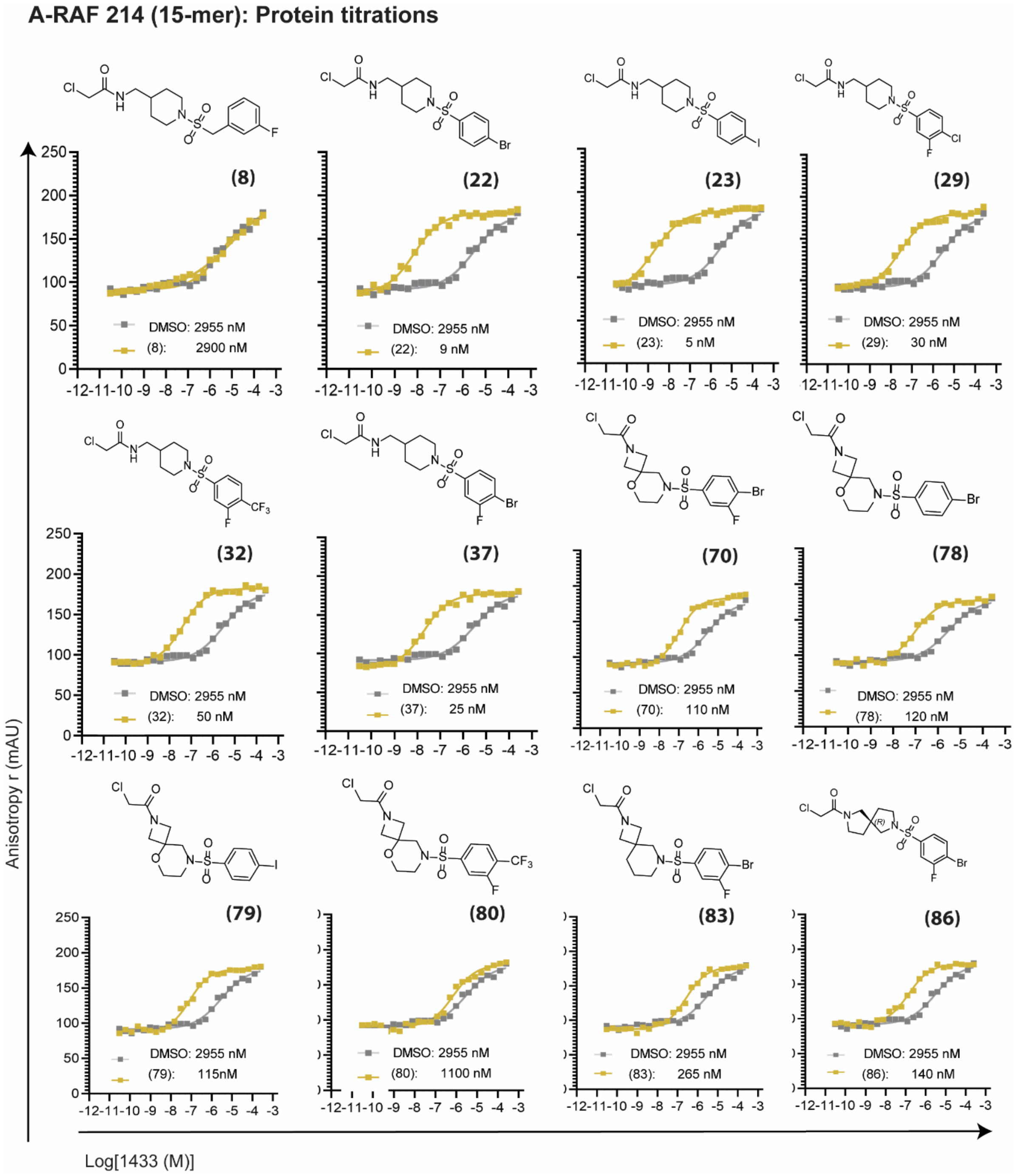

**Figure S28.** FA protein titrations (A-RAF214 15-mer peptide).

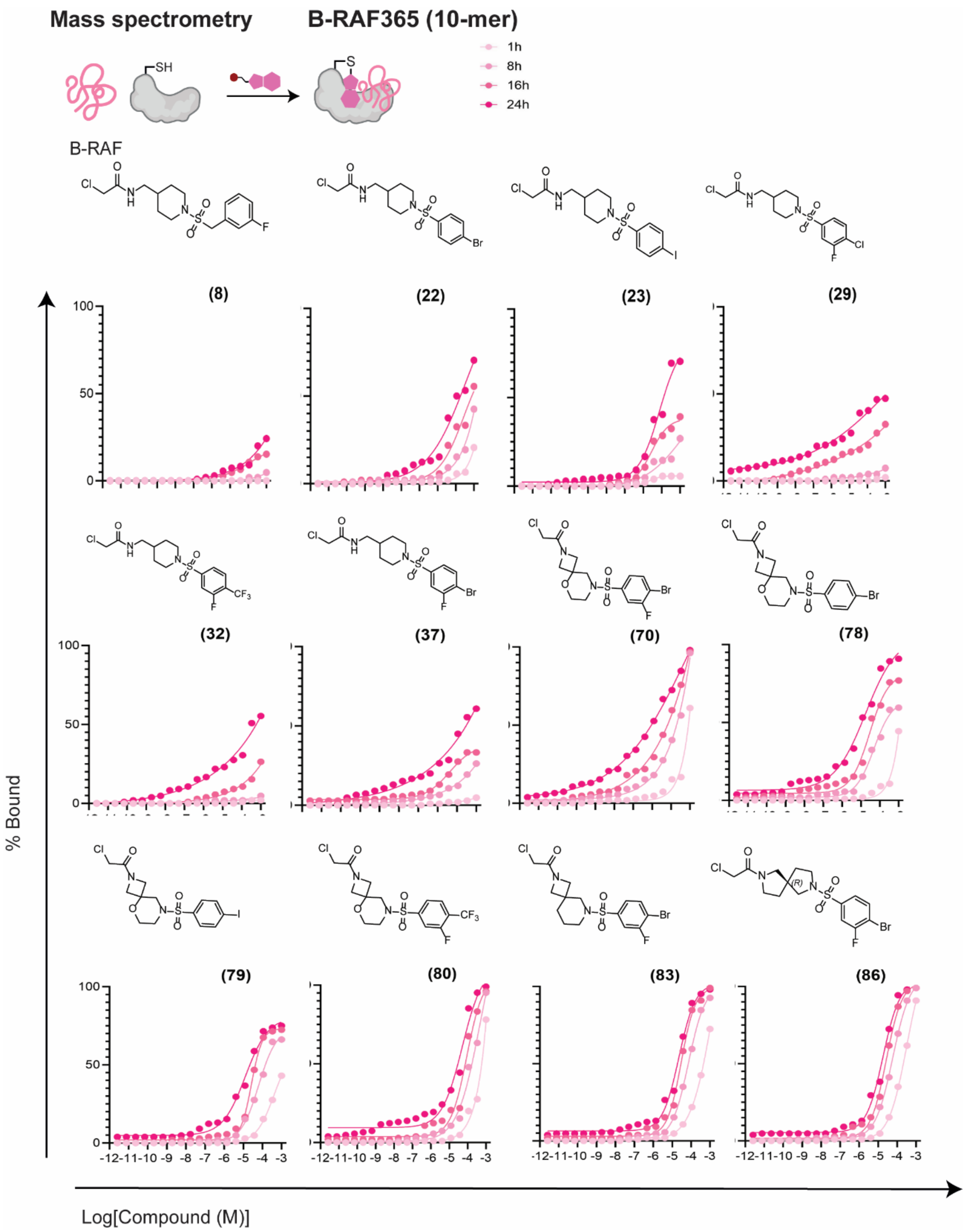

**Figure S29.** MS dose-response curves (B-RAF365 10-mer peptide).

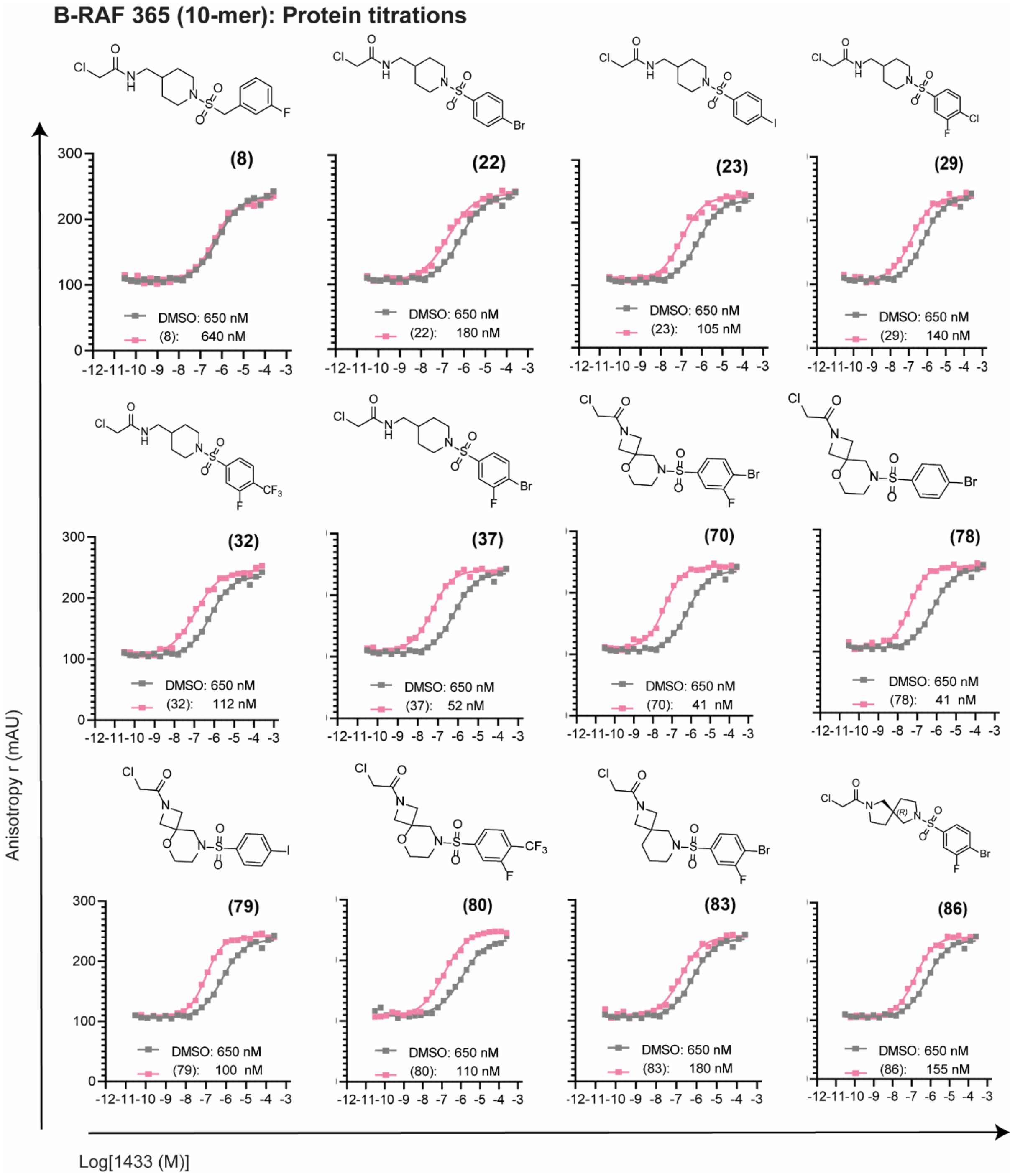

**Figure S30.** FA protein titrations (B-RAF365 10-mer peptide).

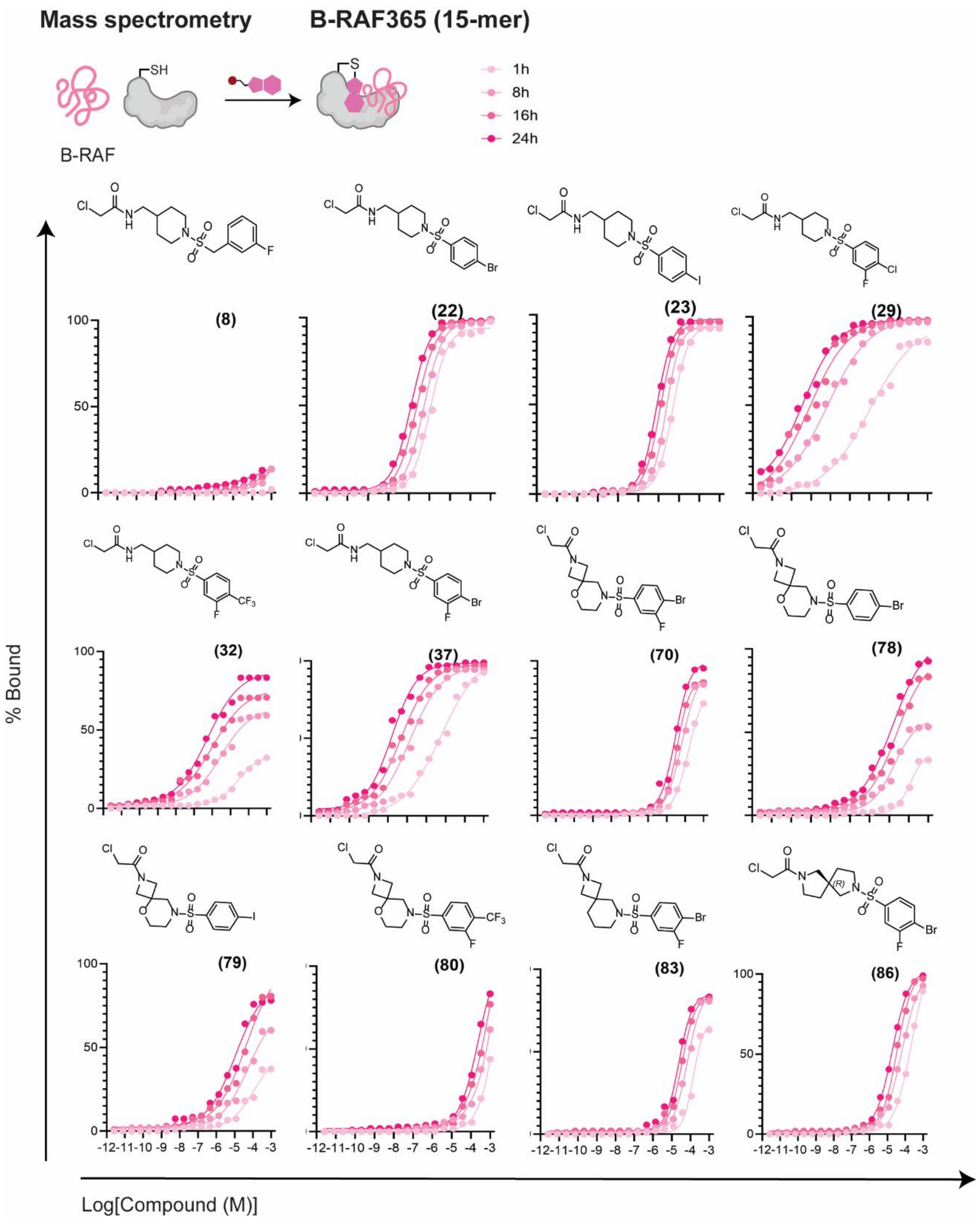

**Figure S31.** MS dose-response curves (B-RAF365 15-mer peptide).

**B-RAF 365 (15-mer): Protein titrations**

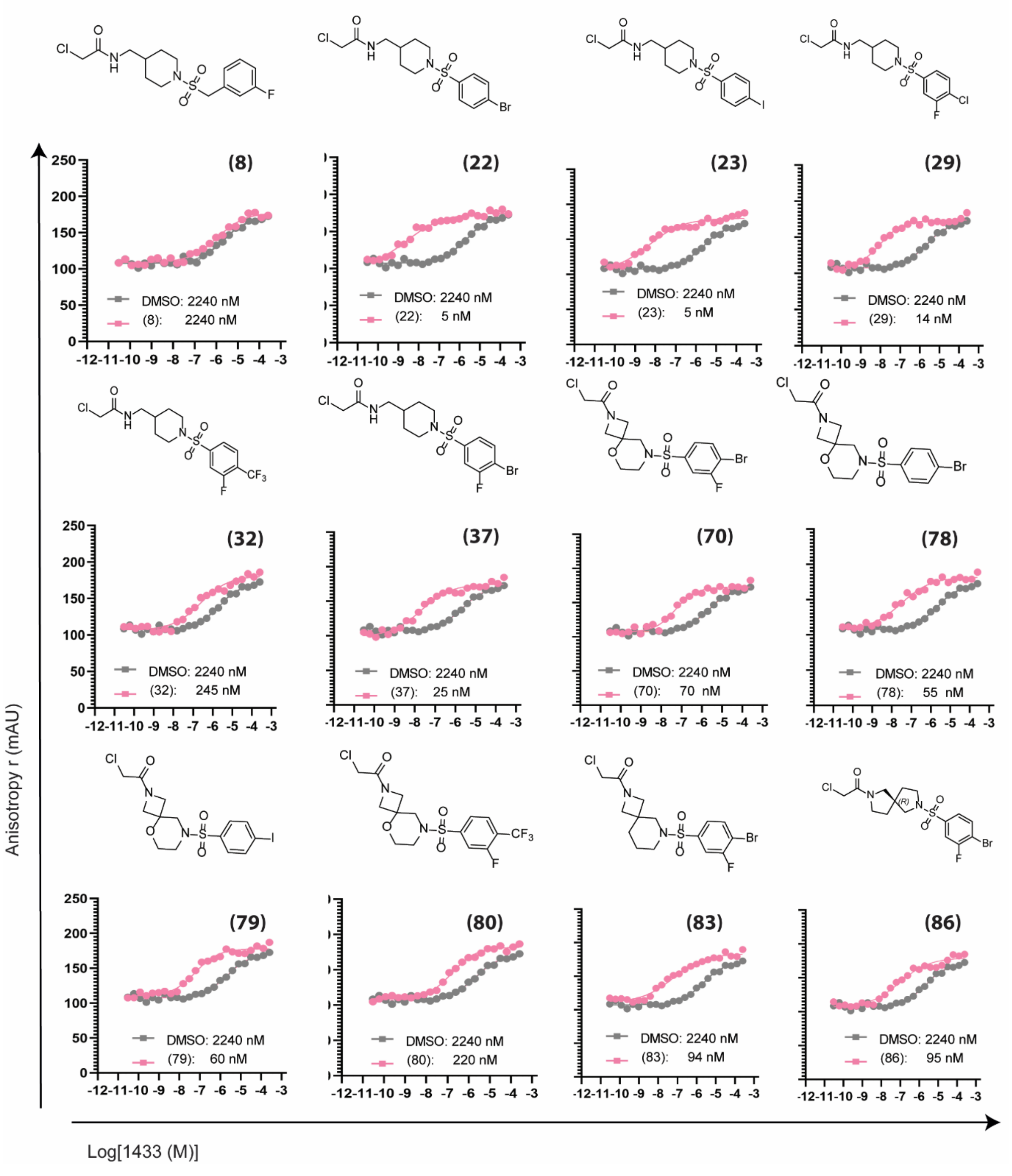

**Figure S32.** FA protein titrations (B-RAF365 15-mer peptide).

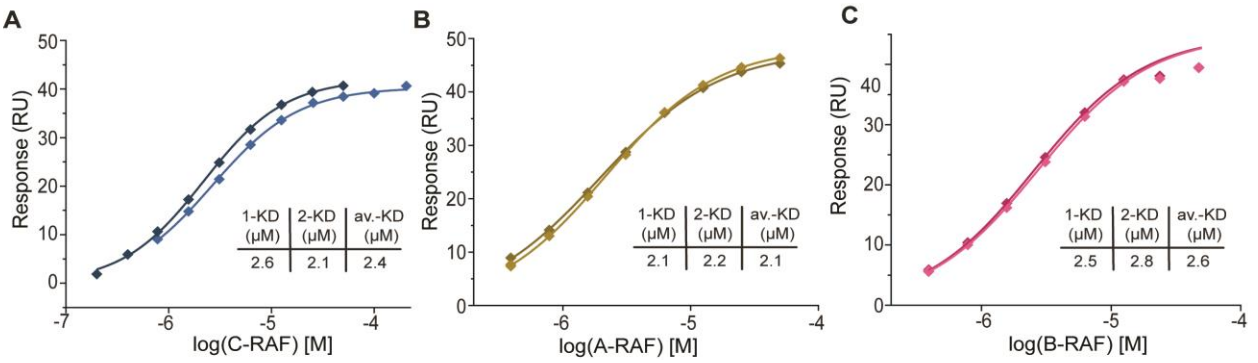

**Figure S33.** Affinity fitting of SPR data: acetylated 12-mer A-, B-, C-RAF peptides binding to 14-3-3σ captured on a chip (n=2). Response at equilibrium is plotted against peptide concentration to determine the affinity constant (only possible for binary interactions as for ternary interactions no equilibrium is reached during association).

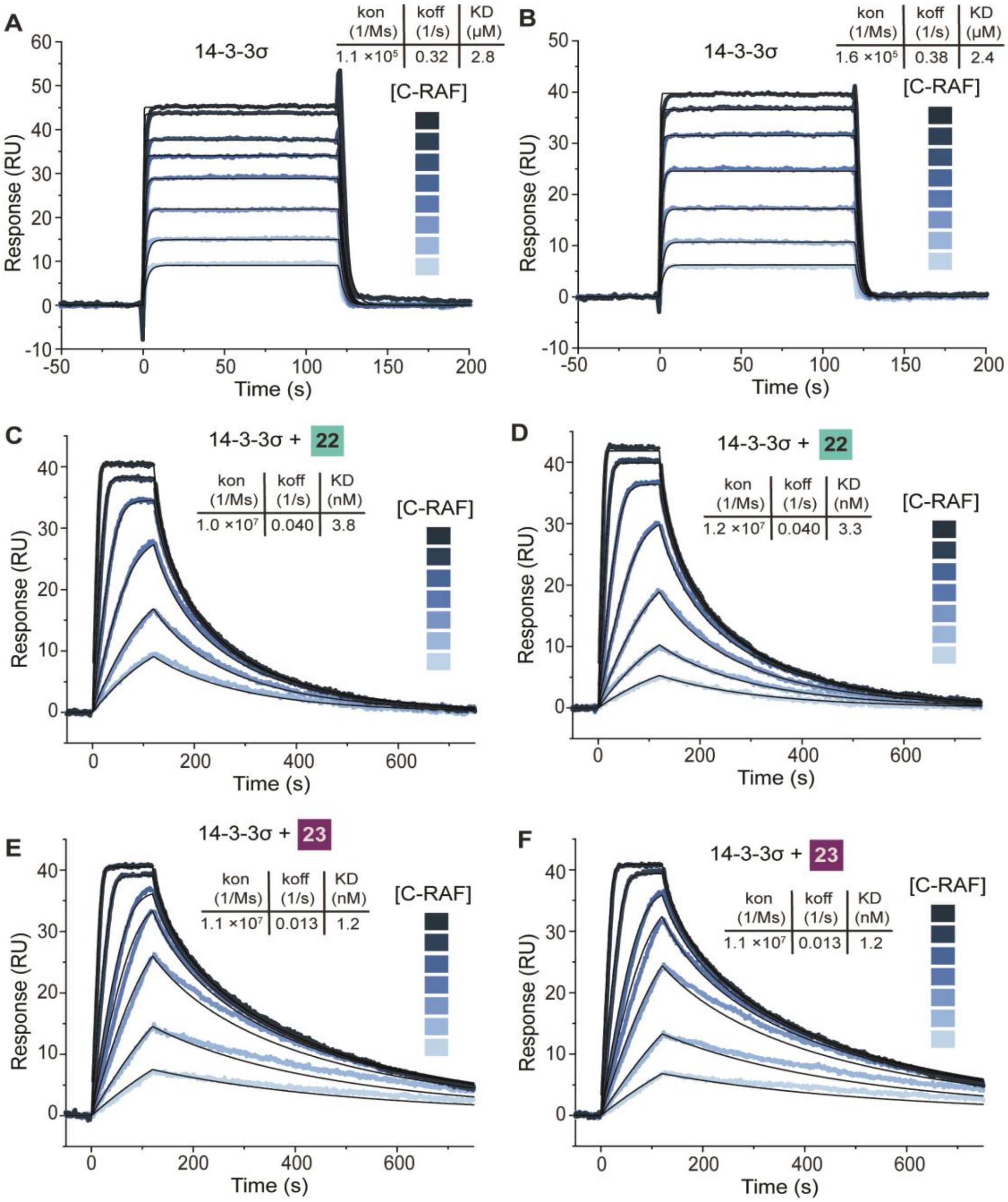

**Figure S34.** Kinetic fitting of SPR data: acetylated 12-mer C-RAF peptide binding to 14-3-3σ captured on a chip without molecular glue and in the presence of compounds **22** and **23** (n=2).

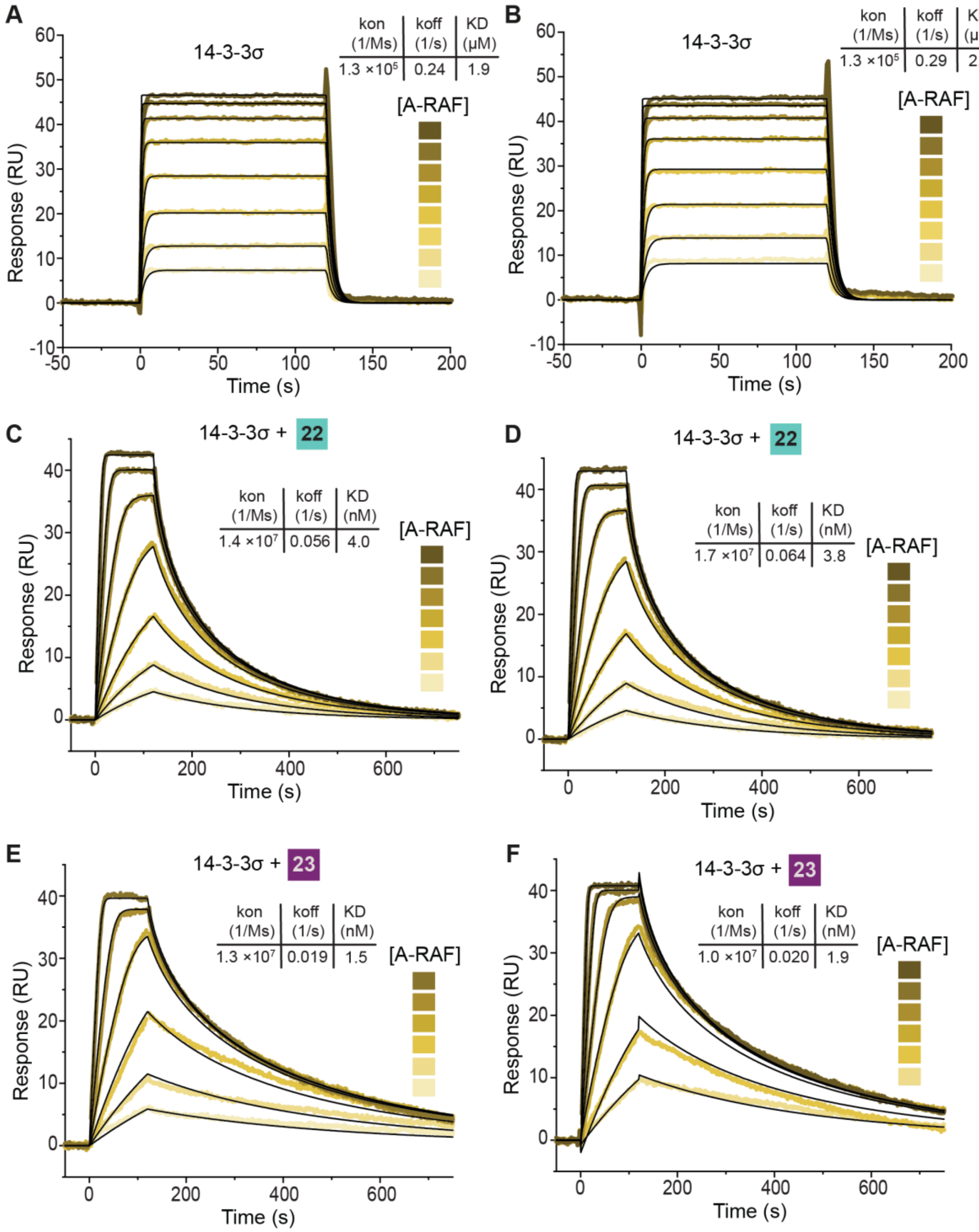

**Figure S35.** Kinetic fitting of SPR data: acetylated 12-mer A-RAF peptide binding to 14-3-3σ captured on a chip without molecular glue and in the presence of compounds **22** and **23** (n=2).

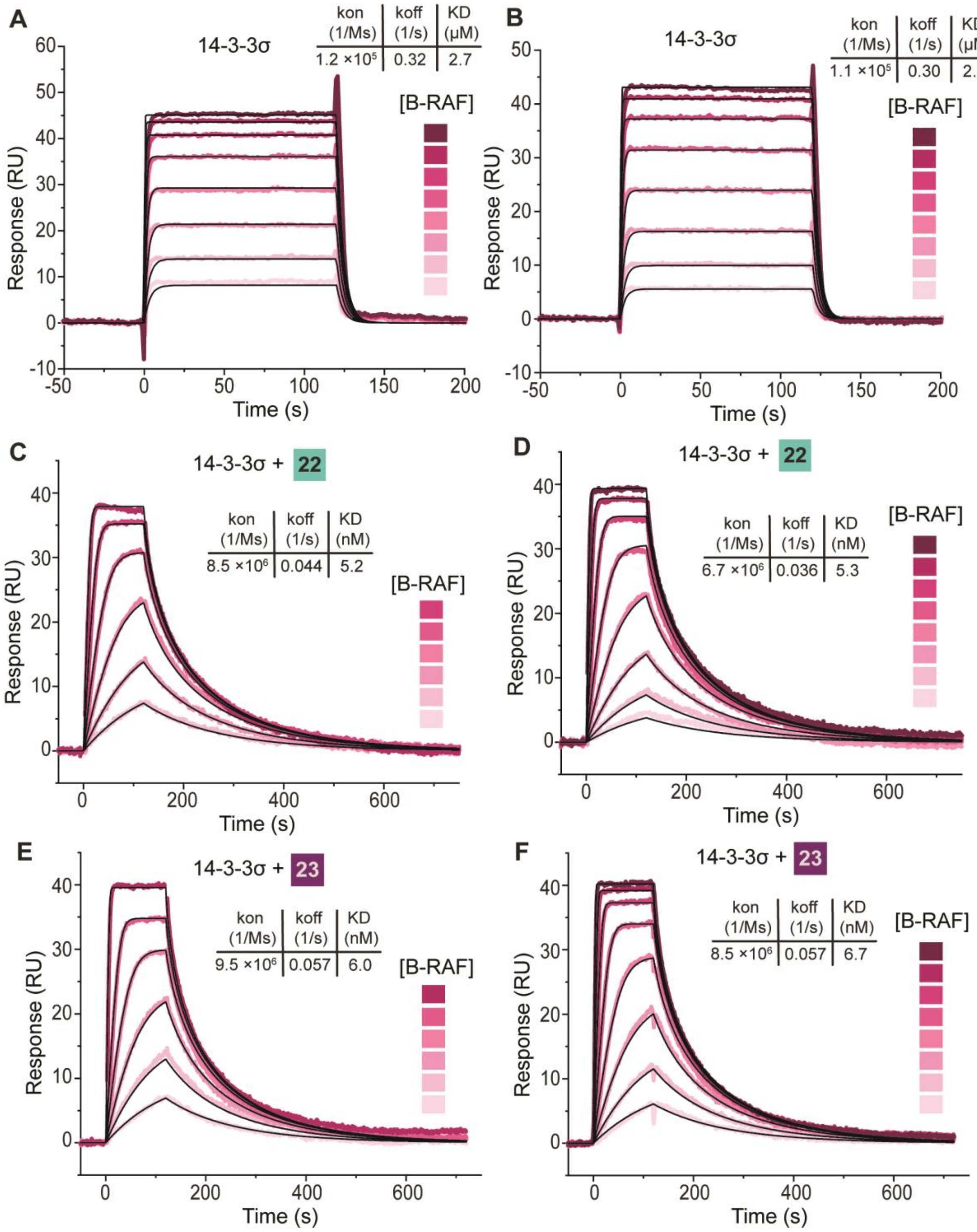

**Figure S36.** Kinetic fitting of SPR data: acetylated 12-mer B-RAF peptide binding to 14-3-3σ captured on a chip without molecular glue and in the presence of compound **22** and **23** (n=2).

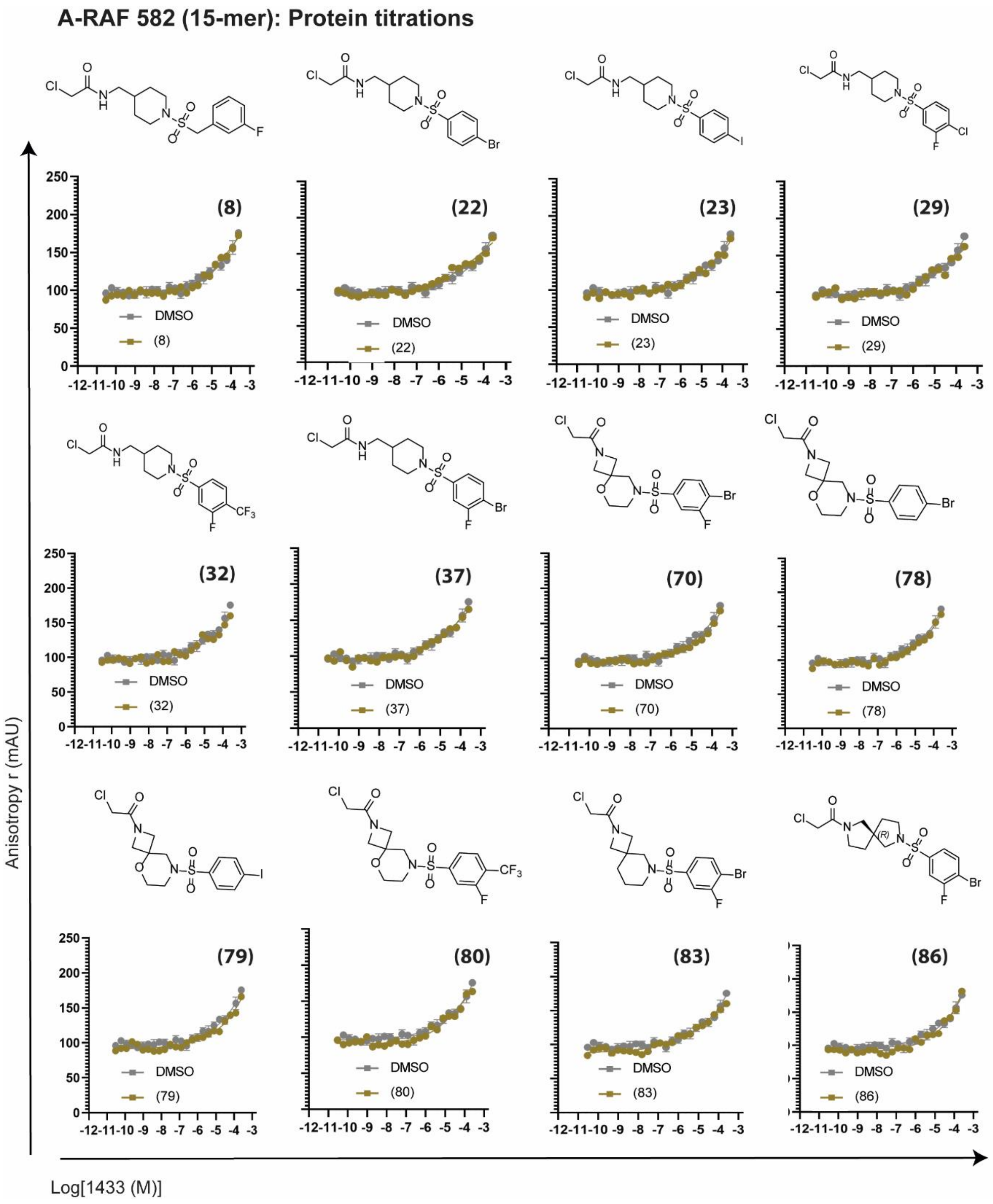

**Figure S37.** FA protein titrations (A-RAF582 15-mer peptide).

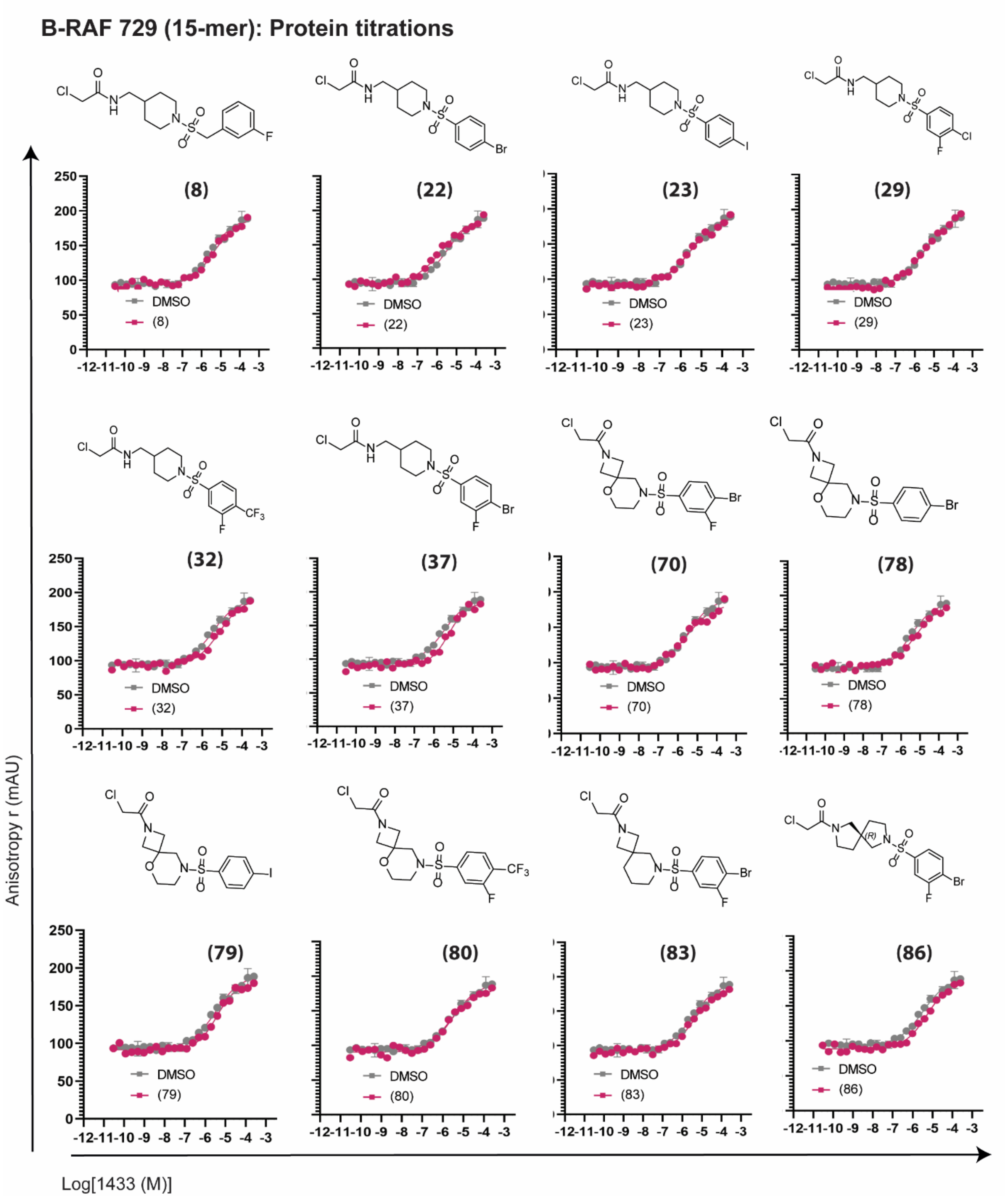

**Figure S38.** FA protein titrations (B-RAF729 15-mer peptide).

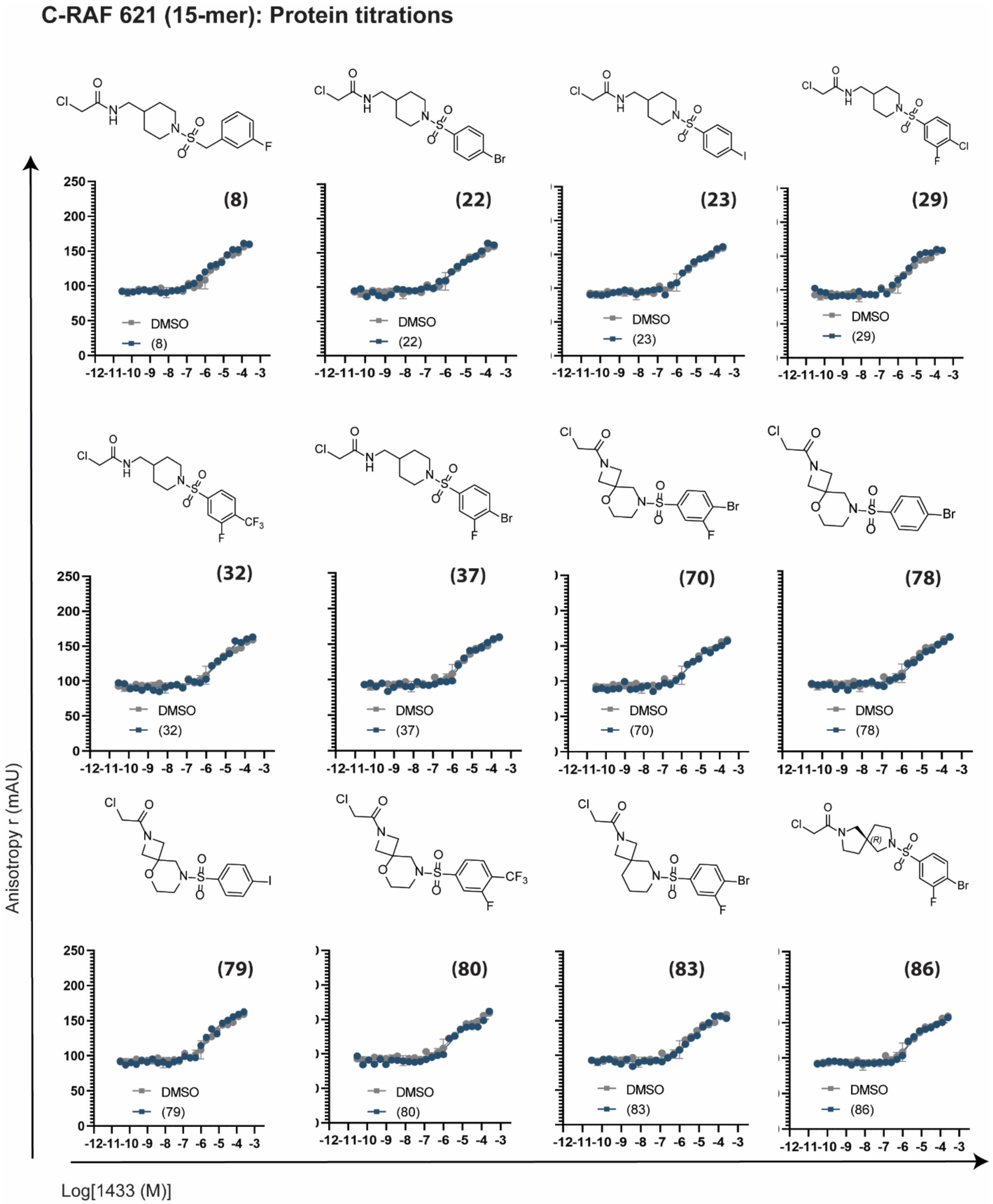

**Figure S39.** FA protein titrations (C-RAF621 15-mer peptide).

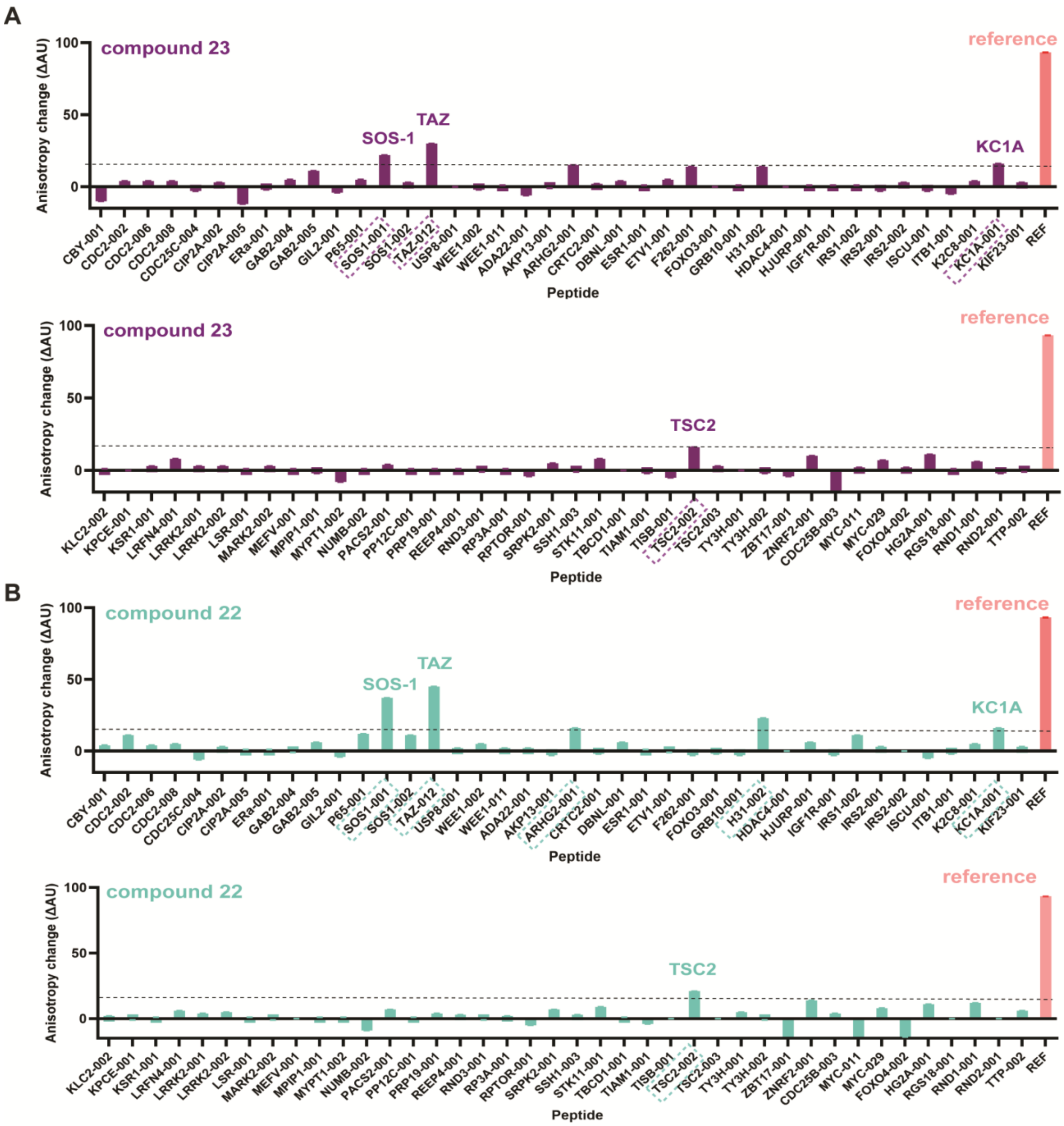

**Figure S40.** FA selectivity panel for compounds **23** (A) and **22** (B) with 14-3-3σ and 80 client peptides. Six client hits were identified: SOS1, TAZ, ARGH2, H31, KC1A and TSC2.

A

| Peptide | Phosphosite | -2 | -1 | Phospho | +1 | +2 | +3 |
|---|---|---|---|---|---|---|---|
| C-RAF | pS259 | S | T | pS | T | P | N |
| A-RAF | pS214 | S | T | pS | T | P | N |
| B-RAF | pS359 | S | S | pS | A | P | N |
| SOS1 | pS1161 | P | E | pS | A | P | A |
| TAZ | pS89 | S | H | pS | S | P | A |
| KC1A | pS218 | R | T | pS | L | P | W |
| TSC2 | pS1254 | S | L | pS | V | P | A |
| ARGH2 | pS886 | R | R | pS | L | P | A |
| H31 | pS29 | R | K | pS | A | P | A |

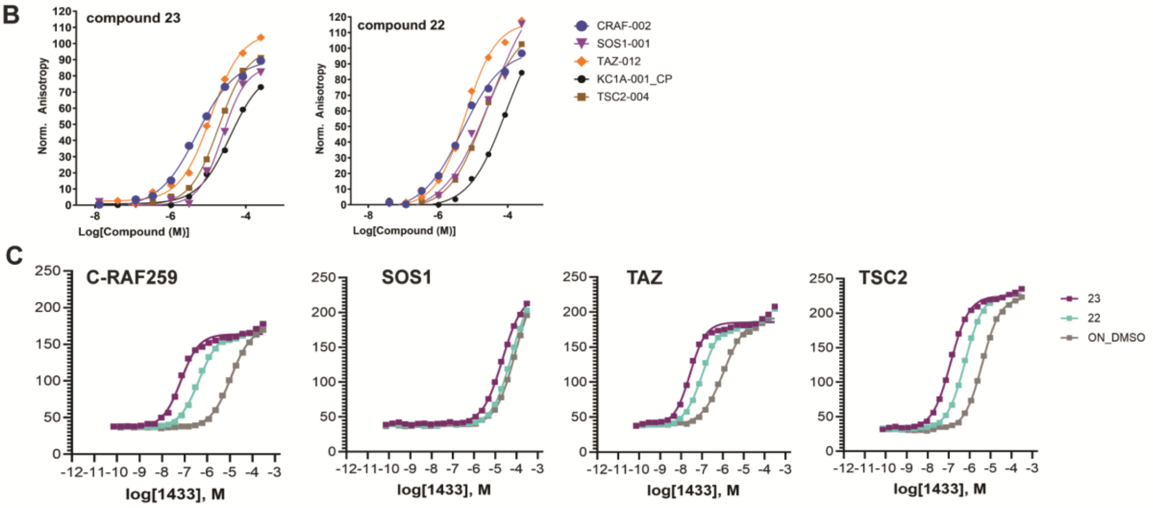

**Figure S41.** A) Phospho-sites and sequences around the phospho-site for client peptides that emerged as hits. B) FA compound dose-response follow-up for the four common clients. C) FA protein titrations follow-up. $EC_{50}$ values, *app*Kds and fold-stabilization are shown in tables S7 and S8.

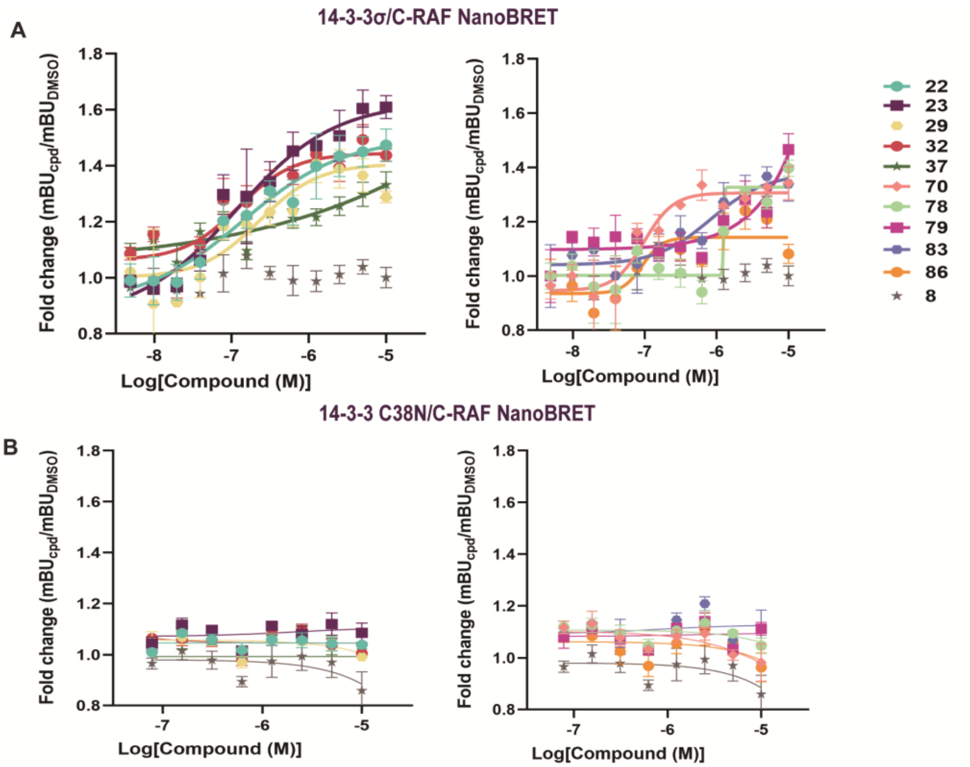

**Figure S42.** A) 14-3-3σ/C-RAF NanoBRET dose-response data in HEK293T cells. Left: piperidine-containing compounds, rights: spirocycles. B) 14-3-3 C38N/C-RAF NanoBRET dose-response data in HEK293T cells. Left: piperidine-containing compounds, rights: spirocycles

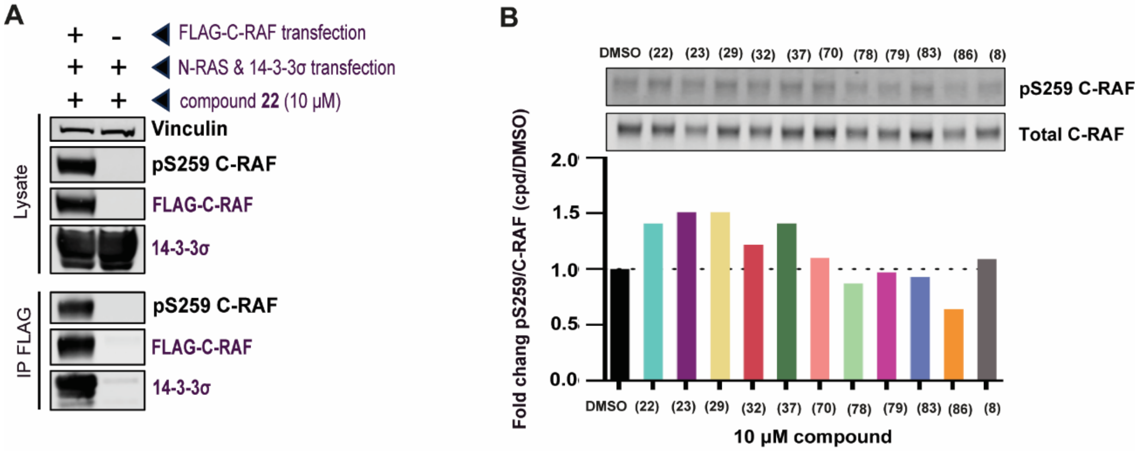

**Figure S43.** A) Co-IP for **22**, comparing endogenous and transfected C-RAF. B) Protection of phosphorylation of C-RAF pS259 site in MIA PaCa-2 cells.

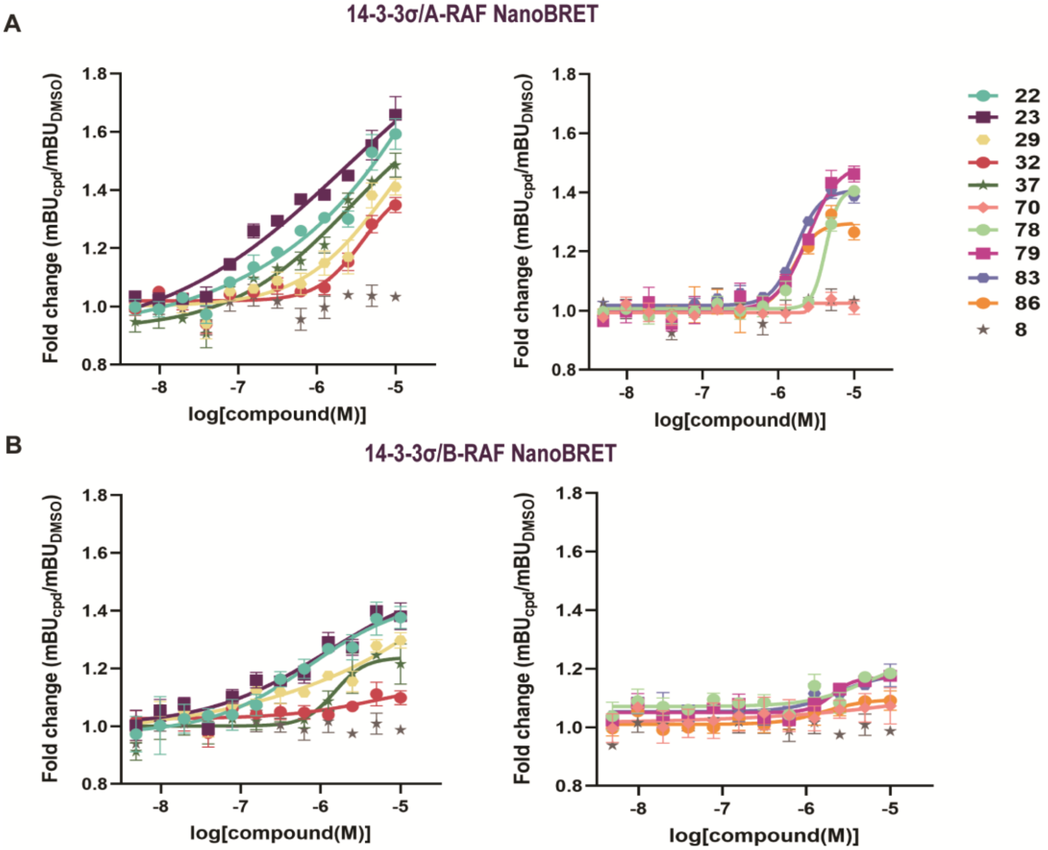

**Figure S44.** A) 14-3-3σ/A-RAF NanoBRET dose-response data in HEK293T cells. Left: piperidine-containing compounds, rights: spirocycles. B) 14-3-3σ/B-RAF NanoBRET dose-response data in HEK293T cells. Left: piperidine-containing compounds, rights: spirocycles.

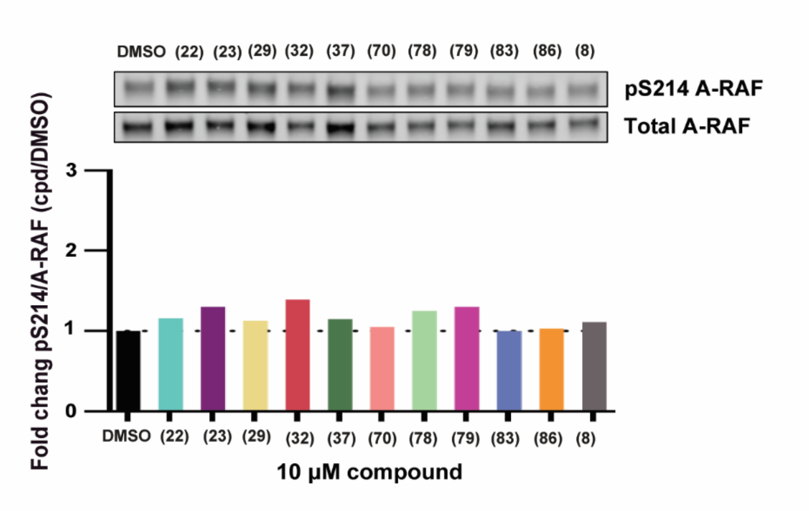

**Figure S45.** Protection of phosphorylation of A-RAF pS214 site in HEK293T cells. The assay was performed with the pS259 C-RAF antibody.

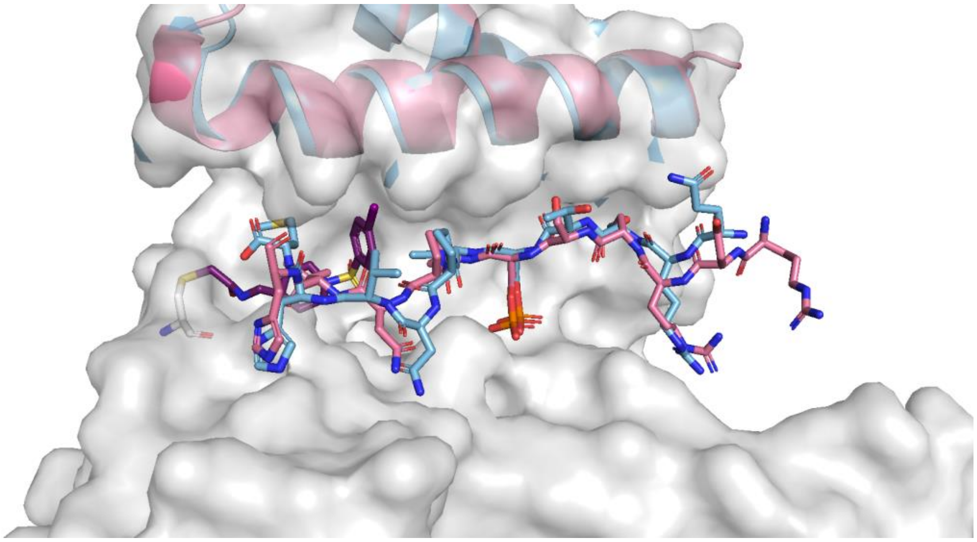

**Figure S46.** Overlay of the cryoEM 14-3-3/B-RAF structure (PDB 6NYB) with the crystal structure of compound **23**/14-3-3σ/CRAF259 12-mer peptide. Compound **23** is shown as purple sticks, B-RAF365 as pink sticks and C-RAF259 as cyan sticks. Helix 9 is in the same position for both structures.

### 3. SUPPLEMENTARY TABLES

**Table S1.** Molecular structure of compounds, assays performed and table number for the assays.

| Compound No. | SMDC ID | Structure | MS [a] | FA – Compound titration [b] | FA – Protein titration [c] | Crystallography [d] |
|---|---|---|---|---|---|---|
| **(6)** | **1075353** | 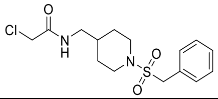 | S2 | S3 | - | - |
| **(7)** | **1075476** | 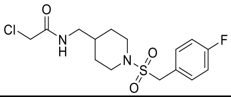 | S2 | S3 | - | - |
| **(8)** | **1075477** | 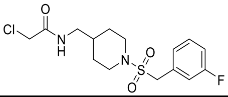 | S2 | S3 | - | - |
| **(9)** | **1075352** | 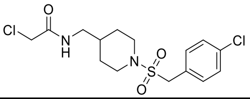 | S2 | S3 | - | - |
| **(10)** | **1076407** | 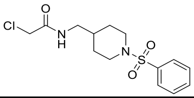 | S2 | S3 | - | - |
| **(11)** | **1076408** | 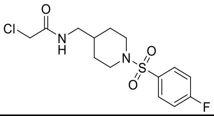 | S2 | S3 | S3 | - |
| **(12)** | **1075475** | 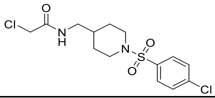 | S2 | S3 | S3 | C-RAF 259 |
| **(13)** | **1076390** | 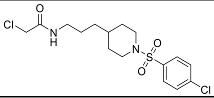 | S2 | S3 | - | - |
| **(14)** | **1076413** | 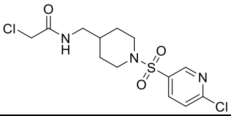 | S2 | S3 | - | - |
| **(15)** | **1083849** | 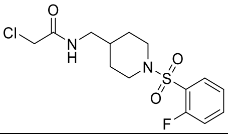 | S2 | S3 | - | - |
| **(16)** | **1083843** | 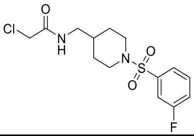 | S2 | S3 | - | - |
| **(17)** | **1083844** | 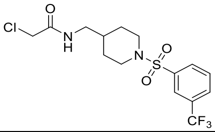 | S2 | S3 | - | - |
| **(18)** | **1076412** | 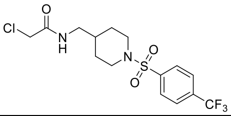 | S2 | S3 | S3 | - |
| **(19)** | **1083842** | 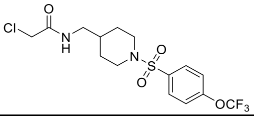 | S2 | S3 | S3 | - |

| | | | | | | |
|---|---|---|---|---|---|---|
| **(20)** | **1083841** | 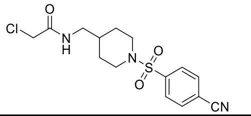 | S2 | S3 | - | - |
| **(21)** | **1075354** | 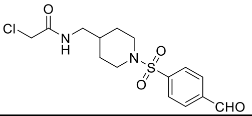 | S2 | S3 | S3 | C-RAF 259 |
| **(22)** | **1083853** | 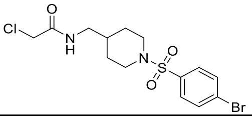 | S2 | S3 | S3 | C-RAF 259<br>A-RAF 214<br>B-RAF 365 |
| **(23)** | **1083848** | 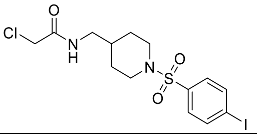 | S2 | S3 | S3 | C-RAF 259<br>A-RAF 214<br>B-RAF 365 |
| **(24)** | **1083852** | 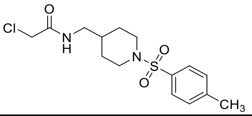 | S2 | S3 | S3 | - |
| **(25)** | **1076411** | 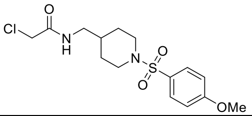 | S2 | S3 | - | - |
| **(26)** | **1124380** | 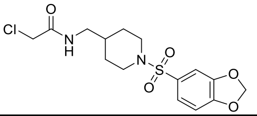 | S2 | S3 | - | - |
| **(27)** | **1083846** | 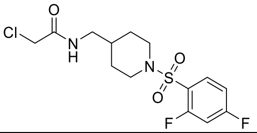 | S2 | S3 | - | - |
| **(28)** | **1083845** | 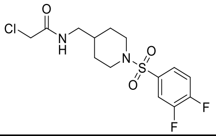 | S2 | S3 | - | - |
| **(29)** | **1076409** | 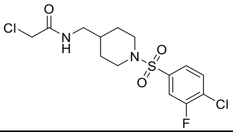 | S2 | S3 | S3 | C-RAF 259 |
| **(30)** | **1076410** | 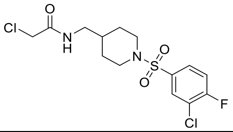 | S2 | S3 | S3 | - |
| **(31)** | **1083850** | 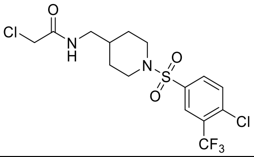 | S2 | S3 | - | - |
| **(32)** | **1083854** | 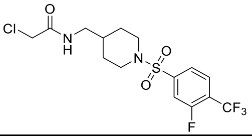 | S2 | S3 | S3 | C-RAF 259 |

| | | | | | | |
|---|---|---|---|---|---|---|
| **(33)** | **1083847** | 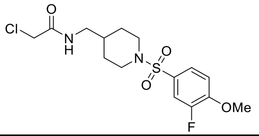 | S2 | S3 | - | - |
| **(34)** | **1083851** | 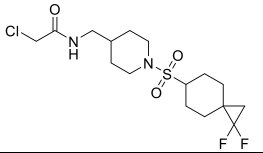 | S2 | S3 | - | - |
| **(35)** | **1083855** | 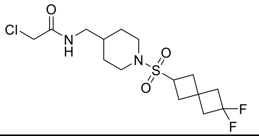 | S2 | S3 | - | - |
| **(36)** | **1083916** | 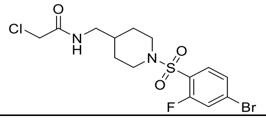 | S2 | S3 | S3 | - |
| **(37)** | **1083917** | 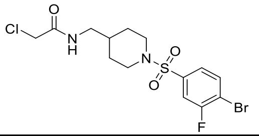 | S2 | S3 | S3 | C-RAF 259 |
| **(38)** | **1083918** | 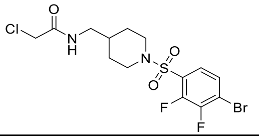 | S2 | S3 | S3 | - |
| **(39)** | **1083919** | 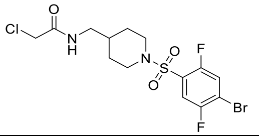 | S2 | S3 | S3 | - |
| **(40)** | **1083920** | 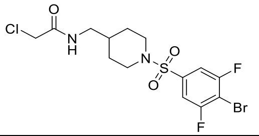 | S2 | S3 | S3 | - |
| **(43)** | **1124381** | 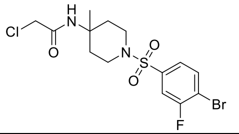 | S2 | S3 | - | - |
| **(46)** | **1124382** | 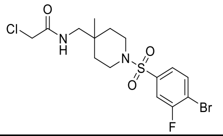 | S2 | S3 | S3 | - |
| **(49)** | **1084346** | 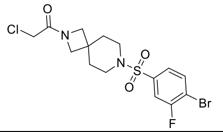 | S2 | S3 | - | - |
| **(52)** | **1084353** | 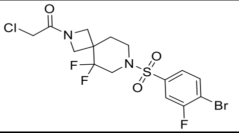 | S2 | S3 | - | - |

| | | | | | | |
|---|---|---|---|---|---|---|
| **(55)** | **1084347** | 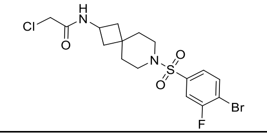 | S2 | S3 | - | - |
| **(58)** | **1084348** | 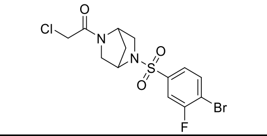 | S2 | S3 | S3 | - |
| **(61)** | **1084349** | 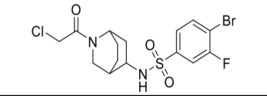 | S2 | S3 | - | - |
| **(64)** | **1084350** | 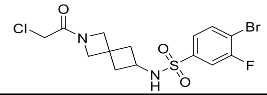 | S2 | S3 | S3 | - |
| **(67)** | **1084351** | 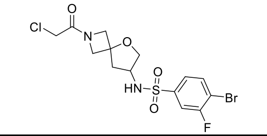 | S2 | S3 | S3 | - |
| **(70)** | **1084352** | 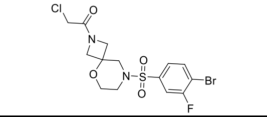 | S2 | S3 | S3 | C-RAF 259 |
| **(73)** | **1084354** | 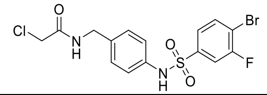 | S2 | S3 | - | - |
| **(74)** | **1084757** | 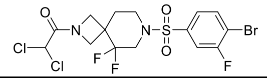 | S2 | S3 | - | - |
| **(75)** | **1084758** | 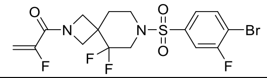 | S2 | S3 | - | - |
| **(78)** | **1124378** | 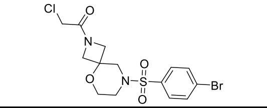 | S2 | S3 | S3 | C-RAF 259<br>A-RAF 214<br>B-RAF 365 |
| **(79)** | **1124379** | 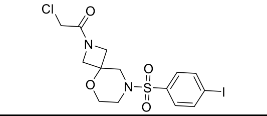 | S2 | S3 | S3 | C-RAF 259<br>B-RAF 365 |
| **(80)** | **1124898** | 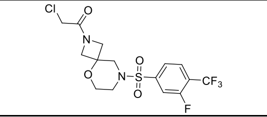 | S2 | S3 | S3 | C-RAF 259 |
| **(83)** | **1124383** | 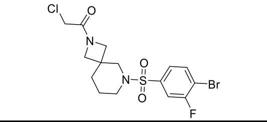 | S2 | S3 | S3 | C-RAF 259 |
| **(86)** | **1124384** | 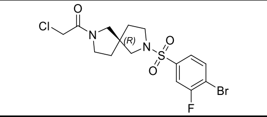 | S2 | S3 | S3 | C-RAF 259<br>A-RAF 214<br>B-RAF 365 |
| **(89)** | **1124385** | 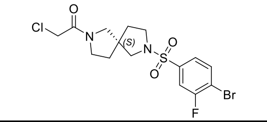 | S2 | S3 | S3 | - |

**Table S2.** Percentage (%) bound of compound (1 μM) to 14-3-3σ (100 nM) measured by mass spectrometry in the absence of peptide (apo) or with C-RAF259 peptide (18 μM) after 1, 8, 16 and 24 hours.

| | APO | | | | C-RAF259 | | | |
|---|---|---|---|---|---|---|---|---|
| **Compound No.** | **1h** | **8h** | **16h** | **24h** | **1h** | **8h** | **16h** | **24h** |
| **(6)** | 0 | 1 | 4.7 | 4.7 | 0.9 | 2.9 | 6.5 | 7 |
| **(7)** | 1 | 1.9 | 1.9 | 2.9 | 1.5 | 1.5 | 2.5 | 4.7 |
| **(8)** | 0 | 0 | 1.9 | 2.9 | 2.9 | 5.6 | 5.6 | 5.6 |
| **(9)** | 0 | 0 | 1.5 | 1.9 | 0 | 0.5 | 2.9 | 3.2 |
| **(10)** | 4.7 | 4.7 | 4.7 | 4.7 | 4.7 | 5.6 | 5.7 | 13 |
| **(11)** | 0 | 1.5 | 4.7 | 6.9 | 6.9 | 15 | 26 | 50.3 |
| **(12)** | 1 | 1.5 | 2.9 | 4.3 | 12.1 | 32.8 | 47.8 | 64.4 |
| **(13)** | 0 | 1.5 | 2.5 | 4.3 | 0.9 | 2.9 | 5.6 | 14.1 |
| **(14)** | 1.9 | 2.5 | 3 | 15 | 4.7 | 7.4 | 9.9 | 20.9 |
| **(15)** | 0.5 | 1.9 | 2.4 | 2.4 | 3.3 | 8.2 | 13.7 | 23.3 |
| **(16)** | 0 | 1 | 1.9 | 2.9 | 3.3 | 8.2 | 18 | 32.4 |
| **(17)** | 0 | 0 | 0.5 | 0.9 | 1.9 | 3.4 | 6.5 | 12.3 |
| **(18)** | 0 | 0.9 | 6.5 | 12.7 | 8.2 | 17.9 | 28.4 | 46.8 |
| **(19)** | 0 | 0.5 | 2.5 | 6.5 | 20.5 | 41.3 | 69.5 | 91.3 |
| **(20)** | 0 | 0.5 | 1.5 | 2.9 | 4.7 | 12.1 | 23.9 | 42.3 |
| **(21)** | 1,5 | 2.9 | 4.4 | 7.4 | 2.9 | 6.5 | 10.7 | 32.2 |
| **(22)** | 6.5 | 10.7 | 16 | 20.3 | 61 | 85.5 | 94.8 | 98.5 |
| **(23)** | 0 | 0 | 0.5 | 0.9 | 55.6 | 78.5 | 88.9 | 91 |
| **(24)** | 0 | 0.5 | 0.5 | 1.5 | 7.3 | 13.7 | 26.7 | 51.8 |
| **(25)** | 0.5 | 1.9 | 7.4 | 13.8 | 10.6 | 15 | 27.1 | 44.4 |
| **(26)** | 0 | 0 | 1.9 | 5.2 | 4.3 | 11.3 | 22.6 | 34.1 |
| **(27)** | 1 | 2,5 | 3.8 | 5.2 | 6.9 | 14.1 | 27.5 | 50.3 |
| **(28)** | 1.9 | 4.7 | 6.5 | 9.1 | 4.3 | 15.2 | 27.5 | 40 |
| **(29)** | 0 | 1.9 | 4.7 | 8.2 | 20.5 | 38.1 | 57.1 | 80.1 |
| **(30)** | 1 | 2.5 | 6.5 | 22.5 | 11 | 24.8 | 41.8 | 60.3 |
| **(31)** | 0.9 | 1.9 | 2.5 | 3.8 | 21 | 38.1 | 59.1 | 81.4 |
| **(32)** | 0 | 0.5 | 0.9 | 0.9 | 14.1 | 21.6 | 39.3 | 74.1 |
| **(33)** | 0 | 0 | 0.9 | 0.9 | 7.7 | 18.5 | 31.8 | 50.3 |
| **(34)** | 0 | 0.5 | 0.9 | 0.9 | 0.5 | 0.9 | 0.9 | 3.8 |
| **(35)** | 0 | 0 | 0 | 0.5 | 0 | 0 | 0 | 3.8 |
| **(36)** | 0 | 0.9 | 2.9 | 4.5 | 20.6 | 41.1 | 69.5 | 91.7 |
| **(37)** | 0 | 0.9 | 2.5 | 7.4 | 48.7 | 77.1 | 95.2 | 98 |
| **(38)** | 0 | 0.9 | 2.9 | 4.4 | 28 | 43.9 | 61.9 | 83.3 |
| **(39)** | 0 | 1.9 | 3.8 | 3.8 | 39.3 | 64.6 | 78.1 | 96.7 |
| **(40)** | 0 | 0.5 | 1.5 | 4.7 | 35.9 | 60.5 | 78.9 | 93.9 |
| **(43)** | 0 | 0.5 | 3.5 | 6.1 | 4.3 | 9.1 | 16.7 | 32.9 |
| **(46)** | 0.5 | 1.9 | 4.7 | 8.2 | 25.8 | 39.4 | 49 | 84 |
| **(49)** | 1.9 | 5.2 | 13.8 | 15.9 | 11.1 | 32.4 | 53.8 | 86.6 |
| **(52)** | 2.9 | 11.3 | 18.5 | 28.6 | 25 | 45.1 | 64.3 | 80.2 |
| **(55)** | 0 | 0.9 | 1.9 | 7.4 | 6.1 | 9.9 | 20.6 | 34.6 |
| **(58)** | 4.7 | 17.3 | 28 | 51.8 | 16.5 | 45.2 | 71.2 | 80.5 |
| **(61)** | 5.2 | 13.3 | 21.8 | 36.8 | 37.1 | 68 | 80.9 | 83.3 |
| **(64)** | 9.9 | 27.2 | 47.2 | 65 | 27.5 | 53.6 | 79 | 80.8 |
| **(67)** | 2.9 | 9.1 | 15.9 | 23 | 15 | 30.4 | 55.6 | 81.5 |
| **(70)** | 2.9 | 7.4 | 15.9 | 20 | 30 | 47.5 | 68 | 87.2 |
| **(73)** | 0 | 1.5 | 1.5 | 3.8 | 1.9 | 2.9 | 12.1 | 31 |
| **(74)** | 0 | 0 | 0 | 0 | 0 | 0 | 0 | 4.7 |
| **(75)** | 0 | 0 | 0 | 0 | 0 | 0 | 0 | 2.9 |
| **(78)** | 1.9 | 3.8 | 12.2 | 14.1 | 8.3 | 25.3 | 35.6 | 70 |
| **(79)** | 1.5 | 5.6 | 10.6 | 32.4 | 15 | 29.5 | 52.6 | 72 |
| **(80)** | 5.6 | 5.6 | 7.8 | 11 | 6 | 7 | 15.2 | 25.1 |

| | | | | | | | | |
|---|---|---|---|---|---|---|---|---|
| **(83)** | 1.9 | 3.8 | 14.5 | 25.6 | 27 | 52.5 | 66.3 | 84 |
| **(86)** | 3.3 | 10.7 | 23 | 36.8 | 64.6 | 93.1 | 97.1 | 98.5 |
| **(89)** | 5.6 | 14.5 | 36.2 | 53.9 | 43 | 65.8 | 82 | 92 |

**Table S3.** $EC_{50}$ values derived from FA compound titrations in the presence of 10 nM of C-RAF259-FAM labeled peptide and 5 µM 14-3-3σ. Protein titrations with C-RAF-259-FAM for compounds with $EC_{50}$ < 20 µM (10 nM of C-RAF259-FAM, 100 µM compound, 250 µM 14-3-3 starting concentration, 2-fold dilution). $EC_{50}$ and *app*Kd values refer to overnight measurements.

| | **Compound titrations** | **Protein titrations** | | |
|---|---|---|---|---|
| **Compound No.** | **$EC_{50}$ value C-RAF259 (µM)** | **Apparent Kd (nM) (100 µM compound)** | **Kd with DMSO reference** | **Fold stabilization (100 µM compound)** |
| **(6)** | >1000 | NA | NA | NA |
| **(7)** | >1000 | NA | NA | NA |
| **(8)** | >1000 | 6600 nM | 6700 nM | 1 |
| **(9)** | >1000 | NA | NA | NA |
| **(10)** | 66 ± 4 µM | NA | NA | NA |
| **(11)** | 17 ± 3 µM | 400 nM | 1325 nM | 4 |
| **(12)** | 3 ± 1 µM | 190 nM | 4200 nM | 22 |
| **(13)** | >1000 | NA | NA | NA |
| **(14)** | 82 ± 6 µM | NA | NA | NA |
| **(15)** | 87 ± 9 µM | NA | NA | NA |
| **(16)** | > 150 | NA | NA | NA |
| **(17)** | > 150 | NA | NA | NA |
| **(18)** | 17 ± 1 µM | 280 nM | 4200 nM | 15 |
| **(19)** | 13 ± 3 µM | 213 nM | 3050 nM | 14 |
| **(20)** | >150 | NA | NA | NA |
| **(21)** | 5 ± 1.0 µM | 480 nM | 4200 nM | 9 |
| **(22)** | 1 ± 0.5 µM | 47 nM | 8427 nM | 179 |
| **(23)** | 1 ± 0.5 µM | 30 nM | 8427 nM | 280 |
| **(24)** | 14 ± 1 µM | 110 nM | 2277 nM | 21 |
| **(25)** | 57 ± 9 µM | NA | NA | NA |
| **(26)** | 26 ± 3 µM | NA | NA | NA |
| **(27)** | >150 µM | NA | NA | NA |
| **(28)** | 88 ± 2 µM | NA | NA | NA |
| **(29)** | 1 ± 0.5 µM | 100 nM | 8427 nM | 84 |
| **(30)** | 12 ± 2 µM | 780 nM | 2277 nM | 3 |
| **(31)** | 8 ± 1 µM | 1420 nM | 2277 nM | 2 |
| **(32)** | 4 ± 1 µM | 140 nM | 8427 nM | 60 |
| **(33)** | 9 ± 1 µM | 230 nM | 3050 nM | 13 |
| **(34)** | > 1000 | NA | NA | NA |
| **(35)** | > 1000 | NA | NA | NA |
| **(36)** | 25 ± 2 µM | 240 nM | 4200 nM | 18 |
| **(37)** | 5 ± 1 µM | 92 nM | 8427 nM | 92 |
| **(38)** | 2 ± 1 µM | 218 nM | 8427 nM | 39 |
| **(39)** | 17 ± 3 µM | 420 nM | 4200 nM | 7 |
| **(40)** | 2 ± 1 µM | 320 nM | 4200 nM | 13 |
| **(43)** | > 1000 | NA | NA | NA |
| **(46)** | 11 ± 1 µM | 170 nM | 8427 nM | 49 |
| **(49)** | 47 ± 4 µM | NA | NA | NA |
| **(52)** | 94 ± 9 µM | NA | NA | NA |
| **(55)** | > 1000 | NA | NA | NA |

| | | | | |
|---|---|---|---|---|
| **(58)** | 12 ± 1 µM | 450 nM | 2700 nM | 6 |
| **(61)** | 78 ± 7 µM | NA | NA | NA |
| **(64)** | 15 ± 2 µM | 1160 nM | 2277 nM | 2 |
| **(67)** | 14 ± 1 µM | 250 nM | 2700 nM | 11 |
| **(70)** | 2 ± 1 µM | 58 nM | 5514 nM | 95 |
| **(73)** | > 1000 | NA | NA | NA |
| **(74)** | > 1000 | NA | NA | NA |
| **(75)** | > 1000 | NA | NA | NA |
| **(78)** | 2 ± 1 µM | 40 nM | 8427 nM | 210 |
| **(79)** | 2 ± 1 µM | 35 nM | 8427 nM | 240 |
| **(80)** | 9 ± 2 µM | 420 nM | 4600 nM | 11 |
| **(83)** | 5 ± 2 µM | 102 nM | 8427 nM | 83 |
| **(86)** | 1 ± 1 µM | 28 nM | 8427 nM | 300 |
| **(89)** | > 250 | 620 nM | 5514 nM | 9 |

**Table S4.** Percentage (%) bound of compound (1 µM) to 14-3-3σ (100 nM) measured by mass spectrometry in the presence of C-RAF259 peptide (18 µM) or A-RAF214 peptide (16 µM) or B-RAF365 peptide (5 µM) after 1, 8, 16 and 24 hours. Peptide lengths are described in the tables below.

| | **A-RAF214 (15mer)** | | | | **B-RAF365 (10mer)** | | | | **C-RAF259 (11mer)** | | | |
|---|---|---|---|---|---|---|---|---|---|---|---|---|
| **Compound No.** | **1h** | **8h** | **16h** | **24h** | **1h** | **8h** | **16h** | **24h** | **1h** | **8h** | **16h** | **24h** |
| **(8)** | 0 | 0 | 0 | 0 | 0 | 0 | 2.5 | 2.5 | 0 | 0 | 1.9 | 2.9 |
| **(22)** | 54.1 | 75 | 81.4 | 88.5 | 1 | 1.9 | 1.9 | 11.6 | 61 | 85.5 | 94.8 | 98.5 |
| **(23)** | 92 | 95.5 | 97.5 | 97.5 | 0 | 1.4 | 2.4 | 5.6 | 55.6 | 78.5 | 88.9 | 91 |
| **(29)** | 79 | 96.5 | 97 | 97 | 0 | 1.4 | 11.4 | 21.2 | 20.5 | 38.1 | 57.1 | 80.1 |
| **(32)** | 58.2 | 82.1 | 94.3 | 95.2 | 0 | 1 | 3.4 | 15.9 | 14.1 | 21.6 | 39.3 | 74.1 |
| **(37)** | 90.9 | 98 | 99 | 99 | 0 | 4.2 | 7 | 17.3 | 48.7 | 77.1 | 95.2 | 98 |
| **(70)** | 5.2 | 12.6 | 18.5 | 32.3 | 2.4 | 9.9 | 19.3 | 32.2 | 30 | 47.5 | 68 | 87.2 |
| **(78)** | 2.9 | 9 | 15.5 | 28 | 1 | 5.2 | 11.9 | 23 | 8.3 | 25.3 | 35.6 | 70 |
| **(79)** | 4.7 | 8.2 | 15.2 | 23.5 | 0 | 1.9 | 5.1 | 14.1 | 15 | 29.5 | 52.6 | 72 |
| **(80)** | 1.9 | 3.8 | 6.5 | 9 | 1 | 4.7 | 11.4 | 18.6 | 6 | 7 | 15.2 | 25.1 |
| **(83)** | 4.7 | 9.5 | 16.5 | 24.5 | 1.9 | 4.2 | 8.6 | 12.6 | 27 | 52.5 | 66.3 | 84 |
| **(86)** | 11.3 | 21.7 | 35.2 | 44.5 | 1 | 5.2 | 9.9 | 11.8 | 64.6 | 93.1 | 97.1 | 98.5 |

| | **A-RAF214 (15mer)** | | | | **B-RAF365 (15mer)** | | | | **C-RAF259 (15mer)** | | | |
|---|---|---|---|---|---|---|---|---|---|---|---|---|
| **Compound No.** | **1h** | **8h** | **16h** | **24h** | **1h** | **8h** | **16h** | **24h** | **1h** | **8h** | **16h** | **24h** |
| **(8)** | 0 | 0 | 0 | 0 | 0 | 0 | 0.9 | 3.8 | 0 | 0 | 0 | 0 |
| **(22)** | 54.1 | 75 | 81.4 | 88.5 | 39 | 56 | 71 | 83 | 48 | 58.9 | 84.5 | 89.3 |
| **(23)** | 92 | 95.5 | 97.5 | 97.5 | 12.8 | 21.2 | 36.2 | 46.7 | 50 | 66 | 72 | 91 |
| **(29)** | 79 | 96.5 | 97 | 97 | 45.5 | 87.8 | 93.9 | 96.2 | 56.8 | 85.9 | 92.2 | 95.3 |
| **(32)** | 58.2 | 82.1 | 94.3 | 95.2 | 4.7 | 23.1 | 36.4 | 51.6 | 59.7 | 86.7 | 89.9 | 94.4 |
| **(37)** | 90.9 | 98 | 99 | 99 | 32.1 | 69.5 | 81.7 | 92.7 | 64.3 | 91.3 | 92 | 94.4 |
| **(70)** | 5.2 | 12.6 | 18.5 | 32.3 | 1.4 | 2.9 | 3.4 | 4.3 | 2 | 3.8 | 5.6 | 5.6 |
| **(78)** | 2.9 | 9 | 15.5 | 28 | 1.9 | 9.8 | 16.8 | 21.8 | 2 | 4.7 | 14 | 15 |
| **(79)** | 4.7 | 8.2 | 15.2 | 23.5 | 3.4 | 8.3 | 15.2 | 19.2 | 7.4 | 9 | 16.8 | 24.5 |
| **(80)** | 1.9 | 3.8 | 6.5 | 9 | 0 | 1.9 | 2.9 | 4.3 | 1.9 | 1.9 | 3.8 | 5.6 |
| **(83)** | 4.7 | 9.5 | 16.5 | 24.5 | 1.4 | 2.9 | 2.9 | 6.5 | 2.9 | 4.7 | 8.3 | 9.09 |
| **(86)** | 11.3 | 21.7 | 35.2 | 44.5 | 2.9 | 3.4 | 5.2 | 7.3 | 7.4 | 16.7 | 18.5 | 23.1 |

**Table S5.** Protein titrations in the presence of 10 nM FAM-labeled peptides: A-RAF214, B-RAF365 or C-RAF259, 100 µM compound, 250 µM 14-3-3σ starting concentration, 2-fold dilution. Apparent Kd (*app*Kd) values refer to overnight measurements.

| | ***A-RAF 214 (15mer)*** | | ***B-RAF 365 (10mer)*** | | ***C-RAF 259 (10mer)*** | | |
|---|---|---|---|---|---|---|---|
| **No.** | ***App* Kd** | **fold stab.** | ***App* Kd** | **fold stab.** | ***App* Kd** | ***App* Kd DMSO** | **fold stab.** |
| **(8)** | NA | NA | NA | NA | NA | 6700 nM | NA |
| **(22)** | 9 nM | 328 | 180 nM | 4 | 47 nM | 8427 nM | 179 |
| **(23)** | 5 nM | 591 | 105 nM | 6 | 30 nM | 8427 nM | 280 |
| **(29)** | 30 nM | 99 | 140 nM | 5 | 100 nM | 8427 nM | 84 |
| **(32)** | 50 nM | 59 | 112 nM | 6 | 140 nM | 8427 nM | 60 |
| **(37)** | 25 nM | 118 | 52 nM | 13 | 92 nM | 8427 nM | 92 |
| **(70)** | 110 nM | 27 | 41 nM | 15 | 58 nM | 5514 nM | 95 |
| **(78)** | 120 nM | 25 | 41 nM | 15 | 40 nM | 8427 nM | 210 |
| **(79)** | 115 nM | 26 | 100 nM | 6 | 35 nM | 8427 nM | 240 |
| **(80)** | 1100 nM | 3 | 110 nM | 6 | 420 nM | 4600 nM | 11 |
| **(83)** | 265 nM | 11 | 180 nM | 4 | 102 nM | 8427 nM | 83 |
| **(86)** | 140 nM | 21 | 155 nM | 4 | 28 nM | 8427 nM | 300 |
| **DMSO** | 2955 nM | - | 650 nM | - | - | - | - |

| | ***A-RAF 214 (15mer)*** | | ***B-RAF 365 (15mer)*** | | ***C-RAF 259 (15mer)*** | | |
|---|---|---|---|---|---|---|---|
| **No.** | ***App* Kd** | **fold stab.** | ***App* Kd** | **fold stab.** | ***App* Kd** | ***App* Kd DMSO** | **fold stab.** |
| **(8)** | NA | NA | NA | NA | NA | 12000 nM | NA |
| **(22)** | 9 nM | 328 | 5 nM | 448 | 5 nM | 12000 nM | 2400 |
| **(23)** | 5 nM | 591 | 5 nM | 448 | 5 nM | 12000 nM | 2400 |
| **(29)** | 30 nM | 99 | 14 nM | 160 | 154 nM | 12000 nM | 78 |
| **(32)** | 50 nM | 59 | 245 nM | 9 | 126 nM | 12000 nM | 95 |
| **(37)** | 25 nM | 118 | 25 nM | 89 | 136 nM | 12000 nM | 88 |
| **(70)** | 110 nM | 27 | 70 nM | 32 | 100 nM | 12000 nM | 120 |
| **(78)** | 120 nM | 25 | 55 nM | 40 | 110 nM | 12000 nM | 110 |
| **(79)** | 115 nM | 26 | 60 nM | 37 | 135 nM | 12000 nM | 90 |
| **(80)** | 1100 nM | 3 | 220 nM | 10 | 332 nM | 12000 nM | 36 |
| **(83)** | 265 nM | 11 | 94 nM | 24 | 145 nM | 12000 nM | 83 |
| **(86)** | 140 nM | 21 | 95 nM | 24 | 140 nM | 12000 nM | 85 |
| **DMSO** | 2955 nM | - | 2240 nM | - | 12000 nM | - | - |

**Table S6.** SPR experiments of acetylated 12-mer A-RAF, B-RAF, and C-RAF injections (2-fold dilutions) to 14-3-3σ-Twinstrep captured on StrepTactin XT coated chips. Binary interactions and ternary interactions (with compounds **22** and **23**) were measured, resulting in the kinetic parameters, their SE (standard error) and fitting parameters.

| | | A-RAF binary | | | B-RAF binary | | |
|---|---|---|---|---|---|---|---|
| | | Replicate 1 | Replicate 2 | Average ± SD | Replicate 1 | Replicate 2 | Average ± SD |
| Kinetic fit | ka (1/Ms) | 1.32E+05 | 1.32E+05 | 1.3E+05± 4.9E+02 | 1.21E+05 | 1.07E+05 | 1.1E+05± 1.0E+04 |
| | SE (ka) | 1.70E+03 | 1.30E+03 | - | 1.20E+03 | 1.10E+03 | - |
| | kd (1/s) | 0.2444 | 0.2914 | 0.268 ± 0.033 | 0.3206 | 0.2998 | 0.310 ± 0.015 |
| | SE (kd) | 0.0028 | 0.0026 | - | 0.0029 | 0.0029 | - |
| | KD (M) | 1.86E-06 | 2.20E-06 | 2.0E-06± 2.5E-07 | 2.65E-06 | 2.80E-06 | 2.7E-06± 1.1E-07 |
| | Rmax (RU) | 46.8 | 48.6 | - | 45.6 | 45.5 | - |
| | tc | 1.63E+15 | 9.25E+16 | - | 9.57E+15 | 4.48E+14 | - |
| | Chi² (RU²) | 1.37 | 0.73 | - | 0.55 | 0.67 | - |
| | U-value | 5 | 4 | - | 4 | 4 | - |
| Affinity fit | KD (M) | 2.10E-06 | 2.18E-06 | 2.1E-06± 5.7E-08 | 2.53E-06 | 2.76E-06 | 2.6E-06± 1.6E-07 |
| | Rmax (RU) | 45.5 | 48.2 | - | 45.4 | 45.0 | - |

| | | C-RAF binary | | | A-RAF + 23 | | |
|---|---|---|---|---|---|---|---|
| | | Replicate 1 | Replicate 2 | Average ± SD | Replicate 1 | Replicate 2 | Average ± SD |
| Kinetic fit | ka (1/Ms) | 1.14E+05 | 1.60E+05 | 1.4E+05± 3.3E+04 | 1.25E+07 | 1.02E+07 | 1.1E+07± 1.6E+06 |
| | SE (ka) | 1.70E+03 | 1.90E+03 | - | 3.10E+05 | 3.00E+05 | - |
| | kd (1/s) | 0.3201 | 0.3835 | 0.352 ± 0.045 | 0.0189 | 0.0196 | 0.019± 0.001 |
| | SE (kd) | 0.0044 | 0.0043 | - | 4.80E-04 | 6.00E-04 | - |
| | KD (M) | 2.82E-06 | 2.40E-06 | 2.6E-06± 3.0E-07 | 1.51E-09 | 1.92E-09 | 1.7E-09± 2.9E-10 |
| | Rmax (RU) | 41.8 | 43.6 | - | 41.6 | 44.6 | - |
| | tc | 1.05E+16 | 3.30E+17 | - | 2.12E+07 | 1.94E+07 | - |
| | Chi² (RU²) | 0.80 | 0.51 | - | 0.58 | 0.66 | - |
| | U-value | 5 | 5 | - | 5 | 4 | - |
| Affinity fit | KD (M) | 2.61E-06 | 2.14E-06 | 2.4E-06± 3.3E-07 | - | - | - |
| | Rmax (RU) | 41.0 | 43.9 | - | - | - | - |

| | | B-RAF + 23 | | | C-RAF + 23 | | |
|---|---|---|---|---|---|---|---|
| | | Replicate 1 | Replicate 2 | Average ± SD | Replicate 1 | Replicate 2 | Average ± SD |
| Kinetic fit | ka (1/Ms) | 9.53E+06 | 8.53E+06 | 9.0E+06± 7.1E+05 | 1.12E+07 | 1.09E+07 | 1.1E+07± 2.6E+05 |
| | SE (ka) | 2.50E+05 | 1.80E+05 | - | 1.80E+05 | 2.10E+05 | - |
| | kd (1/s) | 0.0572 | 0.0569 | 0.057± 0.000 | 0.0131 | 0.0133 | 0.013± 0.000 |
| | SE (kd) | 0.0015 | 0.0012 | - | 2.00E-04 | 2.60E-04 | - |
| | KD (M) | 6.00E-09 | 6.67E-09 | 6.3E-09± 4.7E-10 | 1.17E-09 | 1.22E-09 | 1.2E-09± 3.9E-11 |
| | Rmax (RU) | 41.5 | 41.3 | - | 42.0 | 42.4 | - |
| | tc | 1.65E+07 | 1.41E+07 | - | 2.90E+07 | 2.55E+07 | - |
| | Chi² (RU²) | 0.24 | 0.22 | - | 0.68 | 0.70 | - |
| | U-value | 4 | 4 | - | 3 | 4 | - |

| | | A-RAF + 22 | | | B-RAF + 22 | | |
|---|---|---|---|---|---|---|---|
| | | Replicate 1 | Replicate 2 | Average ± SD | Replicate 1 | Replicate 2 | Average ± SD |
| Kinetic fit | ka (1/Ms) | 1.41E+07 | 1.70E+07 | 1.6E+07± 2.1E+06 | 8.53E+06 | 6.69E+06 | 7.6E+06± 1.3E+06 |
| | SE (ka) | 3.10E+05 | 4.00E+05 | - | 1.50E+05 | 9.80E+04 | - |
| | kd (1/s) | 0.0562 | 0.0645 | 0.060± 0.006 | 0.0444 | 0.0357 | 0.040± 0.006 |
| | SE (kd) | 0.0012 | 0.0015 | - | 8.00E-04 | 5.00E-04 | - |
| | KD (M) | 3.97E-09 | 3.78E-09 | 3.9E-09± 1.3E-10 | 5.20E-09 | 5.33E-09 | 5.3E-09± 9.2E-11 |
| | Rmax (RU) | 45.1 | 45.5 | - | 41.1 | 41.0 | - |
| | tc | 1.80E+07 | 1.76E+07 | - | 1.75E+07 | 1.84E+07 | - |
| | Chi² (RU²) | 0.14 | 0.12 | - | 0.14 | 0.22 | - |
| | U-value | 4 | 5 | - | 3 | 3 | - |

| | | C-RAF + 22 | | |
|---|---|---|---|---|
| | | Replicate 1 | Replicate 2 | Average ± SD |
| Kinetic fit | ka (1/Ms) | 1.04E+07 | 1.22E+07 | 1.1E+07± 1.3E+06 |
| | SE (ka) | 1.80E+05 | 8.40E+04 | - |
| | kd (1/s) | 0.0396 | 0.0397 | 0.040± 0.000 |
| | SE (kd) | 6.80E-04 | 2.60E-04 | - |
| | KD (M) | 3.82E-09 | 3.26E-09 | 3.5E-09± 4.0E-10 |
| | Rmax (RU) | 43.0 | 44.0 | - |
| | tc | 2.00E+07 | 2.19E+07 | - |
| | Chi² (RU²) | 0.18 | 0.10 | - |
| | U-value | 3 | 2 | - |

**Table S7.** Compound titration follow-up from the selectivity panel. 100 nM FAM-labeled peptides: C-RAF259, SOS1, TAZ, KC1A, TSC2, 14-3-3σ concentration at $EC_{20}$ for each protein-peptide complex, 250 µM compound starting concentration, 3-fold dilution. $EC_{50}$ values refer to overnight measurements.

| **No.** | **C-RAF259** | **SOS1** | **TAZ** | **KC1A** | **TSC2** |
|---|---|---|---|---|---|
| **(22)** | 5 µM | 35.7 µM | 6.2 µM | 87.4 µM | 20.3 µM |
| **(23)** | 5.1 µM | 24.2 µM | 11.7 µM | 38.1 µM | 19.2 µM |

**Table S8.** Protein titration follow-up from the selectivity panel. 10 nM FAM-labeled peptides: C-RAF259, SOS1, TAZ, TSC2, 100 µM compound, 300 µM 14-3-3σ starting concentration, 2-fold dilution. Apparent Kd (*app*Kd) values refer to overnight measurements.

| | **C-RAF259** | | **SOS1** | | **TAZ** | | **TSC2** | |
|---|---|---|---|---|---|---|---|---|
| **No.** | ***App* Kd** | **fold stab.** | ***App* Kd** | **fold stab.** | ***App* Kd** | **fold stab.** | ***App* Kd** | **fold stab.** |
| **(22)** | 400 nM | 28 | 54 µM | 1.5 | 105 nM | 11 | 600 nM | 7 |
| **(23)** | 68 nM | 162 | 21 µM | 4 | 27 nM | 41 | 100 nM | 40 |
| **DMSO** | 11 µM | - | 84 µM | - | 1.1 µM | - | 4 µM | - |

**Table S9.** Cell data for 14-3-3/C-RAF: NanoBRET, co-IP, pS259 protection of phosphorylation in HEK293T and MIA PaCa-2 cells. NA = non-applicable, NT = not tested.

| **No.** | **14-3-3/C-RAF NanoBRET $EC_{50}$ (µM)** | **14-3-3/C-RAF NanoBRET fold stab.** | **14-3-3/C-RAF co-IP fold change** | **C-RAF pS259 protection fold change (HEK293T)** | **C-RAF pS259 protection fold change (MIA PaCa-2)** |
|---|---|---|---|---|---|
| **(22)** | 0.2 | 1.5 | 5.1 | 2.4 | 1.4 |
| **(23)** | 0.18 | 1.6 | 5.3 | 3.2 | 1.5 |
| **(29)** | 0.14 | 1.3 | NT | 2.2 | 1.5 |
| **(32)** | 0.1 | 1.4 | NT | 1.9 | 1.2 |
| **(37)** | NA | 1.3 | NT | 2.4 | 1.4 |
| **(70)** | 0.12 | 1.3 | NT | 2.1 | 1.1 |
| **(78)** | 1.2 | 1.4 | 1.6 | 2.3 | 0.9 |
| **(79)** | NA | 1.5 | NT | 1.3 | 1.0 |
| **(83)** | 1.1 | 1.4 | NT | 1.3 | 0.9 |
| **(86)** | 0.3 | 1.2 | NT | 1.0 | 0.6 |

**Table S10.** Cell data for 14-3-3/A-RAF and 14-3-3/B-RAF: NanoBRET and pS214 protection of phosphorylation in HEK293T cells. NA = non-applicable, NT = not tested.

| **No.** | **14-3-3/A-RAF NanoBRET fold stab.** | **14-3-3/B-RAF NanoBRET fold stab.** | **A-RAF pS214 protection fold change (HEK293T)** |
|---|---|---|---|
| **(22)** | 1.6 | 1.4 | 1.2 |
| **(23)** | 1.7 | 1.4 | 1.3 |
| **(29)** | 1.4 | 1.3 | 1.1 |
| **(32)** | 1.4 | 1.1 | 1.4 |
| **(37)** | 1.5 | 1.3 | 1.2 |
| **(70)** | 1.0 | 1.1 | 1.1 |
| **(78)** | 1.4 | 1.2 | 1.3 |
| **(79)** | 1.5 | 1.2 | 1.3 |
| **(83)** | 1.4 | 1.2 | 1.0 |
| **(86)** | 0.3 | 1.2 | 1.0 |

**Table S11: Crystallography**

| PDB | 8Q5C | 8Q55 | 8QS4 |
|---|---|---|---|
| Protein | 14-3-3σΔC | 14-3-3σΔC | 14-3-3σΔC |
| Peptide | C-RAF pS259 10-mer | C-RAF pS259 10-mer | C-RAF pS259 10-mer |
| Compound | **12 (1075475)** | **21 (1075354)** | **22 (1083853)** |
| Beam | ESRF ID23-1 | ESRF ID23-1 | ESRF ID23-1 |
| *Data collection* | | | |
| Wavelength (Å) | 0.885603 | 0.885603 | 0.885603 |
| Space group | C 2 2 2 | C 2 2 21 | C 2 2 2 |
| Cell dimensions<br>a, b, c (Å)<br>α, β, γ (°) | <br>61.8, 149.4, 76.6<br>90, 90, 90 | <br>82.0, 112.1, 62.5<br>90, 90, 90 | <br>62.3, 149.2, 76.4<br>90, 90, 90 |
| Resolution (Å) | 76.62 – 2.00 (2.05 – 2.00) | 66.16 – 1.30 (1.32 – 1.30) | 76.44 – 2.00 (2.05 – 2.00) |
| $I$ / σ($I$) | 17.6 (1.8) | 14.0 (1.7) | 12.9 (2.4) |
| Completeness (%) | 100.0 (100.0) | 99.8 (99.8) | 98.6 (100.0) |
| Redundancy | 13.5 (14.3) | 11.7 (11.6) | 13.0 (14.2) |
| $CC_{1/2}$ | 0.999 (0.810) | 0.999 (0.693) | 0.998 (0.916) |
| *Refinement* | | | |
| No. reflections | 24411 | 70744 | 24170 |
| $R_{work}/R_{free}$ | 0.215/0.267 | 0.156/0.182 | 0.215/0.248 |
| No. atoms<br>Protein<br>Ligand/ion<br>Water | <br>1881<br>26<br>128 | <br>1933<br>25<br>265 | <br>1885<br>27<br>117 |
| *B*-factors<br>Protein<br>Ligand/ion<br>Water | <br>51.84<br>51.72<br>52.67 | <br>19.95<br>29.57<br>31.73 | <br>48.20<br>52.41<br>49.71 |
| R.m.s. deviations<br>Bond lengths (Å)<br>Bond angles (°) | <br>0.173<br>5.89 | <br>0.019<br>1.41 | <br>0.008<br>1.14 |
| Ramachandran<br>favored (%)<br>outliers (%) | <br>97.83<br>0.00 | <br>97.87<br>0.00 | <br>96.96<br>0.00 |

| **PDB** | **8QS3** | **9EW5** | **8QS2** |
|---|---|---|---|
| Protein | 14-3-3σΔC | 14-3-3σΔC | 14-3-3σΔC |
| Peptide | C-RAF pS259 10-mer | C-RAF pS259 12-mer | C-RAF pS259 10-mer |
| Compound | **23 (1083848)** | **23 (1083848)** | **29 (1076409)** |
| Beam | ESRF ID23-1 | ESRF ID23-2 | ESRF ID23-1 |
| *Data collection* | | | |
| Wavelength (Å) | 0.885603 | 0.873128 | 0.885603 |
| Space group | P 2 21 21 | P 2 21 21 | C 2 2 2 |
| Cell dimensions<br>a, b, c (Å)<br>α, β, γ (°) | <br>60.7, 89.3, 117.0<br>90, 90, 90 | <br>61.07, 90.02, 118.05<br>90, 90, 90 | <br>62.3, 149.0, 76.5<br>90, 90, 90 |
| Resolution (Å) | 117 – 1.60 (1.63 – 1.30) | 61.07 – 1.50 (1.53 – 1.50) | 76.53 – 2.00 (2.05 – 2.00) |
| $I$ / σ($I$) | 16.6 (1.8) | 12.8 (1.2) | 17.2 (1.5) |
| Completeness (%) | 98.2 (100.0) | 100.0 (100.0) | 100.0 (100.0) |
| Redundancy | 13.1 (13.8) | 13.3 (13.0) | 13.5 (14.3) |
| $CC_{1/2}$ | 0.999 (0.727) | 0.999 (0.697) | 0.999 (0.790) |
| *Refinement* | | | |
| No. reflections | 82935 | 104665 | 24495 |
| $R_{work}/R_{free}$ | 0.196/0.215 | 0.198 / 0.222 | 0.203/0.233 |
| No. atoms<br>Protein<br>Ligand/ion<br>Water | <br>3787<br>57<br>552 | <br>3820<br>46<br>437 | <br>1885<br>28<br>111 |
| *B*-factors<br>Protein<br>Ligand/ion<br>Water | <br>31.96<br>34.36<br>39.94 | <br>29.09<br>27.32<br>36.04 | <br>56.05<br>57.21<br>54.60 |
| R.m.s. deviations<br>Bond lengths (Å)<br>Bond angles (°) | <br>0.007<br>1.15 | <br>0.173<br>5.93 | <br>0.173<br>5.80 |
| Ramachandran<br>favored (%)<br>outliers (%) | <br>98.49<br>0.00 | <br>99.14<br>0.00 | <br>98.25<br>0.00 |

| **PDB** | **8QS5** | **8QS6** | **8QS7** |
| --- | --- | --- | --- |
| Protein | 14-3-3σΔC | 14-3-3σΔC | 14-3-3σΔC |
| Peptide | C-RAF pS259 10-mer | C-RAF pS259 10-mer | C-RAF pS259 10-mer |
| Compound | **32 (1083854)** | **37 (1083917)** | **70 (1084352)** |
| Beam | ESRF ID23-1 | ESRF ID23-1 | ESRF ID23-1 |
| *Data collection* | | | |
| Wavelength (Å) | 0.885603 | 0.885603 | 0.885603 |
| Space group | C 2 2 2 | C 2 2 2 | C 2 2 2 |
| Cell dimensions<br>a, b, c (Å)<br>α, β, γ (°) | <br>62.0, 150.6, 75.8<br>90, 90, 90 | <br>62.4, 149.5, 76.5<br>90, 90, 90 | <br>62.4, 149.4, 76.7<br>90, 90, 90 |
| Resolution (Å) | 75.82 – 2.00 (2.05 – 2.00) | 74.73 – 1.80 (1.84 – 1.80) | 76.70 – 1.80 (1.84– 1.80) |
| $I$ / σ($I$) | 15.3 (1.7) | 16.2 (1.2) | 21.9 (1.4) |
| Completeness (%) | 96.6 (100.0) | 93.0 (99.9) | 100.0 (100.0) |
| Redundancy | 13.4 (14.3) | 12.9 (12.7) | 13.3 (12.7) |
| $CC_{1/2}$ | 0.999 (0.867) | 0.998 (0.689) | 0.998 (0.890) |
| *Refinement* | | | |
| No. reflections | 23557 | 31231 | 33604 |
| $R_{work}/R_{free}$ | 0.203/0.231 | 0.239/0.283 | 0.206/0.240 |
| No. atoms<br>Protein<br>Ligand/ion<br>Water | <br>1859<br>30<br>110 | <br>1894<br>27<br>97 | <br>1894<br>27<br>193 |
| *B*-factors<br>Protein<br>Ligand/ion<br>Water | <br>52.70<br>52.90<br>51.40 | <br>35.49<br>51.14<br>49.04 | <br>44.88<br>42.87<br>49.96 |
| R.m.s. deviations<br>Bond lengths (Å)<br>Bond angles (°) | <br>0.008<br>0.87 | <br>0.011<br>1.15 | <br>0.010<br>1.35 |
| Ramachandran<br>favored (%)<br>outliers (%) | <br>97.38<br>0.00 | <br>96.97<br>0.00 | <br>97.84<br>0.00 |

| PDB | 8QS8 | 9EW3 | 9EW1 | 8S42 |
|---|---|---|---|---|
| Protein | 14-3-3σΔC | 14-3-3σΔC | 14-3-3σΔC | 14-3-3σΔC |
| Peptide | C-RAF pS259 10-mer | C-RAF pS259 12-mer | C-RAF pS259 10-mer | C-RAF pS259 10-mer |
| Compound | **78 (1124378)** | **78 (1124378)** | **79 (1124379)** | **80 (1124898)** |
| Beam | ESRF ID23-1 | ESRF ID23-2 | ESRF ID23-1 | ESRF ID30A-3 |
| *Data collection* | | | | |
| Wavelength (Å) | 0.885603 | 0.873128 | 0.885603 | 0.967697 |
| Space group | C 2 2 2 | C 2 2 21 | C 2 2 21 | C 2 2 21 |
| Cell dimensions<br>a, b, c (Å)<br>α, β, γ (°) | <br>62.2, 149.9, 76.6<br>90, 90, 90 | <br>82.54, 112.9, 63.06<br>90, 90, 90 | <br>81.8, 112.14, 62.71<br>90, 90, 90 | <br>81.70, 112.3, 63.09<br>90, 90, 90 |
| Resolution (Å) | 76.55 – 1.80<br>(1.84 – 1.80) | 45.81 – 1.40<br>(1.42 – 1.40) | 66.10 – 1.40<br>(1.42 – 1.40) | 45.63 – 1.70<br>(1.73 – 1.70) |
| $I$ / σ($I$) | 19.0 (1.6) | 11.0 (1.2) | 18.1 (1.7) | 13.6 (2.2) |
| Completeness (%) | 98.3 (100.0) | 99.6 (99.4) | 99.5 (98.4) | 99.8 (99.8) |
| Redundancy | 13.2 (12.7) | 10.1 (8.4) | 13.6 (13.1) | 13.1 (13.5) |
| $CC_{1/2}$ | 0.999 (0.856) | 0.997 (0.639) | 0.999 (0.662) | 0.998 (0.786) |
| *Refinement* | | | | |
| No. reflections | 32982 | 57742 | 56738 | 32194 |
| $R_{work}/R_{free}$ | 0.202/0.252 | 0.208 / 0.239 | 0.158/0.179 | 0.178/0.198 |
| No. atoms<br>Protein<br>Ligand/ion<br>Water | <br>1903<br>25<br>155 | <br>1948<br>27<br>272 | <br>1937<br>26<br>257 | <br>1887<br>29<br>304 |
| *B*-factors<br>Protein<br>Ligand/ion<br>Water | <br>51.17<br>46.26<br>51.30 | <br>19.15<br>29.56<br>29.39 | <br>21.41<br>27.70<br>32.88 | <br>23.52<br>32.18<br>34.81 |
| R.m.s. deviations<br>Bond lengths (Å)<br>Bond angles (°) | <br>0.007<br>1.16 | <br>0.163<br>5.80 | <br>0.105<br>2.06 | <br>0.221<br>6.06 |
| Ramachandran<br>favored (%)<br>outliers (%) | <br>96.55<br>0.00 | <br>98.32<br>0.00 | <br>98.28<br>0.00 | <br>97.37<br>0.00 |

| PDB | 8QS9 | 8QSA | 9EW4 |
|---|---|---|---|
| Protein | 14-3-3σΔC | 14-3-3σΔC | 14-3-3σΔC |
| Peptide | C-RAF pS259 10-mer | C-RAF pS259 10-mer | C-RAF pS259 12-mer |
| Compound | **83 (1124383)** | **86 (1124384)** | **86 (1124384)** |
| Beam | ESRF ID23-1 | ESRF ID23-1 | ESRF ID23-2 |
| *Data collection* | | | |
| Wavelength (Å) | 0.885603 | 0.885603 | 0.873128 |
| Space group | C 2 2 2 | C 2 2 21 | C 2 2 21 |
| Cell dimensions<br>a, b, c (Å)<br>α, β, γ (°) | <br>62.59, 149.56, 76.69<br>90, 90, 90 | <br>62.25, 149.42, 76.76<br>90, 90, 90 | <br>82.18, 112.42 63.13<br>90, 90, 90 |
| Resolution (Å) | 76.69 – 2.00 (2.05 – 2.00) | 76.76 – 1.80 (1.84 – 1.80) | 66.34 – 1.50 (1.53 – 1.50) |
| $I$ / σ($I$) | 17.9 (3.0) | 20.0 (2.3) | 14.7 (1.8) |
| Completeness (%) | 100.0 (100.0) | 98.3 (100.0) | 100.0 (100.0) |
| Redundancy | 13.6 (14.3) | 13.3 (12.7) | 13.4 (13.0) |
| $CC_{1/2}$ | 0.998 (0.867) | 0.998 (0.890) | 0.999 (0.667) |
| *Refinement* | | | |
| No. reflections | 24764 | 32999 | 38384 |
| $R_{work}/R_{free}$ | 0.195/0.228 | 0.198/0.242 | 0.191 / 0.224 |
| No. atoms<br>Protein<br>Ligand/ion<br>Water | <br>1902<br>27<br>184 | <br>1890<br>27<br>171 | <br>1941<br>28<br>221 |
| *B*-factors<br>Protein<br>Ligand/ion<br>Water | <br>44.90<br>43.91<br>47.72 | <br>47.15<br>44.58<br>48.36 | <br>22.26<br>39.46<br>30.66 |
| R.m.s. deviations<br>Bond lengths (Å)<br>Bond angles (°) | <br>0.010<br>0.95 | <br>0.007<br>1.16 | <br>0.018<br>1.56 |
| Ramachandran<br>favored (%)<br>outliers (%) | <br>97.85<br>0.00 | <br>97.82<br>0.00 | <br>97.87<br>0.00 |

| PDB | 9EW7 | 9EW6 | 8VSL |
|---|---|---|---|
| Protein | 14-3-3σΔC | 14-3-3σΔC | 14-3-3σΔC |
| Peptide | C-RAF pS259 12-mer | B-RAF pS365 12-mer | A-RAF pS214 12-mer |
| Compound | Binary | Binary | Binary |
| Beam | Rigaku MicroMax-003 | ESRF ID30B | ESRF ID23-2 |
| *Data collection* | | | |
| Wavelength (Å) | 1.541870 | 0.873128 | 0.873130 |
| Space group | C 2 2 21 | C 2 2 21 | C 2 2 21 |
| Cell dimensions<br>a, b, c (Å)<br>α, β, γ (°) | <br>82.38, 112.20, 62.71<br>90, 90, 90 | <br>81.78, 96.04, 80.44<br>90, 90, 90 | 82.518, 112.806, 62.713<br>90, 90, 90 |
| Resolution (Å) | 34.43 – 1.80 (1.84 – 1.80) | 80.44 – 1.65 (1.68 – 1.65) | 66.60 – 1.42 (1.44 – 1.42) |
| *I* / σ(*I*) | 15.3 (4.5) | 14.4 (1.4) | 11.8 (1.5) |
| Completeness (%) | 100.0 (99.9) | 99.3 (99.9) | 99.9 (99.9) |
| Redundancy | 6.2 (5.5) | 13.5 (14.9) | 13.4 (13.1) |
| $CC_{1/2}$ | 0.998 (0.932) | 0.998 (0.694) | 0.997 (0.536) |
| *Refinement* | | | |
| No. reflections | 27304 | 37337 | 55397 |
| $R_{work}/R_{free}$ | 0.175 / 0.197 | 0.221 / 0.228 | 0.187 / 0.209 |
| No. atoms<br>Protein<br>Ligand/ion<br>Water | <br>1947<br>5<br>306 | <br>1876<br>3<br>205 | <br>1917<br>4<br>307 |
| *B*-factors<br>Protein<br>Ligand/ion<br>Water | <br>12.92<br>21.68<br>22.79 | <br>37.02<br>68.30<br>42.47 | <br>13.77<br>15.07<br>25.75 |
| R.m.s. deviations<br>Bond lengths (Å)<br>Bond angles (°) | <br>0.016<br>1.53 | <br>0.012<br>1.79 | <br>0.041<br>2.04 |
| Ramachandran<br>favored (%)<br>outliers (%) | <br>98.31<br>0.00 | <br>98.68<br>0.00 | <br>98.28<br>0.00 |

| **PDB** | **8QSC** | **8QSH** | **8VSM** |
|---|---|---|---|
| Protein | 14-3-3σΔC | 14-3-3σΔC | 14-3-3σΔC |
| Peptide | A-RAF pS214 12-mer | A-RAF pS214 12-mer | A-RAF pS214 12-mer |
| Compound | **22 (1083853)** | **23 (1083848)** | **78 (1124378)** |
| Beam | ESRF ID30A-3 | ESRF ID23-2 | ESRF ID30A-3 |
| *Data collection* | | | |
| Wavelength (Å) | 0.967697 | 0.873130 | 0.967697 |
| Space group | P 2 21 21 | P 2 21 21 | C 2 2 21 |
| Cell dimensions<br>a, b, c (Å)<br>α, β, γ (°) | <br>60.85, 89.58, 117.55<br>90, 90, 90 | <br>61.84, 90.00, 118.09<br>90, 90, 90 | 82.357, 112.784, 63.163<br>90, 90, 90 |
| Resolution (Å) | 58.78 – 1.80 (1.84 – 1.80) | 90.00 – 1.80 (1.84 – 1.80) | 56.39 – 1.50 (1.53 – 1.50) |
| $I$ / σ($I$) | 25.7 (5.1) | 16.1 (3.8) | 16.4 (2.3) |
| Completeness (%) | 99.6 (99.9) | 99.9 (99.8) | 100.0 (100.0) |
| Redundancy | 13.4 (14.0) | 12.7 (13.9) | 13.7 (14.0) |
| $CC_{1/2}$ | 0.999 (0.953) | 0.877 (0.663) | 0.998 (0.735) |
| *Refinement* | | | |
| No. reflections | 60003 | 61078 | 47385 |
| $R_{work}/R_{free}$ | 0.187 / 0.211 | 0.198 / 0.223 | 0.181 / 0.207 |
| No. atoms<br>Protein<br>Ligand/ion<br>Water | <br>3818<br>49<br>425 | <br>3843<br>51<br>500 | <br>1950<br>4<br>286 |
| *B*-factors<br>Protein<br>Ligand/ion<br>Water | <br>28.20<br>30.07<br>34.28 | <br>21.57<br>22.68<br>28.77 | <br>19.29<br>25.78<br>31.00 |
| R.m.s. deviations<br>Bond lengths (Å)<br>Bond angles (°) | <br>0.018<br>1.61 | <br>0.168<br>5.94 | <br>0.014<br>1.64 |
| Ramachandran<br>favored (%)<br>outliers (%) | <br>99.14<br>0.00 | <br>99.14<br>0.00 | <br>98.30<br>0.00 |

| **PDB** | **8QSB** | **8QSF** | **8QSE** |
|---|---|---|---|
| Protein | 14-3-3σΔC | 14-3-3σΔC | 14-3-3σΔC |
| Peptide | A-RAF pS214 12-mer | B-RAF pS365 12-mer | B-RAF pS365 12-mer |
| Compound | **86 (1124384)** | **22 (1083853)** | **23 (1083848)** |
| Beam | ESRF ID30A-3 | ESRF ID23-2 | ESRF ID23-2 |
| *Data collection* | | | |
| Wavelength (Å) | 0.967697 | 0.873130 | 0.873130 |
| Space group | I 21 21 21 | P 2 21 21 | P 2 21 21 |
| Cell dimensions<br>a, b, c (Å)<br>α, β, γ (°) | <br>62.67, 148.70, 154.5<br>90, 90, 90 | <br>61.00, 88.19, 118.02<br>90, 90, 90 | <br>61.02, 88.81, 117.98<br>90, 90, 90 |
| Resolution (Å) | 53.57 – 1.90 (1.94 -1.90) | 61.00 – 1.80 (1.84 – 1.80) | 70.95 – 1.80<br>(1.84 – 1.80) |
| $I$ / σ($I$) | 16.7 (1.8) | 18.8 (2.8) | 19.0 (2.7) |
| Completeness (%) | 99.9 (100.0) | 99.6 (99.7) | 100.0 (99.9) |
| Redundancy | 13.5 (13.8) | 13.4 (14.0) | 13.5 (14.0) |
| $CC_{1/2}$ | 0.999 (0.741) | 0.995 (0.864) | 0.999 (0.854) |
| *Refinement* | | | |
| No. reflections | 57084 | 59444 | 60129 |
| $R_{work}/R_{free}$ | 0.198 / 0.241 | 0.227 / 0.244 | 0.223 / 0.243 |
| No. atoms<br>Protein<br>Ligand/ion<br>Water | <br>3802<br>56<br>378 | <br>3880<br>51<br>396 | <br>3797<br>51<br>395 |
| *B*-factors<br>Protein<br>Ligand/ion<br>Water | <br>37.62<br>37.59<br>41.89 | <br>19.80<br>31.57<br>34.56 | <br>21.55<br>35.96<br>36.82 |
| R.m.s. deviations<br>Bond lengths (Å)<br>Bond angles (°) | <br>0.011<br>1.73 | <br>0.168<br>5.89 | <br>0.169<br>5.93 |
| Ramachandran<br>favored (%)<br>outliers (%) | <br>99.14<br>0.00 | <br>98.95<br>0.00 | <br>98.92<br>0.00 |

| **PDB** | **8VSO** | **8QSD** | **8QSG** |
|---|---|---|---|
| Protein | 14-3-3σΔC | 14-3-3σΔC | 14-3-3σΔC |
| Peptide | B-RAF pS365 12-mer | B-RAF pS365 12-mer | B-RAF pS365 12-mer |
| Compound | **78 (1124378)** | **79 (1124379)** | **86 (1124384)** |
| Beam | ESRF ID30A-3 | ESRF ID30A-3 | ESRF ID30A-3 |
| *Data collection* | | | |
| Wavelength (Å) | 0.967697 | 0.967697 | 0.967697 |
| Space group | | C 1 2 1 | P 2 21 21 |
| Cell dimensions | 82.18, 112.45, 62.89 | | |
| a, b, c (Å) | 90, 90, 90 | 148.28, 63.62, 73.29 | 60.72, 88.18, 117.18 |
| α, β, γ (°) | | 90, 100.46, 90 | 90, 90, 90 |
| Resolution (Å) | 45.65 – 1.50 (1.53 – 1.50) | 58.31 – 2.00 (2.05 – 2.00) | 58.59 – 2.00 (2.05 – 2.00) |
| $I$ / σ($I$) | 14.3 (1.8) | 9.1 (1.7) | 15.7 (2.5) |
| Completeness (%) | 100.0 (100.0) | 98.5 (99.5) | 99.9 (100.0) |
| Redundancy | 11.8 (12.2) | 6.5 (6.5) | 12.8 (13.1) |
| $CC_{1/2}$ | 0.999 (0.724) | 0.994 (0.800) | 0.999 (0.800) |
| *Refinement* | | | |
| No. reflections | 46940 | 44779 | 43223 |
| $R_{work}/R_{free}$ | 0.182 / 0.198 | 0.219 / 0.241 | 0.198 / 0.243 |
| No. atoms | | | |
| Protein | 1960 | 3753 | 3785 |
| Ligand/ion | 3 | 52 | 54 |
| Water | 217 | 356 | 264 |
| *B*-factors | | | |
| Protein | 22.59 | 38.88 | 38.69 |
| Ligand/ion | 28.06 | 37.41 | 40.27 |
| Water | 30.97 | 42.01 | 39.67 |
| R.m.s. deviations | | | |
| Bond lengths (Å) | 0.014 | 0.160 | 0.011 |
| Bond angles (°) | 1.25 | 5.74 | 1.68 |
| Ramachandran | | | |
| favored (%) | 98.27 | 98.48 | 98.49 |
| outliers (%) | 0.00 | 0.00 | 0.00 |

### 4. SYNTHETIC PROCEDURES

#### General Remarks

All solvents and reagents were commercially available and used without purification, unless otherwise stated. Deuterated solvents were obtained from Cambridge Isotope Laboratories. Reaction progress was monitored by analytical thin-layer chromatography (TLC, pre-coated silica gel 60 F254 plates, Merck) using ultraviolet (UV) light (254 and 365 nm). Analytical liquid chromatography coupled with mass spectrometry (LC-MS) was performed on a C4 Jupiter SuC4300A 150 x 2.0 mm column (using a 15 min. gradient of 5% to 100% acetonitrile in water with 0.1% formic acid), connected to a ThermoFischer LCQ Fleet Ion Trap Mass Spectrometer. Flash column chromatography was performed on Biotage SP1 using silica columns. NMR data were recorded on a Bruker Advance-III 400 MHz equipped with a BBFO probe from Bruker (400 MHz for $^{1}$H-NMR and 100 MHz for $^{13}$C-NMR). Chemical shifts were reported in parts per million (ppm) referenced to an internal standard (*d*-chloroform; 7.26 ppm for $^{1}$H-NMR and 77 ppm for $^{13}$C-NMR), relative to tetramethylsilane (TMS).

**Scheme 1**. General synthetic route for piperidine analogs [a]

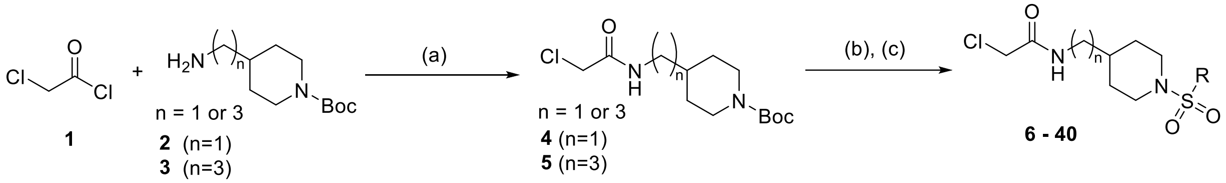

[a] Reagents and conditions: (a) DIPEA, DCM, 0°C to rt, 4h; (b) 4N HCl/dioxane, rt, 3h; (c) appropriate sulfonyl chloride, DIPEA, DCM, 0°C to rt, overnight.

#### General procedures

**Procedure A**: *N*-(*tert*-Butoxycarbonyl)-4-aminomethylpiperidine **2** or 1-Boc-4-(3-aminopropyl)piperidine **3** (1.0 equiv, 3 mmol) was dissolved in 15 ml dry DCM. DIPEA (3 equiv, 9 mmol) was added. The reaction mixture was cooled at 0°C and chloroacetyl chloride **1** (1.2 equiv, 4.2 mmol) was added dropwise. Stirring rt for 4h. The reaction mixture was quenched with sat. $NaHCO_3$ (15 ml) and extracted with DCM (3 x 20 ml). The combined organic phases were dried over $MgSO_4$, filtered and concentrated under reduced pressure. The obtained oil was purified with flash column chromatography [Biotage, hexane – EA, 0-100% EtOAc in hexane].

**Procedure B:** The residue was suspended in 5 ml HCl/dioxane (4N) for Boc-deprotection. Stirring rt for 3h. The solvent was removed under reduced pressure and the obtained oil was used directly in the next step.

**Procedure C:** The obtained HCl salt (1.5 equiv, 0.3 mmol) was suspended in 2 ml dry DCM and cooled at 0 °C. Under stirring, DIPEA (3 equiv, 0.6 mmol) was added. After 10 min, the appropriate sulfonyl chloride (1.0 equiv, 0.2 mmol) was added slowly. Liquid sulfonyl chlorides were added directly the reaction mixture, whereas solid sulfonyl chlorides were dissolved in 1 ml dry DCM and then added to the reaction mixture. Stirring at 0 °C for 30 min and then rt overnight. The reaction mixture was quenched with sat. $NaHCO_3$ (10 ml) and extracted with DCM (3 x 10ml). The combined organic phases were dried over $MgSO_4$, filtered and concentrated under reduced pressure. The obtained crude was purified with flash column chromatography [Biotage, hexane – EA, 0-100% EtOAc in hexane].

*tert-butyl 4-((2-chloroacetamido)methyl)piperidine-1-carboxylate* **(4)**

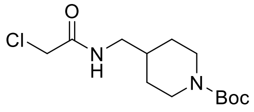

Obtained using procedure A on 3 mmol scale, white solid, 1.8 mmol, 520 mg, 60% yield. $^{1}$H NMR (400 MHz, $CDCl_3$) δ 6.66 (b, 1H), 4.13 – 4.11 (m, 2H), 4.06 (s, 2H), 3.21 (t, *J* = 6.0 Hz, 2H), 2.68 (t, *J* = 12.4 Hz, 2H), 1.72 – 1.69 (m, 3H), 1.45 (s, 9H), 1.14 (qd, *J* = 12.9, 4.7 Hz, 2H). $^{13}$C NMR (100 MHz, $CDCl_3$) δ 166.0, 154.8, 79.5, 45.1, 42.7, 36.2, 29.6, 28.4. LCMS (ESI): m/z calcd for $C_{13}H_{23}ClN_2O_3$; found $[M+Na]^+$ 313.13.

*tert-butyl 4-(3-(2-chloroacetamido)propyl)piperidine-1-carboxylate* **(5)**

Obtained using procedure A on 3 mmol scale, yellow oil, 2.64 mmol, 840 mg, 88% yield. $^{1}$H NMR (400 MHz, $CDCl_3$) δ 6.68 (b, 1H), 3.99 (s, 4H), 3.27 – 3.21 (m, 2H), 2.62 – 2.60 (m, 2H), 1.61 – 1.50 (m, 4H), 1.40 (s, 9H), 1.37 – 1.32 (m, 1H), 1.25 – 1.21 (m, 2H), 1.06 – 1.00 (m, 2H). $^{13}$C NMR (100 MHz, $CDCl_3$) δ 165.8, 154.7, 79.2, 43.8, 42.5, 39.8, 35.5, 33.4, 31.9, 28.3, 26.4. LCMS (ESI): m/z calcd for $C_{15}H_{27}ClN_2O_3$; found $[M+Na]^+$ 341.0.

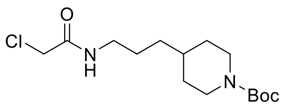

*N-((1-(benzylsulfonyl)piperidin-4-yl)methyl)-2-chloroacetamide* **(6) 1075353**

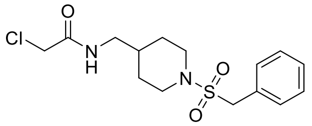

Obtained using procedure C on 0.2 mmol scale, white solid, 0.08 mmol, 29.4 mg, 40% yield. $^{1}$H NMR (400 MHz, $CDCl_3$) δ 7.38 – 7.36 (m, 5H), 6.64 (b, 1H), 4.21 (s, 2H), 4.05 (s, 2H), 3.68 – 3.65 (m, 2H), 3.17 (t, *J* = 6.4 Hz, 2H), 2.54 (td, *J* = 12.4, 2.2 Hz, 2H), 1.67 – 1.61 (m, 2H), 1.24 – 1.14 (m, 3H). $^{13}$C NMR (100 MHz, $CDCl_3$) δ 166.1, 130.6, 128.9, 128.8, 57.1, 45.8, 44.9, 42.6, 35.5, 29.7. LCMS (ESI): m/z calcd for $C_{15}H_{21}ClN_2O_3S$; found $[M+Na]^+$ 367.04.

*2-chloro-N-((1-((4-fluorobenzyl)sulfonyl)piperidin-4-yl)methyl)acetamide* **(7) 1075476**

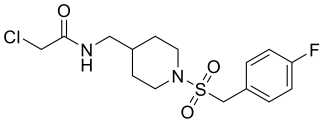

Obtained using procedure C on 0.2 mmol scale, white solid, 0.04 mmol, 15.0 mg, 20% yield. $^{1}$H NMR (400 MHz, $CDCl_3$) δ 7.38 – 7.34 (m, 2H), 7.10 – 7.05 (m, 2H), 6.66 (b, 1H), 4.16 (s, 2H), 4.05 (s, 2H), 3.68 (d, *J* = 12.4 Hz, 2H), 3.18 (t, *J* = 6.4 Hz, 2H), 2.57 (td, *J* = 12.4, 2.3 Hz, 2H), 1.73 – 1.59 (m, 3H), 1.25 – 1.18 (m, 2H). $^{13}$C NMR (100 MHz, $CDCl_3$) δ 166.1, 162.9 (d, *J* = 247.4 Hz), 132.4 (d, *J* = 8.3 Hz), 124.8 (d, *J* = 3.4 Hz), 115.8 (d, *J* = 21.6 Hz), 56.1, 45.9, 44.8, 42.6, 35.5, 29.7. LCMS (ESI): m/z calcd for $C_{15}H_{20}ClFN_2O_3S$; found $[M+Na]^+$ 385.09.

*2-chloro-N-((1-((3-fluorobenzyl)sulfonyl)piperidin-4-yl)methyl)acetamide* **(8) 1075477**

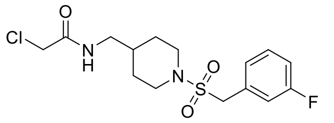

Obtained using procedure C on 0.2 mmol scale, white solid, 0.046 mmol, 16.9 mg, 23.4% yield. $^{1}$H NMR (400 MHz, $CDCl_3$) δ 7.38 – 7.32 (m, 1H), 7.17 – 7.12 (m, 2H), 7.09 – 7.05 (m, 1H), 6.66 (b, 1H), 4.17 (s, 2H), 4.04 (s, 2H), 3.70 – 3.68 (m, 2H), 3.18 (t, *J* = 6.4 Hz, 2H), 2.61 – 2.55 (m, 2H), 1.70 – 1.59 (m, 3H), 1.25 – 1.18 (m, 2H). $^{13}$C NMR (100 MHz, $CDCl_3$) δ 166.0, 162.6 (d, *J* = 246.2 Hz), 131.1 (d, *J* = 7.9 Hz), 130.2 (d, *J* = 8.3 Hz), 126.3 (d, *J* = 3.1 Hz), 117.6 (d, *J* = 17.1 Hz), 115.7 (d, *J* = 20.9 Hz), 56.4, 45.8, 44.7, 42.6, 35.4, 29.6. LCMS (ESI): m/z calcd for $C_{15}H_{20}ClFN_2O_3S$; found $[M+Na]^+$385.04.

*2-chloro-N-((1-((4-chlorobenzyl)sulfonyl)piperidin-4-yl)methyl)acetamide* **(9) 1075352**

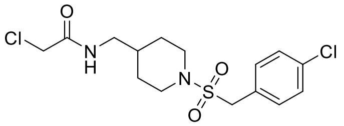

Obtained using procedure C on 0.2 mmol scale, white solid, 0.042 mmol, 16.0 mg, 21.1% yield. $^{1}$H NMR (400 MHz, $CDCl_3$) δ 7.33 (q, *J* = 8.6 Hz, 4H), 6.65 (b, 1H), 4.14 (s, 2H), 4.05 (s, 2H), 3.68 (d, *J* = 12.4 Hz, 2H), 3.18 (t, *J* = 6.4 Hz, 2H), 2.56 (t, J = 12.3 Hz, 2H), 1.69 – 1.63 (m, 2H), 1.24 – 1.17 (m, 3H). $^{13}$C NMR (100 MHz, $CDCl_3$) δ 166.1, 134.9, 131.9, 129.0, 127.5, 56.2, 45.9, 44.8, 42.6, 35.5, 29.7. LCMS (ESI): m/z calcd for $C_{15}H_{20}Cl_2N_2O_3S$; found $[M+Na]^+$ 400.94.

*2-chloro-N-((1-(phenylsulfonyl)piperidin-4-yl)methyl)acetamide* **(10) 1076407**

Obtained using procedure C on 0.2 mmol scale, colorless oil, 0.102 mmol, 33.7 mg, 51% yield. $^{1}$H NMR (400 MHz, $CDCl_3$) δ 7.73 (d, *J* = 7.6 Hz, 2H), 7.60 – 7.50 (m, 3H), 6.69 (b, 1H), 4.00 (s, 2H), 3.80 – 3.78 (m, 2H), 3.16 (t, *J* = 6.4 Hz, 2H), 2.25 (dd, *J* = 16.6, 7.0 Hz, 2H), 1.75 – 1.72 (m, 2H), 1.61 – 1.44 (m, 1H), 1.37 – 1.28 (m, 2H). $^{13}$C NMR (100 MHz, $CDCl_3$) δ 166.0, 136.0, 132.7, 129.0, 127.5, 45.9, 44.6, 42.6, 35.2, 29.0. LCMS (ESI): m/z calcd for $C_{14}H_{19}ClN_2O_3S$; found $[M+H]^+$ 331.13, $[M+Na]^+$ 353.18.

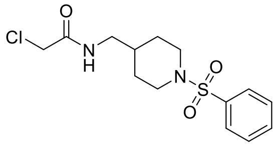

*2-chloro-N-((1-((4-fluorophenyl)sulfonyl)piperidin-4-yl)methyl)acetamide* **(11) 1076408**

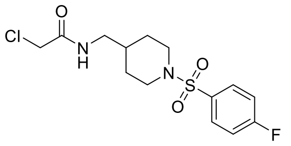

Obtained using procedure C on 0.2 mmol scale, white solid, 0.074 mmol, 26.0 mg, 37% yield. $^{1}$H NMR (400 MHz, $CDCl_3$) δ 7.78 – 7.72 (m, 2H), 7.23 – 7.18 (m, 2H), 6.67 (b, 1H), 4.01 (s, 2H), 3.78 (d, *J* = 11.7 Hz, 2H), 3.17 (t, *J* = 6.5 Hz, 2H), 2.26 (td, *J* = 11.9, 2.3 Hz, 2H), 1.76 – 1.71 (m, 2H), 1.54 – 1.48 (m, 1H), 1.38 – 1.28 (m, 2H).$^{13}$C NMR (100 MHz, $CDCl_3$) δ, 166.06, 165.1 (d, *J* = 254.3 Hz), 132.2 (d, *J* = 3.2 Hz), 130.2 (d, *J* = 9.3 Hz), 116.3 (d, *J* = 22.0 Hz), 45.9, 44.6, 42.6, 35.2, 29.0. LCMS (ESI): m/z calcd for $C_{14}H_{18}ClFN_2O_3S$; found $[M+H]^+$ 349.13, $[M+Na]^+$ 371.14.

*2-chloro-N-((1-((4-chlorophenyl)sulfonyl)piperidin-4-yl)methyl)acetamide* **(12) 1075475**

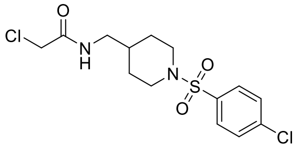

Obtained using procedure C on 0.2 mmol scale, white solid, 0.17 mmol, 61.7 mg, 85.0% yield. $^{1}$H NMR (400 MHz, $CDCl_3$) δ 7.67 (d, *J* = 8.6 Hz, 2H), 7.49 (d, *J* = 8.6 Hz, 2H), 6.67 (b, 1H), 4.02 (s, 2H), 3.78 (d, *J* = 11.7 Hz, 2H), 3.18 (t, *J* = 6.4 Hz, 2H), 2.27 (td, *J* = 11.8, 2.1 Hz, 2H), 1.76 – 1.73 (m, 2H), 1.57 – 1.44 (m, 1H), 1.38 – 1.29 (m, 2H). $^{13}$C NMR (100 MHz, $CDCl_3$) δ 166.0, 139.3, 134.7, 129.3, 129.0, 45.9, 44.6, 42.6, 35.2, 29.0. LCMS (ESI): m/z calcd for $C_{14}H_{18}Cl_2N_2O_3S$; found $[M+H]^+$ 365.04, $[M+Na]^+$ 387.09.

*2-chloro-N-(3-(1-((4-chlorophenyl)sulfonyl)piperidin-4-yl)propyl)acetamide* **(13) 1076390**

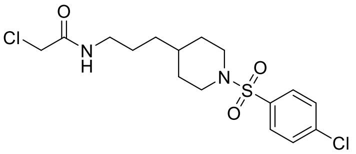

Obtained using procedure C on 0.2 mmol scale, white solid, 0.07 mmol, 26.0 mg, 35.0% yield. $^{1}$H NMR (400 MHz, $CDCl_3$) δ 7.67 (dd, *J* = 8.9, 2.0 Hz, 2H), 7.49 (dd, *J* = 8.9, 2.0 Hz, 2H), 6.59 (b, 1H), 4.01 (s, 2H), 3.75 (d, *J* = 11.7 Hz, 2H), 3.24 (dd, *J* = 13.5, 7.0 Hz, 2H), 2.21 (dd, *J* = 16.6, 6.9 Hz, 2H), 1.71 (d, *J* = 11.7 Hz, 2H), 1.52 – 1.46 (m, 2H), 1.31 – 1.19 (m, 5H).$^{13}$C NMR (100 MHz, $CDCl_3$) δ 165.8, 139.1, 134.7, 129.3, 129.0, 46.3, 42.6, 39.7, 34.6, 32.9, 31.3, 26.4. LCMS (ESI): m/z calcd for $C_{16}H_{22}Cl_2N_2O_3S$; found $[M+H]^+$ 393.14, $[M+Na]^+$ 415.10.

*2-chloro-N-((1-((6-chloropyridin-3-yl)sulfonyl)piperidin-4-yl)methyl)acetamide* **(14) 1076413**

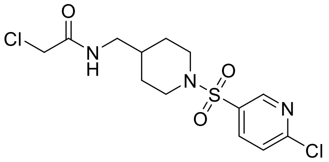

Obtained using procedure C on 0.2 mmol scale, white solid, 0.11 mmol, 40.0 mg, 55.0% yield. $^{1}$H NMR (400 MHz, $CDCl_3$) δ 8.73 (d, *J* = 2.4 Hz, 1H), 7.97 (dd, *J* = 8.3, 2.5 Hz, 1H), 7.49 (d, *J* = 8.3 Hz, 1H), 6.67 (b, 1H), 4.03 (s, 2H), 3.83 (d, *J* = 11.7 Hz, 2H), 3.20 (t, *J* = 6.5 Hz, 2H), 2.35 (td, *J* = 11.9, 2.3 Hz, 2H), 1.80 – 1.77 (m, 2H), 1.59 – 1.52 (m, 1H), 1.40 – 1.30 (m, 2H).$^{13}$C NMR (100 MHz, $CDCl_3$) δ 166.1, 155.6, 148.6, 137.6, 132.2, 124.7, 45.8, 44.5, 42.6, 35.2, 29.0. LCMS (ESI): m/z calcd for $C_{13}H_{17}Cl_2N_3O_3S$; found $[M+H]^+$ 366.09.

*2-chloro-N-((1-((2-fluorophenyl)sulfonyl)piperidin-4-yl)methyl)acetamide* **(15) 1083849**

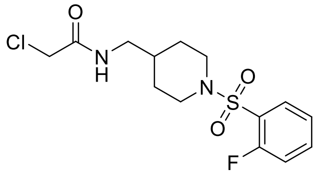

Obtained using procedure C on 0.2 mmol scale, white solid, 0.043 mmol, 15.0 mg, 21.5% yield. $^{1}$H NMR (400 MHz, $CDCl_3$) δ 7.84 (td, *J* = 7.8, 1.8 Hz, 1H), 7.60 – 7.54 (m, 1H), 7.29 – 7.25 (m, 1H), 7.23 – 7.18 (m, 1H), 6.65 (b, 1H), 4.04 (s, 2H), 3.89 (d, *J* = 12.4 Hz, 2H), 3.20 (t, *J* = 6.5 Hz, 2H), 2.55 (t, *J* = 12.2 Hz, 2H), 1.76 (dd, *J* = 12.8, 1.7 Hz, 2H), 1.68 – 1.59 (m, 1H), 1.33 (ddd, *J* = 24.8, 12.2, 4.2 Hz, 2H). $^{13}$C NMR (100 MHz, $CDCl_3$) δ 166.1, 159.0 (d, *J* = 255.0 Hz), 135.0 (d, *J* = 8.4 Hz), 131.2, 125.7 (d, *J* = 14.6 Hz), 124.4 (d, *J* = 3.9 Hz), 117.2 (d, *J* = 22.0 Hz), 45.6, 44.8, 42.6, 35.5, 29.3. LCMS (ESI): m/z calcd for $C_{14}H_{18}ClFN_2O_3S$; found $[M+H]^+$ 349.08.

*2-chloro-N-((1-((3-fluorophenyl)sulfonyl)piperidin-4-yl)methyl)acetamide* **(16) 1083843**

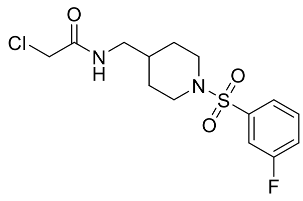

Obtained using procedure C on 0.2 mmol scale, white solid, 0.08 mmol, 28.2 mg, 40.5% yield. $^1$H NMR (400 MHz, $CDCl_3$) δ 7.54 – 7.50 (m, 2H), 7.46 – 7.44 (m, 1H), 7.32 – 7.26 (m, 1H), 6.67 (b, 1H), 4.01 (s, 2H), 3.80 (d, *J* = 11.6 Hz, 2H), 3.18 (t, *J* = 6.4 Hz, 2H), 2.31 (dd, *J* = 16.7, 7.0 Hz, 2H), 1.76 (d, *J* = 12.7 Hz, 2H), 1.58 – 1.50 (m, 1H), 1.38 – 1.31 (m, 2H). $^{13}$C NMR (100 MHz, $CDCl_3$) δ 166.1, 162.4 (d, *J* = 251.0 Hz), 138.3 (d, *J* = 6.0 Hz), 130.8 (d, *J* = 6.0 Hz), 123.3 (d, *J* = 3.0 Hz), 119.9 (d, *J* = 22.0 Hz), 114.8 (d, *J* = 22.0 Hz), 45.9, 44.6, 42.6, 35.2, 29.0. LCMS (ESI): m/z calcd for $C_{14}H_{18}ClFN_2O_3S$; found $[M+H]^+$ 349.08.

*2-chloro-N-((1-((3-(trifluoromethyl)phenyl)sulfonyl)piperidin-4-yl)methyl)acetamide* **(17) 1083844**

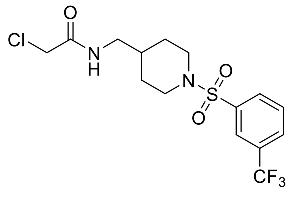

Obtained using procedure C on 0.2 mmol scale, off-white solid, 0.06 mmol, 25.5 mg, 32% yield. $^1$H NMR (400 MHz, $CDCl_3$) δ 8.01 – 8.00 (m, 1H), 7.94 (d, *J* = 7.9 Hz, 1H), 7.86 (d, *J* = 7.8 Hz, 1H), 7.69 (t, *J* = 7.8 Hz, 1H), 6.67 (b, 1H), 4.02 (s, 2H), 3.84 (d, *J* = 11.7 Hz, 2H), 3.18 (t, *J* = 6.5 Hz, 2H), 2.29 (td, *J* = 11.9, 2.5 Hz, 2H), 1.77 (dd, *J* = 12.7, 1.7 Hz, 2H), 1.60 – 1.47 (m, 1H), 1.36 (td, *J* = 12.5, 4.1 Hz, 2H). $^{13}$C NMR (100 MHz, $CDCl_3$) δ 166.1, 137.7, 131.8 (q, *J* = 33.0 Hz), 130.7, 129.9, 129.4 (q, *J* = 3.5 Hz), 124.5 – 124.4 (m), 123.1 (q, *J* = 272.0 Hz), 45.9, 44.6, 42.6, 35.2, 29.0. LCMS (ESI): m/z calcd for $C_{15}H_{18}ClF_3N_2O_3S$; found $[M+H]^+$ 399.10.

*2-chloro-N-((1-((4-(trifluoromethyl)phenyl)sulfonyl)piperidin-4-yl)methyl)acetamide* **(18) 1076412**

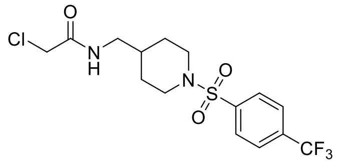

Obtained using procedure C on 0.2 mmol, white solid, 0.065 mmol, 26.2 mg, 33% yield. $^1$H NMR (400 MHz, $CDCl_3$) δ 7.87 (d, *J* = 8.3 Hz, 2H), 7.79 (d, *J* = 8.4 Hz, 2H), 6.67 (b, 1H), 4.02 (s, 2H), 3.83 (d, *J* = 11.8 Hz, 2H), 3.19 (t, *J* = 6.5 Hz, 2H), 2.30 (td, *J* = 11.9, 2.3 Hz, 2H), 1.78 – 1.74 (m, 2H), 1.56 – 1.50 (m, 1H), 1.40 – 1.30 (m, 2H). $^{13}$C NMR (100 MHz, $CDCl_3$) δ 166.1, 139.91, 134.4 (q, *J* = 33.0 Hz),128.0, 126.2 (q, *J* = 3.6 Hz),123.2 (q, *J* = 271.9 Hz), 45.9, 44.6, 42.6, 35.2, 29.0. LCMS (ESI): m/z calcd for $C_{15}H_{18}ClF_3N_2O_3S$; found $[M+H]^+$ 399.20, $[M+Na]^+$ 421.15.

*2-chloro-N-((1-((4-(trifluoromethoxy)phenyl)sulfonyl)piperidin-4-yl)methyl)acetamide* **(19) 1083842**

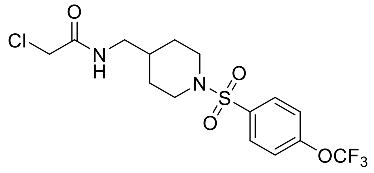

Obtained using procedure C on 0.2 mmol scale, white solid, 0.042 mmol, 17.6 mg, 22% yield. $^1$H NMR (400 MHz, $CDCl_3$) δ 7.81 – 7.79 (m, 2H), 7.35 (d, *J* = 8.1 Hz, 2H), 6.65 (b, 1H), 4.02 (s, 2H), 3.81 (d, *J* = 11.8 Hz, 2H), 3.19 (t, *J* = 6.5 Hz, 2H), 2.30 (td, *J* = 11.9, 2.4 Hz, 2H), 1.76 (dd, *J* = 12.7, 1.7 Hz, 2H), 1.57 – 1.51 (m, 1H), 1.35 (ddd, *J* = 24.7, 12.2, 4.1 Hz, 2H). $^{13}$C NMR (100 MHz, $CDCl_3$) δ 166.1, 152.2, 134.7, 129.7, 120.9, 120.2 (q, *J* = 258.0 Hz), 45.9, 44.7, 42.6, 35.3, 29.0. LCMS (ESI): m/z calcd for $C_{15}H_{18}ClF_3N_2O_4S$; found $[M+H]^+$ 414.15.

*2-chloro-N-((1-((4-cyanophenyl)sulfonyl)piperidin-4-yl)methyl)acetamide* **(20) 1083841**

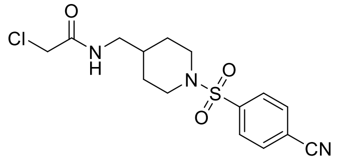

Obtained using procedure C on 0.2 mmol scale, white solid, 0.068 mmol, 24.3 mg, 35% yield. $^1$H NMR (400 MHz, $CDCl_3$) δ 7.87 – 7.81 (m, 4H), 6.67 (b, 1H), 4.02 (s, 2H), 3.82 (d, *J* = 11.8 Hz, 2H), 3.19 (t, *J* = 6.5 Hz, 2H), 2.33 (td, *J* = 11.9, 2.5 Hz, 2H), 1.78 – 1.75 (m, 2H), 1.56 – 1.50 (m, 1H), 1.39 – 1.29 (m, 2H). $^{13}$C NMR (100 MHz, $CDCl_3$) δ 166.1, 140.9, 132.8, 128.1, 117.2, 116.5, 45.8, 44.5, 42.6, 35.2, 29.0. LCMS (ESI): m/z calcd for $C_{15}H_{18}ClN_3O_3S$; found $[M+H]^+$ 356.14.

*2-chloro-N-((1-((4-formylphenyl)sulfonyl)piperidin-4-yl)methyl)acetamide* **(21) 1075354**

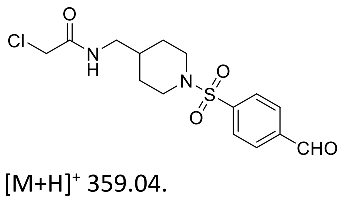

Obtained using procedure C on 0.2 mmol scale, white solid, 0.056 mmol, 20.4 mg, 28.0% yield. $^1$H NMR (400 MHz, $CDCl_3$) δ 10.11 (s, 1H), 8.04 (d, *J* = 8.2 Hz, 2H), 7.92 (d, *J* = 8.2 Hz, 2H), 6.65 (s, 1H), 4.02 (s, 2H), 3.85 (d, *J* = 11.8 Hz, 2H), 3.18 (t, *J* = 6.4 Hz, 2H), 2.42 – 2.24 (m, 2H), 1.76 (d, *J* = 11.9 Hz, 2H), 1.59 – 1.46 (m, 1H), 1.35 (dd, *J* = 12.3, 3.6 Hz, 2H). $^{13}$C NMR (100 MHz, $CDCl_3$) δ 190.8, 166.1, 141.6, 138.9, 130.1, 128.2, 45.9, 44.6, 42.6, 35.2, 29.0. LCMS (ESI): m/z calcd for $C_{15}H_{19}ClN_2O_4S$; found $[M+H]^+$ 359.04.

*2-chloro-N-((1-((4-bromo-phenyl)sulfonyl)piperidin-4-yl)methyl)acetamide* **(22) 1083853**

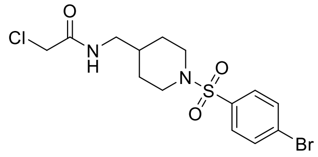

Obtained using procedure C on 0.2 mmol scale, white solid, 0.11 mmol, 45.8 mg, 56% yield. $^1$H NMR (400 MHz, $CDCl_3$) δ 7.65 (d, *J* = 8.5 Hz, 2H), 7.58 (d, *J* = 8.5 Hz, 2H), 6.69 (b, 1H), 4.00 (s, 2H), 3.76 (d, *J* = 11.7 Hz, 2H), 3.16 (t, *J* = 6.4 Hz, 2H), 2.25 (dd, *J* = 11.7, 10.2 Hz, 2H), 1.75 – 1.72 (m, 2H), 1.58 – 1.42 (m, 1H), 1.33 (ddd, *J* = 24.7, 12.2, 4.0 Hz, 2H). $^{13}$C NMR (100 MHz, $CDCl_3$) δ 166.0, 135.3, 132.3, 129.1, 127.8, 45.9, 44.6, 42.6, 35.2, 29.0. LCMS (ESI): m/z calcd for $C_{14}H_{18}BrClN_2O_3S$; found $[M+H]^+$ 410.90.

*2-chloro-N-((1-((4-iodophenyl)sulfonyl)piperidin-4-yl)methyl)acetamide* **(23) 1083848**

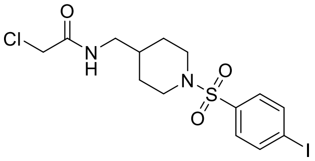

Obtained using procedure C on 0.2 mmol scale, white solid, 0.056 mmol, 25.5 mg, 28.0% yield. $^1$H NMR (400 MHz, $CDCl_3$) δ 7.89 – 7.86 (m, 2H), 7.46 – 7.43 (m, 2H), 6.67 (b, 1H), 4.02 (b, 2H), 3.79 – 3.75 (m, 2H), 3.18 (t, *J* = 6.5 Hz, 2H), 2.27 (t, *J* = 11.8 Hz, 2H), 1.74 (d, *J* = 12.9 Hz, 2H), 1.52 – 1.49 (m, 1H), 1.33 (ddd, *J* = 24.7, 12.2, 4.1 Hz, 2H).$^{13}$C NMR (100 MHz, $CDCl_3$) δ 166.0, 138.3, 135.9, 129.0, 100.2, 45.9, 44.6, 42.6, 35.3, 29.0. LCMS (ESI): m/z calcd for $C_{14}H_{18}ClIN_2O_3S$; found $[M+H]^+$ 457.01.

*2-chloro-N-((1-tosylpiperidin-4-yl)methyl)acetamide* **(24) 1083852**

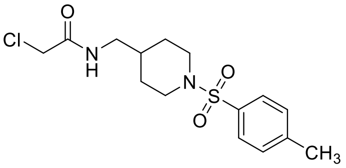

Obtained using procedure C on 0.2 mmol scale, white solid, 0.17 mmol, 58.9 mg, 85.6% yield. $^1$H NMR (400 MHz, $CDCl_3$) δ 7.59 (d, *J* = 8.3 Hz, 2H), 7.29 (d, *J* = 8.0 Hz, 2H), 6.71 (b, 1H), 3.98 (s, 2H), 3.74 (d, *J* = 12.0 Hz, 2H), 3.14 (t, *J* = 6.5 Hz, 2H), 2.40 (s, 3H), 2.21 (td, *J* = 11.8, 2.4 Hz, 2H), 1.71 (dd, *J* = 12.7, 1.9 Hz, 2H), 1.59 – 1.41 (m, 1H), 1.30 (ddd, *J* = 24.6, 12.2, 4.0 Hz, 2H). $^{13}$C NMR (100 MHz, $CDCl_3$) δ 166.0, 143.5, 132.9, 129.5, 127.5, 45.8, 44.6, 42.5, 35.1, 28.9, 21.4. LCMS (ESI): m/z calcd for $C_{15}H_{21}ClN_2O_3S$; found $[M+H]^+$ 345.08.

*2-chloro-N-((1-((4-methoxyphenyl)sulfonyl)piperidin-4-yl)methyl)acetamide* **(25) 1076411**

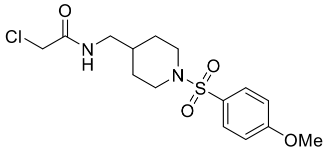

Obtained using procedure C on 0.2 mmol, colorless oil, 0.056 mmol, 20.2 mg, 28% yield. $^1$H NMR (400 MHz, $CDCl_3$) δ 7.70 – 7.66 (m, 2H), 7.01 – 6.97 (m, 2H), 6.67 (b, 1H), 4.02 (s, 2H), 3.87 (s, 3H), 3.78 – 3.76 (m, 2H), 3.20 – 3.16 (m, 2H), 2.27 – 2.24 (m, 2H), 1.75 – 1.67 (m, 2H), 1.52 – 1.50 (m, 1H), 1.38 – 1.32 (m, 2H).$^{13}$C NMR (100 MHz, $CDCl_3$) δ 166.0, 162.9, 129.7, 127.6, 114.1, 55.6, 45.9, 44.7, 42.6, 35.3, 29.0. LCMS (ESI): m/z calcd for $C_{15}H_{21}ClN_2O_4S$; found $[M+H]^+$ 361.09.

*N-((1-(benzo[d][1,3]dioxol-5-ylsulfonyl)piperidin-4-yl)methyl)-2-chloroacetamide* **(26) 1124380**

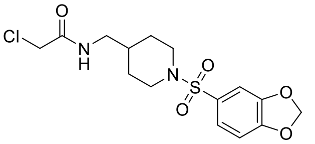

Obtained using procedure C on 0.6 mmol scale, white solid, 0.12 mmol, 46.1 mg, 20.0 % yield. $^1$H NMR (400 MHz, $CDCl_3$) δ 7.28 (dd, *J* = 8.2, 1.8 Hz, 1H), 7.13 (d, *J* = 1.8 Hz, 1H), 6.88 (d, *J* = 8.2 Hz, 1H), 6.72 (b, 1H), 6.07 (s, 2H), 4.01 (s, 2H), 3.73 (d, *J* = 11.7 Hz, 2H), 3.16 (t, *J* = 6.5 Hz, 2H), 2.26 (td, *J* = 11.8, 2.4 Hz, 2H), 1.73 (dd, *J* = 12.7, 1.7 Hz, 2H), 1.55 – 1.50 (m, 1H), 1.32 (ddd, *J* = 24.6, 12.2, 4.0 Hz, 2H). $^{13}$C NMR (100 MHz, $CDCl_3$) δ 166.1, 151.3, 148.1, 129.2, 123.2, 108.2, 107.7, 102.3, 45.9, 44.6, 42.6, 35.2, 28.9. LCMS (ESI): m/z calcd for $C_{15}H_{19}ClN_2O_5S$; found $[M+H]^+$ 375.17.

*2-chloro-N-((1-((2,4-difluorophenyl)sulfonyl)piperidin-4-yl)methyl)acetamide* **(27) 1083846**

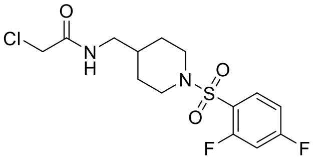

Obtained using procedure C on 0.2 mmol scale, white solid, 0.062 mmol, 22.9 mg, 31.2% yield. $^1$H NMR (400 MHz, $CDCl_3$) δ 7.88 – 7.82 (m, 1H), 7.02 – 6.92 (m, 2H), 6.68 (b, 1H), 4.03 (s, 2H), 3.86 (d, *J* = 12.3 Hz, 2H), 3.20 (t, *J* = 6.5 Hz, 2H), 2.54 (t, *J* = 12.2 Hz, 2H), 1.76 (dd, *J* = 12.8, 1.8 Hz, 2H), 1.71 – 1.54 (m, 1H), 1.32 (ddd, *J* = 24.8, 12.2, 4.2 Hz, 2H). $^{13}$C NMR (100 MHz, $CDCl_3$) δ 166.1, 165.6 (dd, *J* = 256, 11.4 Hz), 159.7 (dd, *J* = 257, 12.7 Hz), 132.9 (dd, *J* = 10.4, 2.1 Hz), 122.2 (d, *J* = 14.9 Hz), 111.9 (dd, *J* = 21.8, 3.8 Hz), 105.7 (t, *J* = 26.0 Hz), 45.5, 44.7, 42.6, 35.4, 29.2. LCMS (ESI): m/z calcd for $C_{14}H_{17}ClF_2N_2O_3S$; found $[M+H]^+$ 367.09.

*2-chloro-N-((1-((3,4-difluorophenyl)sulfonyl)piperidin-4-yl)methyl)acetamide* **(28) 1083845**

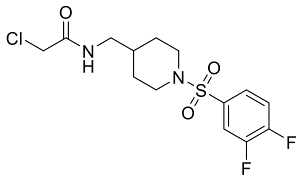

Obtained using procedure C on 0.2 mmol scale, white solid, 0.057 mmol, 21.1 mg, 29.0% yield. $^{1}$H NMR (400 MHz, $CDCl_3$) δ 7.59 (ddd, *J* = 9.3, 7.2, 2.2 Hz, 1H), 7.55 – 7.50 (m, 1H), 7.36 – 7.29 (m, 1H), 6.67(b, 1H), 4.02 (s, 2H), 3.79 (d, *J* = 11.8 Hz, 2H), 3.19 (t, *J* = 6.5 Hz, 2H), 2.31 (td, *J* = 11.9, 2.5 Hz, 2H), 1.77 (dd, *J* = 12.7, 1.8 Hz, 2H), 1.53 (dtt, *J* = 13.8, 6.7, 3.6 Hz, 1H), 1.34 (ddd, *J* = 24.8, 12.2, 4.1 Hz, 2H). $^{13}$C NMR (100 MHz, CDCl3) δ 166.1, 153.1 (dd, *J* = 256, 12.5 Hz), 150.1 (dd, *J* = 254, 13.3 Hz), 133.2 (t, *J* = 4.3 Hz), 124.6 (dd, *J* = 7.4, 4.1 Hz), 118.2 (d, *J* = 18.4 Hz), 117.4 (d, *J* = 19.6 Hz), 45.9, 44.6 42.6, 35.2, 29.0. LCMS (ESI): m/z calcd for $C_{14}H_{17}ClF_2N_2O_3S$; found $[M+H]^+$ 367.14.

*2-chloro-N-((1-((4-chloro-3-fluorophenyl)sulfonyl)piperidin-4-yl)methyl)acetamide* **(29) 1076409**

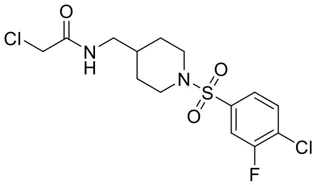

Obtained using procedure C on 0.2 mmol, white solid, 0.067 mmol, 25.7 mg, 33.5% yield. $^{1}$H NMR (400 MHz, $CDCl_3$) δ 7.59 – 7.46 (m, 3H), 6.67 (b, 1H), 4.02 (s, 2H), 3.79 (d, *J* = 11.8 Hz, 2H), 3.19 (t, *J* = 6.5 Hz, 2H), 2.32 (td, *J* = 11.9, 2.3 Hz, 2H), 1.76 (d, *J* = 11.2 Hz, 2H), 1.62 – 1.50 (m, 1H), 1.39 – 1.30 (m, 2H). $^{13}$C NMR (100 MHz, $CDCl_3$) δ 166.1, 157.8 (d, *J* = 253.0 Hz), 136.6 (d, *J* = 5.5 Hz), 131.5, 126.4 (d, *J* = 17.7 Hz), 123.9 (d, *J* = 4.2 Hz), 116.0 (d, *J* = 24.0 Hz), 45.9, 44.6, 42.6, 35.2, 29.0. LCMS (ESI): m/z calcd for $C_{14}H_{17}Cl_2FN_2O_3S$; found $[M+H]^+$ 383.09, $[M+Na]^+$ 405.20.

*2-chloro-N-((1-((3-chloro-4-fluorophenyl)sulfonyl)piperidin-4-yl)methyl)acetamide* **(30) 1076410**

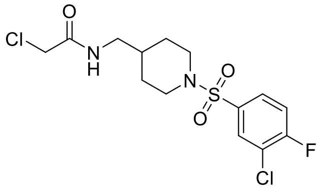

Obtained using procedure C on 0.2 mmol, white solid, 0.078 mmol, 30.0 mg, 39.0% yield. $^{1}$H NMR (400 MHz, $CDCl_3$) δ 7.82 (dd, *J* = 6.7, 2.2 Hz, 1H), 7.64 (ddd, *J* = 8.6, 4.3, 2.2 Hz, 1H), 7.29 (t, *J* = 8.5 Hz, 1H), 6.67 (b, 1H), 4.02 (s, 2H), 3.80 (d, *J* = 11.7 Hz, 2H), 3.19 (t, *J* = 6.5 Hz, 2H), 2.31 (td, *J* = 11.9, 2.4 Hz, 2H), 1.78 – 1.74 (m, 2H), 1.56 – 1.51 (m, 1H), 1.40 – 1.30 (m, 2H).$^{13}$C NMR (100 MHz, $CDCl_3$) δ 166.1, 160.6 (d, *J* = 256.1 Hz), 133.4 (d, *J* = 4.0 Hz), 130.3, 128.0 (d, *J* = 8.4 Hz), 122.5 (d, *J* = 18.6 Hz), 117.3 (d, *J* = 22.1 Hz), 45.9, 44.6, 42.6, 35.2, 29.0. LCMS (ESI): m/z calcd for $C_{14}H_{17}Cl_2FN_2O_3S$; found $[M+H]^+$ 383.14.

*2-chloro-N-((1-((4-chloro-3-(trifluoromethyl)phenyl)sulfonyl)piperidin-4-yl)methyl)acetamide* **(31) 1083850**

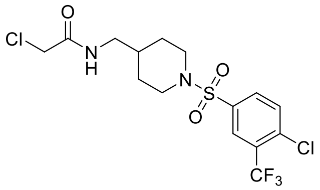

Obtained using procedure C on 0.2 mmol scale, white solid, 0.11 mmol, 49.4 mg, 57% yield. $^{1}$H NMR (400 MHz, $CDCl_3$) δ 8.03 (d, *J* = 2.0 Hz, 1H), 7.84 (dd, *J* = 8.4, 2.1 Hz, 1H), 7.68 (d, *J* = 8.4 Hz, 1H), 6.68 (b, 1H), 4.01 (s, 2H), 3.81 (d, *J* = 11.7 Hz, 2H), 3.19 (t, *J* = 6.5 Hz, 2H), 2.31 (td, *J* = 11.9, 2.4 Hz, 2H), 1.78 (dd, *J* = 12.8, 1.7 Hz, 2H), 1.60 – 1.50 (m, 1H), 1.35 (ddd, *J* = 24.8, 12.2, 4.1 Hz, 2H). $^{13}$C NMR (100 MHz, $CDCl_3$) δ 166.1, 137.3 (d, *J* = 1.5 Hz), 135.7, 132.4, 131.6, 129.44 (q, *J* = 32.4 Hz), 126.68 (q, *J* = 5.4 Hz), 121.93 (q, *J* = 274.1 Hz), 45.8, 44.5, 42.6, 35.2, 29.0. LCMS (ESI): m/z calcd for $C_{15}H_{17}Cl_2F_3N_2O_3S$; found $[M+H]^+$ 433.06.

*2-chloro-N-((1-((3-fluoro-4-(trifluoromethyl)phenyl)sulfonyl)piperidin-4-yl)methyl)acetamide* **(32) 1083854**

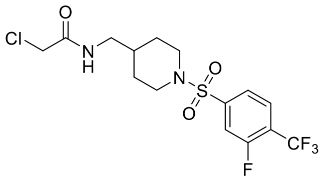

Obtained using procedure C on 0.2 mmol scale, white solid, 0.12 mmol, 51.6 mg, 62.0% yield. $^{1}$H NMR (400 MHz, $CDCl_3$) δ 7.81 – 7.77 (m, 1H), 7.62 – 7.58 (m, 2H), 6.67 (b, 1H), 4.02 (s, 2H), 3.82 (d, *J* = 11.8 Hz, 2H), 3.19 (t, *J* = 6.5 Hz, 2H), 2.36 (td, *J* = 11.9, 2.4 Hz, 2H), 1.78 (dd, *J* = 12.8, 1.7 Hz, 2H), 1.61 – 1.50 (m, 1H), 1.35 (qd, *J* = 12.3, 4.1 Hz, 2H). $^{13}$C NMR (100 MHz, $CDCl_3$) δ 166.1, 159.5 (d, *J* = 264.3 Hz), 142.5 (d, *J* = 7.1 Hz), 128.3 (d, *J* = 4.5 Hz), 123.1 (d, *J* = 4.3 Hz), 122.3 (dd, *J* = 33.7, 12.5 Hz), 120.3, 116.3 (d, *J* = 23.4 Hz), 45.9, 44.5, 42.6, 35.2, 29.0. LCMS (ESI): m/z calcd for $C_{15}H_{17}ClF_4N_2O_3S$; found $[M+H]^+$ 417.15.

*2-chloro-N-((1-((3-fluoro-4-methoxyphenyl)sulfonyl)piperidin-4-yl)methyl)acetamide* **(33) 1083847**

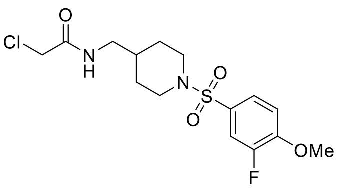

Obtained using procedure C on 0.2 mmol scale, white solid, 0.047 mmol, 18.0 mg, 24.0% yield. $^1$H NMR (400 MHz, $CDCl_3$) δ 7.52 – 7.50 (m, 1H), 7.48 – 7.45 (m, 1H), 7.05 (t, $J$ = 8.2 Hz, 1H), 6.64 (b, 1H), 4.03 (s, 2H), 3.96 (s, 3H), 3.78 (d, $J$ = 11.7 Hz, 2H), 3.19 (t, $J$ = 6.5 Hz, 2H), 2.29 (td, $J$ = 11.8, 2.5 Hz, 2H), 1.76 (dd, $J$ = 12.7, 1.9 Hz, 2H), 1.56 – 1.45 (m, 1H), 1.35 (ddd, $J$ = 24.6, 12.1, 4.1 Hz, 2H). $^{13}$C NMR (100 MHz, $CDCl_3$) δ 166.0, 151.7 (d, $J$ = 251.0 Hz), 151.5 (d, $J$ = 10.5 Hz), 128.2 (d, $J$ = 5.4 Hz), 124.8 (d, $J$ = 3.8 Hz), 115.7 (d, $J$ = 21.0 Hz), 112.9 (d, $J$ = 2.0 Hz), 56.4, 45.9, 44.7, 42.6, 35.3, 29.0. LCMS (ESI): m/z calcd for $C_{15}H_{20}ClFN_2O_4S$; found $[M+H]^+$ 379.14.

*2-chloro-N-((1-((1,1-difluorospiro[2.5]octan-6-yl)sulfonyl)piperidin-4-yl)methyl)acetamide* **(34) 1083851**

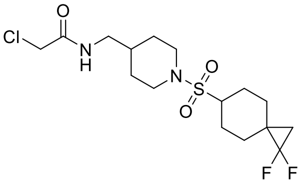

Obtained using procedure C on 0.2 mmol scale, white solid, 0.02 mmol, 9.3 mg, 10% yield. $^1$H NMR (400 MHz, $CDCl_3$) δ 6.68 (b, 1H), 4.07 (s, 2H), 3.86 – 3.83 (m, 2H), 3.24 (t, $J$ = 6.4 Hz, 2H), 2.95 – 2.84 (m, 3H), 2.17 – 2.14 (m, 2H), 1.78 – 1.60 (m, 8H), 1.35 – 1.29 (m, 3H), 1.05 (t, $J$ = 8.3 Hz, 2H). $^{13}$C NMR (100 MHz, $CDCl_3$) δ 166.1, 115.9, 60.4, 46.0, 44.9, 42.7, 35.8, 30.1, 27.9 (t, $J$ = 11.1 Hz), 27.4, 24.4, 21.7 (t, $J$ = 10.1 Hz). LCMS (ESI): m/z calcd for $C_{16}H_{25}ClF_2N_2O_3S$; found $[M+H]^+$ 399.05.

*2-chloro-N-((1-((6,6-difluorospiro[3.3]heptan-2-yl)sulfonyl)piperidin-4-yl)methyl)acetamide* **(35) 1083855**

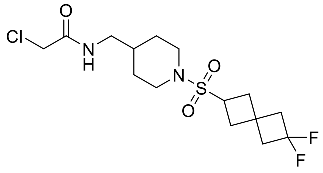

Obtained using procedure C on 0.20 mmol scale, white solid, 0.014 mmol, 5.4 mg, 7.1% yield. $^1$H NMR (400 MHz, $CDCl_3$) δ 6.67 (b, 1H), 4.06 (s, 2H), 3.79 (d, $J$ = 12.3 Hz, 2H), 3.68 – 3.63 (m, 1H), 3.23 (t, $J$ = 6.5 Hz, 2H), 2.76 – 2.58 (m, 8H), 2.44 – 2.38 (m, 2H), 1.78 – 1.75 (m, 2H), 1.72 – 1.65 (m, 1H), 1.28 (ddd, $J$ = 15.9, 12.2, 4.2 Hz, 2H). $^{13}$C NMR (100 MHz, $CDCl_3$) δ 166.1, 118.8 (t, $J$ =279.4 Hz), 47.6 (t, $J$ = 22.4 Hz), 46.94 (t, $J$ = 22.4 Hz), 45.9, 44.8, 42.7, 35.9, 35.6, 29.7, 28.7 (t, $J$ = 8.6 Hz). LCMS (ESI): m/z calcd for $C_{15}H_{23}ClF_2N_2O_3S$; $[M+H]^+$ 385.19.

*N-((1-((4-bromo-2-fluorophenyl)sulfonyl)piperidin-4-yl)methyl)-2-chloroacetamide* **(36) 1083916**

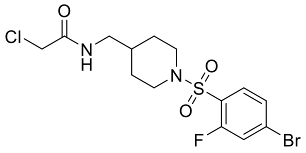

Obtained using procedure C on 0.2 mmol scale, white solid, 0.04 mmol, 17.1 mg, 20.0 % yield. $^1$H NMR (400 MHz, $CDCl_3$) δ 7.70 (t, $J$ = 7.9 Hz, 1H), 7.44 – 7.39 (m, 2H), 6.67 (b, 1H), 4.04 (s, 2H), 3.89 – 3.86 (m, 2H), 3.21 (t, $J$ = 6.5 Hz, 2H), 2.56 (t, $J$ = 12.2 Hz, 2H), 1.78 – 1.75 (m, 2H), 1.61 – 1.59 (m, 1H), 1.32 (ddd, $J$ = 24.7, 12.2, 4.1 Hz, 2H). $^{13}$C NMR (100 MHz, $CDCl_3$) δ 166.1, 158.6 (d, $J$ = 260.1 Hz), 132.1, 128.3 (d, $J$ = 9.1 Hz), 127.9 (d, $J$ = 3.8 Hz), 125.1 (d, $J$ = 14.9 Hz), 120.9 (d, $J$ = 25.1 Hz), 45.5, 44.7, 42.6, 35.41, 29.3. LCMS (ESI): m/z calcd for $C_{14}H_{17}BrClFN_2O_3S$; found $[M+H]^+$ 427.05.

*N-((1-((4-bromo-3-fluorophenyl)sulfonyl)piperidin-4-yl)methyl)-2-chloroacetamide* **(37) 1083917**

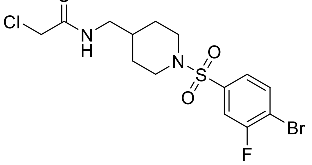

Obtained using procedure C on 0.2 mmol scale, white solid, 0.056 mmol, 24 mg, 28.1 % yield. $^1$H NMR (400 MHz, $CDCl_3$) δ $^1$H NMR (400 MHz, $CDCl_3$) δ 7.74 (dd, $J$ = 8.3, 6.5 Hz, 1H), 7.50 (dd, $J$ = 7.7, 2.0 Hz, 1H), 7.43 – 7.40 (m, 1H), 6.64 (b, 1H), 4.03 (s, 2H), 3.81 (d, $J$ = 11.8 Hz, 2H), 3.20 (t, $J$ = 6.5 Hz, 2H), 2.33 (td, $J$ = 11.9, 2.5 Hz, 2H), 1.77 (dd, $J$ = 12.8, 1.8 Hz, 2H), 1.54 – 1.51 (m, 1H), 1.35 (ddd, $J$ = 24.7, 12.1, 4.1 Hz, 2H). $^{13}$C NMR (100 MHz, $CDCl_3$) δ 166.1, 158.9 (d, $J$ = 253.1 Hz), 137.6 (d, $J$ = 5.6 Hz), 134.4, 124.2 (d, $J$ = 4.1 Hz), 115.7 (d, $J$ = 25.0 Hz), 114.8 (d, $J$ = 21.0 Hz), 45.9, 44.6, 42.6, 35.3, 29.0. LCMS (ESI): m/z calcd for $C_{14}H_{17}BrClFN_2O_3S$; found $[M+H]^+$ 427.01.

*N-((1-((4-bromo-2,3-difluorophenyl)sulfonyl)piperidin-4-yl)methyl)-2-chloroacetamide* **(38) 1083918**

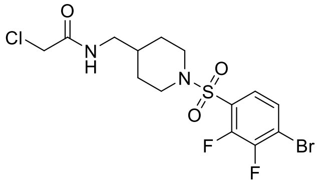

Obtained using procedure C on 0.2 mmol scale, white solid, 0.04 mmol, 18.0 mg, 20% yield. $^1$H NMR (400 MHz, $CDCl_3$) δ 7.51 – 7.43 (m, 2H), 6.69 (b, 1H), 4.03 (s, 2H), 3.87 (d, *J* = 12.3 Hz, 2H), 3.21 (t, *J* = 6.5 Hz, 2H), 2.58 (t, *J* = 12.2 Hz, 2H), 1.79 – 1.76 (m, 2H), 1.66 – 1.59 (m, 1H), 1.33 (ddd, *J* = 24.8, 12.3, 4.2 Hz, 2H). $^{13}$C NMR (100 MHz, $CDCl_3$) δ 166.1, 148.7 (dd, *J* = 253.0, 14.7 Hz), 147.6 (dd, *J* = 260, 15.2 Hz), 128.0 (d, *J* = 4.2 Hz), 127.3 (d, *J* = 12.5 Hz), 125.5 (d, *J* = 4.5 Hz), 116.1 (d, *J* = 18.1 Hz), 45.5, 44.6, 42.6, 35.3, 29.2. LCMS (ESI): m/z calcd for $C_{14}H_{16}BrClF_2N_2O_3S$; found $[M+H]^+$ 447.06.

*N-((1-((4-bromo-2,5-difluorophenyl)sulfonyl)piperidin-4-yl)methyl)-2-chloroacetamide* **(39) 1083919**

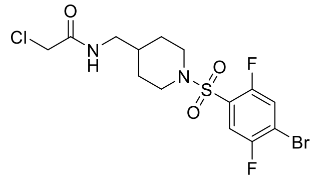

Obtained using procedure C on 0.2 mmol scale, white solid, 0.08 mmol, 35.0 mg, 40.0 % yield. $^1$H NMR (400 MHz, $CDCl_3$) δ 7.60 – 7.57 (m, 1H), 7.47 – 7.44 (m, 1H), 6.68 (b, 1H), 4.03 (s, 2H), 3.87 (d, *J* = 12.4 Hz, 2H), 3.20 (t, *J* = 6.5 Hz, 2H), 2.59 (t, *J* = 12.2 Hz, 2H), 1.79 – 1.75 (m, 2H), 1.66 – 1.59 (m, 1H), 1.31 (ddd, *J* = 24.8, 12.3, 4.2 Hz, 2H).$^{13}$C NMR (100 MHz, $CDCl_3$) δ 166.1, 155.0 (dd, *J* = 250.0, 3.2 Hz), 154.2 (dd, *J* = 260.0, 3.2 Hz), 126.5 (dd, *J* = 17.6, 5.4 Hz), 122.3 (d, *J* = 27.2 Hz), 117.9 (dd, *J* = 27.3, 1.7 Hz), 115.1 (dd, *J* = 23.7, 9.5 Hz), 45.6, 44.6, 42.6, 35.3, 29.24. LCMS (ESI): m/z calcd for $C_{14}H_{16}BrClF_2N_2O_3S$; found $[M+H]^+$ 447.01.

*N-((1-((4-bromo-3,5-difluorophenyl)sulfonyl)piperidin-4-yl)methyl)-2-chloroacetamide* **(40) 1083920**

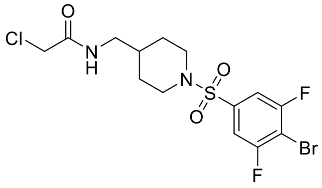

Obtained using procedure C on 0.2 mmol scale, white solid, 0.096 mmol, 43.0 mg, 48% yield. $^1$H NMR (400 MHz, $CDCl_3$) δ 7.35 – 7.31 (m, 2H), 6.67 (b, 1H), 4.03 (s, 2H), 3.81 (d, *J* = 11.8 Hz, 2H), 3.20 (t, *J* = 6.5 Hz, 2H), 2.37 (td, *J* = 11.9, 2.5 Hz, 2H), 1.78 (dd, *J* = 12.8, 1.8 Hz, 2H), 1.61 – 1.51 (m, 1H), 1.35 (ddd, *J* = 24.8, 12.1, 4.1 Hz, 2H). $^{13}$C NMR (100 MHz, $CDCl_3$) δ 166.1, 159.9 (dd, *J* = 255.0, 3.6 Hz), 138.0 (t, *J* = 7.3 Hz), 111.40 – 111.1 (m), 103.8 (t, *J* = 24.3 Hz), 45.9, 44.5, 42.6, 35.2, 29.0. LCMS (ESI): m/z calcd for $C_{14}H_{16}BrClF_2N_2O_3S$; found $[M+H]^+$ 447.01.

**Procedure D**

*Tert*-butyl (4-methylpiperidin-4-yl)carbamate **41** or *tert*-butyl ((4-methylpiperidin-4-yl)methyl)carbamate **44** (1.1 equiv, 0.5 mmol) was dissolved in 3 ml dry DCM and then DIPEA (3 equiv, 1.5 mmol) was added. The reaction mixture was cooled at 0°C and 4-bromo-3-fluorobenzenesulfonyl chloride (1.0 equiv, 0.45 mmol) was added slowly. Stirring rt overnight. The reaction mixture was quenched with sat. $NaHCO_3$ (15 ml) and extracted with DCM (3 x 20 ml). The combined organic phases were dried over $MgSO_4$, filtered and concentrated under reduced pressure. The obtained crude was purified with flash column chromatography [Biotage, hexane – EA, 0-100% EtOAc in hexane] and used directly in the next step. The intermediate was cooled at 0°C, 3ml dry DCM were added, followed by 300 µl TFA (~15 equiv). The reaction mixture was stirred at 0° for 30 min and then rt overnight. Solvents were removed under reduced pressure and the obtained TFA salt was used directly in the next step. The residue was suspended in 3 ml dry DCM and cooled at 0°C. Under stirring, DIPEA (3 equiv) was added. After 10 min, chloroacetyl chloride (1.0 equiv) was added to the reaction mixture. Stirring at 0°C for 30min and then rt overnight. The reaction mixture was quenched with sat. $NaHCO_3$ (10 ml) and extracted with DCM (3 x 10 ml). The combined organic phases were dried over $MgSO_4$, filtered and concentrated under reduced pressure. The obtained crude was purified with flash column chromatography [Biotage, hexane – EA, 0-100% EtOAc in hexane].

**Scheme 2**. Synthetic route for analog **43 (1124381)** [a]

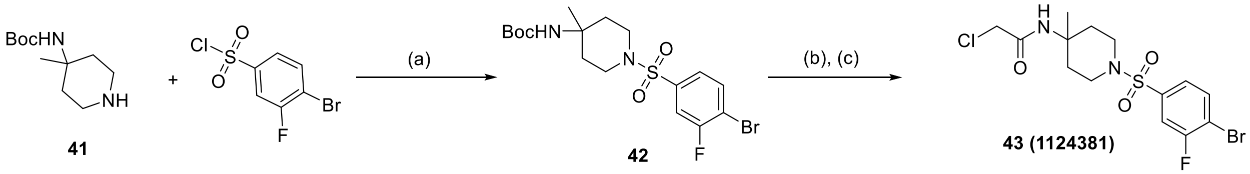

[a] Reagents and conditions: (a) DIPEA, DCM, 0°C to rt, overnight; (b) TFA, DCM, rt, overnight; (c) 4-bromo-3-fluorobenzenesulfonyl chloride, DIPEA, DCM, 0°C to rt, overnight.

*N-(1-((4-bromo-3-fluorophenyl)sulfonyl)-4-methylpiperidin-4-yl)-2-chloroacetamide* **(43) 1124381**

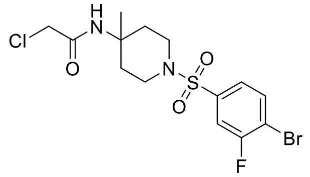

Obtained using procedure D on 0.5 mmol scale, off-white solid, 0.15 mmol, 65.5 mg, 30.0 % yield (over 3 steps). $^{1}$H NMR (400 MHz, $CDCl_3$) δ 7.75 (dd, *J* = 8.3, 6.5 Hz, 1H), 7.50 (dd, *J* = 7.7, 2.0 Hz, 1H), 7.47 – 7.36 (m, 1H), 6.08 (b, 1H), 3.89 (s, 2H), 3.37 (dt, *J* = 11.9, 4.4 Hz, 2H), 2.88 – 2.64 (m, 2H), 2.29 – 2.13 (m, 2H), 1.76 (ddd, *J* = 14.2, 10.3, 4.1 Hz, 2H), 1.37 (s, 3H). $^{13}$C NMR (100 MHz, $CDCl_3$) δ 165.3, 158.9 (d, *J* = 253.1 Hz), 137.48 (d, *J* = 5.6 Hz), 134.5, 124.1 (d, *J* = 4.1 Hz), 115.7 (d, *J* = 25.1 Hz), 115.0 (d, *J* = 21.0 Hz), 51.5, 42.9, 42.1, 35.2, 25.2. LCMS (ESI): m/z calcd for $C_{14}H_{17}BrClFN_2O_3S$; found $[M+H]^+$ 429.14.

**Scheme 3**. Synthetic route for analog **46 (1124382)** [a]

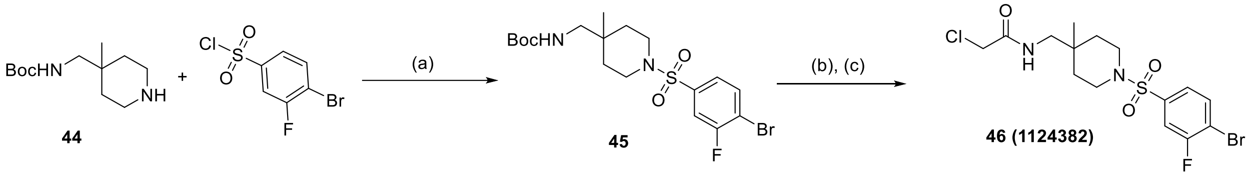

[a] Reagents and conditions: (a) DIPEA, DCM, 0°C to rt, overnight; (b) TFA, DCM, rt, overnight; (c) 4-bromo-3-fluorobenzenesulfonyl chloride, DIPEA, DCM, 0°C to rt, overnight.

*N-((1-((4-bromo-3-fluorophenyl)sulfonyl)-4-methylpiperidin-4-yl)methyl)-2-chloroacetamide* **(46) 1124382**

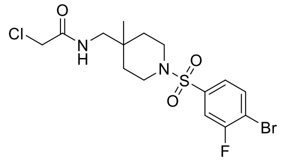

Obtained using procedure D on 0.5 mmol scale, white solid, 0.18 mmol, 84.1 mg, 36.0 % yield (over 3 steps). $^{1}$H NMR (400 MHz, $CDCl_3$) δ 7.73 (dd, *J* = 8.3, 6.5 Hz, 1H), 7.48 (dd, *J* = 7.7, 2.0 Hz, 1H), 7.40 (dd, *J* = 8.3, 1.5 Hz, 1H), 6.65 (t, *J* = 5.9 Hz, 1H), 4.03 (s, 2H), 3.34 – 3.21 (m, 2H), 3.12 (d, *J* = 6.6 Hz, 2H), 2.87 (ddd, *J* = 12.2, 9.0, 3.5 Hz, 2H), 1.54 (ddd, *J* = 13.1, 8.9, 4.0 Hz, 2H), 1.42 (ddd, *J* = 13.5, 6.0, 3.7 Hz, 2H), 0.85 (s, 3H). $^{13}$C NMR (100 MHz, $CDCl_3$) δ 166.1, 158.8 (d, *J* = 253.0 Hz), 137.7 (d, *J* = 5.6 Hz), 134.5, 124.0 (d, *J* = 4.1 Hz), 115.6 (d, *J* = 25.0 Hz), 114.8 (d, *J* = 21.0 Hz), 48.1, 42.6, 41.9, 33.7, 32.7, 22.1. LCMS (ESI): m/z calcd for $C_{15}H_{19}BrClFN_2O_3S$; found $[M+H]^+$ 443.14.

**Scheme 4**. Synthetic route for analog **49 (1084346)** [a]

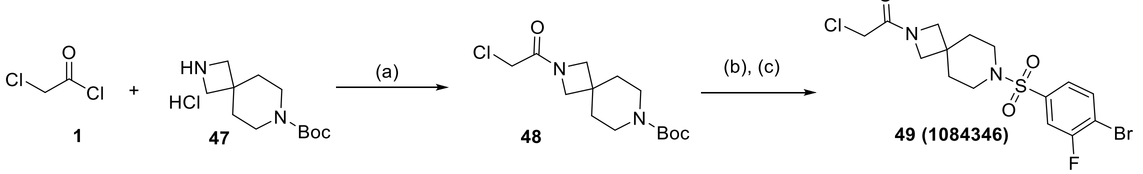

[a] Reagents and conditions: (a) DIPEA, DCM, 0°C to rt, 4h; (b) TFA, DCM, rt, overnight; (c) 4-bromo-3-fluorobenzenesulfonyl chloride, DIPEA, DCM, 0°C to rt, overnight.

**Procedure E:** *Tert*-butyl 2,7-diazaspiro[3.5]nonane-7-carboxylate hydrochloride **47** (1.0 equiv, 1 mmol, 262 mg) was dissolved in 5 ml dry DCM. DIPEA (4 equiv, 4 mmol, 710 µl) was added. The reaction mixture was cooled at 0°C and chloroacetyl chloride **1** (1.2 equiv, 1.2 mmol, 95 µl) was added dropwise. Stirring rt for 4h. The reaction mixture was quenched with sat. $NaHCO_3$ (15 ml) and extracted with DCM (3 x 20 ml). The combined organic phases were dried over $MgSO_4$, filtered and concentrated under reduced pressure. The obtained oil was purified with flash column chromatography [Biotage, hexane – EA, 0-100% EtOAc in hexane].

**Procedure F:** The residue was cooled at 0°C, 3ml dry DCM were added, followed by 300 µl TFA (15 equiv). The reaction mixture was stirred at 0° for 30 min and then rt overnight. Solvents were removed under reduced pressure and the obtained TFA salt was used directly in the next step.

**Procedure G:** The obtained TFA salt (1.1 equiv, 0.583 mmol, 118.0 mg) was suspended in 5 ml dry DCM and cooled at 0°C. Under stirring, DIPEA (3 equiv, 1.59 mmol, 282.5 µl) was added. After 5 min, the 4-bromo-3-fluorobenzenesulfonyl chloride (1.0 equiv, 0.53 mmol, 145 mg) was added to the reaction mixture. Stirring at 0 °C for 30min and then rt overnight. The reaction mixture was quenched with sat. $NaHCO_3$ (10 ml) and extracted with DCM (3 x 10ml). The combined organic phases were dried over $MgSO_4$, filtered and concentrated under reduced pressure. The obtained crude was purified with flash column chromatography [Biotage, hexane – EA, 0-100% EtOAc in hexane].

*1-(7-((4-bromo-3-fluorophenyl)sulfonyl)-2,7-diazaspiro[3.5]nonan-2-yl)-2-chloroethan-1-one* **(49) 1084346**

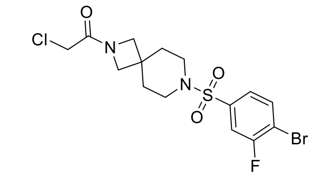

Obtained using procedure G on 0.53 mmol scale, white solid, 0.15 mmol, 67.6 mg, 30.0 % yield.[1]H NMR (400 MHz, $CDCl_3$) δ 7.74 (dd, *J* = 8.3, 6.5 Hz, 1H), 7.47 (dd, *J* = 7.6, 2.0 Hz, 1H), 7.41 – 7.38 (m, 1H), 3.89 (s, 2H), 3.83 (s, 2H), 3.66 (s, 2H), 3.05 – 2.95 (m, 4H), 1.87 (t, *J* = 5.6 Hz, 4H).[13]C NMR (100 MHz, $CDCl_3$) δ 166.2, 158.9 (d, *J* = 253.3 Hz), 137.3 (d, *J* = 5.6 Hz), 134.6, 124.0 (d, *J* = 4.1 Hz), 115.6 (d, *J* = 25.0 Hz), 115.1 (d, *J* = 21.0 Hz), 60.1, 57.7, 43.0, 39.4, 34.4, 33.3. LCMS (ESI): m/z calcd for $C_{15}H_{17}BrClFN_2O_3S$; found $[M+H]^+$ 441.04.

**Procedure H**

Tert-butyl 5,5-difluoro-2,7-diazaspiro[3.5]nonane-2-carboxylate **50** (1.1 equiv, 0.5 mmol) was dissolved in 3 ml dry DCM and then DIPEA (3 equiv, 1.5 mmol) was added. The reaction mixture was cooled at 0°C and 4-bromo-3-fluorobenzenesulfonyl chloride (1.0 equiv, 0.45 mmol) was added slowly. Stirring rt overnight. The reaction mixture was quenched with sat. $NaHCO_3$ (15 ml) and extracted with DCM (3 x 20 ml). The combined organic phases were dried over $MgSO_4$, filtered and concentrated under reduced pressure. The obtained crude was purified with flash column chromatography [Biotage, hexane – EA, 0-100% EtOAc in hexane] and used directly in the next step. The intermediate was cooled at 0°C, 3ml dry DCM were added, followed by 300 µl TFA (~15 equiv). The reaction mixture was stirred at 0° for 30 min and then rt overnight. Solvents were removed under reduced pressure and the obtained TFA salt was used directly in the next step. The residue was suspended in 3 ml dry DCM and cooled at 0°C. Under stirring, DIPEA (3 equiv) was added. After 10 min, chloroacetyl chloride (1.0 equiv) was added to the reaction mixture. Stirring at 0°C for 30min and then rt overnight. The reaction mixture was quenched with sat. $NaHCO_3$ (10 ml) and extracted with DCM (3 x 10 ml). The combined organic phases were dried over $MgSO_4$, filtered and concentrated under reduced pressure. The obtained crude was purified with flash column chromatography [Biotage, hexane – EA, 0-100% EtOAc in hexane].

### **Scheme 5.** Synthetic route for analog **52 (1084353)** [a]

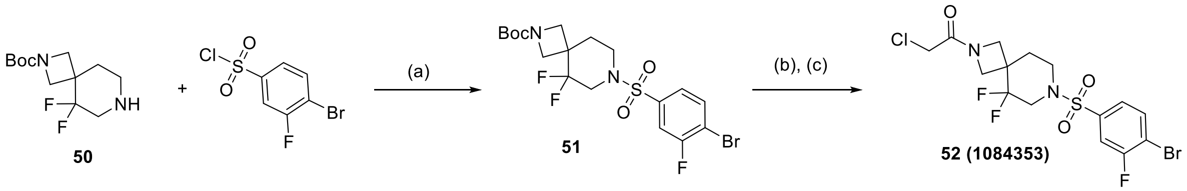

[a] Reagents and conditions: (a) DIPEA, DCM, 0°C to rt, overnight; (b) TFA, DCM, rt, overnight; (c) 4-bromo-3-fluorobenzenesulfonyl chloride, DIPEA, DCM, 0°C to rt, overnight.

*1-(7-((4-bromo-3-fluorophenyl)sulfonyl)-5,5-difluoro-2,7-diazaspiro[3.5]nonan-2-yl)-2-chloroethan-1-one* **(52) 1084353**

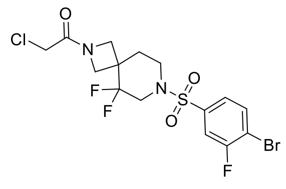

Obtained using procedure H on 0.45 mmol scale, white solid, 0.13 mmol, 57.6 mg, 28% yield (over 3 steps). $^1$H NMR (400 MHz, $CDCl_3$) δ 7.76 (dd, *J* = 8.3, 6.5 Hz, 1H), 7.51 (dd, *J* = 7.6, 2.0 Hz, 1H), 7.45 – 7.43 (m, 1H), 4.38 (d, *J* = 9.1 Hz, 1H), 4.08 (d, *J* = 10.2 Hz, 1H), 3.92 (d, *J* = 9.0 Hz, 1H), 3.87 (d, *J* = 1.9 Hz, 2H), 3.67 (d, *J* = 10.2 Hz, 1H), 3.53 – 3.45 (m, 1H), 3.33 – 3.17 (m, 2H), 3.07 – 3.01 (m, 1H), 2.07 (t, *J* = 5.5 Hz, 2H). $^{13}$C NMR (100 MHz, $CDCl_3$) δ 166.0, 159.0 (d, *J* = 253.7 Hz), 137.9 (d, *J* = 5.7 Hz), 134.9, 124.0 (d, *J* = 4.1 Hz), 120.6, 118.2, 115.8 – 115.4 (m), 54.8 (t, *J* = 4.6 Hz), 52.9 (t, *J* = 5.0 Hz), 47.6 (t, *J* = 32.6 Hz), 41.8, 39.4 – 37.6 (m), 32.0 (t, *J* = 2.7 Hz). LCMS (ESI): m/z calcd for $C_{15}H_{15}BrClF_3N_2O_3S$; found $[M+H]^+$ 447.06.

**Scheme 6**. Synthetic route for analog **55 (1084347)** [a]

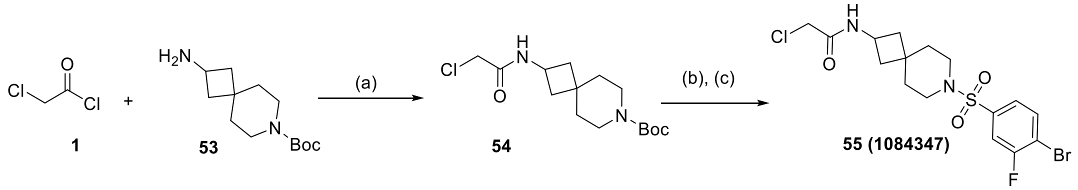

[a] Reagents and conditions: (a) DIPEA, DCM, 0°C to rt, 4h; (b) 4N HCl/dioxane, rt, 3h; (c) 4-bromo-3-fluorobenzenesulfonyl chloride, DIPEA, DCM, 0°C to rt, overnight.

**Procedure I:** *Tert*-Butyl 2-amino-7-azaspiro[3.5]nonane-7-carboxylate **53** (1.0 equiv, 1 mmol, 240 mg) was dissolved in 5 ml dry DCM. DIPEA (4 equiv, 4 mmol, 710 µl) was added. The reaction mixture was cooled at 0°C and chloroacetyl chloride **1** (1.1 equiv, 1.1 mmol, 86 µl) was added dropwise. Stirring rt for 4h. The reaction mixture was quenched with sat. $NaHCO_3$ (15 ml) and extracted with DCM (3 x 20 ml). The combined organic phases were dried over $MgSO_4$, filtered and concentrated under reduced pressure. The obtained oil was purified with flash column chromatography [Biotage, hexane – EA, 0-100% EtOAc in hexane].

**Procedure J:** The residue was suspended in 3 ml HCl/dioxane (4N) for Boc-deprotection. Stirring rt for 3h. The solvent was removed under reduced pressure and the white solid oil was used directly in the next step.

**Procedure K:** The obtained HCl salt (1.1 equiv, 0.65 mmol, 140.6 mg) was suspended in 5 ml dry DCM and cooled at 0°C. Under stirring, DIPEA (3 equiv, 1.75 mmol, 312 µl) was added. After 5 min, 4-bromo-3-fluorobenzenesulfonyl chloride (1.0 equiv, 0.585 mmol, 159.7 mg) was added to the reaction mixture. Stirring at 0 °C for 30min and then rt overnight. The reaction mixture was quenched with sat. $NaHCO_3$ (10 ml) and extracted with DCM (3 x 10ml). The combined organic phases were dried over $MgSO_4$, filtered and concentrated under reduced pressure. The obtained crude was purified with flash column chromatography [Biotage, hexane – EA, 0-100% EtOAc in hexane].

*N-(7-((4-bromo-3-fluorophenyl)sulfonyl)-7-azaspiro[3.5]nonan-2-yl)-2-chloroacetamide* **(55) 1084347**

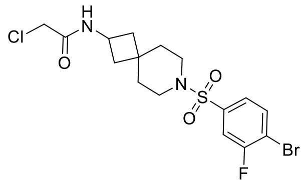

Obtained using procedure K on 0.585 mmol scale, white solid, 0.12 mmol, 53.0 mg, 20.0% yield. $^1$H NMR (400 MHz, $CDCl_3$) δ 7.73 (dd, *J* = 8.3, 6.5 Hz, 1H), 7.48 (dd, *J* = 7.7, 2.0 Hz, 1H), 7.40 (dd, *J* = 8.3, 1.5 Hz, 1H), 6.61 (b, 1H), 4.36 – 4.26 (m, 1H), 3.98 (s, 2H), 3.02 – 2.99 (m, 2H), 2.95 – 2.92 (m, 2H), 2.26 – 2.21 (m, 2H), 1.75 – 1.70 (m, 2H), 1.69 – 1.64 (m, 4H). $^{13}$C NMR (100 MHz, $CDCl_3$) δ 165.1, 158.9 (d, *J* = 253.0 Hz), 137.7 (d, *J* = 5.6 Hz), 134.4, 124.1 (d, *J* = 4.0 Hz), 115.7 (d, *J* = 25.0 Hz), 114.8 (d, *J* = 21.0 Hz), 43.2, 43.0, 42.4, 40.1, 39.6, 38.5, 34.9, 31.7. LCMS (ESI): m/z calcd for $C_{16}H_{19}BrClFN_2O_3S$; found $[M+H]^+$ 455.01.

**Procedure L**

The appropriate boc-protected diamine **56, 59, 62, 65, 68** or **71** (1.1 equiv, 0.5 mmol) was dissolved in 3 ml dry DCM and then DIPEA (3 equiv, 1.5 mmol) was added. The reaction mixture was cooled at 0°C and 4-bromo-3-fluorobenzenesulfonyl chloride (1.0 equiv, 0.45 mmol) was added slowly. Stirring rt overnight. The reaction mixture was quenched with sat. $NaHCO_3$ (15 ml) and extracted with DCM (3 x 20 ml). The combined organic phases were dried over $MgSO_4$, filtered and concentrated under reduced pressure. The obtained crude was purified with flash column chromatography [Biotage, hexane – EA, 0-100% EtOAc in hexane] and used directly in the next step. The intermediate was cooled at 0°C, 3ml dry DCM were added, followed by 300 µl TFA (~15 equiv). The reaction mixture was stirred at 0° for 30 min and then rt overnight. Solvents were removed under reduced pressure and the obtained TFA salt was used directly in the next step. The residue was suspended in 3 ml dry DCM and cooled at 0°C. Under stirring, DIPEA (3 equiv) was added. After 10 min, chloroacetyl chloride (1.0 equiv) was added to the reaction mixture. Stirring at 0°C for 30min and then rt overnight. The reaction mixture was quenched with sat. $NaHCO_3$ (10 ml) and extracted with DCM (3 x 10 ml). The combined organic phases were dried over $MgSO_4$, filtered and concentrated under reduced pressure. The obtained crude was purified with flash column chromatography [Biotage, hexane – EA, 0-100% EtOAc in hexane].

**Scheme 7**. Synthetic route for analog **58 (1084348)** [a]

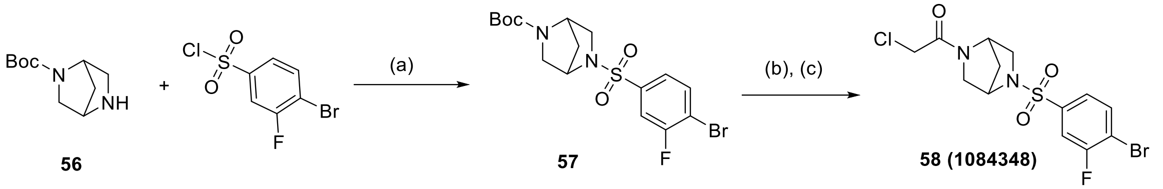

[a] Reagents and conditions: (a) DIPEA, DCM, 0°C to rt, overnight; (b) TFA, DCM, rt, overnight; (c) 4-bromo-3-fluorobenzenesulfonyl chloride, DIPEA, DCM, 0°C to rt, overnight.

*1-(5-((4-bromo-3-fluorophenyl)sulfonyl)-2,5-diazabicyclo[2.2.1]heptan-2-yl)-2-chloroethan-1-one* **(58) 1084348**

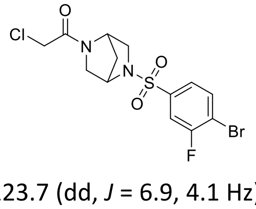

Obtained using procedure L on 0.45 mmol scale, off-white semi-solid, 0.09 mmol, 37.2 mg, 20% yield (over 3 steps). $^1$H NMR (400 MHz, $CDCl_3$) δ 7.75 (dd, *J* = 8.2, 6.6 Hz, 1H), 7.58 (dd, *J* = 7.6, 1.9 Hz, 1H), 7.50 (d, *J* = 8.3 Hz, 1H), 4.88 (s, 0.6 H), 4.58 – 4.55 (m, 1.4 H), 4.00 (q, *J* = 12.2 Hz, 0.6 H), 3.87 (q, *J* = 12.2 Hz, 1.3 H), 3.67 – 3.65 (m, 0.6 H), 3.58 – 3.37 (m, 3H), 3.23 (dd, *J* = 9.4, 2.0 Hz, 0.6 H), 1.85 – 1.76 (m, 1H), 1.56 – 1.48 (m, 1H). $^{13}$C NMR (100 MHz, $CDCl_3$) δ 164.8, 164.1, 159.0 (dd, *J* = 253.5, 3.9 Hz), 139.6 (dd, *J* = 13.9, 5.6 Hz), 134.8, 123.7 (dd, *J* = 6.9, 4.1 Hz), 115.34 (dt, *J* = 21.0, 6.0 Hz), 60.3, 59.1, 58.7, 56.5, 55.2, 54.6, 54.3, 54.0, 41.3, 41.1, 37.4, 35.8. LCMS (ESI): m/z calcd for $C_{13}H_{13}BrClFN_2O_3S$; found $[M+H]^+$ 413.08.

**Scheme 8**. Synthetic route for analog **61 (1084349)** [a]

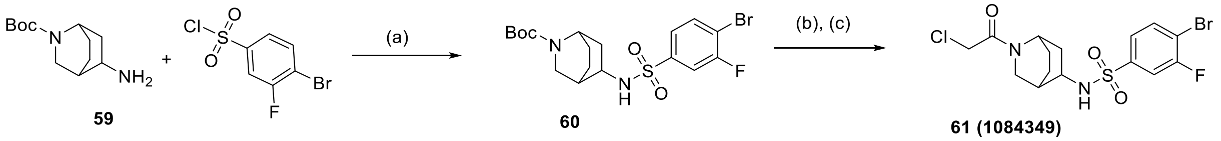

[a] Reagents and conditions: (a) DIPEA, DCM, 0°C to rt, overnight; (b) TFA, DCM, rt, overnight; (c) 4-bromo-3-fluorobenzenesulfonyl chloride, DIPEA, DCM, 0°C to rt, overnight.

*4-bromo-N-(2-(2-chloroacetyl)-2-azabicyclo[2.2.2]octan-5-yl)-3-fluorobenzenesulfonamide* **(61) 1084349**

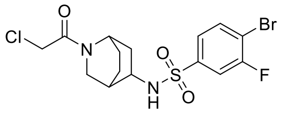

Obtained using procedure L on 0.45 mmol scale, white solid, 0.07 mmol, 32.7 mg, 16% yield (over 3 steps). Racemic mixture, conformers are observed. $^1$H NMR (400 MHz, $CDCl_3$) δ 7.74 – 7.68 (m, 2H), 7.65 – 7.61 (m, 2H), 7.57 – 7.53 (m, 2H), 6.79 (d, *J* = 8.0 Hz, 0.5H), 5.96 – 5.93 (m, 1H), 4.46 – 4.44 (m, 1H), 4.07 (d, *J* = 1.8 Hz, 0.5H), 4.02 (d, *J* = 2.5 Hz, 1H), 3.99 (s, 1H), 3.96 (s, 1.5H), 3.77 – 3.70 (m, 2H), 3.58 – 3.48 (m, 3H), 3.40 – 3.38 (m, 2H), 3.29 – 3.26 (m, 0.5H), 2.33 – 2.26 (m, 1H), 2.23 – 2.14 (m, 1H), 2.10 – 2.02 (m, 2H), 2.01 – 1.47 (m, 12H), 1.40 – 1.35 (m, 0.5H). $^{13}$C NMR (100 MHz, $CDCl_3$) δ 165.5 – 165.2 (m), 158.9 (dd, *J* = 252.7, 12.4 Hz), 142.6 (d, *J* = 5.8 Hz), 141.6 (dd, *J* = 19.6, 5.8 Hz), 134.64 – 134.4 (m), 123.7 – 123.5 (m), 115.3 – 114.8 (m), 50.2, 50.0, 49.3, 48.3, 47.5, 47.1, 46.9, 43.8, 43.2, 42.2, 41.5, 41.1, 40.8, 36.0, 35.7, 34.7, 31.7, 31.3, 30.9, 26.4, 25.3, 24.5, 22.5, 22.1, 17.3. LCMS (ESI): m/z calcd for $C_{15}H_{17}BrClFN_2O_3S$; found $[M+H]^+$ 411.09.

**Scheme 9**. Synthetic route for analog **64 (1084350)** [a]

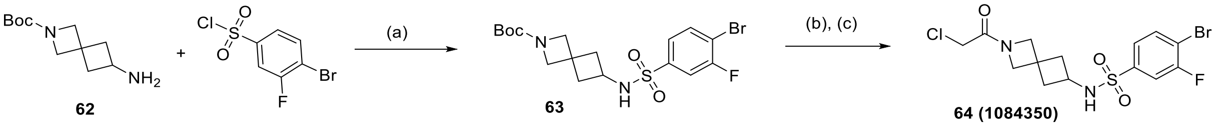

[a] Reagents and conditions: (a) DIPEA, DCM, 0°C to rt, overnight; (b) TFA, DCM, rt, overnight; (c) 4-bromo-3-fluorobenzenesulfonyl chloride, DIPEA, DCM, 0°C to rt, overnight.

*4-bromo-N-(2-(2-chloroacetyl)-2-azaspiro[3.3]heptan-6-yl)-3-fluorobenzenesulfonamide* **(64) 1084350**

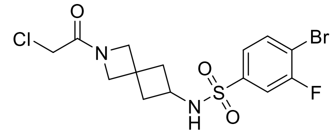

Obtained using procedure L on 0.45 mmol scale, white solid, 0.07 mmol, 31.4 mg, 17% yield (over 3 steps). $^1$H NMR (400 MHz, $CDCl_3$) δ 7.74 – 7.69 (m, 1H), 7.60 (dd, *J* = 7.7, 1.7 Hz, 1H), 7.52 (dd, *J* = 8.3, 0.7 Hz, 1H), 5.99 – 5.80 (m, 1H), 4.23 (s, 1H), 4.14 (s, 1H), 4.02 (s, 1H), 3.94 (s, 1H), 3.84 – 3.83 (m, 2H), 3.74 – 3.66 (m, 1H), 2.49 – 2.42 (m, 2H), 2.15 – 2.08 (m, 2H). $^{13}$C NMR (100 MHz, $CDCl_3$) δ 165.9, 158.90 (d, *J* = 252.9 Hz), 141.7 (dd, *J* = 23.2, 5.7 Hz), 134.6 (d, *J* = 3.0 Hz), 123.6 (t, *J* = 4.6 Hz), 115.2 (d, *J* = 25.2 Hz), 114.9 – 114.6 (m), 62.9, 61.5, 60.6, 59.3, 43.1, 41.3, 39.4, 32.3. LCMS (ESI): m/z calcd for $C_{14}H_{15}BrClFN_2O_3S$; found $[M+H]^+$ 427.09.

**Scheme 10**. Synthetic route for analog **67 (1084351)** [a]

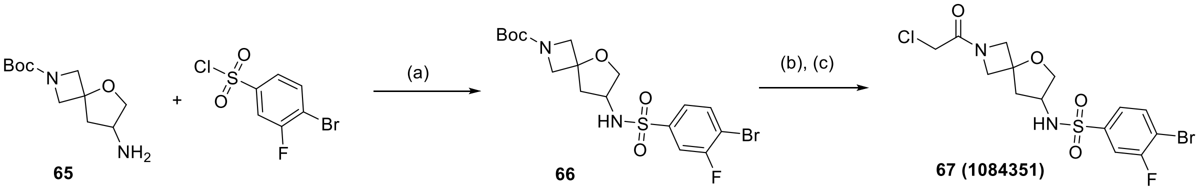

[a] Reagents and conditions: (a) DIPEA, DCM, 0°C to rt, overnight; (b) TFA, DCM, rt, overnight; (c) 4-bromo-3-fluorobenzenesulfonyl chloride, DIPEA, DCM, 0°C to rt, overnight.

*4-bromo-N-(2-(2-chloroacetyl)-5-oxa-2-azaspiro[3.4]octan-7-yl)-3-fluorobenzenesulfonamide* **(67) 1084351**

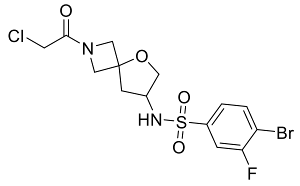

Obtained using procedure L on 0.45 mmol scale, white solid, 0.12 mmol, 54.5 mg, 27.4% yield (over 3 steps). $^1$H NMR (400 MHz, , DMSO-$d_6$) δ 8.20 (b, 1H), 8.01 (dd, *J* = 8.2, 6.9 Hz, 1H), 7.78 – 7.75 (m, 1H), 7.61 – 7.59 (m, 1H), 4.28 – 4.10 (m, 4H), 4.02 – 3.79 (m, 4H), 3.54 (d, *J* = 4.7 Hz, 1H), 2.25 (dd, *J* = 12.6, 6.3 Hz, 1H), 2.03 – 1.96 (m, 1H).$^{13}$C NMR (100 MHz, , DMSO-$d_6$) δ 165.6, 158.1 (d, *J* = 249.5 Hz), 142.0, 134.9, 124.1, 114.9 (d, *J* = 25.1 Hz), 113.4 (d, *J* = 21.0 Hz), 77.8, 71.9, 62.6, 60.8, 52.8, 40.8, 40.3. LCMS (ESI): m/z calcd for $C_{14}H_{15}BrClFN_2O_4S$; found $[M+H]^+$ 443.09.

**Scheme 11**. Synthetic route for analog **70 (1084352)** [a]

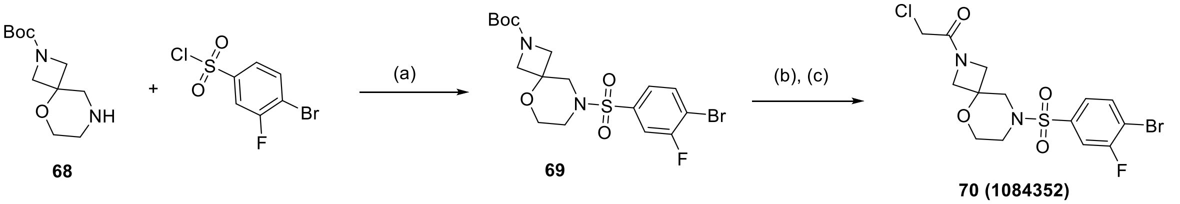

[a] Reagents and conditions: (a) DIPEA, DCM, 0°C to rt, overnight; (b) TFA, DCM, rt, overnight; (c) 4-bromo-3-fluorobenzenesulfonyl chloride, DIPEA, DCM, 0°C to rt, overnight.

*1-(8-((4-bromo-3-fluorophenyl)sulfonyl)-5-oxa-2,8-diazaspiro[3.5]nonan-2-yl)-2-chloroethan-1-one* **(70) 1084352**

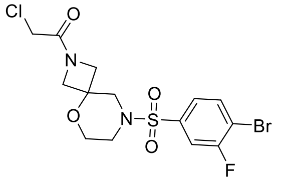

Obtained using procedure L on 0.45 mmol scale, white solid, 0.24 mmol, 107.0 mg, 53.3% yield (over 3 steps). $^{1}$H NMR (400 MHz, DMSO-$d_6$) δ 8.05 (dd, *J* = 8.3, 6.8 Hz, 1H), 7.75 (dd, *J* = 8.1, 2.0 Hz, 1H), 7.53 (dd, *J* = 8.3, 1.9 Hz, 1H), 4.17 (d, *J* = 1.5 Hz, 2H), 4.09 (q, *J* = 9.5 Hz, 2H), 3.83 – 3.76 (m, 1H), 3.75 – 3.65 (m, 3H), 3.12 (q, *J* = 11.9 Hz, 2H), 3.00 – 2.85 (m, 2H). $^{13}$C NMR (100 MHz, DMSO-$d_6$) δ 166.1, 158.3 (d, *J* = 250.0 Hz), 136.0 (d, *J* = 6.1 Hz), 135.0, 125.0 (d, *J* = 3.9 Hz), 115.9 (d, *J* = 25.3 Hz), 114.6 (d, *J* = 20.9 Hz), 70.6, 61.5, 58.5, 56.7, 50.7, 44.7, 40.2. LCMS (ESI): m/z calcd for $C_{14}H_{15}BrClFN_2O_4S$; found $[M+H]^+$ 443.04.

**Scheme 12**. Synthetic route for analog **73 (1084354)** [a]

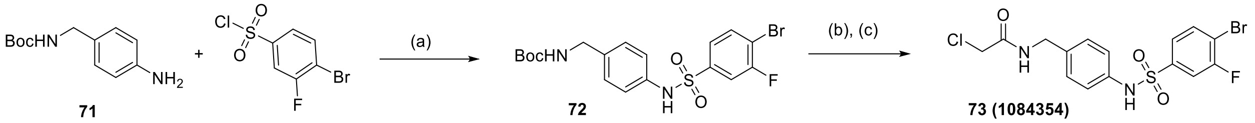

[a] Reagents and conditions: (a) DIPEA, DCM, 0°C to rt, overnight; (b) TFA, DCM, rt, overnight; (c) 4-bromo-3-fluorobenzenesulfonyl chloride, DIPEA, DCM, 0°C to rt, overnight.

*N-(4-((4-bromo-3-fluorophenyl)sulfonamido)benzyl)-2-chloroacetamide* **(73) 1084354**

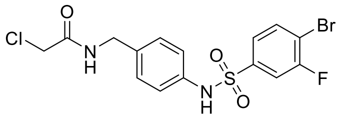

Obtained using procedure L on 0.45 mmol scale, white solid, 0.05 mmol, 25.0 mg, 12% yield (over 3 steps). $^{1}$H NMR (400 MHz, Acetone-$d_6$) δ 9.21 (b, 1H), 7.91 (b, 1H), 7.86 (dd, *J* = 8.4, 6.7 Hz, 1H), 7.64 (dd, *J* = 8.2, 2.0 Hz, 1H), 7.55 (dd, *J* = 8.4, 1.5 Hz, 1H), 7.23 (d, *J* = 8.6 Hz, 2H), 7.18 (d, *J* = 8.6 Hz, 2H), 4.37 (d, *J* = 6.0 Hz, 2H), 4.11 (s, 2H). $^{13}$C NMR (100 MHz, Acetone-$d_6$) δ 166.8, 159.5 (d, *J* = 250.2 Hz), 142.3 (d, *J* = 5.9 Hz), 137.0 (d, *J* = 5.2 Hz), 135.6, 129.4, 125.3 (d, *J* = 4.0 Hz), 122.3, 116.1 (d, *J* = 25.4 Hz), 114.7 (d, *J* = 21.0 Hz), 43.3, 43.2. LCMS (ESI): m/z calcd for $C_{15}H_{13}BrClFN_2O_3S$; found $[M+H]^+$ 435.04.

**Procedure M**

*Tert*-butyl 5,5-difluoro-2,7-diazaspiro[3.5]nonane-2-carboxylate **50** (1.1 equiv, 0.5 mmol) was dissolved in 3 ml dry DCM and then DIPEA (3 equiv, 1.5 mmol) was added. The reaction mixture was cooled at 0°C and 4-bromo-3-fluorobenzenesulfonyl chloride (1.0 equiv, 0.45 mmol) was added slowly. Stirring rt overnight. The reaction mixture was quenched with sat. $NaHCO_3$ (15 ml) and extracted with DCM (3 x 20 ml). The combined organic phases were dried over $MgSO_4$, filtered and concentrated under reduced pressure. The obtained crude was purified with flash column chromatography [Biotage, hexane – EA, 0-100% EtOAc in hexane] and used directly in the next step. The intermediate was deprotected with 5 ml HCl/dioxane (4N). Stirring rt for 3h. Solvents were removed under reduced pressure and the obtained white solid (HCl salt) was used directly in the next step. The appropriate carboxylic acid (1 equiv, 0.24 mmol) and HATU (1.2 equiv, 0.28 mmol) were dissolved in 1.5 ml dry DMF at 0°C. The deprotected amine (1.1 equiv, 0.26 mmol) and DIPEA (3 equiv, 0.72 mmol) were dissolved in 1.5 ml DMF and added to the reaction mixture under stirring. Stirring rt for 4h. The reaction mixture was diluted with sat. $NaHCO_3$ (20 ml) and extracted with ethyl acetate (3x20ml). The combined organic phases were washed with Brine, dried over $MgSO_4$, filtered and concentrated under reduced pressure. The obtained crude was purified by flash column chromatography [Biotage, hexane – EA, 0-100% EtOAc in hexane].

**Scheme 13**. Synthetic route for analogs with varying warheads [a]

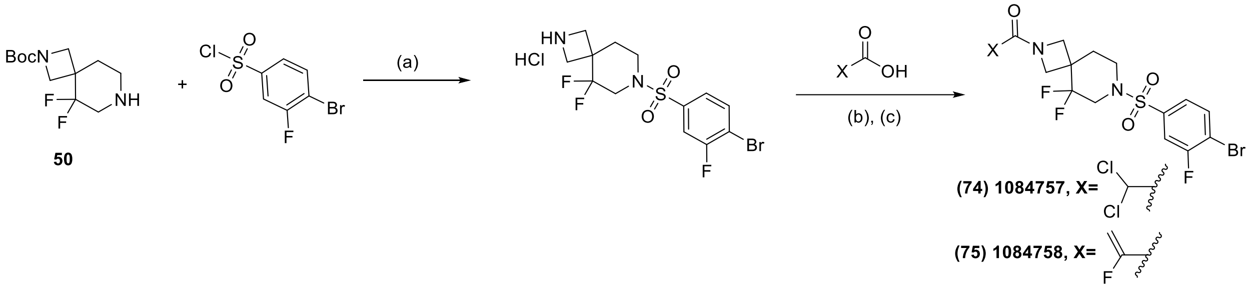

[a] Reagents and conditions: (a) DIPEA, DCM, 0°C to rt, overnight; (b) 4N HCl/dioxane, rt, 3h; (c) appropriate carboxylic acid, HATU, DIPEA, DMF, 0°C to rt, 4h.

*1-(7-((4-bromo-3-fluorophenyl)sulfonyl)-5,5-difluoro-2,7-diazaspiro[3.5]nonan-2-yl)-2,2-dichloroethan-1-one* **(74) 1084757**

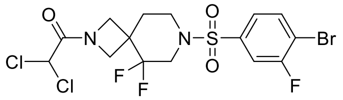

Obtained using procedure M on 0.24 mmol scale, white solid, 0.08 mmol, 38.0 mg, 32% yield. $^{1}$H NMR (400 MHz, $CDCl_3$) δ 7.77 (dd, *J* = 8.3, 6.5 Hz, 1H), 7.52 (dd, *J* = 7.6, 2.0 Hz, 1H), 7.46 – 7.43 (m, 1H), 5.87 (s, 1H), 4.58 (d, *J* = 9.5 Hz, 1H), 4.16 – 4.09 (m, 2H), 3.73 (d, *J* = 10.4 Hz, 1H), 3.50 – 3.42 (m, 1H), 3.33– 3.24 (m, 2H), 3.13 – 3.07 (m, 1H), 2.10 (t, *J* = 5.5 Hz, 2H).$^{13}$C NMR (100 MHz, $CDCl_3$) δ 162.7, 159.1 (d, *J* = 253.8 Hz), 137.9 (d, *J* = 5.7 Hz), 134.9, 124.0 (d, *J* = 4.1 Hz), 120.5, 118.0, 115.8 – 115.4 (m), 64.6, 55.9 (t, *J* = 4.7 Hz), 53.5 (t, *J* = 4.9 Hz), 47.7 (t, *J* = 32.5 Hz), 41.8, 39.8 (t, *J* = 23.6 Hz), 32.1. LCMS (ESI): m/z calcd for $C_{15}H_{14}BrCl_2F_3N_2O_3S$ found $[M+H]^+$ 511.03.

*1-(7-((4-bromo-3-fluorophenyl)sulfonyl)-5,5-difluoro-2,7-diazaspiro[3.5]nonan-2-yl)-2-fluoroprop-2-en-1-one* **(75) 1084758**

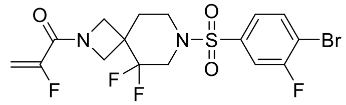

Obtained using procedure M on 0.24 mmol scale, white solid, 0.05 mmol, 22.0 mg, 21% yield. $^{1}$H NMR (400 MHz, $CDCl_3$) δ 7.77 (dd, *J* = 8.3, 6.5 Hz, 1H), 7.52 (dd, *J* = 7.6, 2.0 Hz, 1H), 7.46 – 7.43 (m, 1H), 5.68 – 5.56 (m, 1H), 5.10 (dd, *J* = 15.7, 3.3 Hz, 1H), 4.48 (dd, *J* = 10.2, 3.5 Hz, 1H), 4.13 (d, *J* = 10.7 Hz, 1H), 4.00 (dd, *J* = 10.1, 3.9 Hz, 1H), 3.70 (d, *J* = 10.6 Hz, 1H), 3.45 – 3.40 (m, 1H), 3.31 – 3.23 (m, 2H), 3.10 (dd, *J* = 12.4, 6.2 Hz, 1H), 2.07 (t, *J* = 5.5 Hz, 2H).$^{13}$C NMR (100 MHz, $CDCl_3$) δ 159.9 (t, *J* = 35.9 Hz), 156.9 (d, *J* = 269.0 Hz), 138.01 (d, *J* = 5.7 Hz), 134.9, 124.0 (d, *J* = 4.2 Hz), 120.7, 118.2, 115.8 – 115.5 (m), 100.6 (d, *J* = 14.1 Hz), 56.2 – 56.0 (m), 52.60 (t, *J* = 4.6 Hz), 47.7 (t, *J* = 32.6 Hz), 41.81, 39.60 (td, *J* = 23.6, 4.2 Hz), 32.10 (t, *J* = 2.7 Hz). LCMS (ESI): m/z calcd for $C_{16}H_{15}BrF_4N_2O_3S$ found $[M+H]^+$ 473.06

**Scheme 14**. Synthetic route for analogs **78 (1124378)**, **79 (1124379)** and **80 (1124898)** [a]

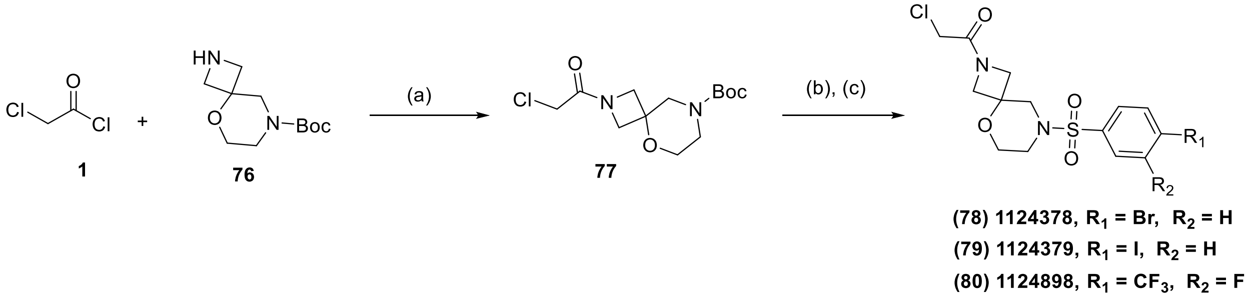

[a] Reagents and conditions: (a) DIPEA, DCM, 0°C to rt, 4h; (b) 4N HCl/dioxane, rt, 3h; (c) appropriate sulfonyl chloride, DIPEA, DCM, 0°C to rt, overnight.

**Procedure N:** *tert*-butyl 5-oxa-2,8-diazaspiro[3.5]nonane-8-carboxylate **76** (1.0 equiv, 1.2 mmol) was dissolved in 10 ml dry DCM. DIPEA (3 equiv, 3.6 mmol) was added. The reaction mixture was cooled at 0°C and chloroacetyl chloride **1** (1.2 equiv, 1.44 mmol) was added dropwise. Stirring rt for 4h. The reaction mixture was quenched with sat. $NaHCO_3$ (15 ml) and extracted with DCM (3 x 15 ml). The combined organic phases were dried over $MgSO_4$, filtered and concentrated under reduced pressure. The obtained crude was purified with flash column chromatography [Biotage, hexane – EA, 0-100% EtOAc in hexane]. The obtained intermediate was suspended in 3 ml HCl/dioxane (4N) for Boc-deprotection. Stirring rt for 3h. The solvent was removed under reduced pressure and the obtained salt was used directly in the next step. The obtained HCl salt (1.5 equiv, 0.3 mmol) was suspended in 2 ml dry DCM and cooled at 0 °C. Under stirring, DIPEA (3 equiv, 0.6 mmol) was added. After 10 min, the appropriate sulfonyl chloride (1.0 equiv, 0.2 mmol) was added slowly. Stirring at 0 °C for 30 min and then rt overnight. The reaction mixture was quenched with sat. $NaHCO_3$ (10 ml) and extracted with DCM (3 x 10ml). The combined organic phases were dried over $MgSO_4$, filtered and concentrated under reduced pressure. The obtained crude was purified with flash column chromatography [Biotage, hexane – EA, 0-100% EtOAc in hexane].

*1-(8-((4-bromophenyl)sulfonyl)-5-oxa-2,8-diazaspiro[3.5]nonan-2-yl)-2-chloroethan-1-one* **(78) 1124378**

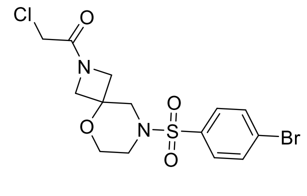

Obtained using procedure N on 0.16 mmol scale, yellow oil, 0.078 mmol, 33 mg mg, 48% yield. $^1$H NMR (400 MHz, $CDCl_3$) δ 7.73 – 7.71 (m, 2H), 7.62 – 7.60 (m, 2H), 4.20 (q, *J* = 9.4 Hz, 2H), 3.93 (b, 3H), 3.78 – 3.61 (m, 3H), 3.27 (d, *J* = 11.5 Hz, 1H), 3.15 – 3.13 (m, 1H), 2.92 (d, *J* = 11.4 Hz, 1H), 2.85 – 2.77 (m, 1H). $^{13}$C NMR (100 MHz, $CDCl_3$) δ 166.4, 134.0, 132.7, 129.2, 128.7, 70.9, 62.1, 59.4, 57.3, 51.0, 44.8, 39.7. LCMS (ESI): m/z calcd for $C_{14}H_{16}BrClN_2O_4S$; found $[M+H]^+$ 425.14.

*1-(8-((4-iodophenyl)sulfonyl)-5-oxa-2,8-diazaspiro[3.5]nonan-2-yl)-2-chloroethan-1-one* **(79) 1124379**

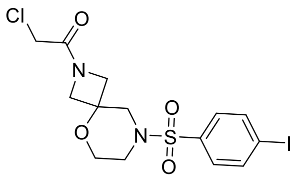

Obtained using procedure N on 0.16 mmol scale, yellow oil, 0.068 mmol, 32 mg, 43% yield. $^1$H NMR (400 MHz, $CDCl_3$) δ 7.94 – 7.91 (m, 2H), 7.46 – 7.43 (m, 2H), 4.19 (q, *J* = 9.4 Hz, 2H), 3.92 (b, 3H), 3.77 – 3.68 (m, 2H), 3.65 – 3.60 (m, 1H), 3.26 (d, *J* = 11.5 Hz, 1H), 3.13 – 3.11 (m, 1H), 2.92 (d, *J* = 11.5 Hz, 1H), 2.85 – 2.78 (m, 1H). $^{13}$C NMR (100 MHz, $CDCl_3$) δ 166.4, 138.6, 134.6, 128.9, 101.1, 70.9, 62.1, 59.4, 57.2, 51.0, 44.8, 39.6. LCMS (ESI): m/z calcd for $C_{14}H_{16}ClIN_2O_4S$; found $[M+H]^+$ 471.01.

*2-chloro-1-(8-((3-fluoro-4-(trifluoromethyl)phenyl)sulfonyl)-5-oxa-2,8-diazaspiro[3.5]nonan-2-yl)ethan-1-one* **(80) 1124898**

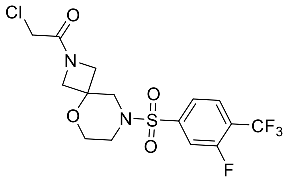

Obtained using procedure N on 0.20 mmol scale, white solid, 0.06 mmol, 26 mg, 30% yield. $^1$H NMR (400 MHz, $CDCl_3$) δ 7.87 – 7.83 (m, 1H), 7.63 (dd, *J* = 15.6, 8.8 Hz, 2H), 4.21 (q, *J* = 9.4 Hz, 2H), 3.95 (s, 2H), 3.93 (s, 2H), 3.81 – 3.63 (m, 2H), 3.32 (d, *J* = 11.5 Hz, 1H), 3.18 (d, *J* = 11.5 Hz, 1H), 3.02 (d, *J* = 11.5 Hz, 1H), 2.95 (dd, *J* = 7.5, 3.2 Hz, 1H). $^{13}$C NMR (100 MHz, $CDCl_3$) δ 166.4, 159.7 (d, *J* = 261.5 Hz), 141.2 (d, *J* = 6.6 Hz), 128.7 (d, *J* = 3.4 Hz), 123.3 (d, *J* = 4.3 Hz), 123.2 – 122.6 (m), 120.2, 116.5 (d, *J* = 23.5 Hz), 70.9, 62.1, 59.3, 57.2, 51.0, 44.8, 39.7. LCMS (ESI): m/z calcd for $C_{15}H_{15}ClF_4N_2O_4S$; found $[M+H]^+$ 431.14

**Procedure O**

The appropriate boc-protected diamine **81, 84** or **87** (1.1 equiv, 0.5 mmol) was dissolved in 3 ml dry DCM and then DIPEA (3 equiv, 1.5 mmol) was added. The reaction mixture was cooled at 0°C and 4-bromo-3-fluorobenzenesulfonyl chloride (1.0 equiv, 0.45 mmol) was added slowly. Stirring rt overnight. The reaction mixture was quenched with sat. $NaHCO_3$ (15 ml) and extracted with DCM (3 x 20 ml). The combined organic phases were dried over $MgSO_4$, filtered and concentrated under reduced pressure. The obtained crude was purified with flash column chromatography [Biotage, hexane – EA, 0-100% EtOAc in hexane] and used directly in the next step. The intermediate was cooled at 0°C, 3ml dry DCM were added, followed by 300 µl TFA (~15 equiv). The reaction mixture was stirred at 0° for 30 min and then rt overnight. Solvents were removed under reduced pressure and the obtained TFA salt was used directly in the next step. The residue was suspended in 3 ml dry DCM and cooled at 0°C. Under stirring, DIPEA (3 equiv) was added. After 10 min, chloroacetyl chloride (1.0 equiv) was added to the reaction mixture. Stirring at 0°C for 30min and then rt overnight. The reaction mixture was quenched with sat. $NaHCO_3$ (10 ml) and extracted with DCM (3 x 10 ml). The combined organic phases were dried over $MgSO_4$, filtered and concentrated under reduced pressure. The obtained crude was purified with flash column chromatography [Biotage, hexane – EA, 0-100% EtOAc in hexane].

**Scheme 15**. Synthetic route for analog **83 (1124383)**[a]

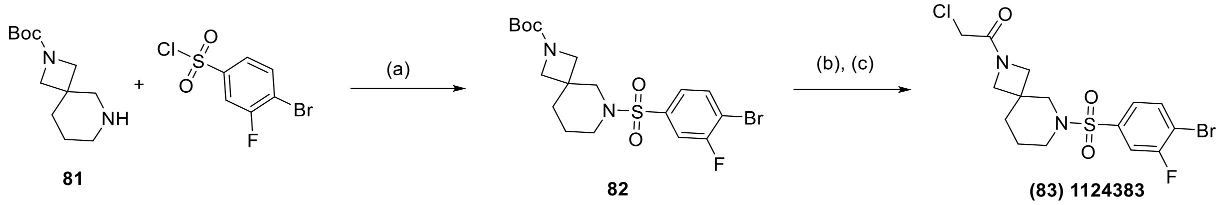

[a] Reagents and conditions: (a) DIPEA, DCM, 0ºC to rt, overnight; (b) TFA, DCM, rt, overnight; (c) 4-bromo-3-fluorobenzenesulfonyl chloride, DIPEA, DCM, 0ºC to rt, overnight.

*N-((1-((4-bromo-3-fluorophenyl)sulfonyl)-4-methylpiperidin-4-yl)methyl)-2-chloroacetamide* **(83) 1124383**

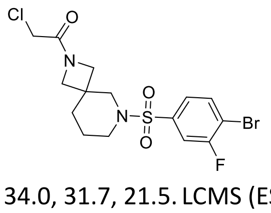

Obtained using procedure O on 0.5 mmol scale, orange solid, 0.17 mmol, 78.4 mg, 34.0 % yield (over 3 steps). $^{1}$H NMR (400 MHz, DMSO) δ 8.03 (dd, *J* = 8.3, 6.8 Hz, 1H), 7.76 (dd, *J* = 8.2, 2.0 Hz, 1H), 7.53 (dd, *J* = 8.3, 1.8 Hz, 1H), 4.14 (d, *J* = 2.3 Hz, 2H), 3.87 (d, *J* = 5.9 Hz, 2H), 3.59 (d, *J* = 5.3 Hz, 2H), 3.05 (q, *J* = 11.5 Hz, 2H), 2.98 – 2.82 (m, 2H), 1.63 – 1.53 (m, 4H). $^{13}$C NMR (100 MHz, DMSO) δ 166.0, 158.2 (d, *J* = 249.8 Hz), 136.9 (d, *J* = 5.9 Hz), 134.9, 124.8 (d, *J* = 3.8 Hz), 115.7 (d, *J* = 25.1 Hz), 114.2 (d, *J* = 20.9 Hz), 58.2, 56.3, 53.1, 45.6, 34.0, 31.7, 21.5. LCMS (ESI): m/z calcd for $C_{15}H_{17}BrClFN_2O_3S$; found $[M+H]^+$ 441.14

**Scheme 16**. Synthetic route for analogs **86 (1124384)**[a] and **89 (1124385)**[a]

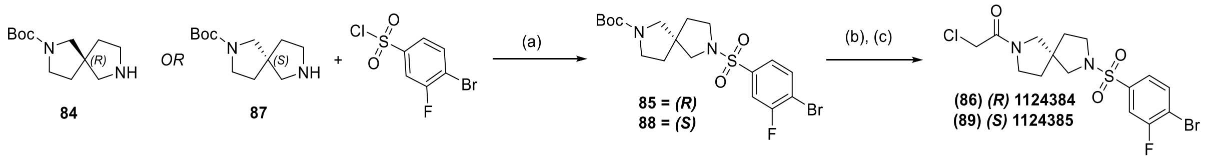

[a] Reagents and conditions: (a) DIPEA, DCM, 0ºC to rt, overnight; (b) TFA, DCM, rt, overnight; (c) 4-bromo-3-fluorobenzenesulfonyl chloride, DIPEA, DCM, 0ºC to rt, overnight. Stereochemistry for final products assigned based on crystallography.

*(R)-1-(7-((4-bromo-3-fluorophenyl)sulfonyl)-2,7-diazaspiro[4.4]nonan-2-yl)-2-chloroethan-1-one* **(86) 1124384**

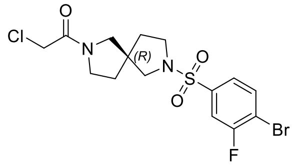

Obtained using procedure O on 0.5 mmol scale, yellow oil, 0.34 mmol, 152 mg, 68.0 % yield (over 3 steps). $^{1}$H NMR (400 MHz, $CDCl_3$) δ 7.75 (ddd, *J* = 8.3, 6.5, 3.4 Hz, 1H), 7.55 (ddd, *J* = 7.5, 5.3, 2.0 Hz, 1H), 7.47 (td, *J* = 8.0, 1.9 Hz, 1H), 3.98 (s, 1H), 3.90 (s, 1H), 3.67 – 3.47 (m, 2H), 3.43 – 3.21 (m, 5H), 3.11 (t, *J* = 10.1 Hz, 1H), 1.98 – 1.74 (m, 4H). $^{13}$C NMR (100 MHz, $CDCl_3$) δ 165, 160.2 (d, *J* = 1.5 Hz), 137.7 (dd, *J* = 8.7, 5.6 Hz), 134.6 (d, *J* = 2.0 Hz), 123.9 (dd, *J* = 11.2, 4.1 Hz), 115.6 – 115.3 (m), 56.0, 55.1, 54.5, 49.6, 47.4, 46.7, 45.3, 41.5, 35.2, 34.5, 33.0. LCMS (ESI): m/z calcd for $C_{15}H_{17}BrClFN_2O_3S$; found $[M+H]^+$ 441.09.

*(S)-1-(7-((4-bromo-3-fluorophenyl)sulfonyl)-2,7-diazaspiro[4.4]nonan-2-yl)-2-chloroethan-1-one* **(89) 1124385**

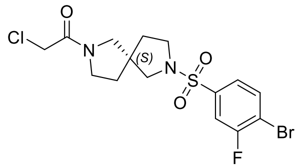

Obtained using procedure O on 0.5 mmol scale, yellow oil, 0.17 mmol, 80 mg, 34.0 % yield (over 3 steps). $^{1}$H NMR (400 MHz, $CDCl_3$) δ 7.76 (ddd, *J* = 8.3, 6.5, 3.3 Hz, 1H), 7.56 (ddd, *J* = 7.5, 5.2, 2.0 Hz, 1H), 7.55 – 7.44 (m, 1H), 3.98 (s, 1H), 3.91 (s, 1H), 3.63 – 3.50 (m, 2H), 3.49 – 3.19 (m, 5H), 3.12 (dd, *J* = 9.8, 8.3 Hz, 1H), 1.99 – 1.74 (m, 4H). $^{13}$C NMR (100 MHz, $CDCl_3$) δ 165.0, 159 (d, *J* = 253.3 Hz), 137.7 (dd, *J* = 8.8, 5.6 Hz), 134.6 (d, *J* = 2.2 Hz), 123.9 (dd, *J* = 11.5, 4.1 Hz), 115.6 – 115.4 (m), 56.1, 55.2, 54.5, 49.6, 47.4, 46.8, 45.4, 41.5, 35.2, 34.4, 33.1. LCMS (ESI): m/z calcd for $C_{15}H_{17}BrClFN_2O_3S$; found $[M+H]^+$ 441.09.

### 5. REPRESENTATIVE NMR SPECTRA

*2-chloro-N-((1-((4-fluorophenyl)sulfonyl)piperidin-4-yl)methyl)acetamide* **(11) 1076408**

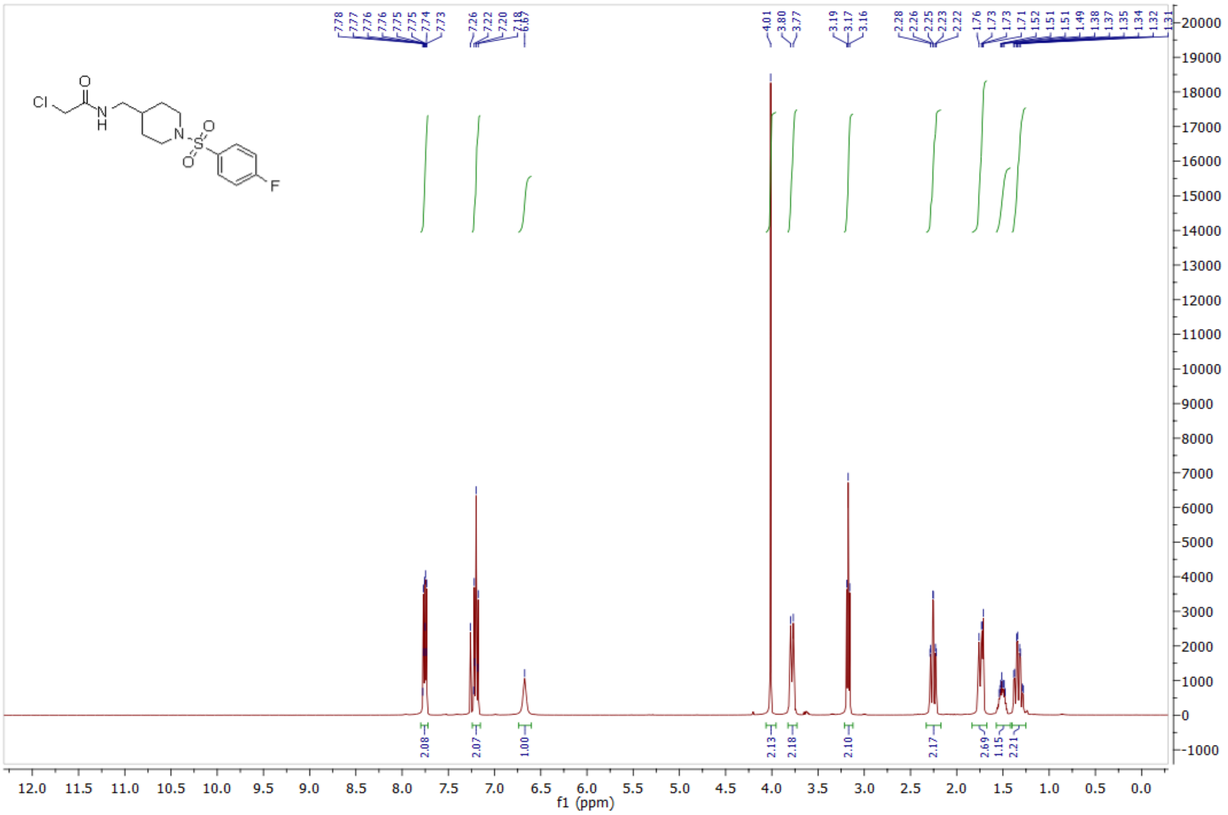

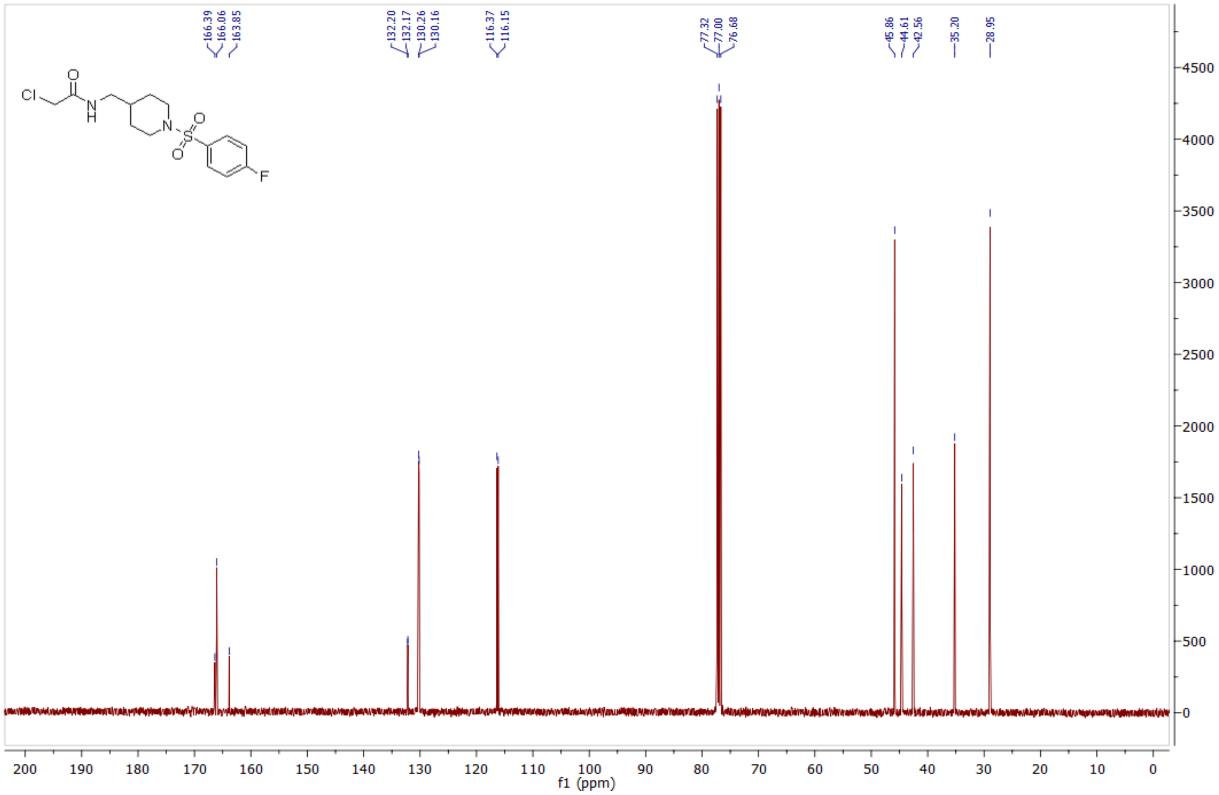

*2-chloro-N-((1-((4-chlorophenyl)sulfonyl)piperidin-4-yl)methyl)acetamide* **(12) 1075475**

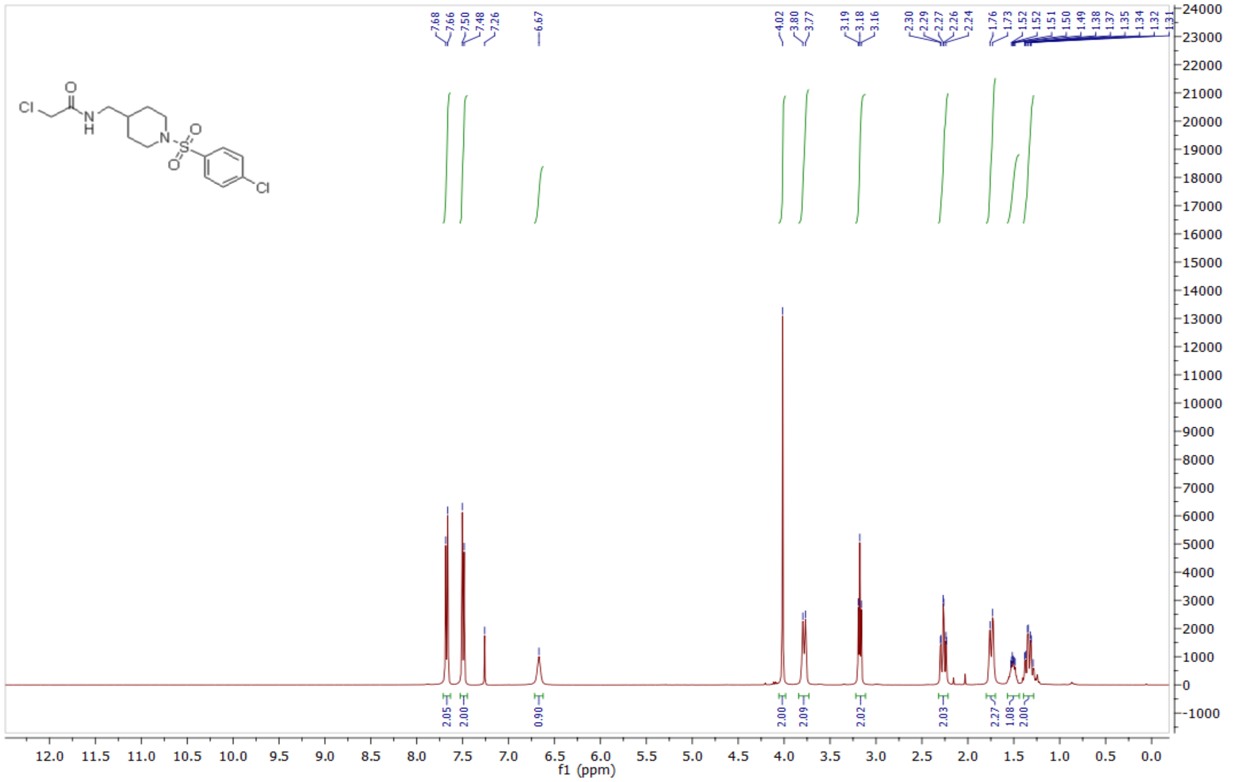

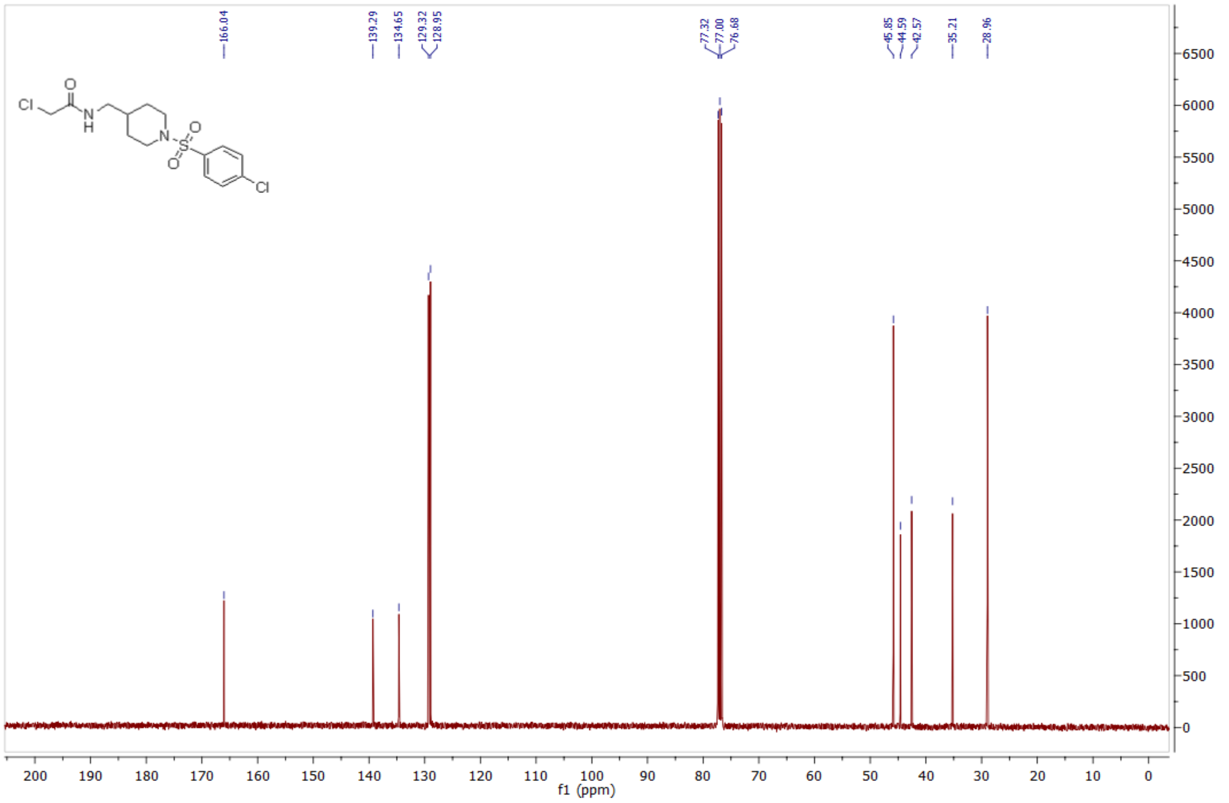

*2-chloro-N-((1-((4-cyanophenyl)sulfonyl)piperidin-4-yl)methyl)acetamide* **(20) 1083841**

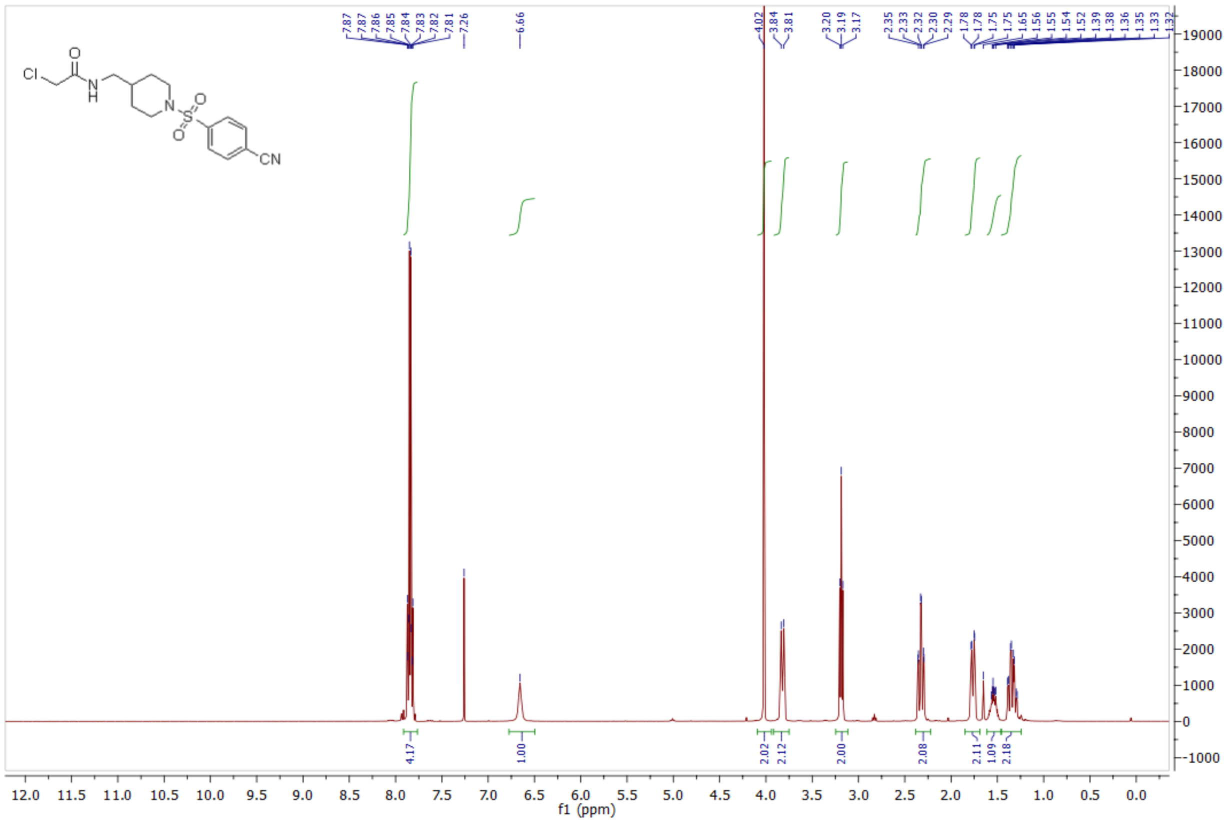

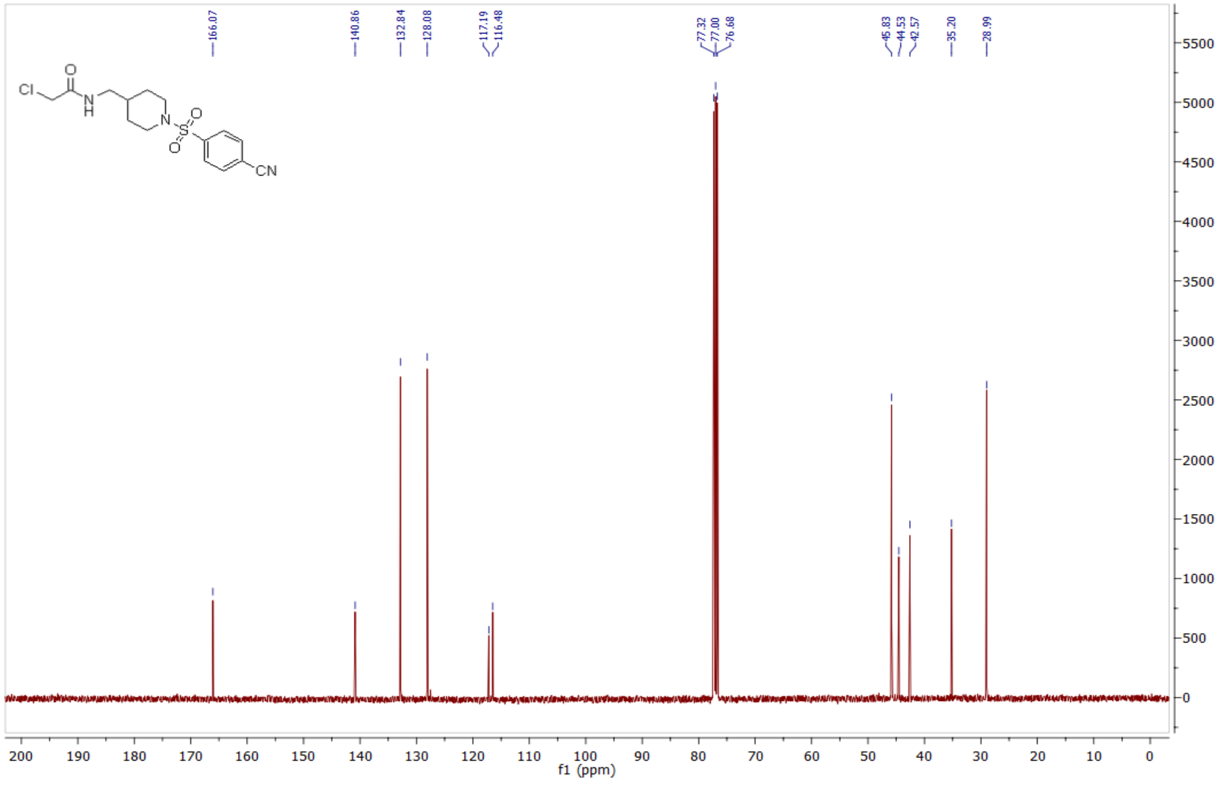

*2-chloro-N-((1-((4-formylphenyl)sulfonyl)piperidin-4-yl)methyl)acetamide* **(21) 1075354**

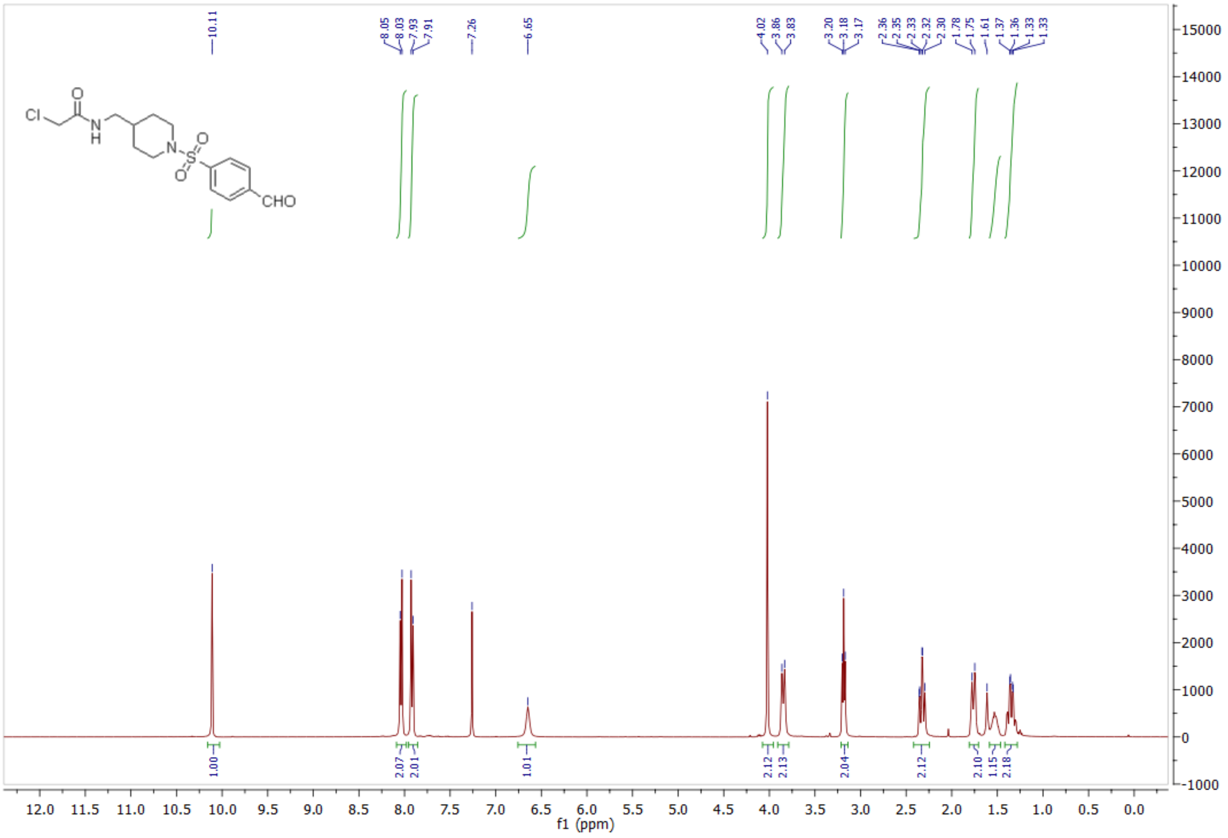

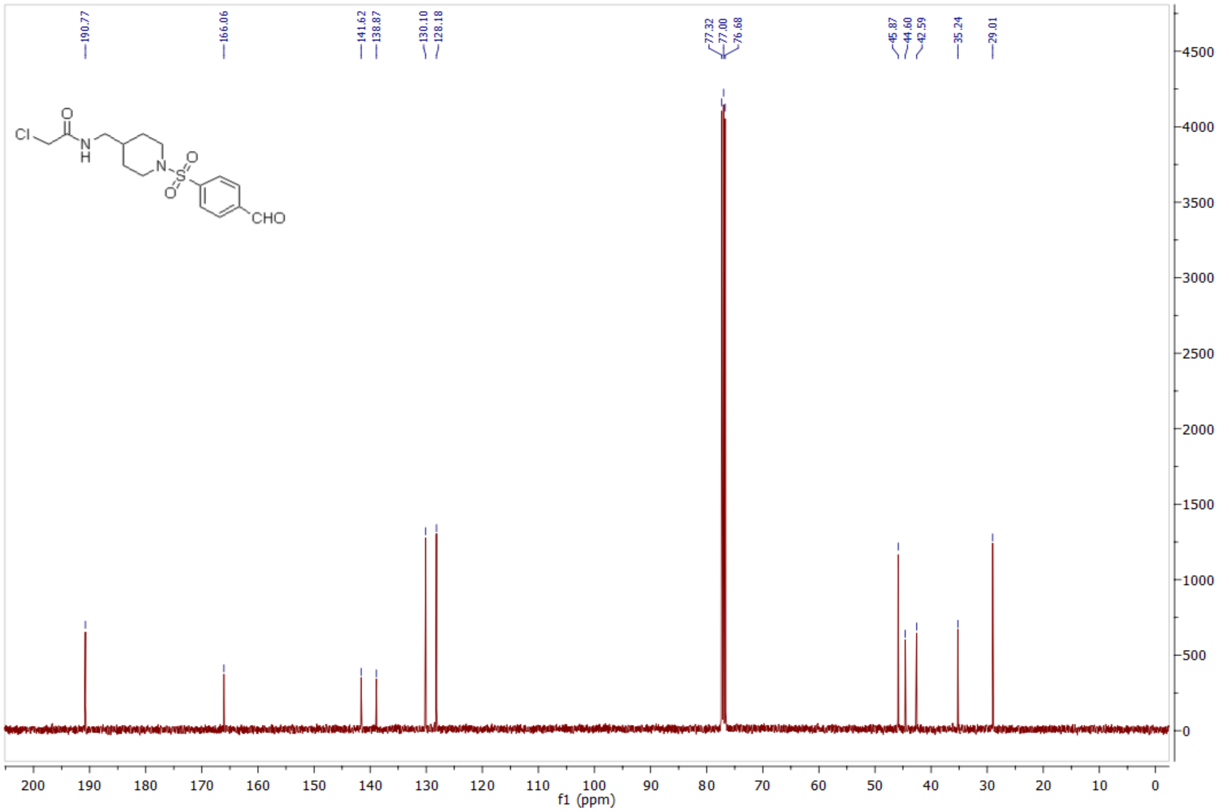

*2-chloro-N-((1-((4-bromo-phenyl)sulfonyl)piperidin-4-yl)methyl)acetamide* **(22) 1083853**

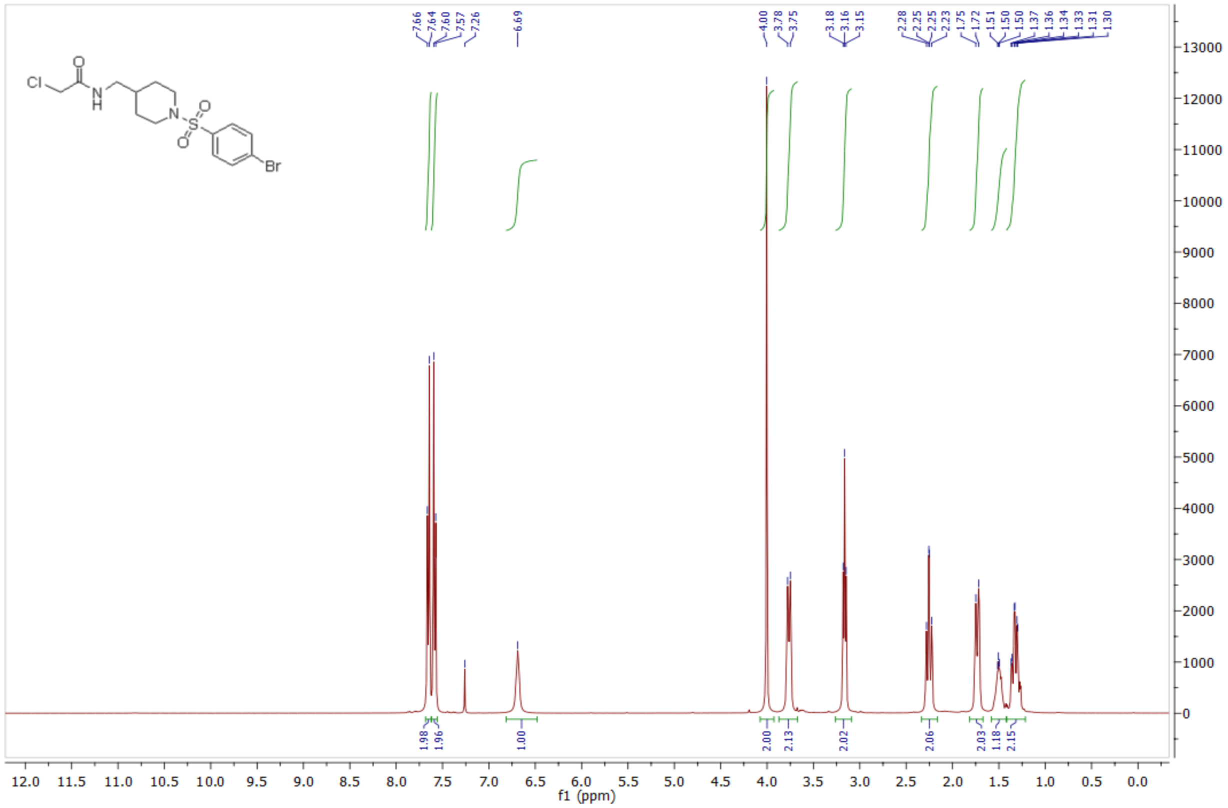

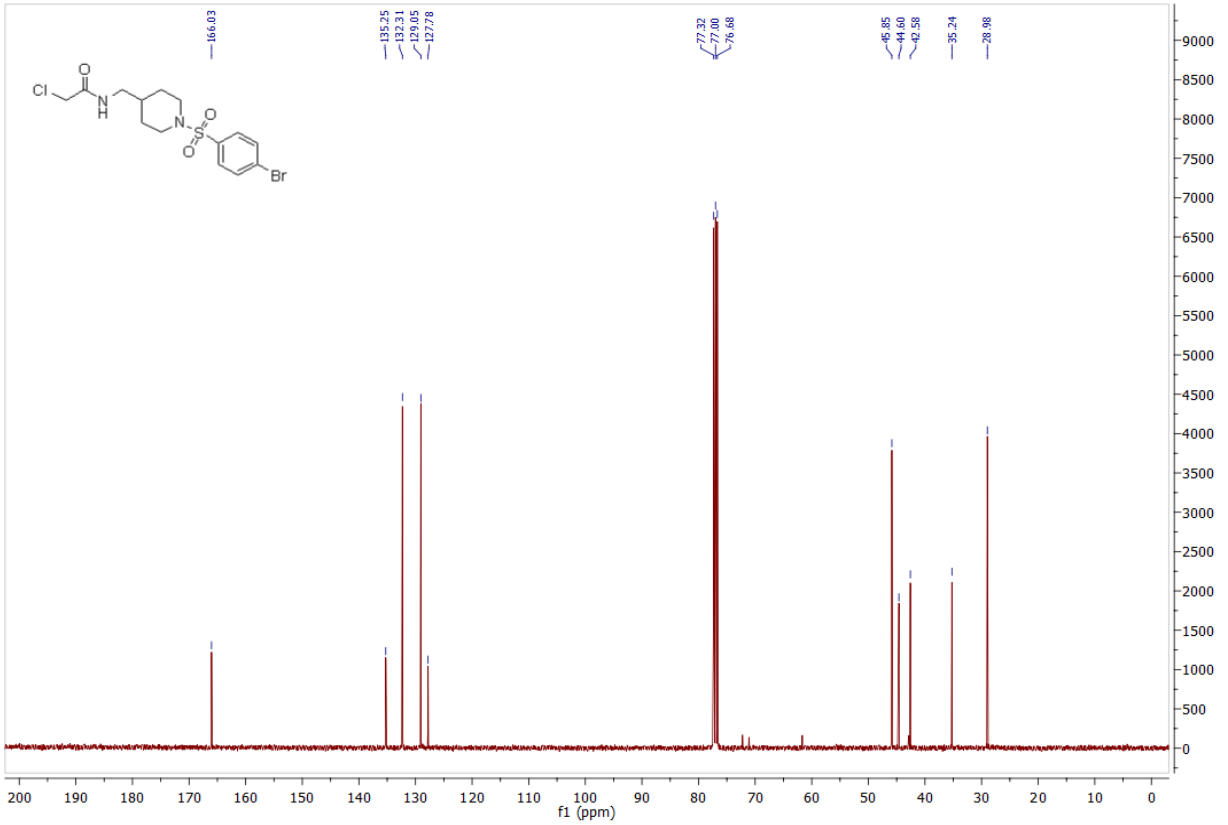

*2-chloro-N-((1-((4-iodophenyl)sulfonyl)piperidin-4-yl)methyl)acetamide* **(23) 1083848**

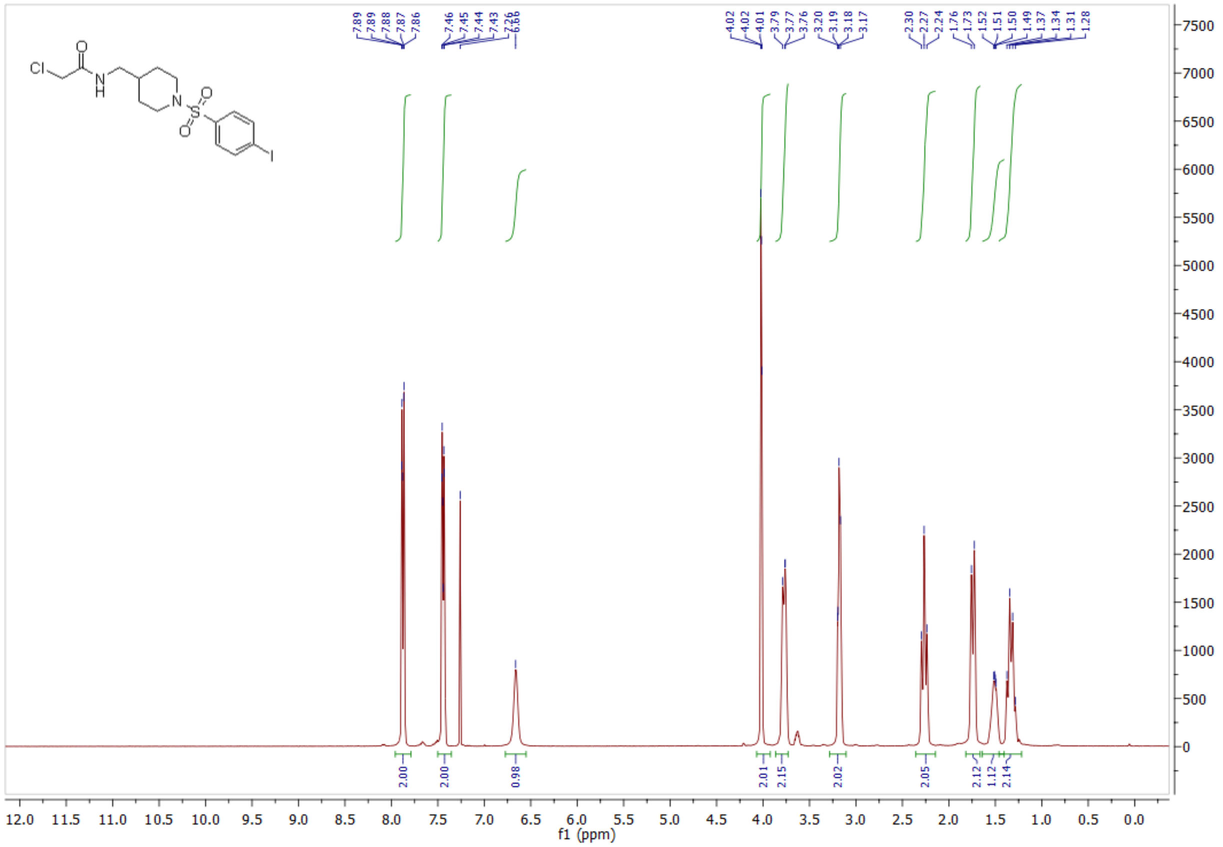

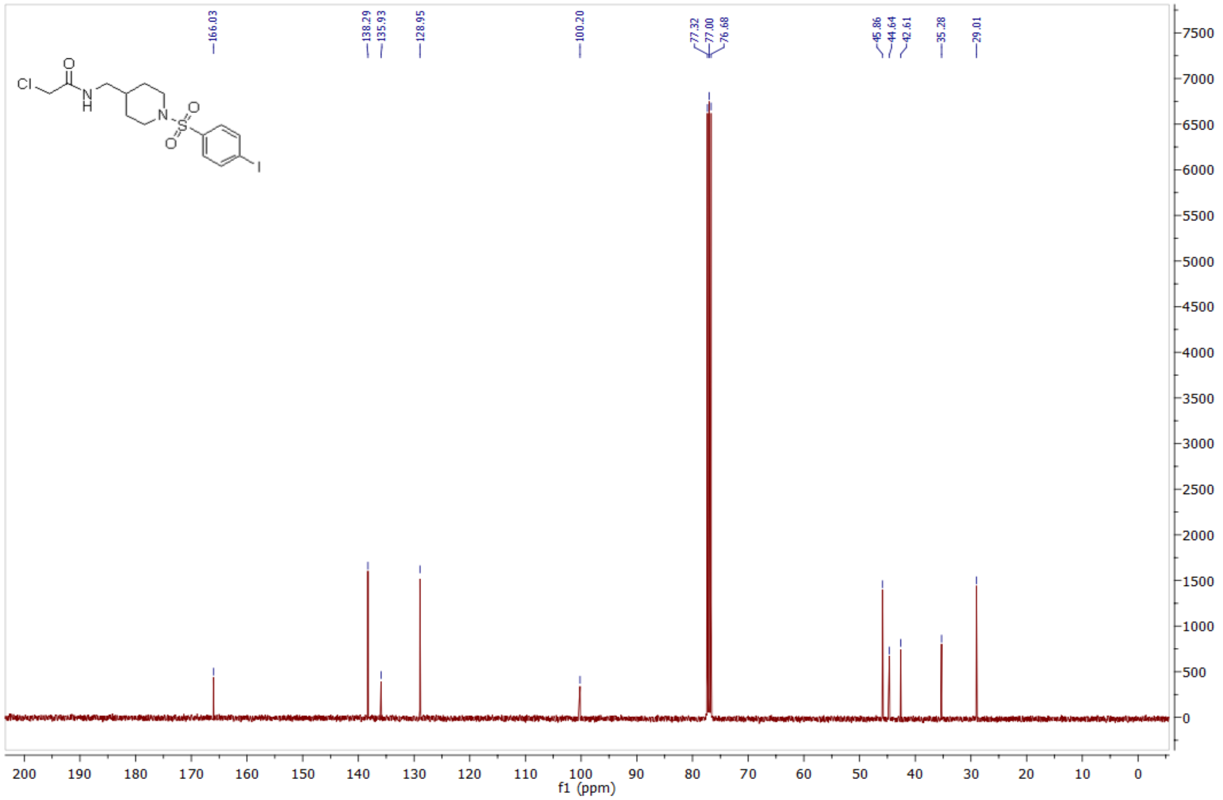

*2-chloro-N-((1-tosylpiperidin-4-yl)methyl)acetamide* **(24) 1083852**

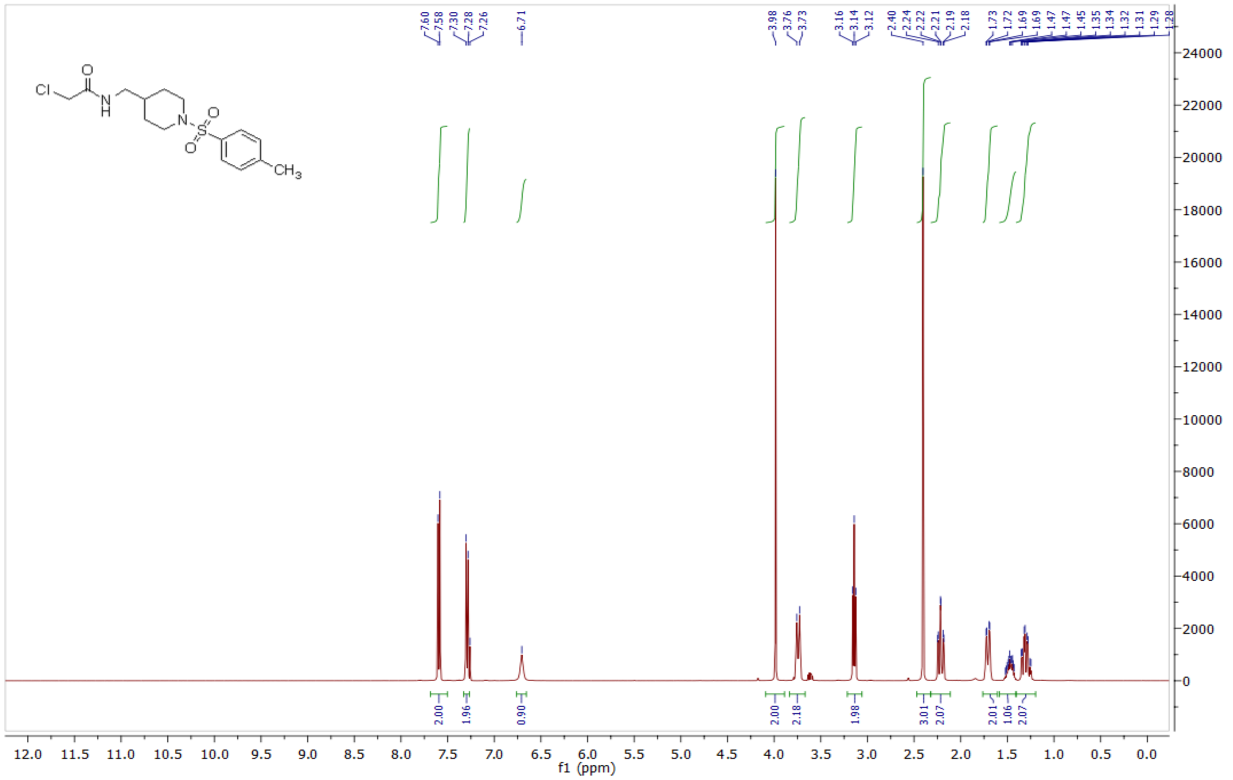

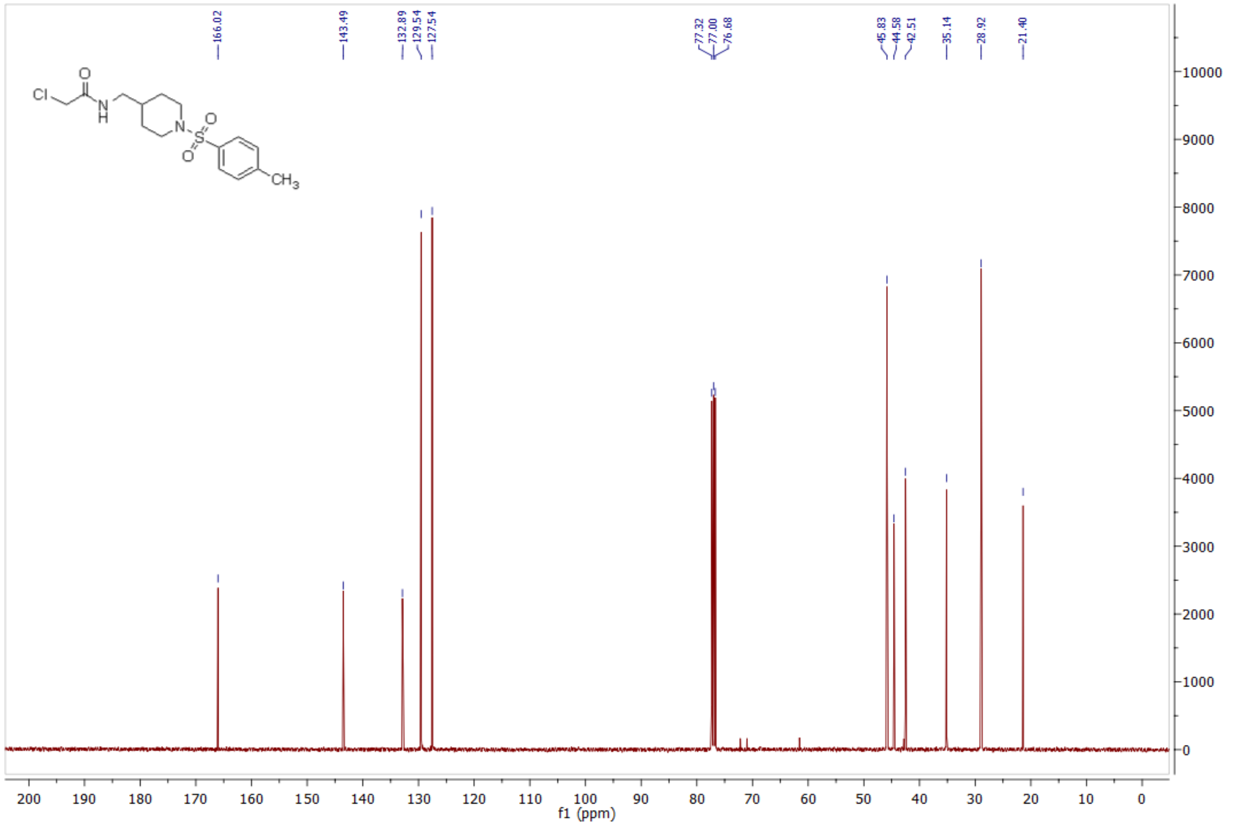

*2-chloro-N-((1-((4-chloro-3-fluorophenyl)sulfonyl)piperidin-4-yl)methyl)acetamide* **(29) 1076409**

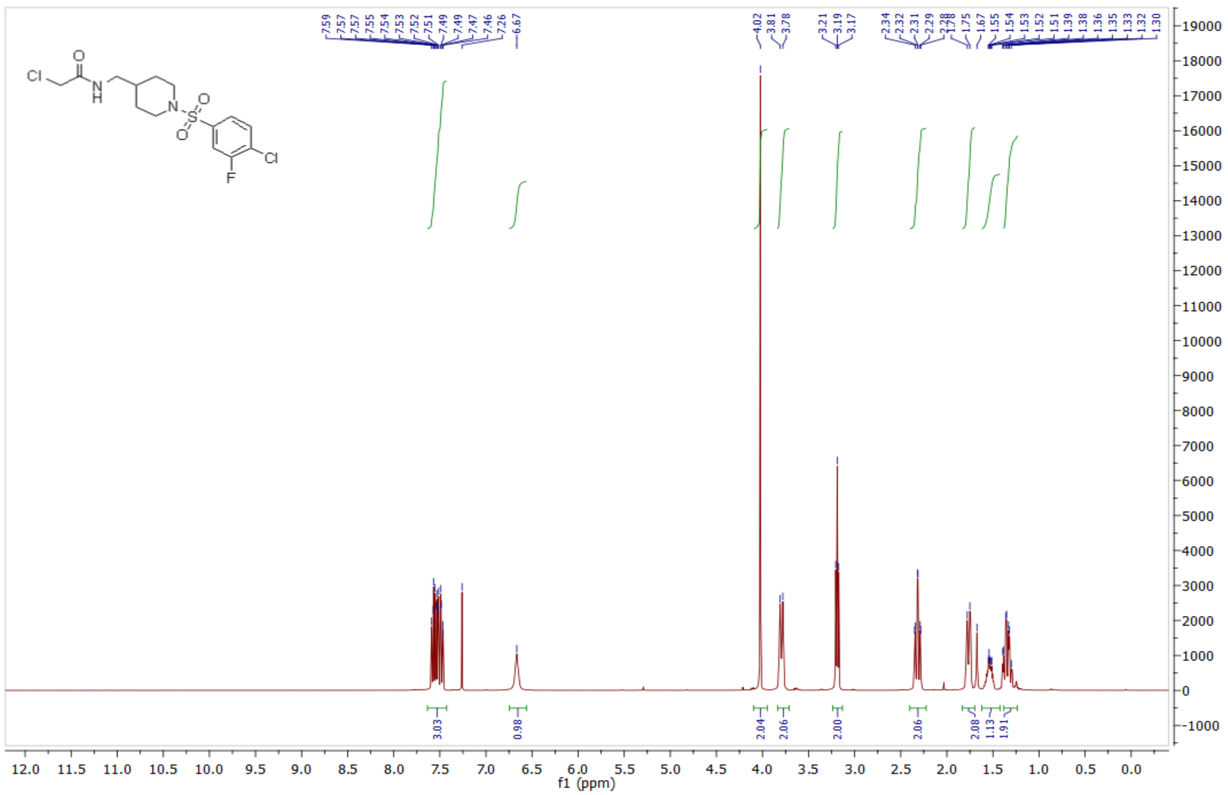

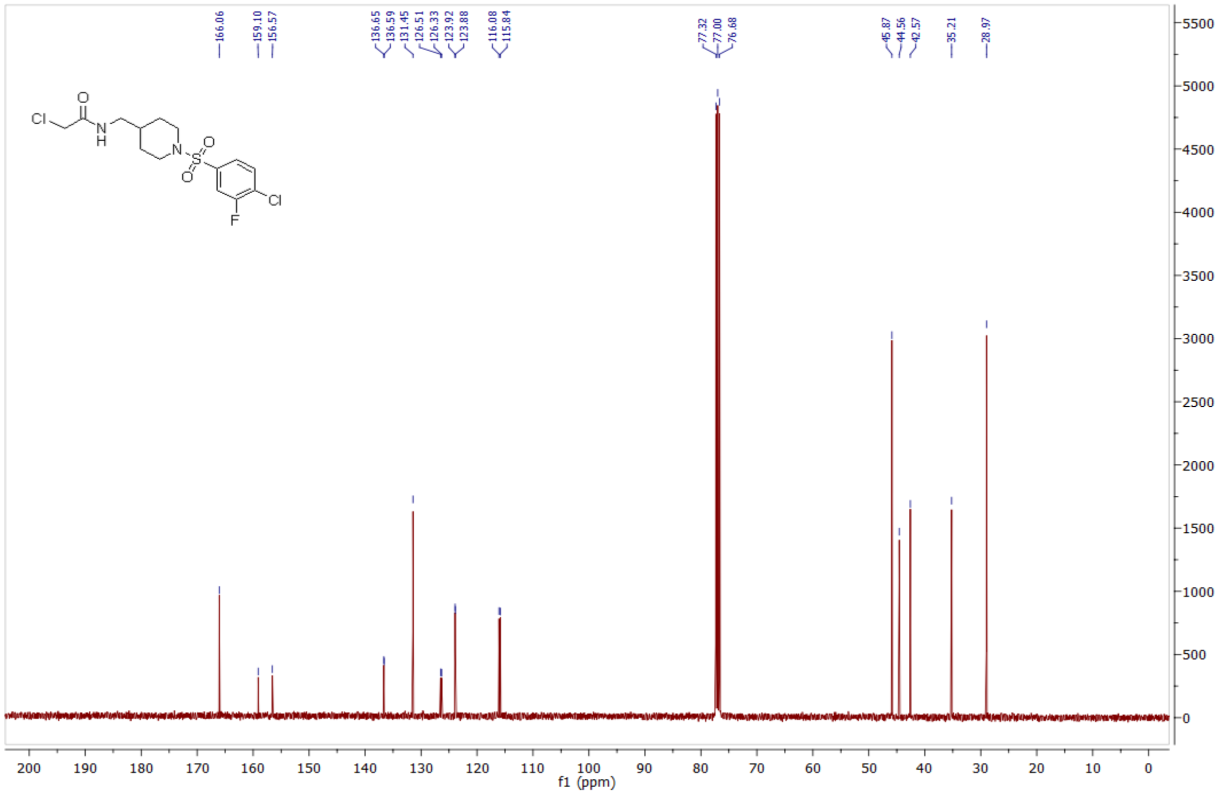

*2-chloro-N-((1-((3-chloro-4-fluorophenyl)sulfonyl)piperidin-4-yl)methyl)acetamide* **(30) 1076410**

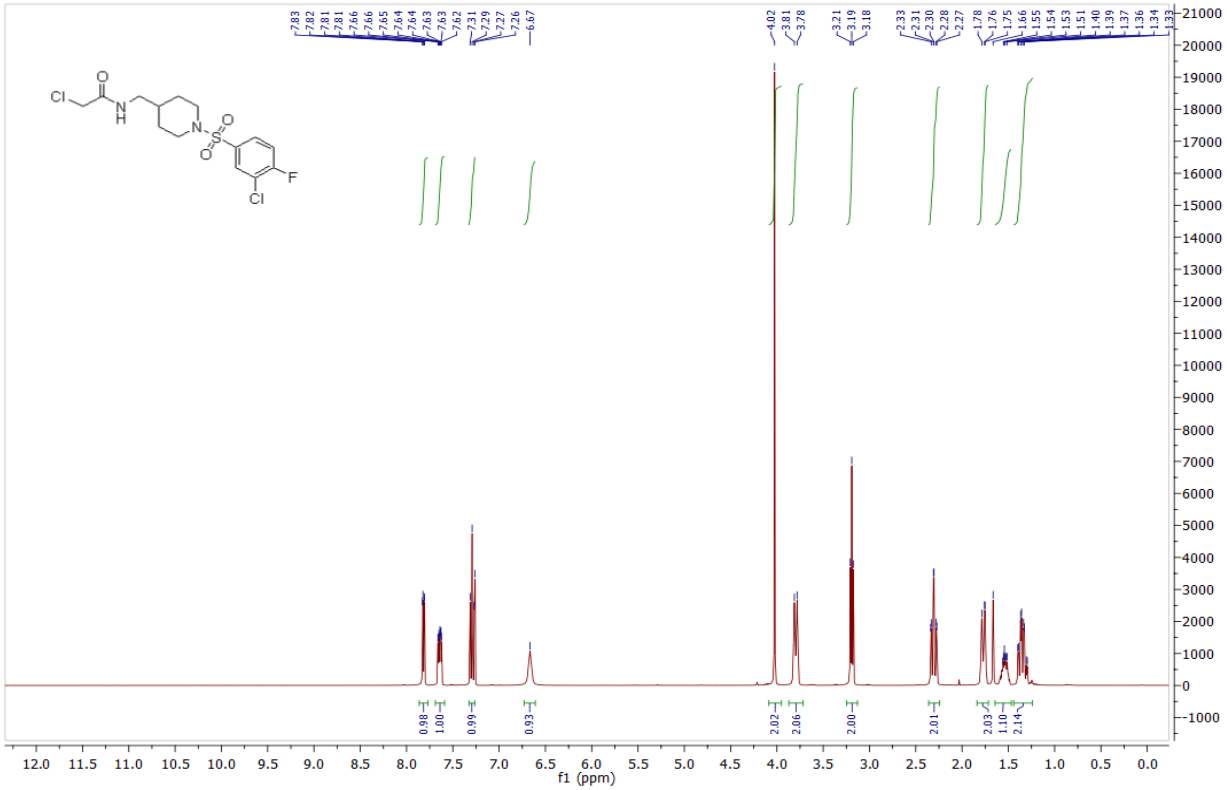

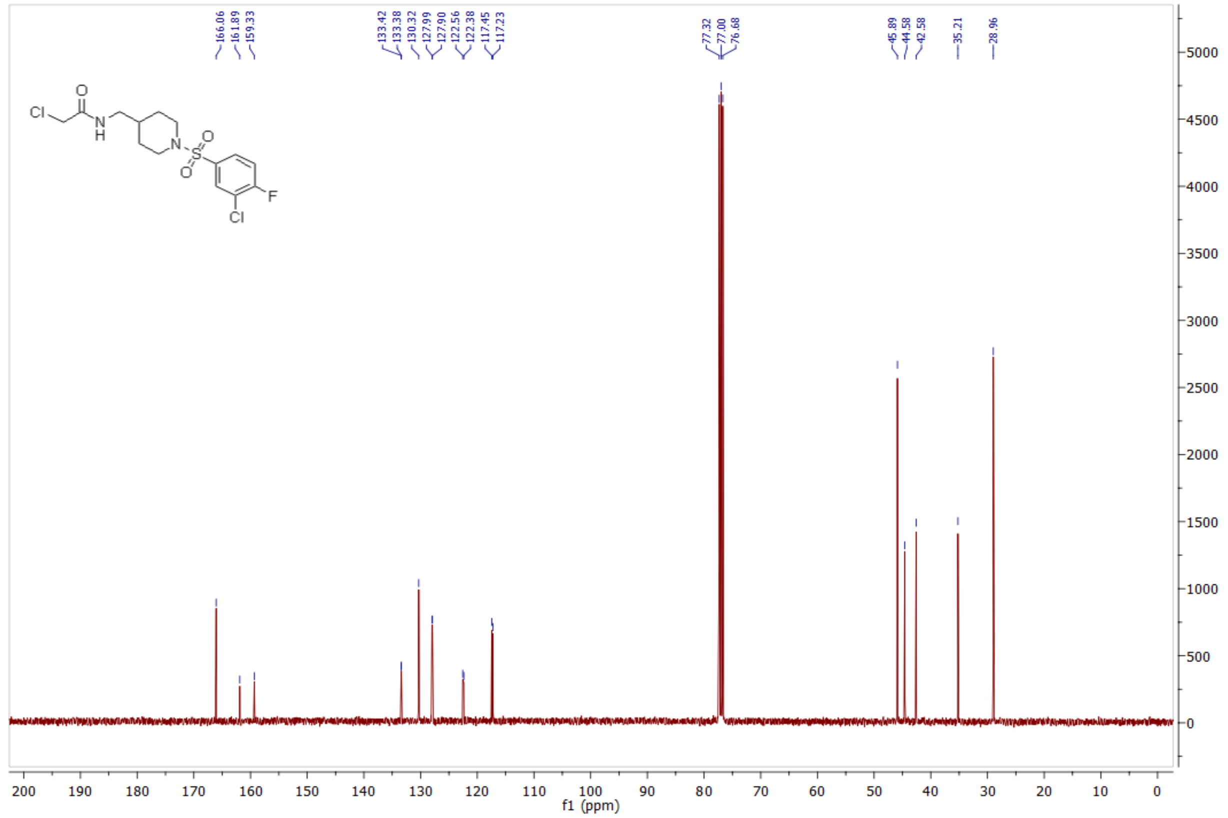

*2-chloro-N-((1-((3-fluoro-4-(trifluoromethyl)phenyl)sulfonyl)piperidin-4-yl)methyl)acetamide* **(32) 1083854**

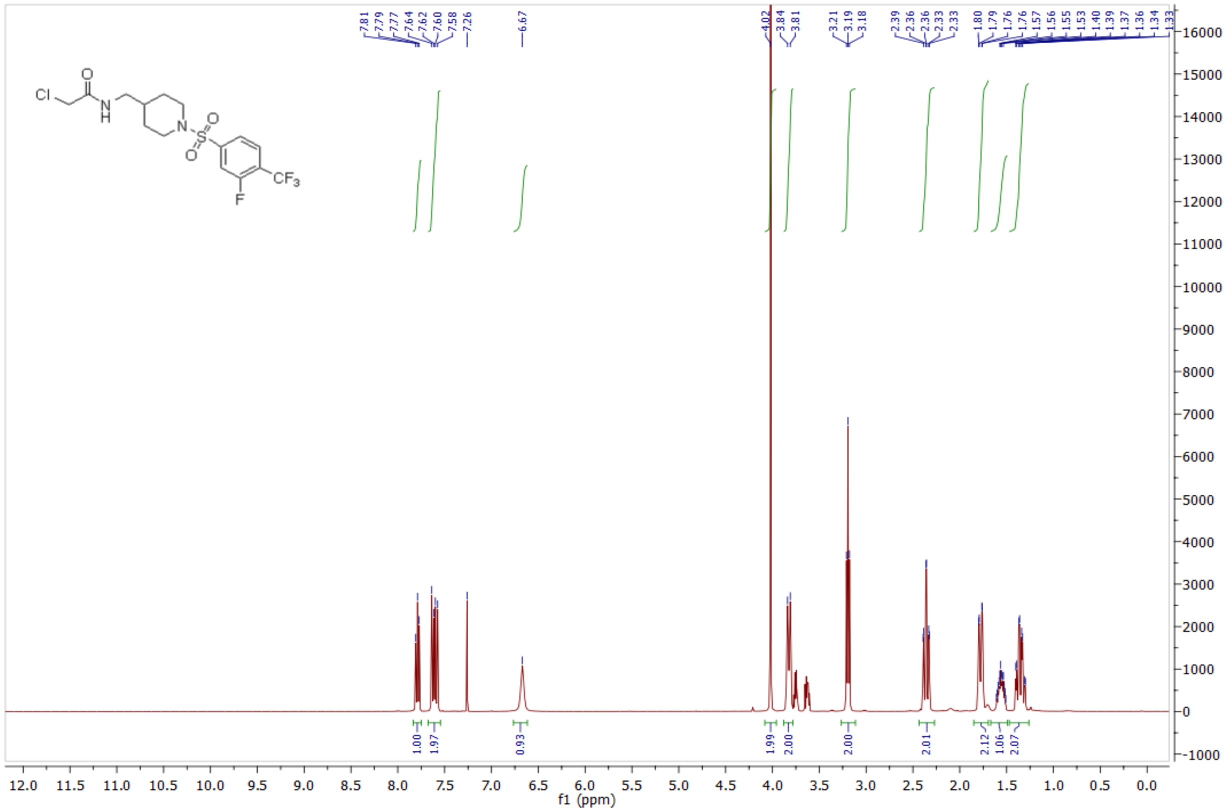

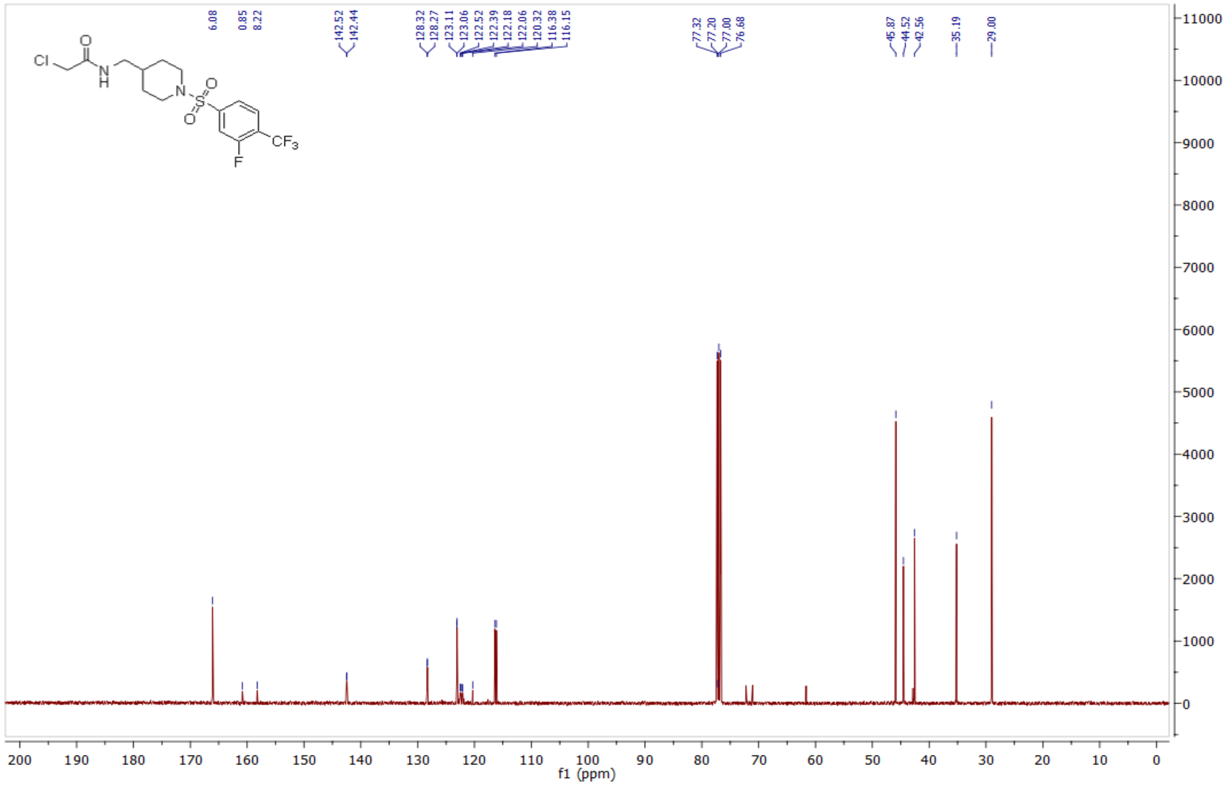

*2-chloro-N-((1-((3-fluoro-4-methoxyphenyl)sulfonyl)piperidin-4-yl)methyl)acetamide* **(33) 1083847**

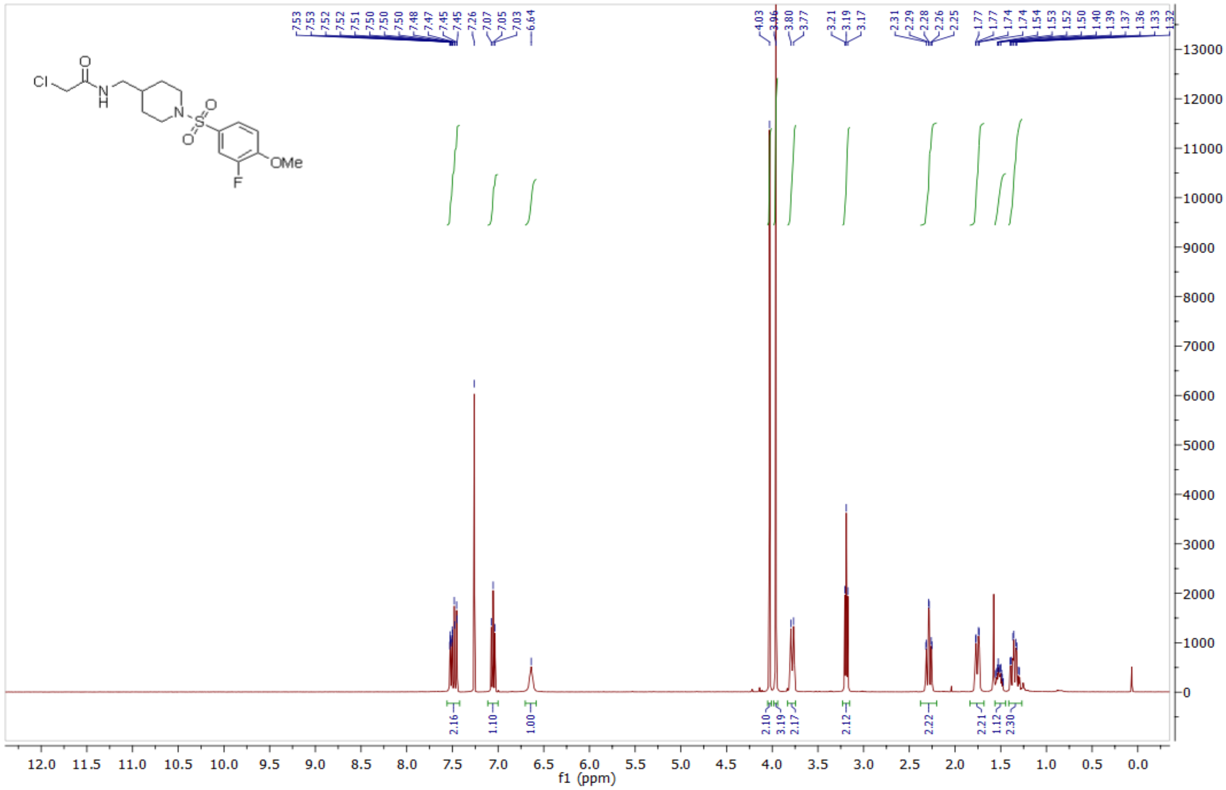

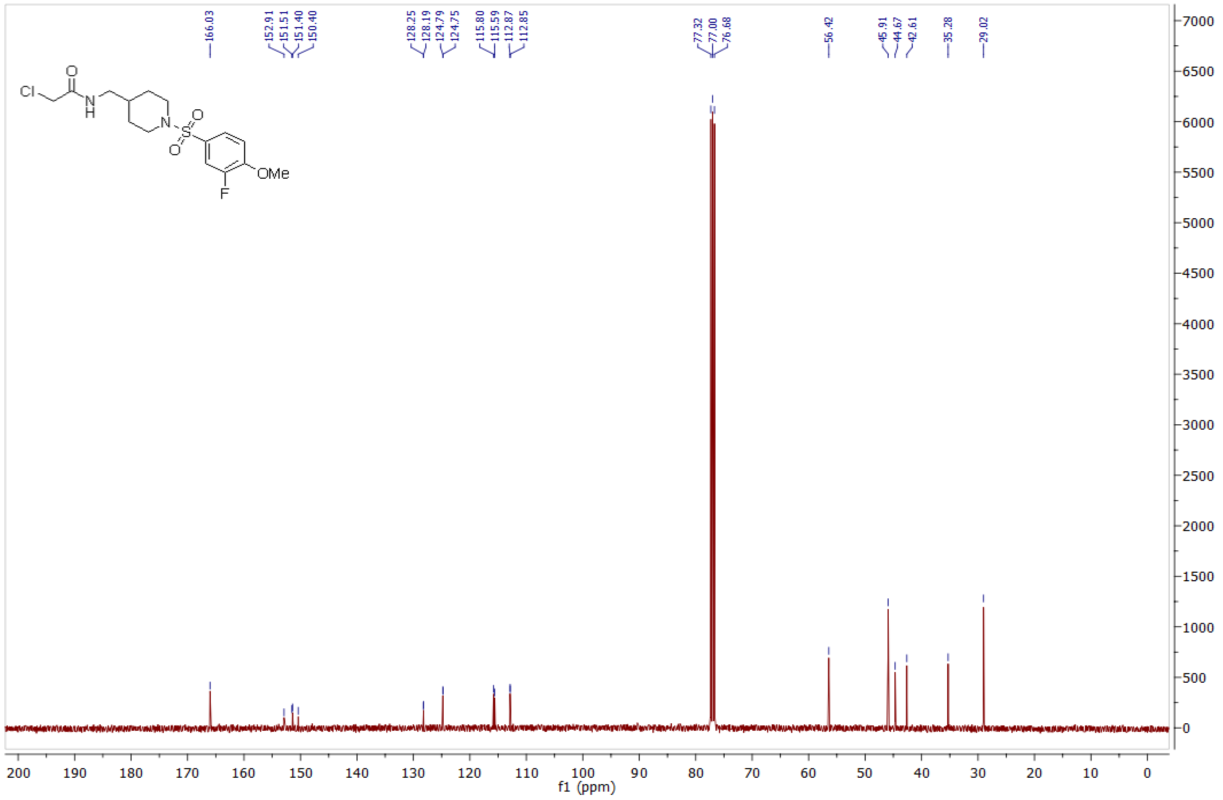

*N-((1-((4-bromo-2-fluorophenyl)sulfonyl)piperidin-4-yl)methyl)-2-chloroacetamide* **(36) 1083916**

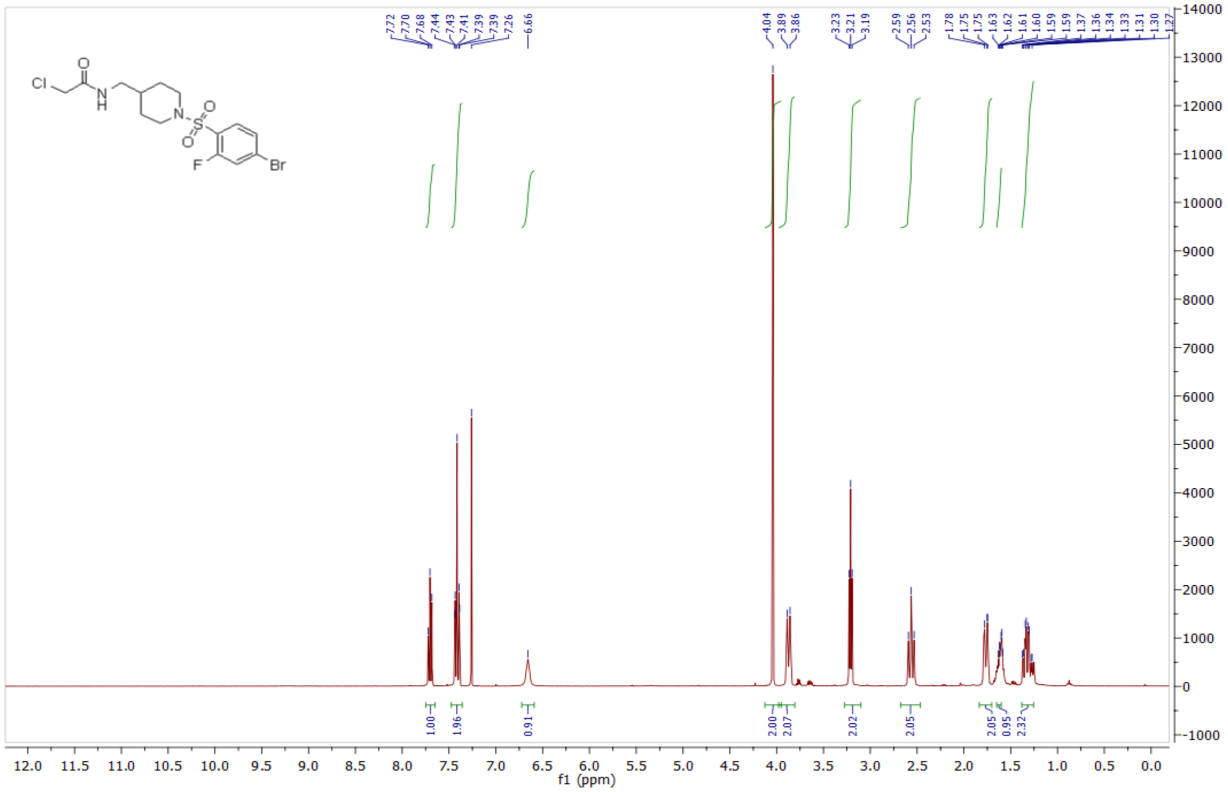

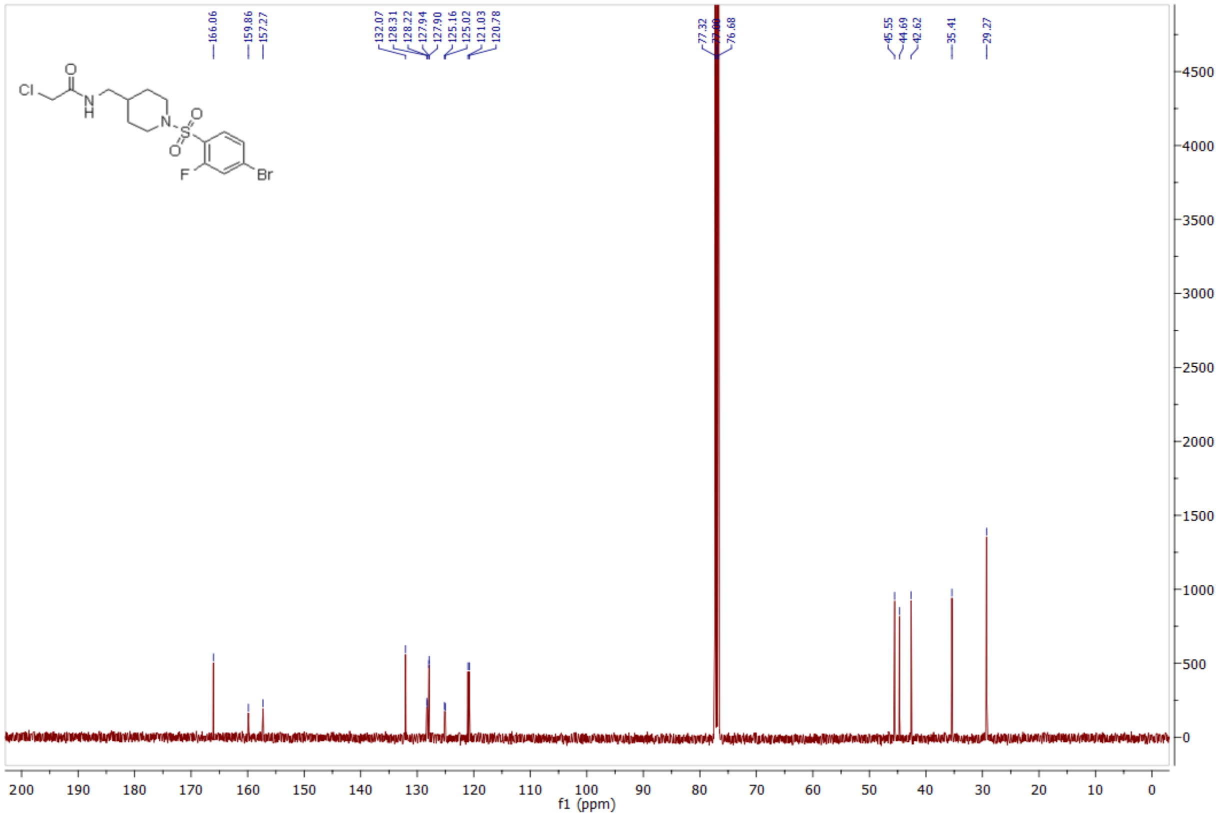

*N-((1-((4-bromo-3-fluorophenyl)sulfonyl)piperidin-4-yl)methyl)-2-chloroacetamide* **(37) 1083917**

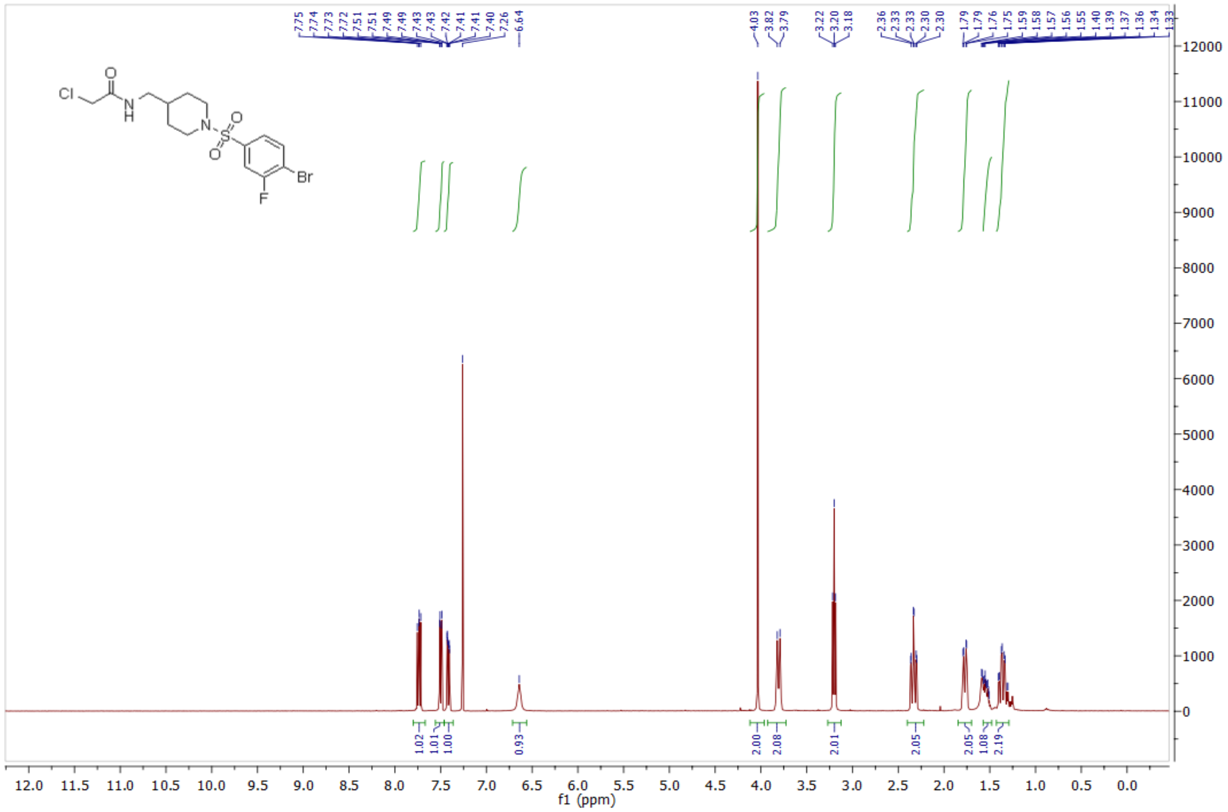

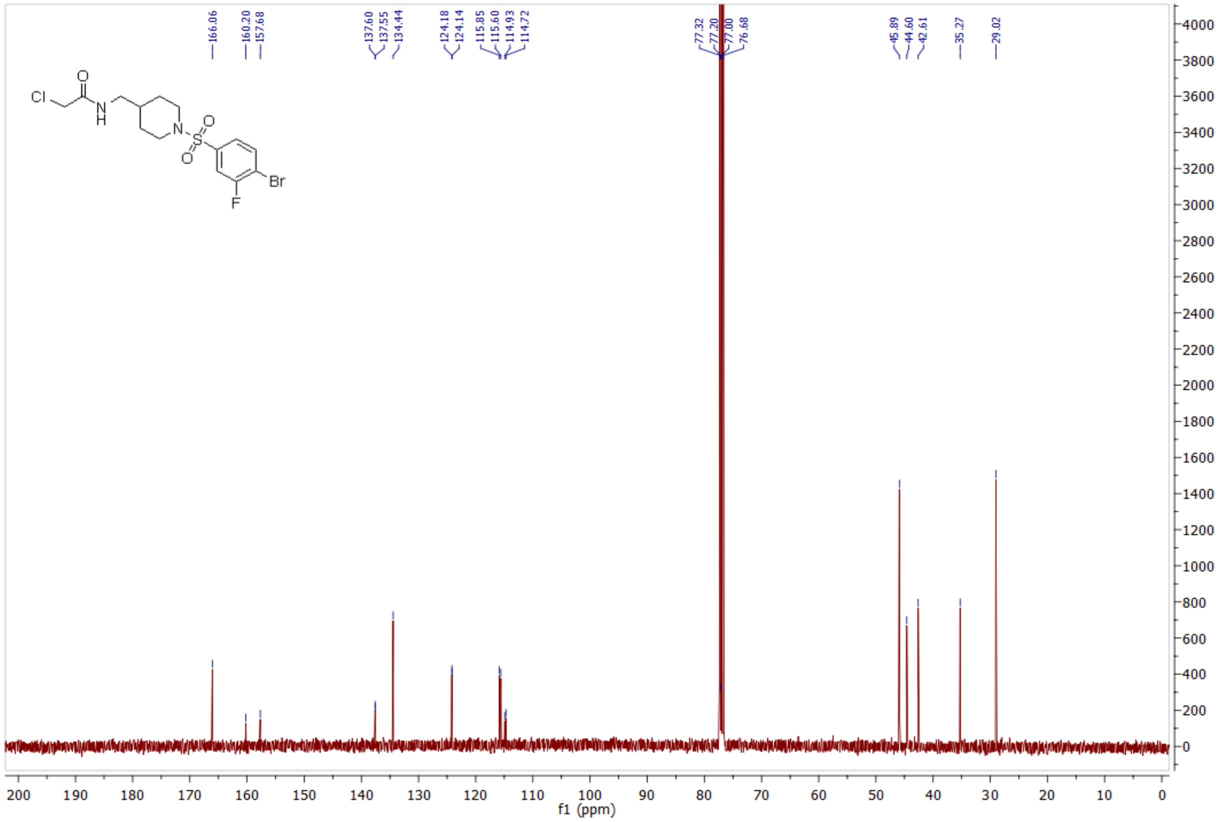

*N-((1-((4-bromo-2,3-difluorophenyl)sulfonyl)piperidin-4-yl)methyl)-2-chloroacetamide* **(38) 1083918**

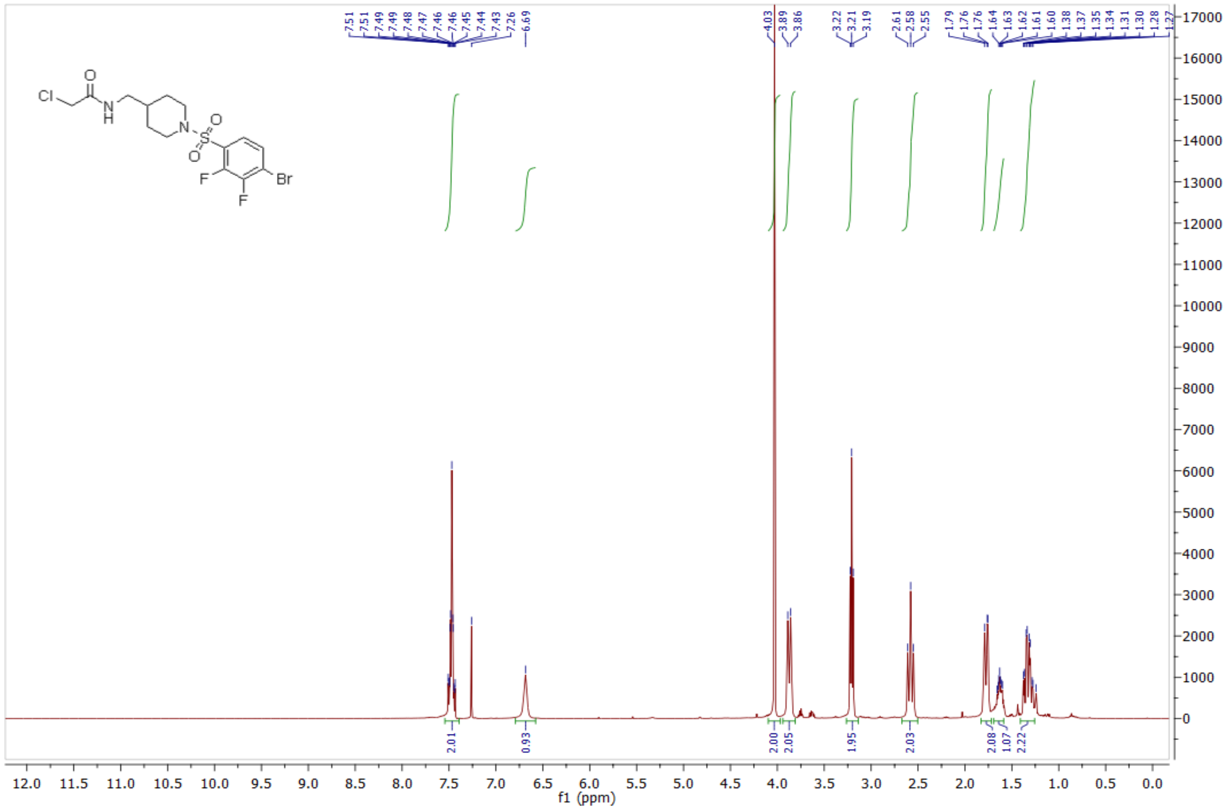

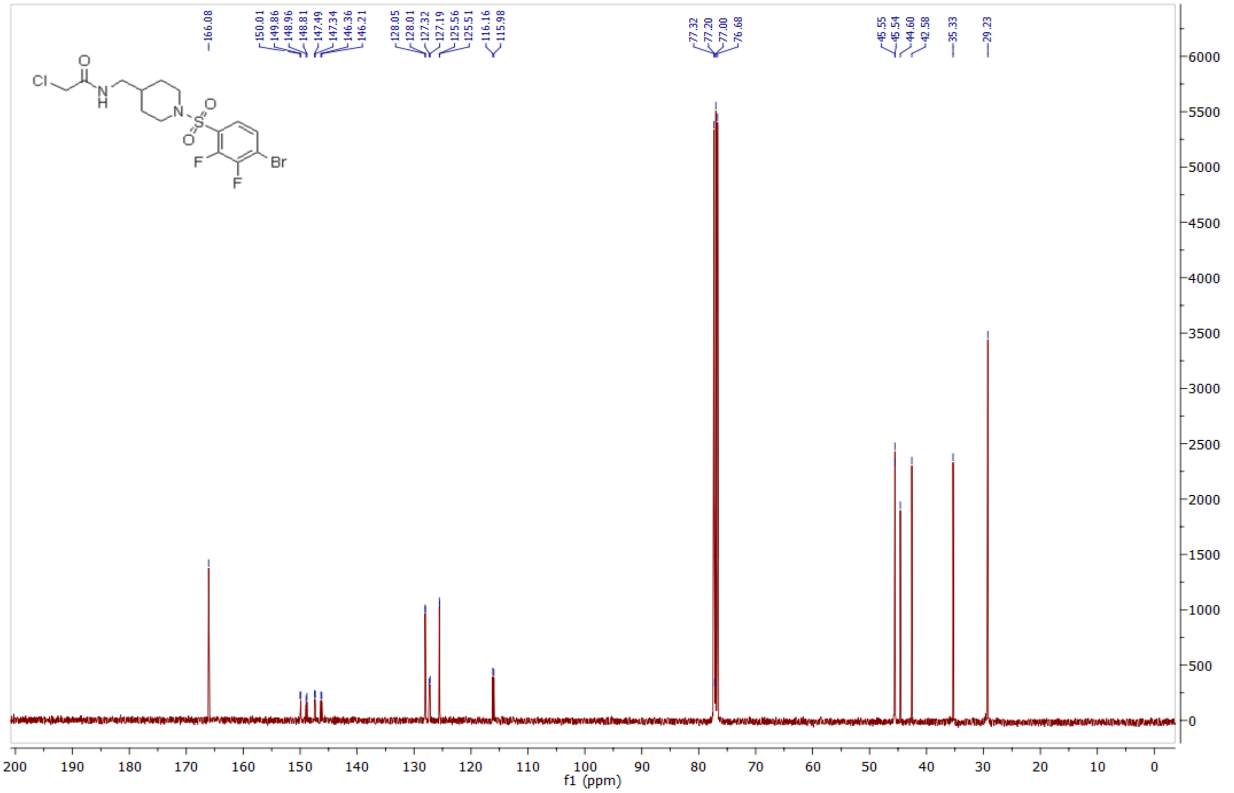

*N-((1-((4-bromo-2,5-difluorophenyl)sulfonyl)piperidin-4-yl)methyl)-2-chloroacetamide* **(39) 1083919**

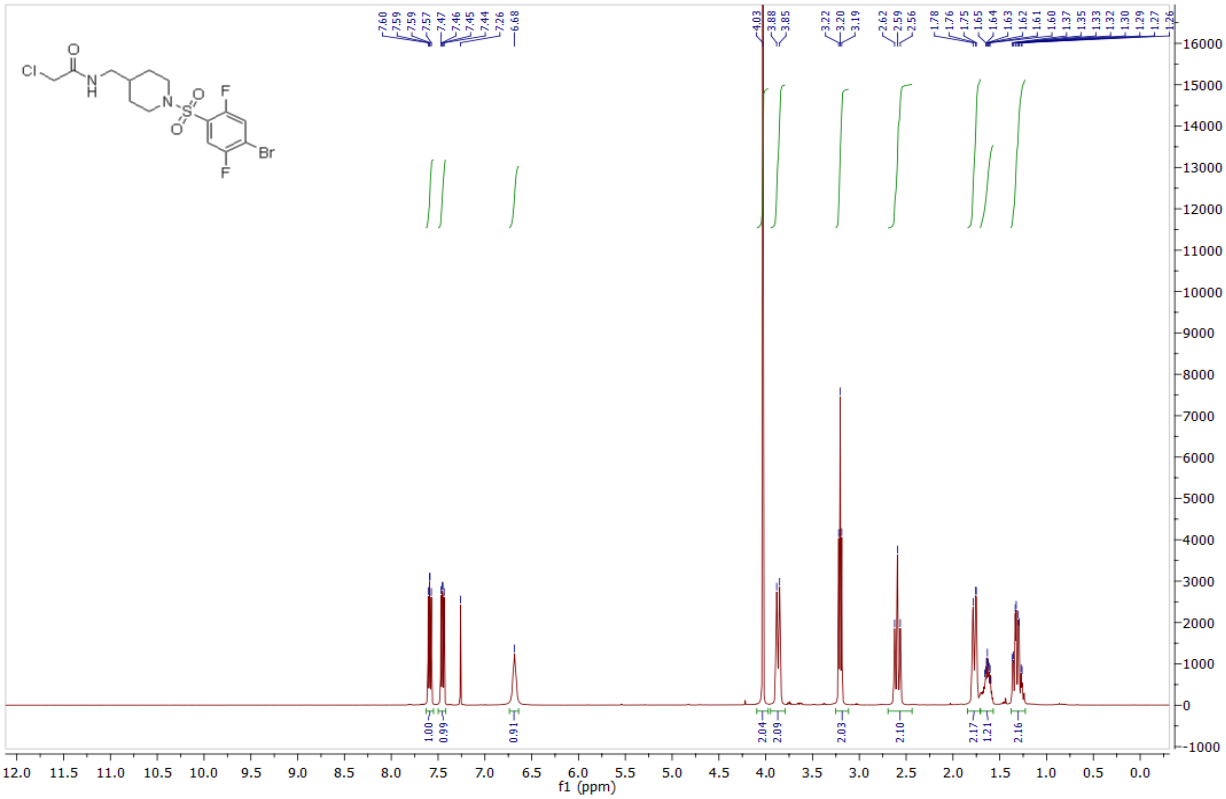

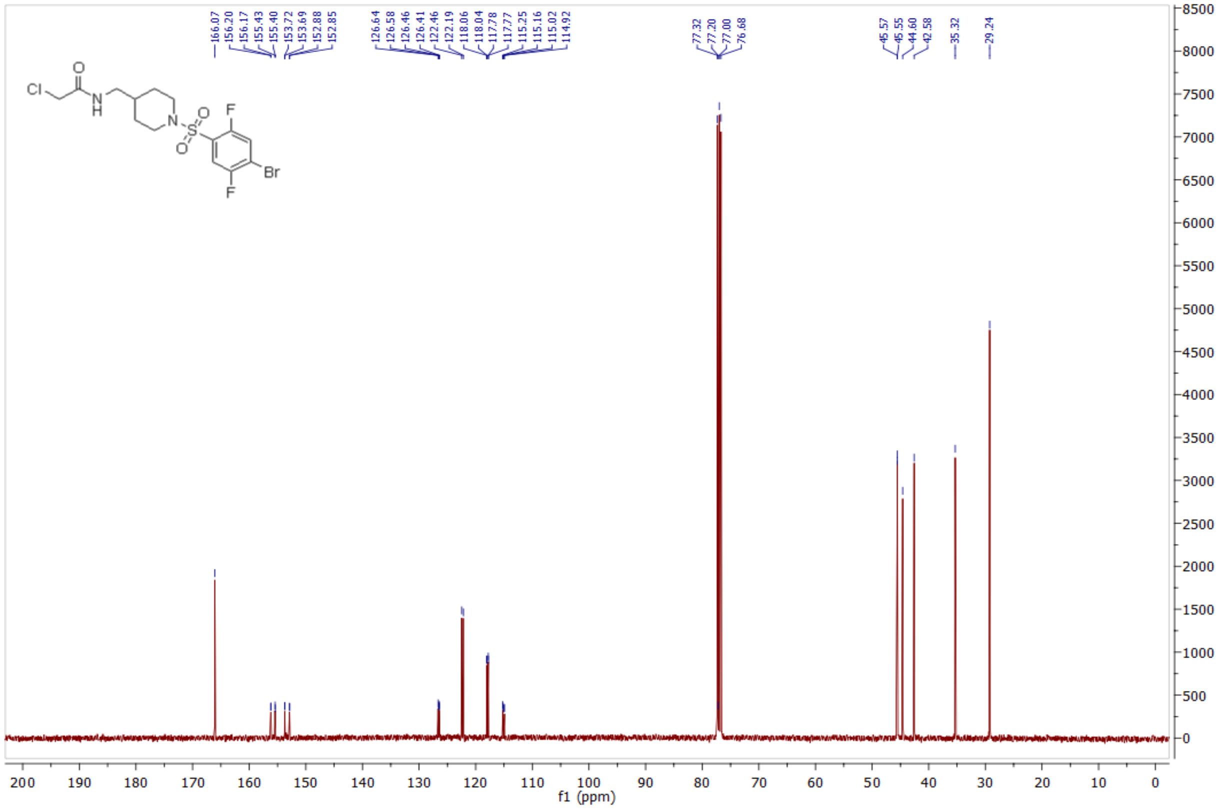

*N-((1-((4-bromo-3,5-difluorophenyl)sulfonyl)piperidin-4-yl)methyl)-2-chloroacetamide* **(40) 1083920**

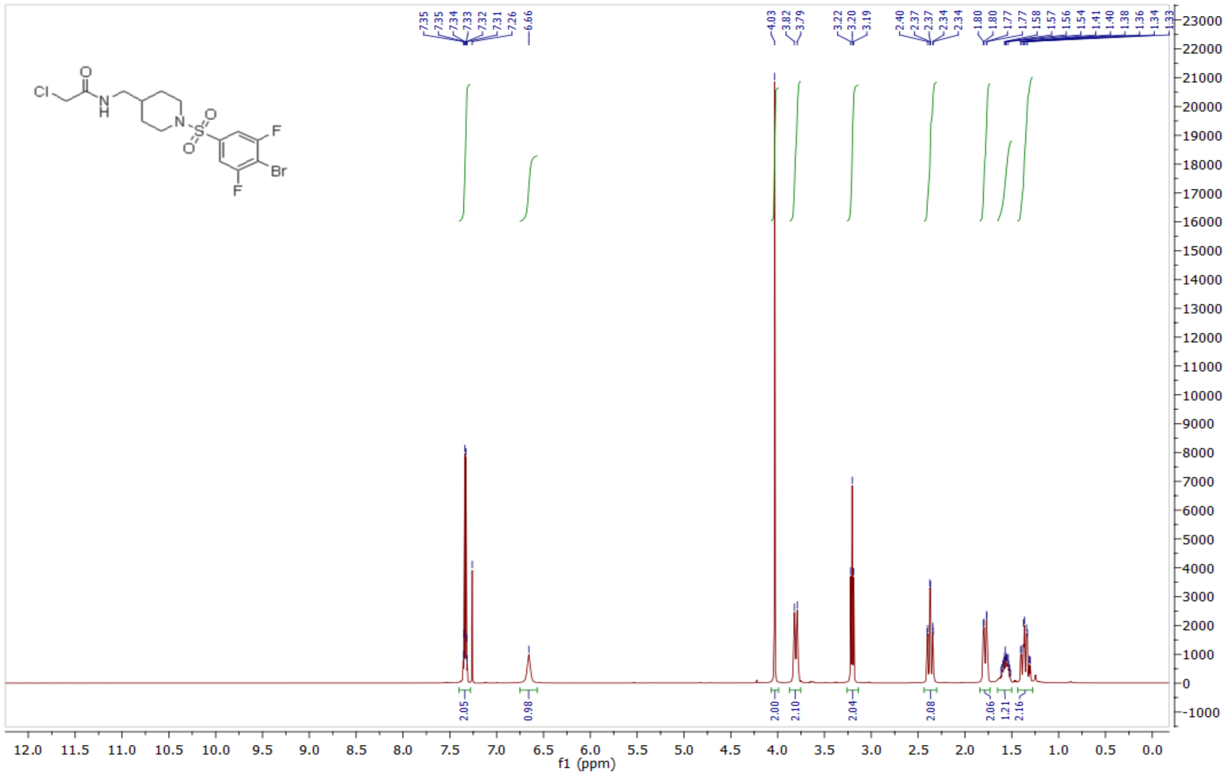

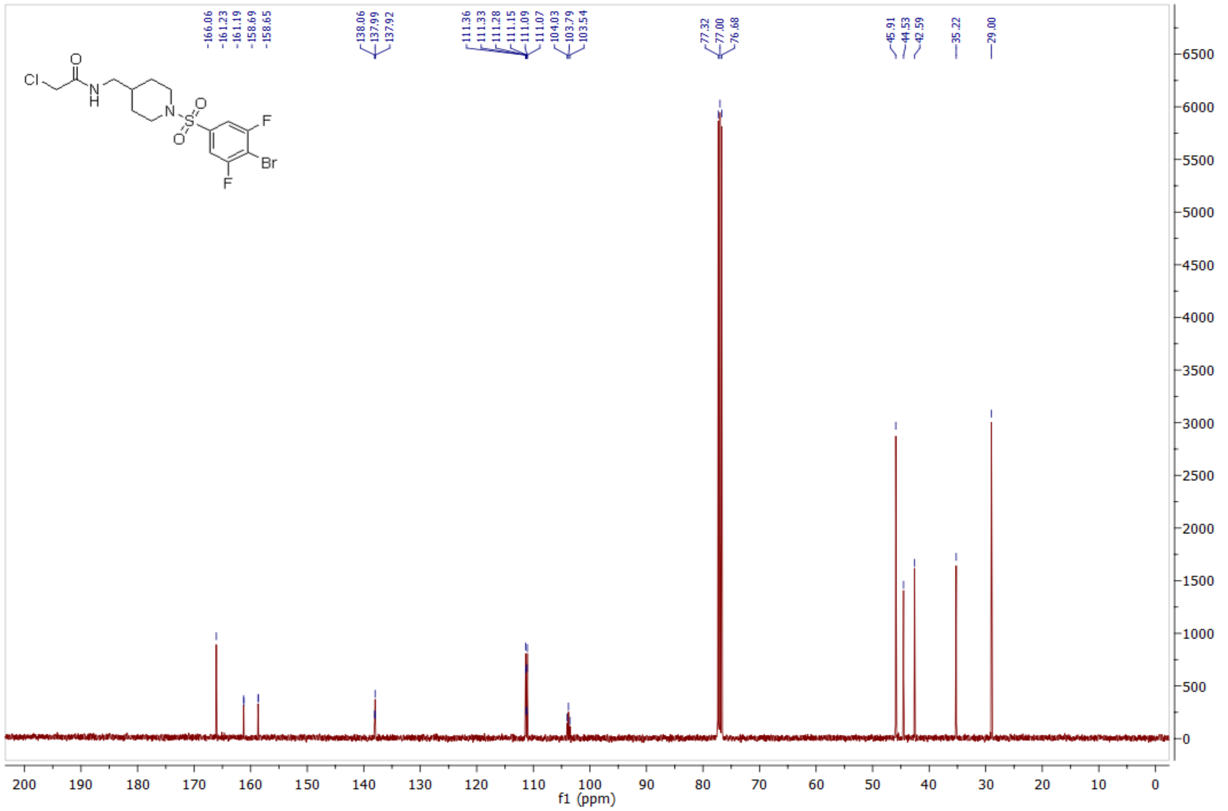

*N-(1-((4-bromo-3-fluorophenyl)sulfonyl)-4-methylpiperidin-4-yl)-2-chloroacetamide* **(43) 1124381**

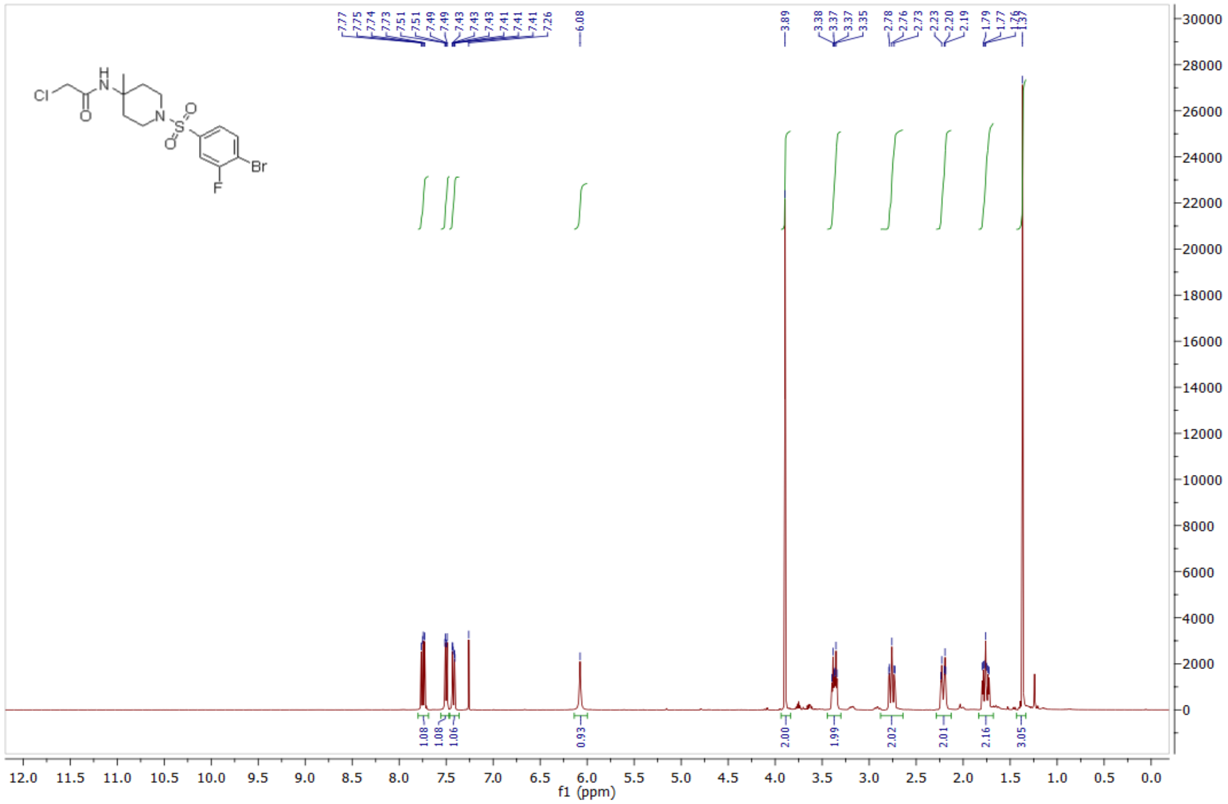

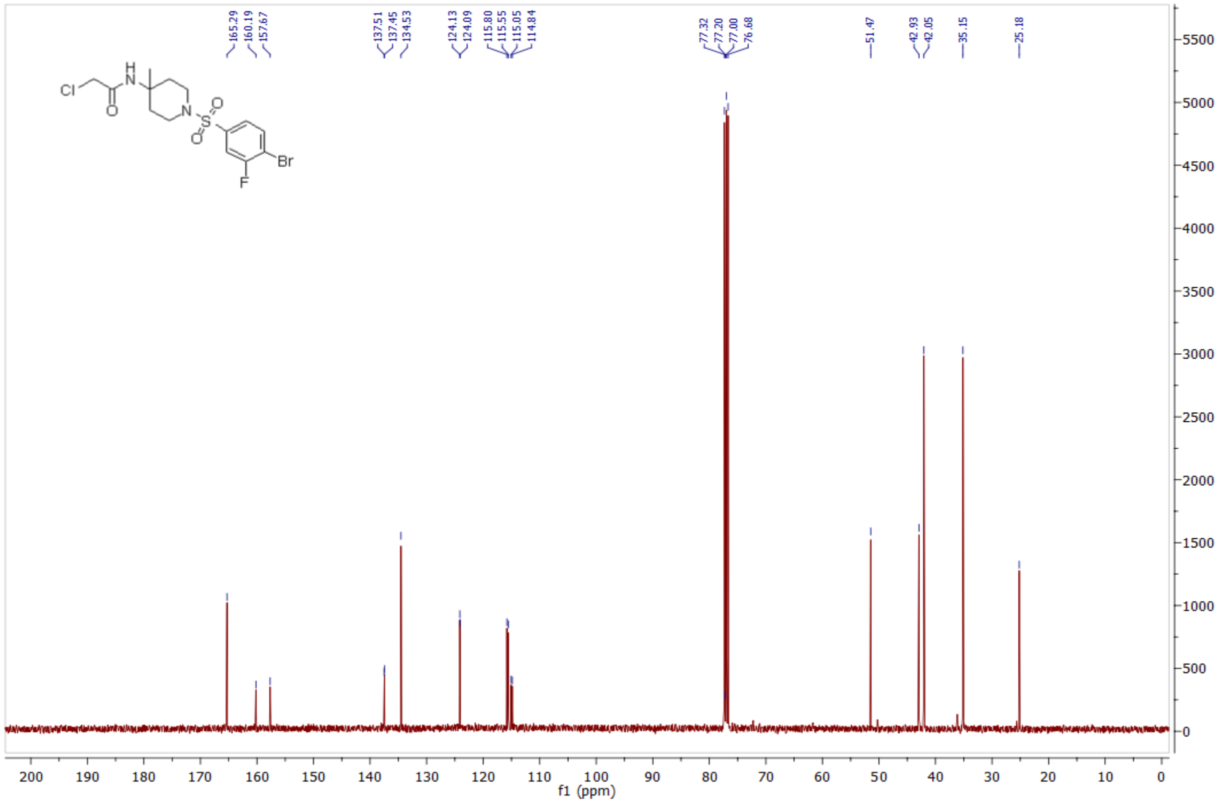

*1-(7-((4-bromo-3-fluorophenyl)sulfonyl)-2,7-diazaspiro[3.5]nonan-2-yl)-2-chloroethan-1-one* **(49) 1084346**

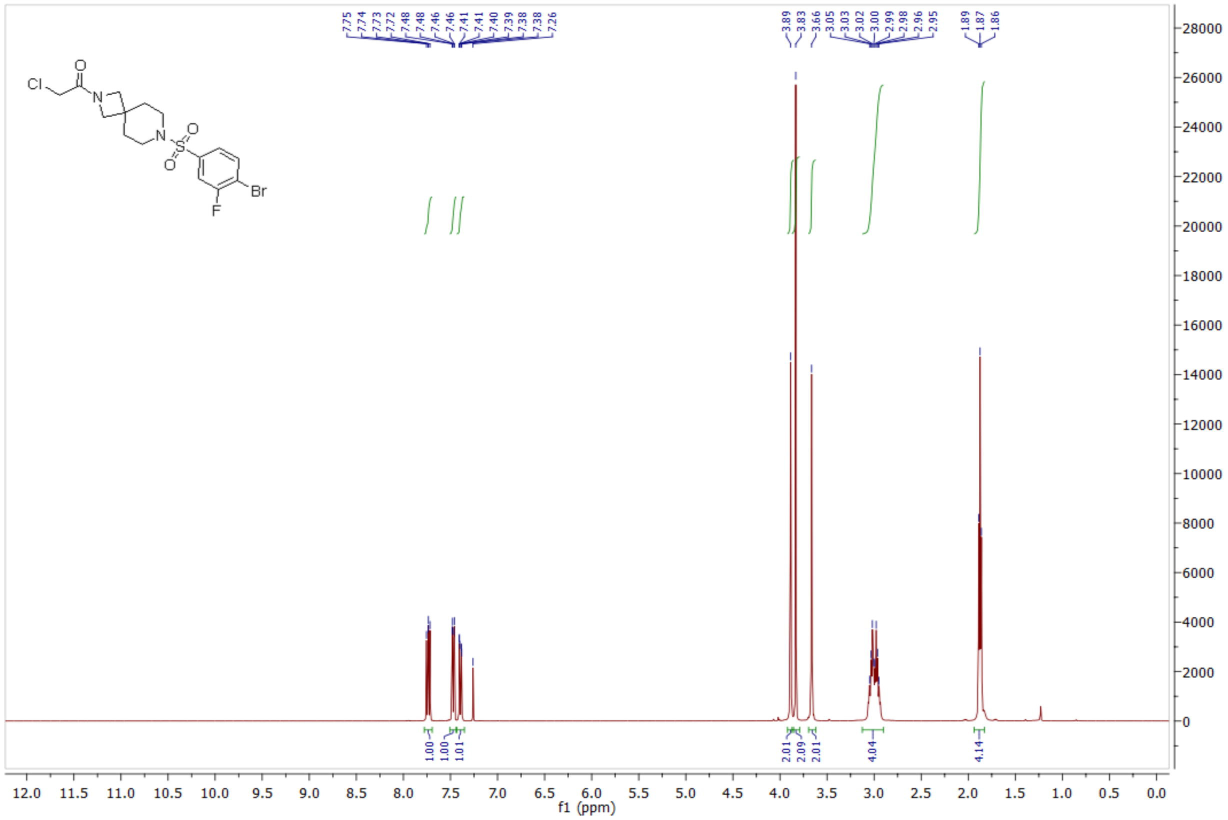

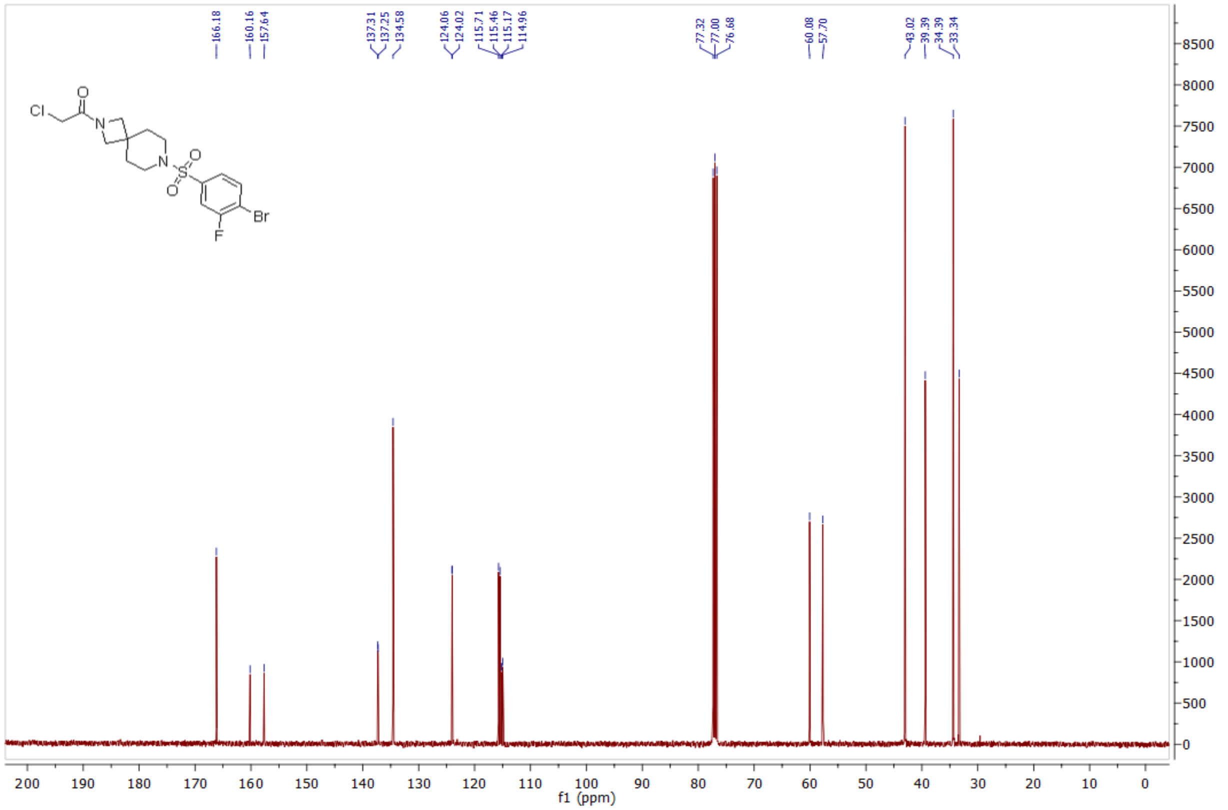

*N-(7-((4-bromo-3-fluorophenyl)sulfonyl)-7-azaspiro[3.5]nonan-2-yl)-2-chloroacetamide* **(55) 1084347**

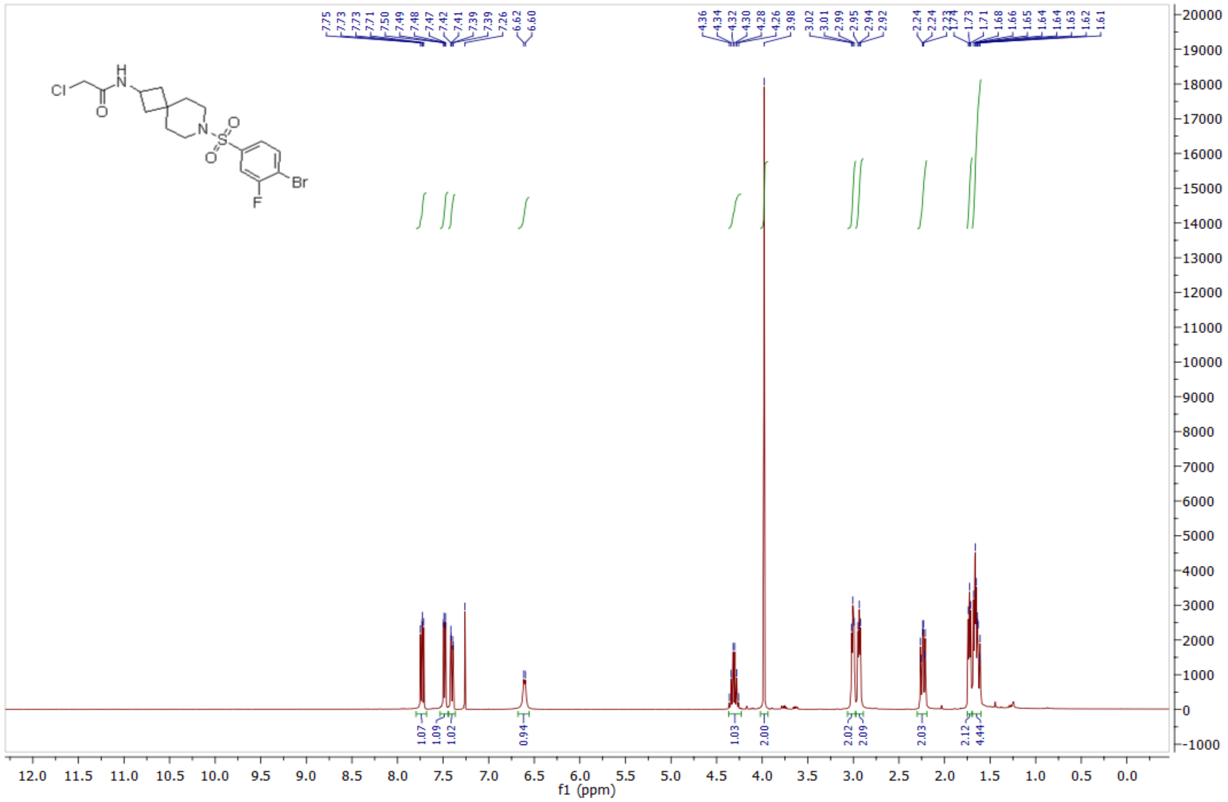

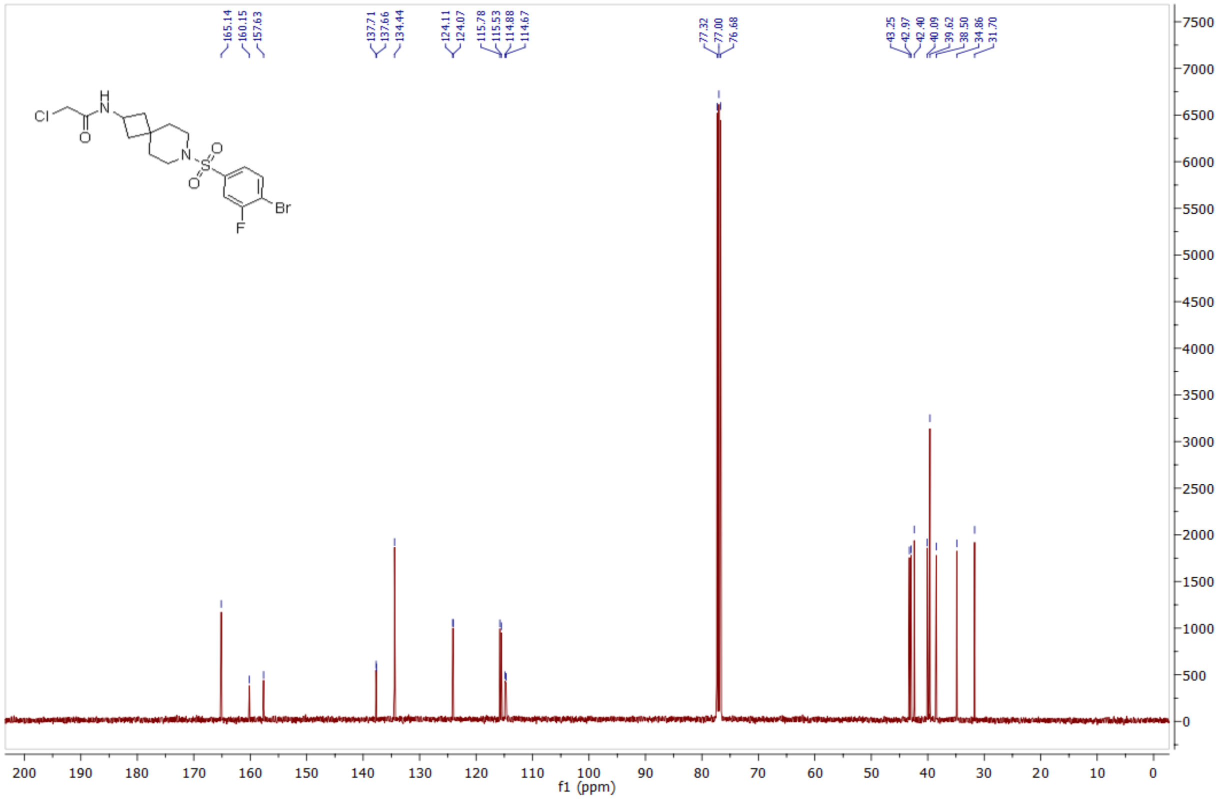

*1-(5-((4-bromo-3-fluorophenyl)sulfonyl)-2,5-diazabicyclo[2.2.1]heptan-2-yl)-2-chloroethan-1-one* **(58) 1084348**

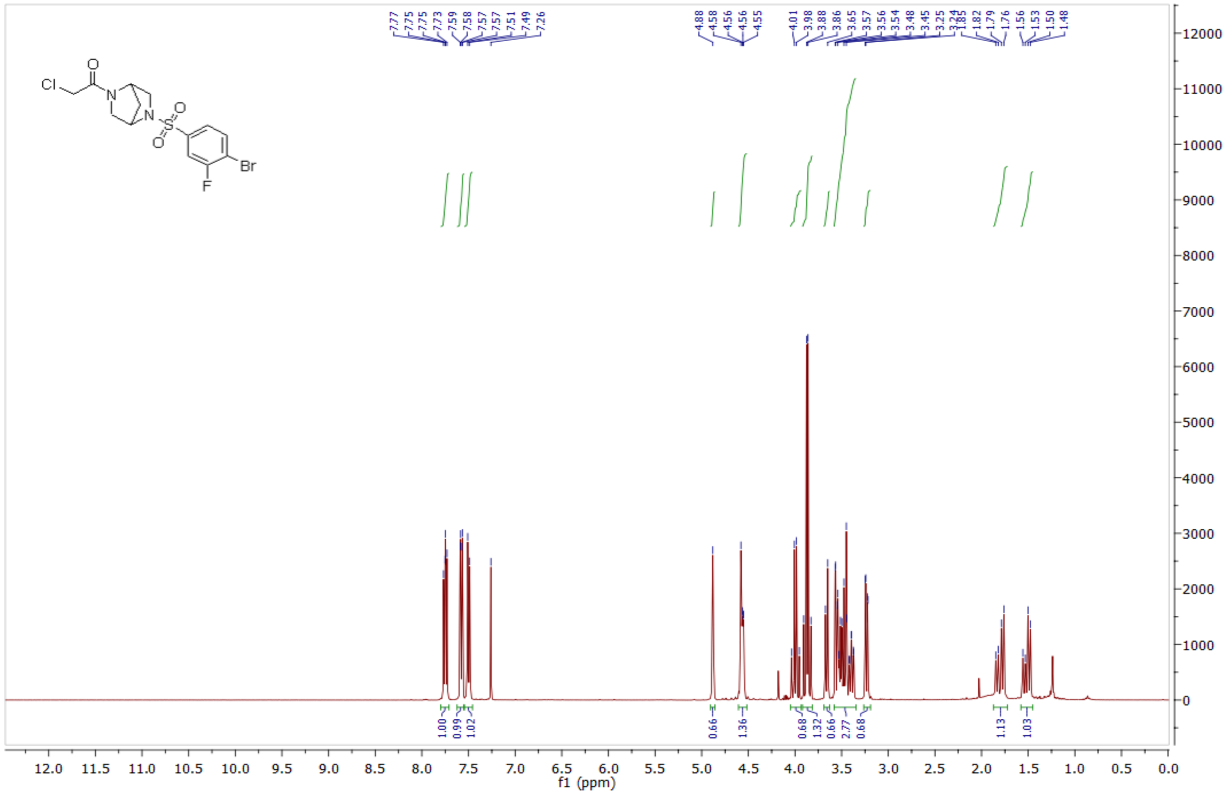

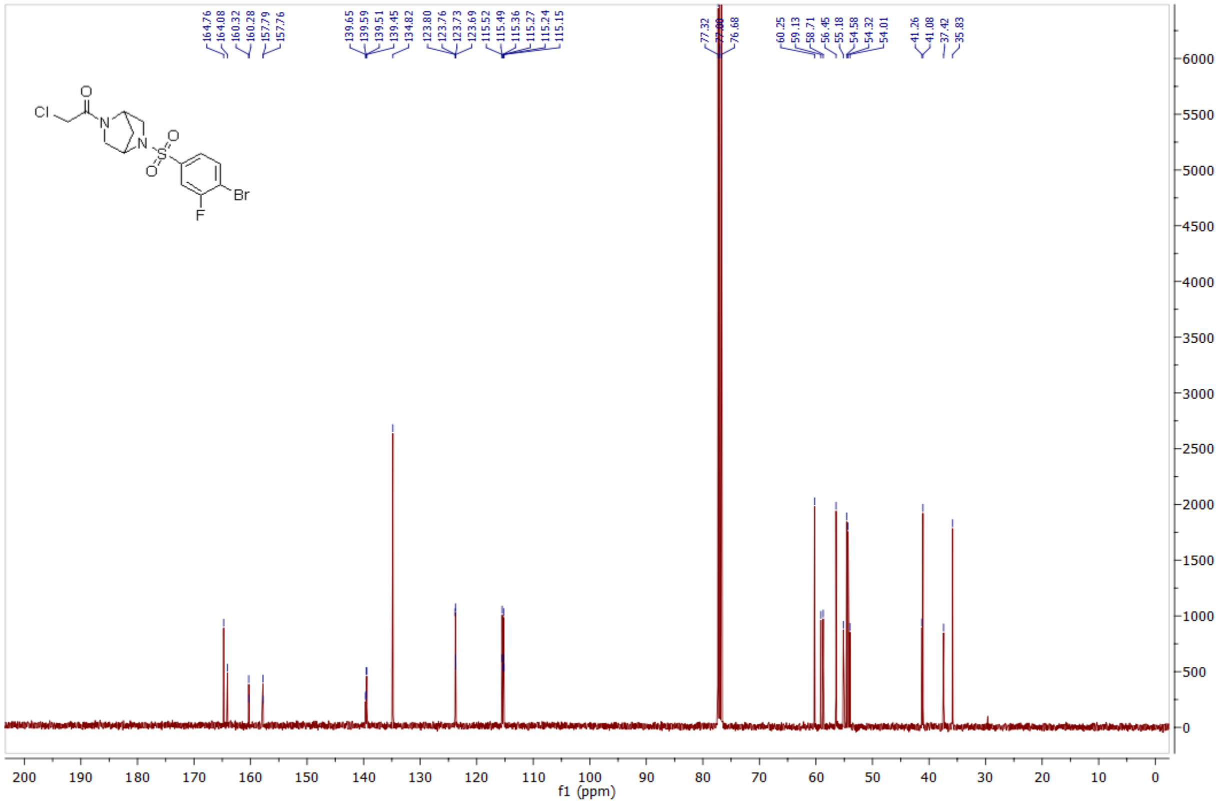

*4-bromo-N-(2-(2-chloroacetyl)-5-oxa-2-azaspiro[3.4]octan-7-yl)-3-fluorobenzenesulfonamide* **(67) 1084351**

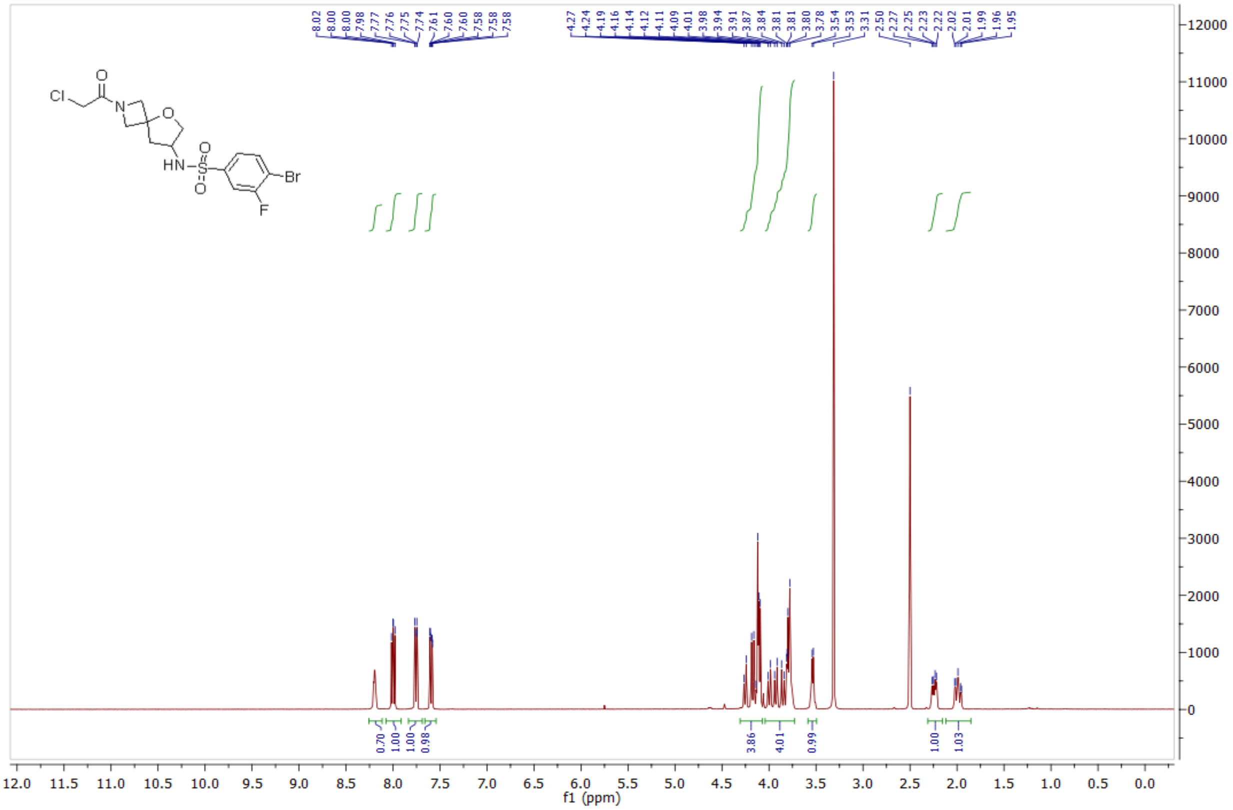

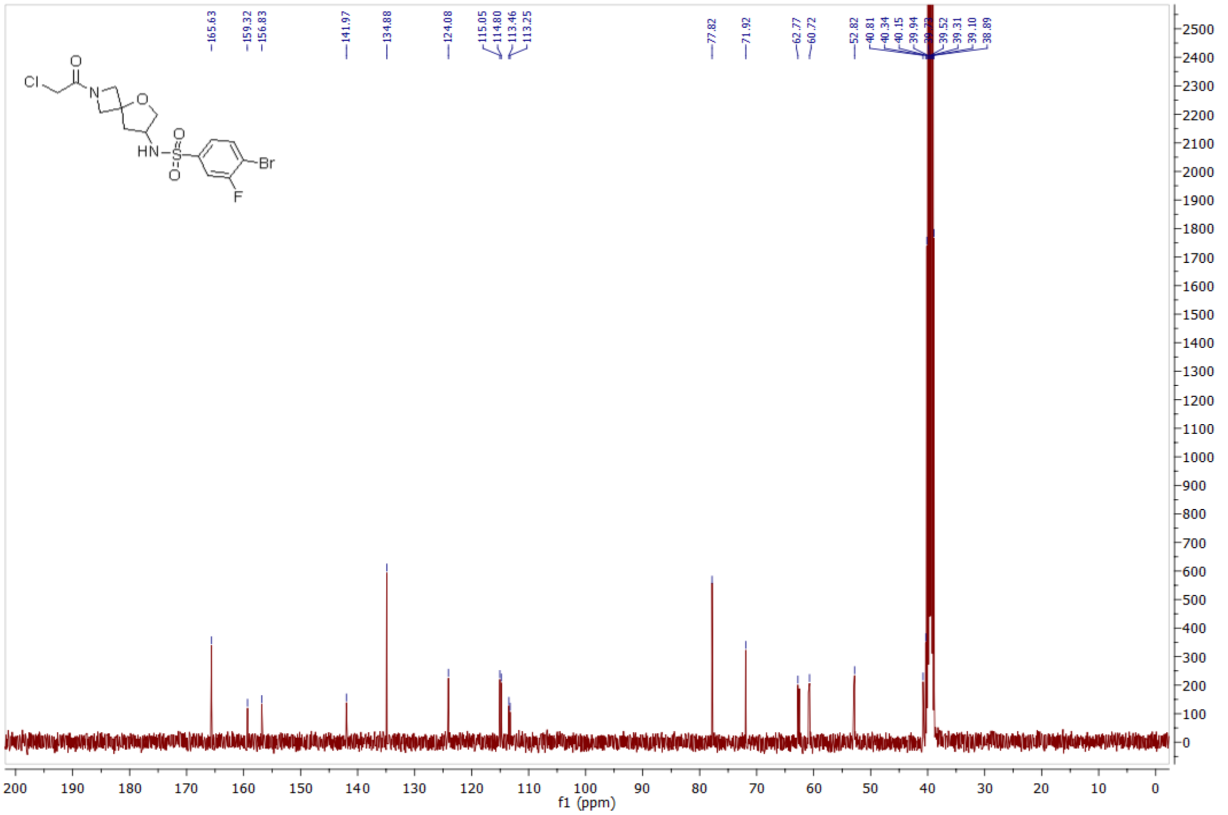

*1-(8-((4-bromo-3-fluorophenyl)sulfonyl)-5-oxa-2,8-diazaspiro[3.5]nonan-2-yl)-2-chloroethan-1-one* **(70) 1084352**

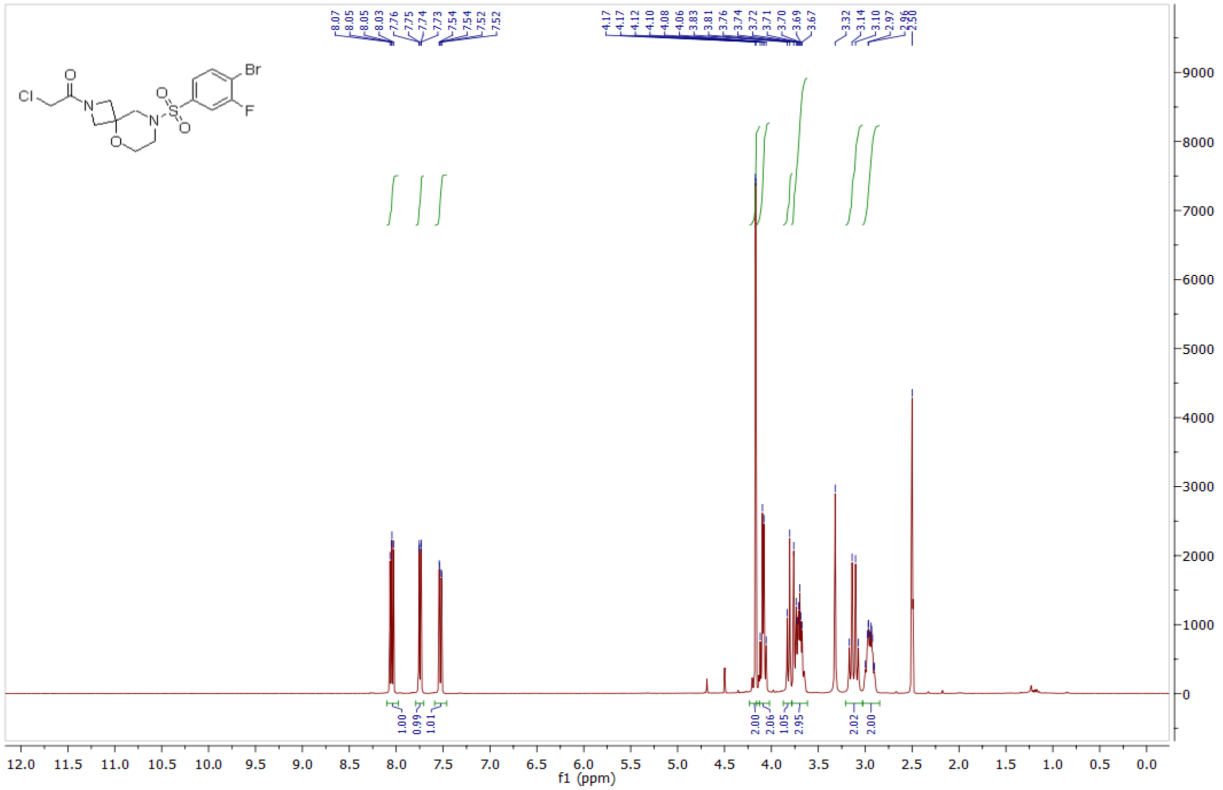

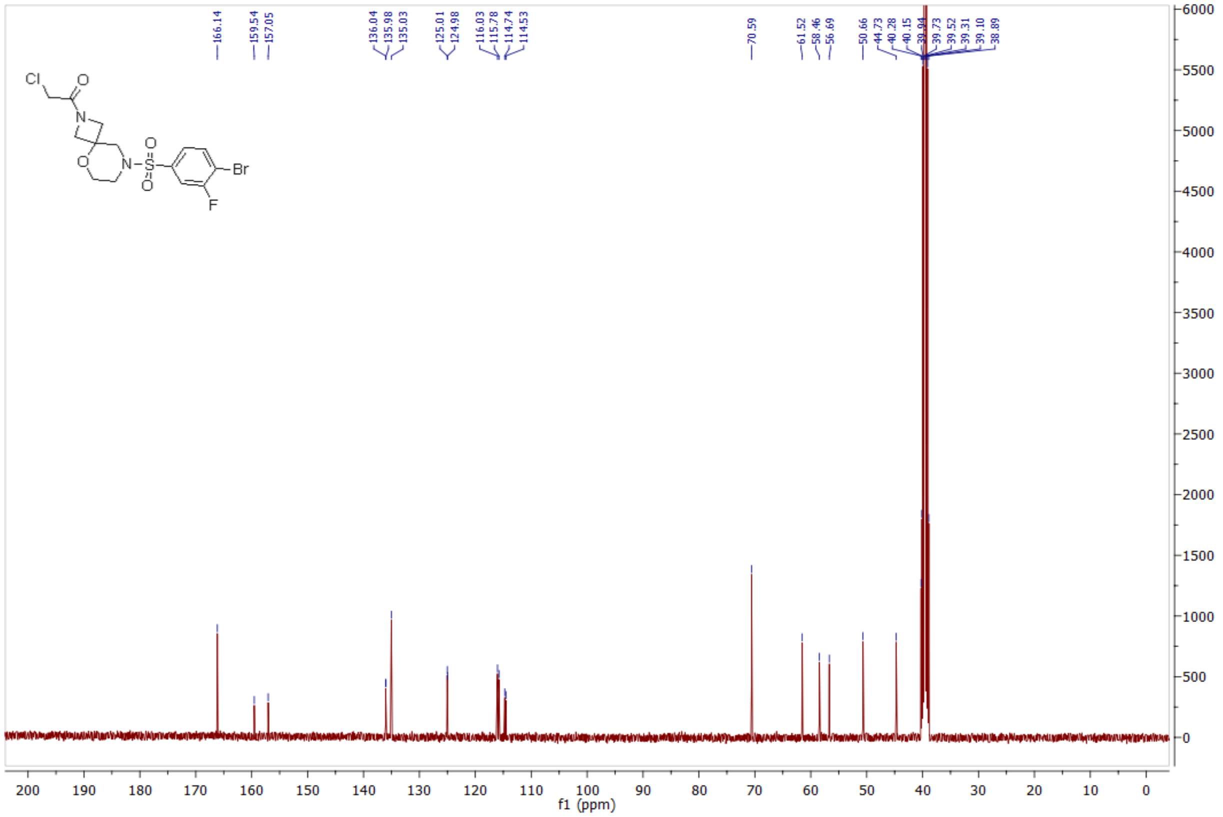

*1-(8-((4-bromophenyl)sulfonyl)-5-oxa-2,8-diazaspiro[3.5]nonan-2-yl)-2-chloroethan-1-one* **(78) 1124378**

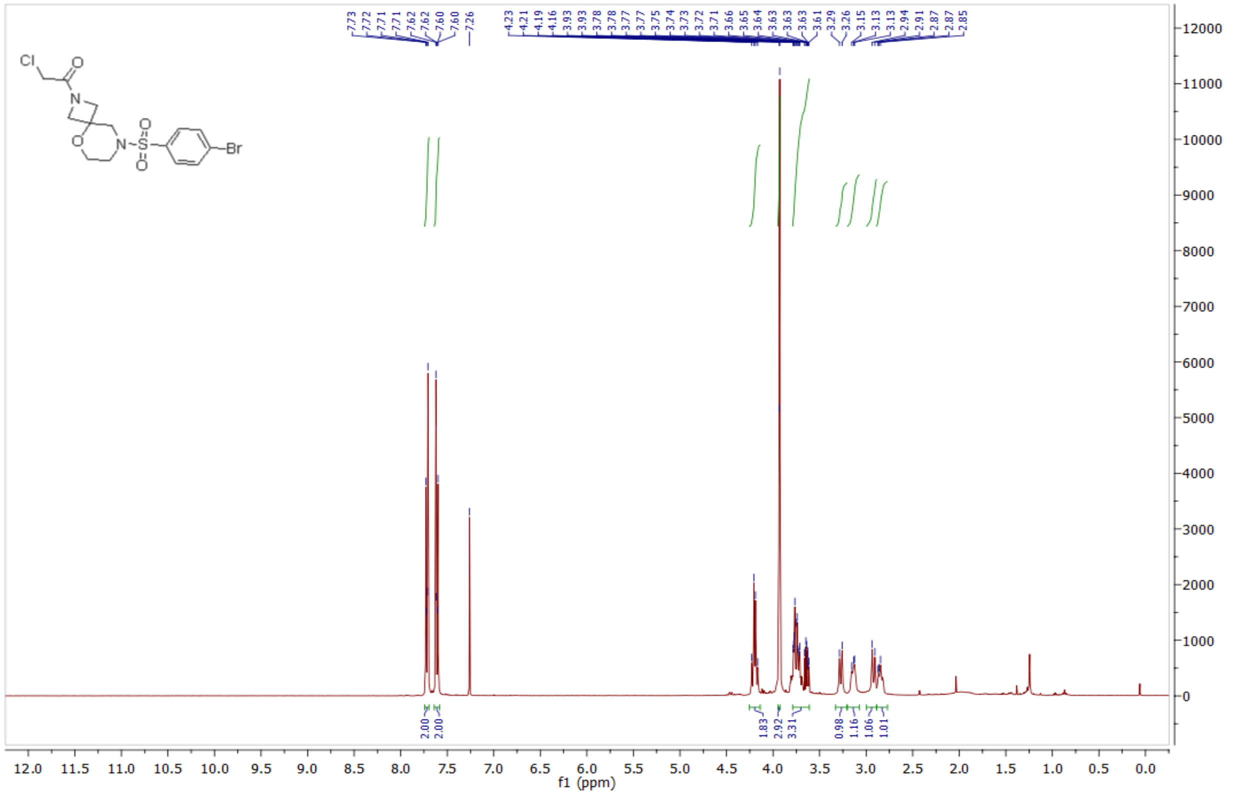

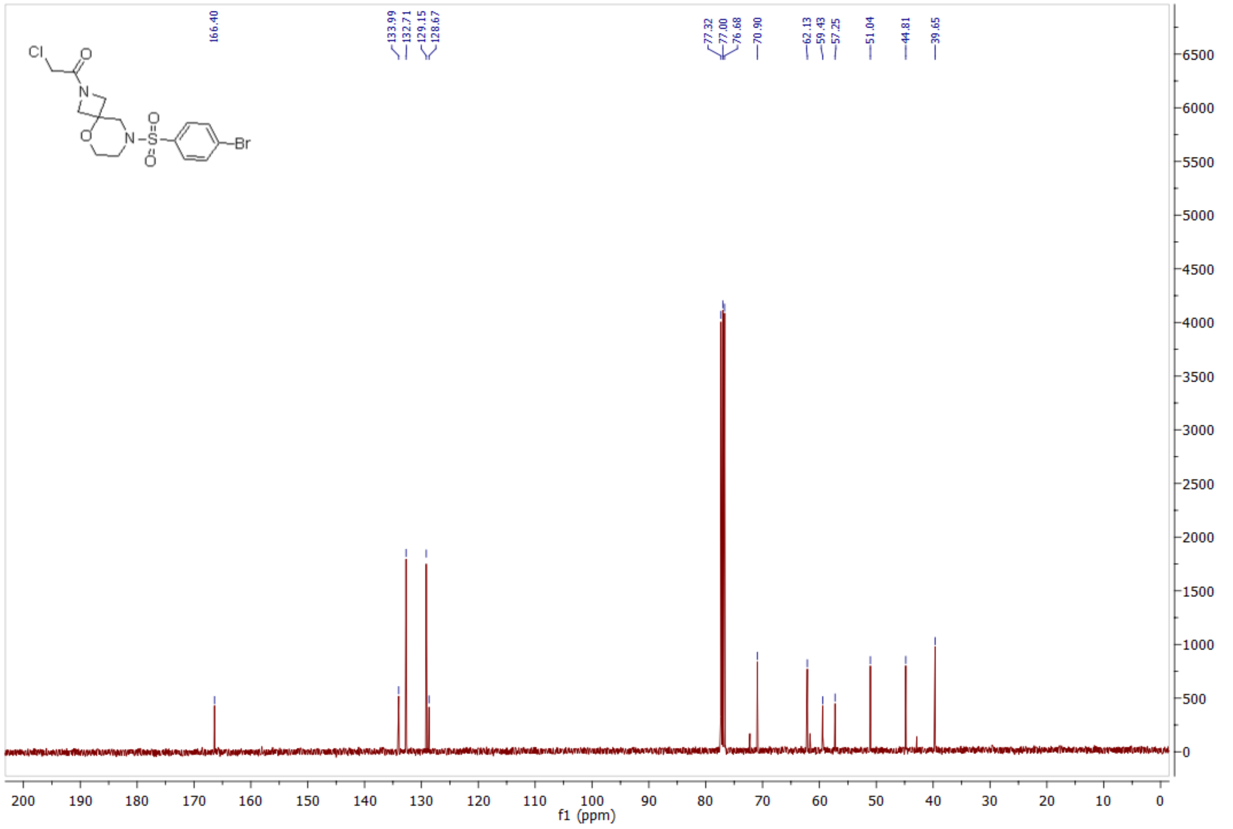

*1-(8-((4-iodophenyl)sulfonyl)-5-oxa-2,8-diazaspiro[3.5]nonan-2-yl)-2-chloroethan-1-one* **(79) 1124379**

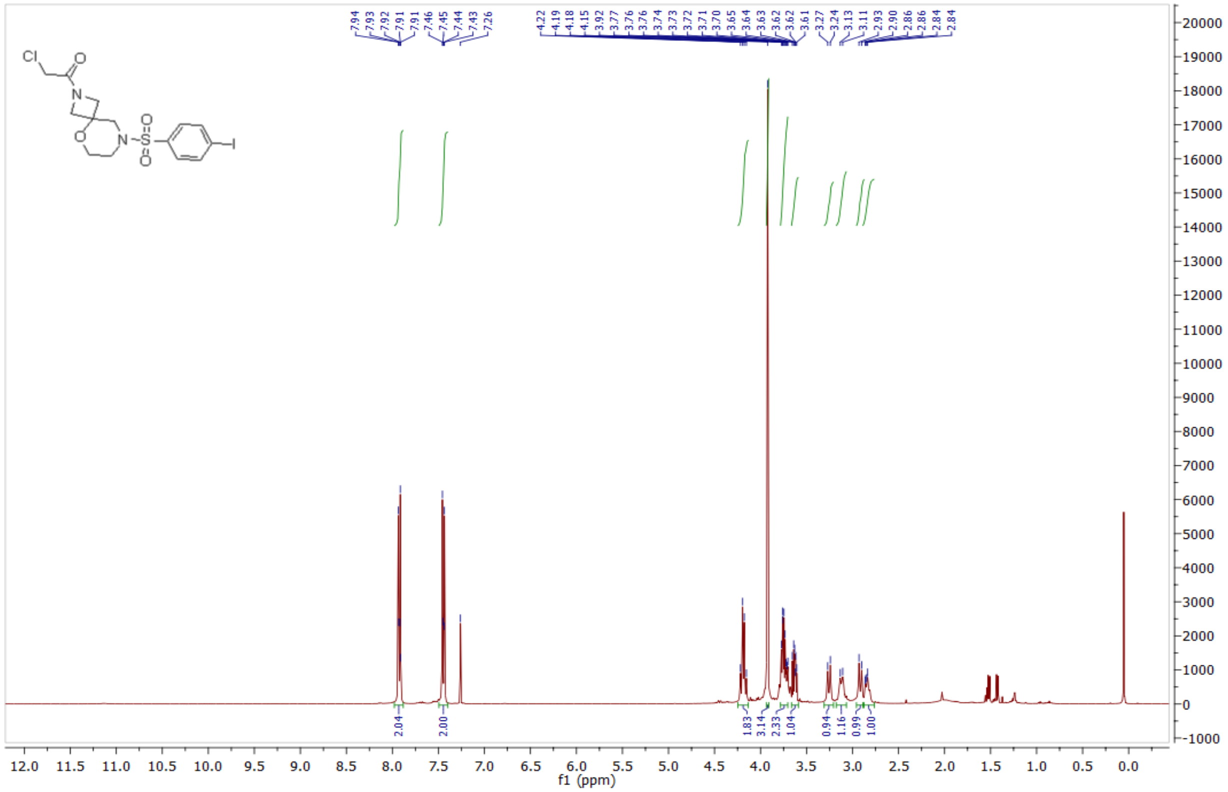

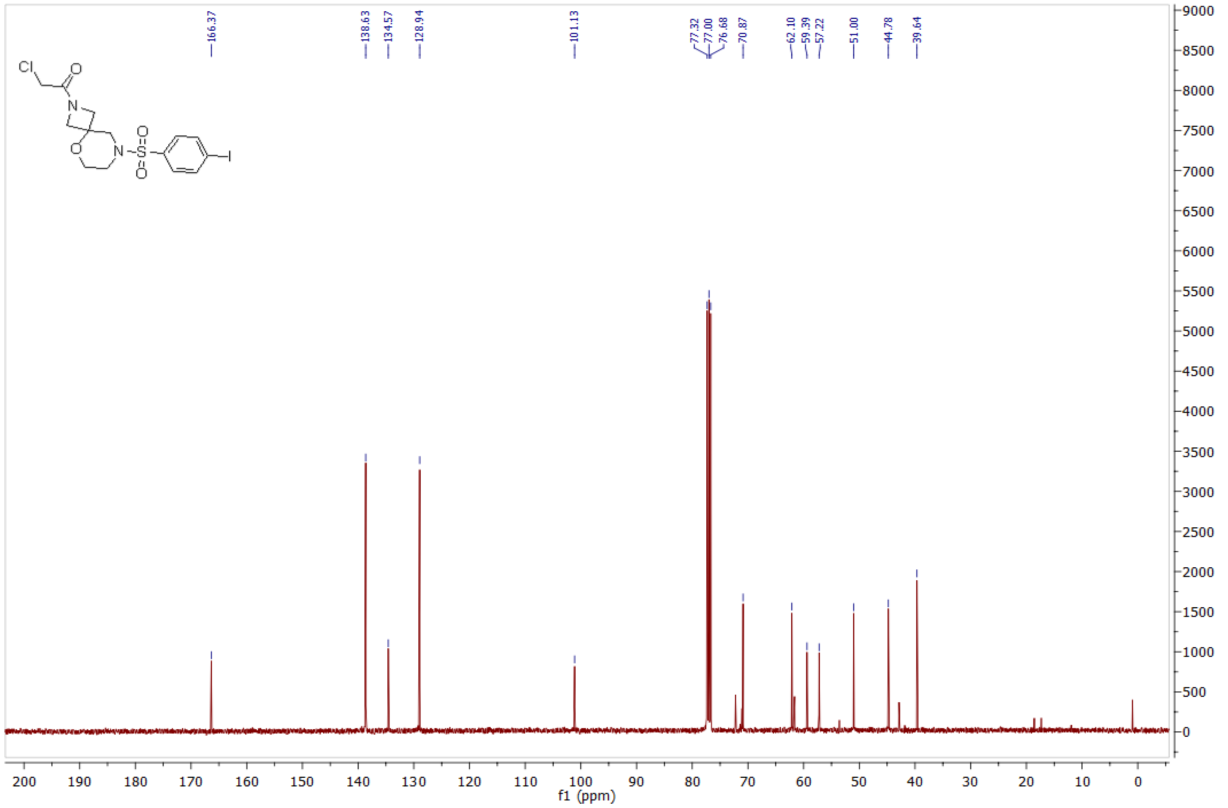

*N-((1-((4-bromo-3-fluorophenyl)sulfonyl)-4-methylpiperidin-4-yl)methyl)-2-chloroacetamide* **(83) 1124383**

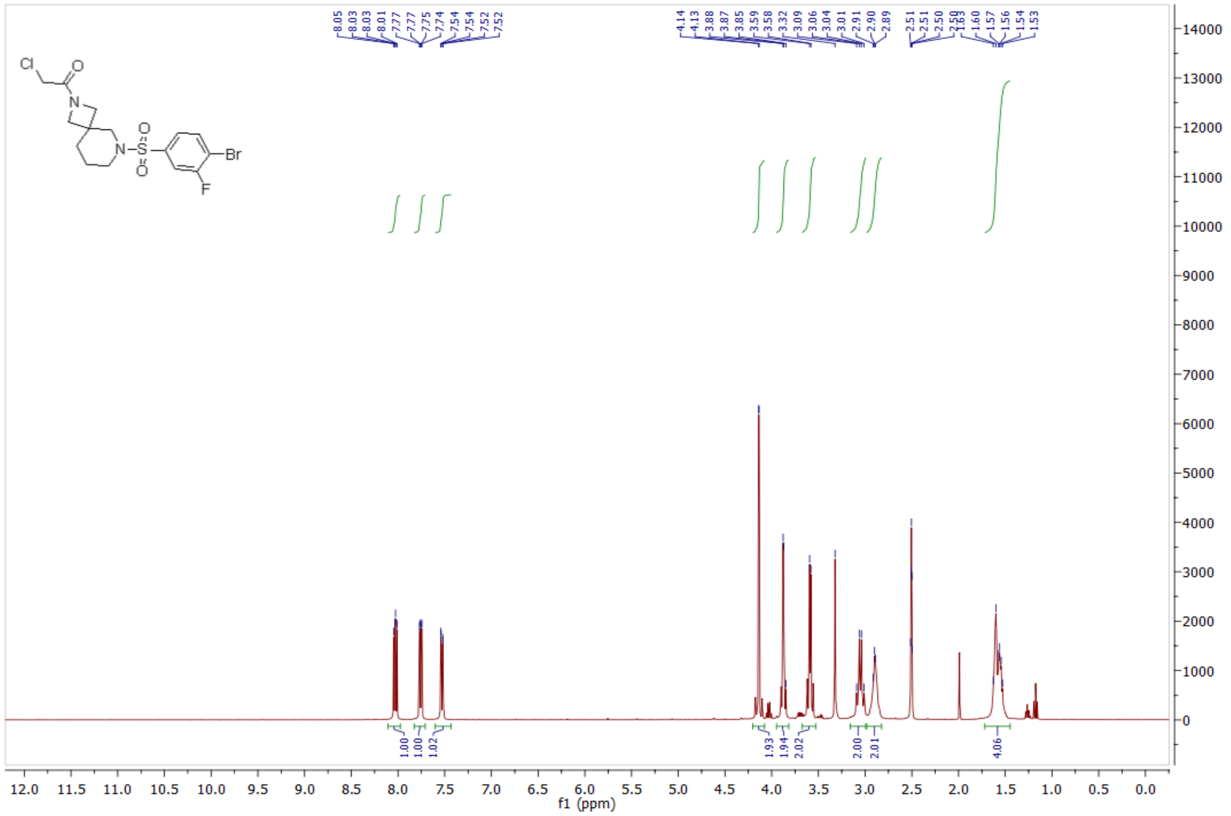

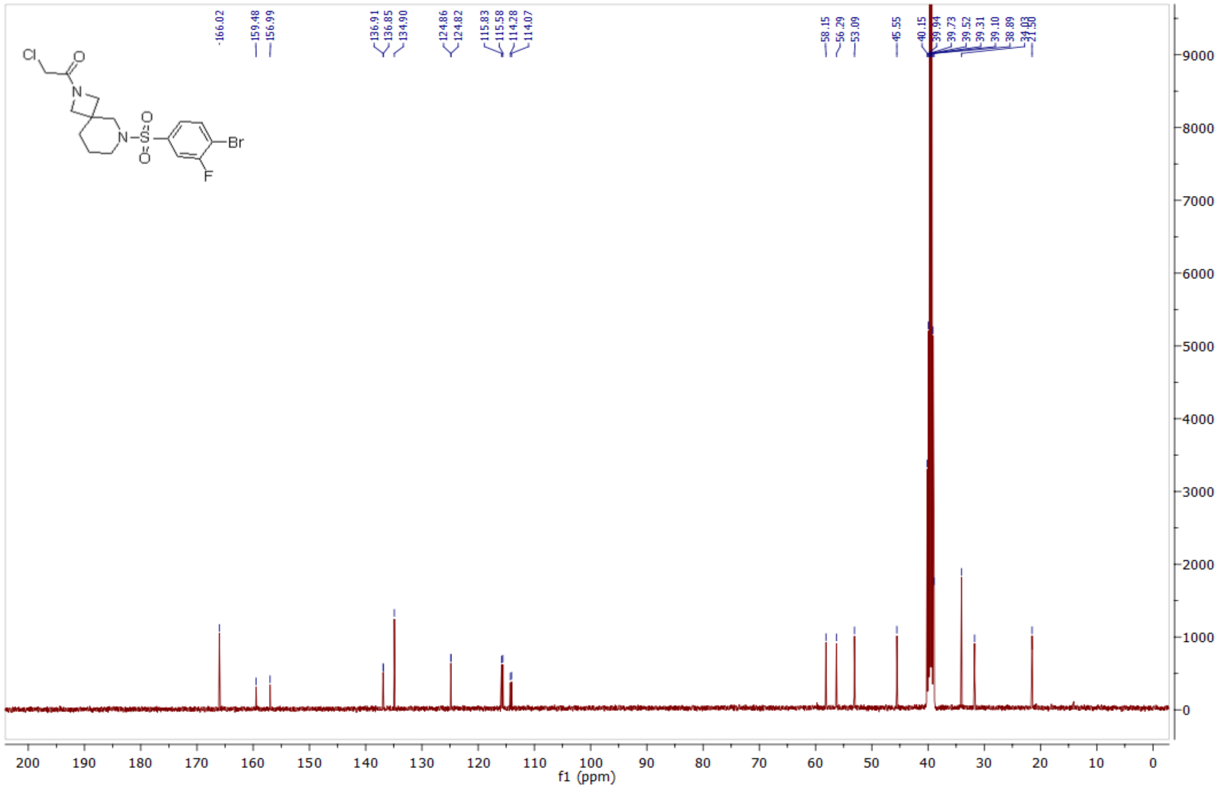

*(R)-1-(7-((4-bromo-3-fluorophenyl)sulfonyl)-2,7-diazaspiro[4.4]nonan-2-yl)-2-chloroethan-1-one* **(86) 1124384**

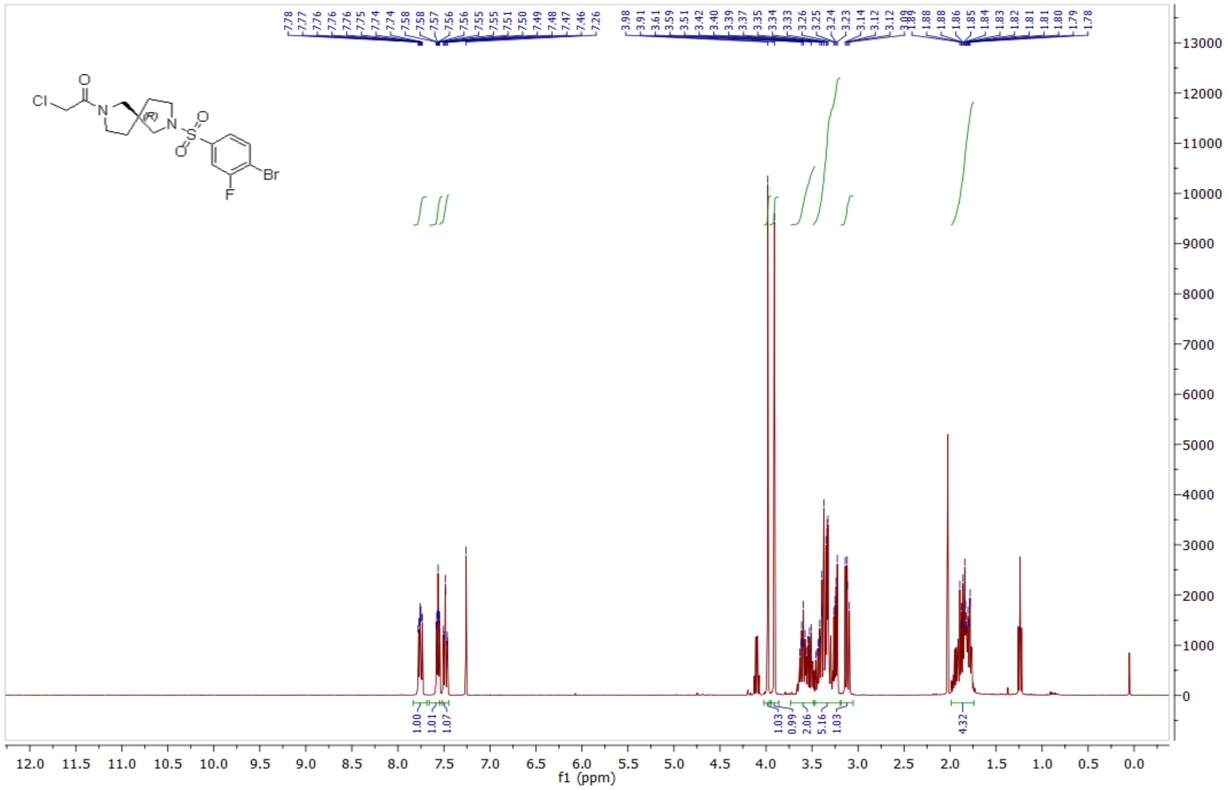

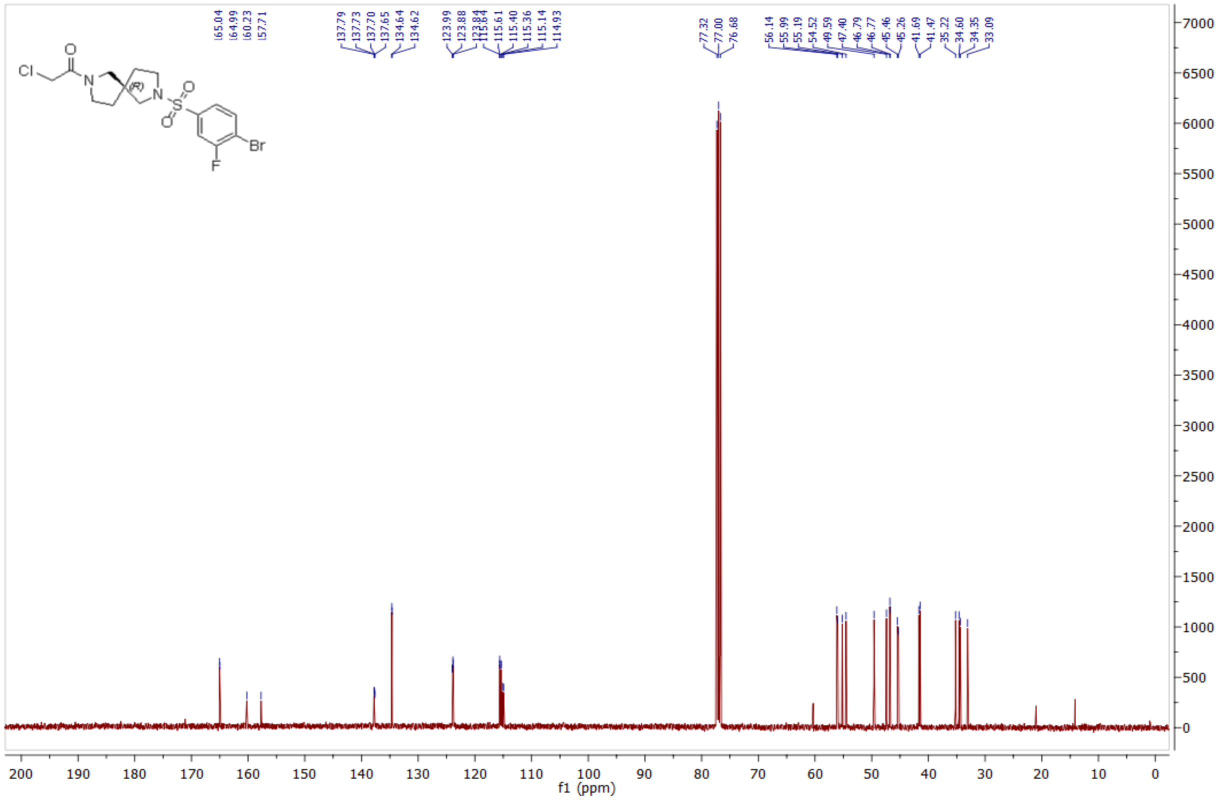

*(S)-1-(7-((4-bromo-3-fluorophenyl)sulfonyl)-2,7-diazaspiro[4.4]nonan-2-yl)-2-chloroethan-1-one* **(89) 1124385**

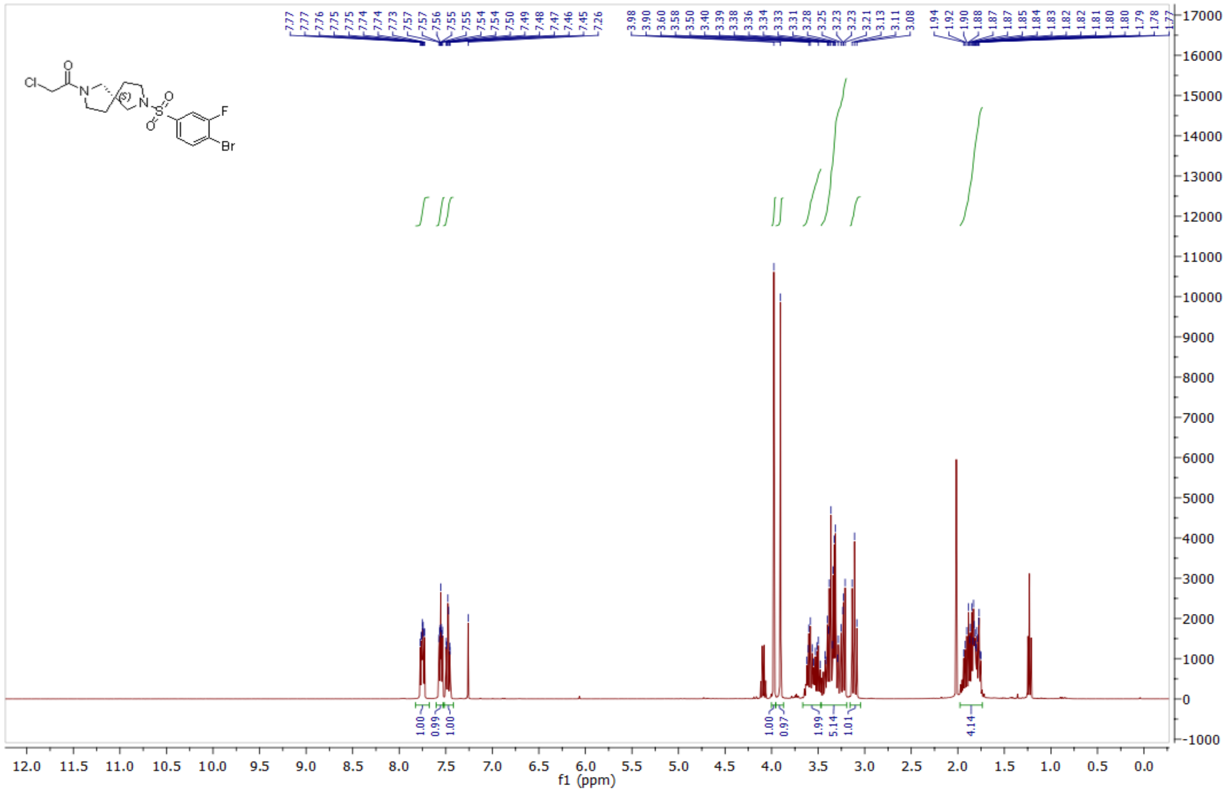

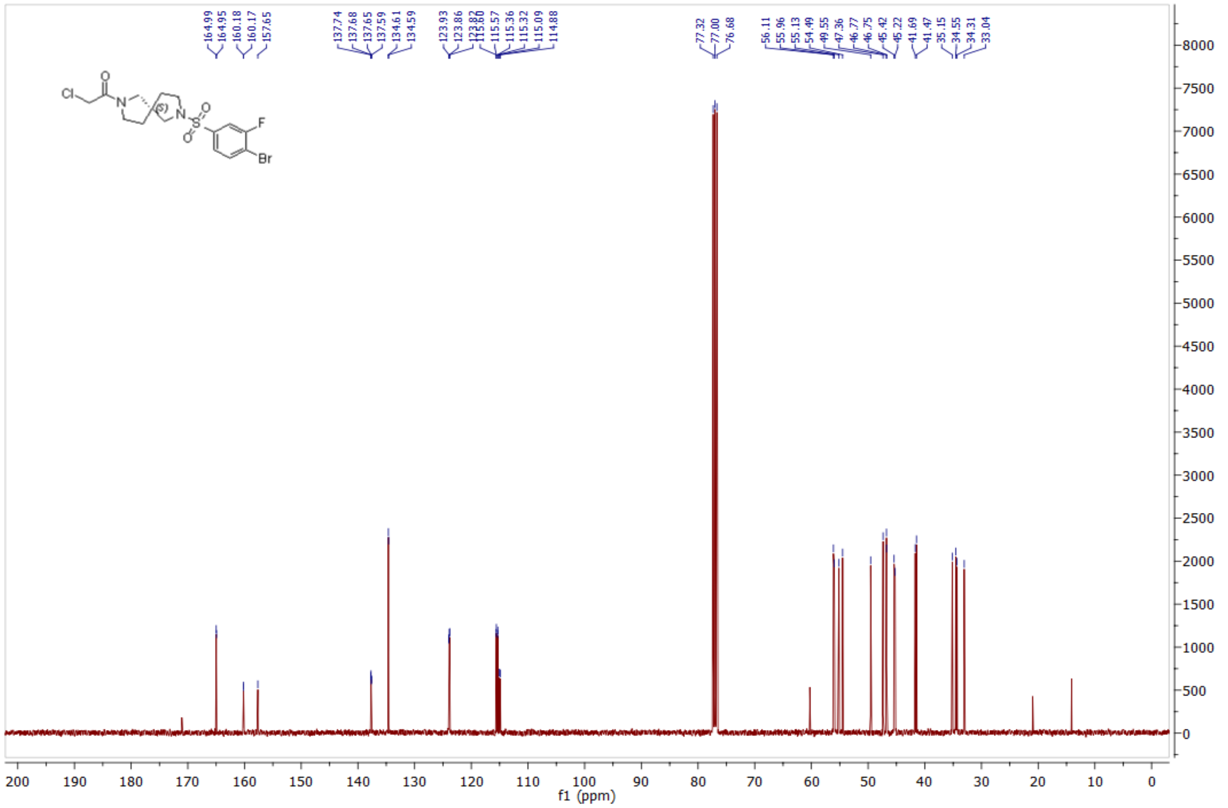

### 6. REFERENCES

(1) Hallenbeck, K. K.; Davies, J. L.; Merron, C.; Ogden, P.; Sijbesma, E.; Ottmann, C.; Renslo, A. R.; Wilson, C.; Arkin, M. R. A Liquid Chromatography/Mass Spectrometry Method for Screening Disulfide Tethering Fragments. *SLAS Discovery* **2018**, *23* (2), 183–192. https://doi.org/10.1177/2472555217732072.

(2) Vickery, H. R.; Virta, J. M.; Konstantinidou, M.; Arkin, M. R. Development of a NanoBRET Assay for Evaluation of 14-3-3σ Molecular Glues. *SLAS Discovery* **2024**, 100165. https://doi.org/10.1016/j.slasd.2024.100165.
